# Establishment of terminal selector combinations in optic lobe neurons

**DOI:** 10.1101/2024.02.05.578975

**Authors:** Félix Simon, Isabel Holguera, Yen-Chung Chen, Jennifer Malin, Priscilla Valentino, Claire Njoo-Deplante, Rana Naja El-Danaf, Katarina Kapuralin, Ted Erclik, Nikolaos Konstantinides, Mehmet Neset Özel, Claude Desplan

## Abstract

The medulla is the part of the *Drosophila* optic lobe with the greatest neuronal diversity, in which the identity of each neuronal type is specified in progenitors and newborn neurons via the integration of temporal, spatial, and Notch-driven patterning mechanisms. This identity is maintained in differentiating and adult neurons by the expression of neuronal type-specific combinations of terminal selectors, which are transcription factors expressed continuously during development and in the adult that are thought to control all neuronal type-specific gene expression. However, how the patterning mechanisms establish terminal selector expression is unknown. We have previously characterized the temporal and Notch origin of medulla neurons. Here we have used single-cell mRNA-sequencing to characterize their spatial origins and identified two new spatial subdomains. Together, this makes the medulla the first complex brain structure for which the patterning mechanisms specifying the identity of each neuronal type are known. This knowledge allowed us to identify correlations between patterning information, terminal selector expression and neuronal features. Our results suggest that different subsets of the patterning information accessible to a given neuronal type control the expression of each of its terminal selectors and of modules of terminal features, including neurotransmitter identity. Therefore, the evolution of new neuronal types could rely on the acquisition of modules of neuronal features pre-determined by their developmental origin.

## Introduction

Brain functions rely on communication between specialized neuronal types that are each characterized by a specific set of features, both molecular (neurotransmitters, ion channels, etc.) and morphological (brain area targeted, synaptic partners, etc.). Understanding how each of these features is established is key to understanding brain development. The stereotyped development and repetitive structure of the *Drosophila* optic lobe has enabled the comprehensive characterization of its neuronal morphological diversity^1–4^ and of its synaptic connectivity^5–10^. Recently, we and others have also characterized its transcriptomic diversity^11–15^. To do so, we performed single-cell mRNA sequencing (scRNA-seq) at six different stages of optic lobe development^11^, and identified ∼170 neuronal and ∼30 glial clusters. Almost ninety of these neuronal clusters and all glial clusters have since been matched one-to-one to known morphological cell types^11,16–21^. Lastly, the factors responsible for the specification of optic lobe neuronal types have been characterized in great detail^18,20,22,23^. In this work, we use these resources as a basis to explore how the information from specification factors in progenitors and in newly produced neurons is relayed to downstream targets during development to establish and maintain the features of optic lobe neurons.

The *Drosophila* optic lobe is constituted of four neuropils (Fig.1A): the lamina, medulla, lobula and lobula plate. Each is subdivided into ∼800 columns that correspond to the ∼800 ommatidia (*i.e.*, unit eyes) of the retina, and in several layers orthogonal to these columns except for the lamina that has no layers. Optic lobe neuron arbors can be restricted to a unique column (narrow-field), in which case the neurons are often present in ∼800 copies (synperiodic, *i.e.,* one per column), or they can span multiple columns (wide-field), in which case the neurons are often present in smaller and variable numbers (infraperiodic), although any combination of these characteristics can be found across cell types.

**Figure 1:**
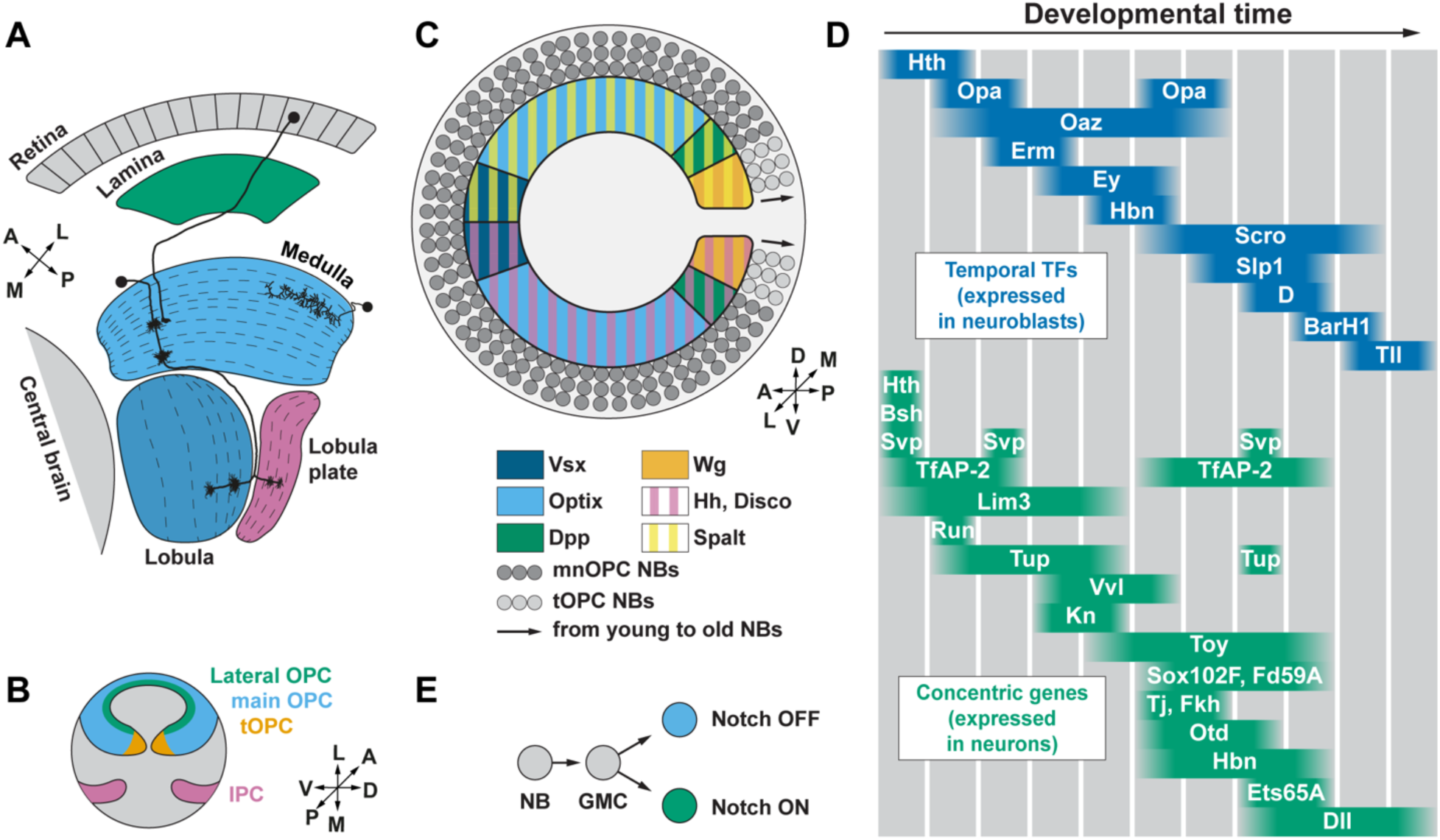
Specification of neuronal diversity in the *Drosophila* optic lobe. A) Cross-section of the adult visual system, with neuropil labels located adjacent to each one. Two narrow-field neurons (left) including one photoreceptor, and one wide-field neuron (right), are represented. Dashed lines: boundaries between neuropil layers. B, C) A: anterior, D: dorsal, L: lateral, M: medial, P: posterior, V: ventral. B) Schematic of the third-instar larval optic lobe, containing the lateral OPC (green), main OPC (blue), tips of the OPC (tOPC, orange) and the inner proliferation center (IPC, purple). C) Lateral view of the main OPC, showing the domain of expression of the spatial specification factors: Vsx (dark blue, anterior), Optix (light blue, middle), Dpp (green, posterior). The tips of the OPC (tOPC) express Wg (orange). Disco and Hh (pink hatched line) define the ventral domain while Spalt (yellow hatched lines) defines the dorsal domain. NB: neuroblast. D) In the main OPC, a series of temporal TFs (blue) defines temporal windows (grey columns) in neuroblasts, leading to the expression of “concentric genes” (green) in neurons (adapted from Konstantinides et al.^28^). E) Diagram of Notch signaling in optic lobe neurons. GMC: ganglion mother cell.

Optic lobe neurons are produced from two crescent-shaped neuroepithelia at the third-instar larval stage (L3, Fig.1B), the inner proliferation center and the outer proliferation center (OPC). The OPC can be further subdivided into the lateral OPC that produces lamina neurons, the tips of the OPC that express *wingless* (*wg*), and the main OPC that produces medulla neurons (Fig.1B). In this work we focus on the main OPC, which generates the greatest neuronal diversity and for which neuronal specification is best understood. At the beginning of the L3 stage, a neurogenic wave starts converting the OPC neuroepithelial cells into neural stem cells called neuroblasts in *Drosophila.* Neuroblasts divide multiple times to self-renew and produce an intermediate progenitor called a ganglion mother cell, which in turn divides once more to generate two neurons. Neuronal identity is specified by the integration of three mechanisms^24–29^. First (Fig.1C), the main OPC neuroepithelium is subdivided into different spatial domains that express either the spatial transcription factors (TFs) Visual system homeobox 1 (Vsx1) or Optix, or the signaling molecule Decapentaplegic (Dpp)^25^. Additionally, the dorsal part of the main OPC neuroepithelium expresses Spalt TFs (Salm and Salr), while the ventral side expresses Disco TFs (Disco and Disco-r) and, only early in development, Hedgehog (Hh)^24,30^. We will hereafter refer to ventral and dorsal Vsx, Optix, and Dpp domains as v/dVsx, v/dOptix and v/dDpp, respectively. Second (Fig.1D), all main OPC neuroblasts sequentially express, as they age, a cascade of temporal transcription factors (tTFs) whose expression partially overlaps^25–29^. Although tTFs often do not maintain their expression in neurons, several downstream TFs termed “concentric genes” (Fig.1D) are expressed in neuronal types emerging from one or several juxtaposed temporal windows, and are therefore expressed in a concentric fashion in the main OPC^28,31^ (reflecting the spatial organization of neuroblasts of identical age in the main OPC, Fig.1D). Third (Fig.1E), the Notch pathway is transiently active in one of the two neurons produced by each ganglion mother cell (Notch^ON^ neurons) but inactive in the other (Notch^OFF^ neurons)^11,26^. We hereafter use “specification factor” to refer to any factor (often transiently expressed) involved in spatial, temporal or Notch patterning, and “developmental origin” for any combination of spatial, temporal and Notch origin.

The expression of spatial and temporal specification factors is mostly not maintained in neurons, and the direct effectors of Notch signaling are present only in newborn neurons^11,26^. To understand how neuronal features are established and maintained, one must therefore understand how the information encoded by the specification factors is propagated to differentiating neurons. This could be achieved by several mechanisms: a) epigenetic mechanisms established in progenitors and maintained in neurons (*e.g.*, chromatin accessibility, histone marks, etc.), b) TF cascades initiated by specification factors and unfolding in neurons, c) continuous expression in neurons of TFs downstream of the specification factors, or d) a combination of these. It has been proposed that continuous TF expression in neurons is a prevalent mechanism for neuronal differentiation and is achieved by terminal selectors (TSs), *i.e.,* neuronal type-specific combinations of TFs that are maintained during development and in adults and control all specific features of a neuronal type by binding the cis-regulatory regions of either terminal effector genes or other TFs controlling terminal effector genes^32,33^. We previously identified candidate TSs (simplified hereafter as “TSs”) in the optic lobe: combinations of TFs, with an average of 10 per neuron, that are unique to each neuronal type and are maintained throughout neuronal life^11,17^. We were able to convert distinct neuronal types that shared very similar TS combinations into one another by rendering these combinations identical^17^.

Although we recently described the temporal origin and the Notch status of all the main OPC clusters in our scRNA-seq atlas^28^, the lack of knowledge of their spatial origin has prevented us from exploring how the specification factors establish together the TS combinations specific to each neuronal type. In this study, we isolated all the neurons produced from different main OPC spatial domains and performed scRNA-seq to identify them and thus comprehensively characterize the spatial origin of main OPC neurons. This complements our earlier characterization of their temporal origin and Notch status. It makes the medulla the first complex brain structure for which the specification mechanisms patterning each neuronal type are known, and can be linked to gene expression during development and adulthood using our scRNA-seq atlas of optic lobe development^11^. Moreover, these data allowed us to describe two new spatial domains. We then used the characterization of the developmental origin of each neuronal type to identify how each temporal window, spatial domain, Notch status, or any combination of these three specification mechanisms, correlates with the expression of TSs (as well as other genes specifically expressed in neuronal types and morphological features) in differentiating and adult neuronal types. Lastly, we trained random forest models that show that the developmental origin of optic lobe neurons is highly predictive of their TS expression, and that developmental origin or TS expression are equally informative in predicting gene expression or morphological features of optic lobe neurons.

## Results

### Profiling the neuronal types produced in each spatial domain

Identifying the spatial origin of neuronal types has traditionally relied on immunostainings against marker TFs, or on the combined use of lineage tracing and reporter lines^20,25,34^. However, such approaches are low-throughput and are not always possible due to the lack of appropriate marker genes, antibodies, or driver lines. We therefore designed a high-throughput approach to identify the spatial origins of all main OPC neurons (Fig.2A). We drove the expression of nuclear GFP in cells produced from various main OPC spatial domains using domain-specific *Gal4* drivers, isolated the labeled cells by Fluorescence-Activated Cell Sorting (FACS), performed scRNA-seq on the sorted cells, and annotated the cells from each population using a neural network classifier trained on our previously published optic lobe scRNA-seq atlas^11^. Whenever possible, we favored sequencing adult neurons by using memory cassettes (MC) (Fig.2B, Fig.S1A) that label all progeny of cells that expressed a given *Gal4* driver earlier in development. Indeed, at early pupal stages some neurons are still too immature to be identified precisely by our classifier^11^. Moreover, when possible, we added a temperature-sensitive (ts) *Gal80^t^*^s^ to prevent later activation of the memory cassettes in postmitotic neurons that may not originate from the domain of interest (Fig.2B, Fig.S1B).

**Figure 2:**
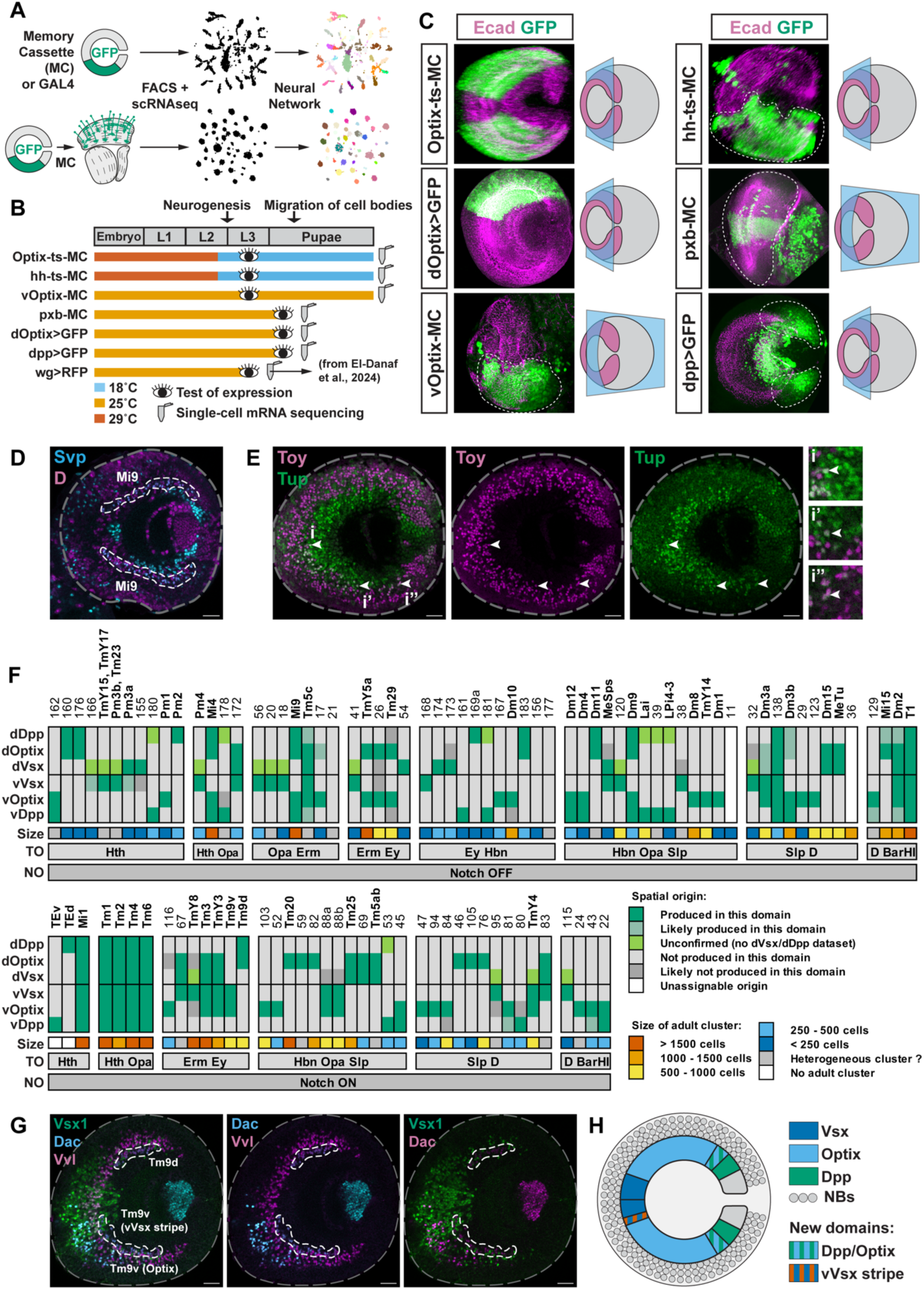
Identification of the spatial origins of main OPC neurons using scRNA-seq. A) Schematic of the approach used to identify the neurons produced in each main OPC domain. Gal4 lines driving the expression of a nuclear GFP were used to isolate cells from a given spatial origin with FACS, either in early pupae (top) when the Gal4 is expressed in neuronal progenitors, or in adults (bottom) using a memory cassette to label permanently the neurons produced in a given spatial domain (Fig.S1A). Sorted cells were then sequenced, and the single-cell transcriptomes were annotated using a neural network classifier^11^. B) Summary of the stages at which we tested the expression of, or sequenced, each of the lines used. The expression pattern of the lines used was tested before migration of cell bodies, either at the L3 stage or in early pupa. Temperature sensitive memory cassettes (ts-MC) are inactive at 18°C, but permanently label the cells expressing them at 29°C as well as their progeny. C) Maximum intensity projections showing that each of the lines sequenced labelled the expected main OPC populations (see Fig.1C), following the protocol summarized on panel B. Labelled neurons outside of the dashed lines are not part of the main OPC. Schematics show the orientation of the maximal intensity projection. D, E, G) Lateral view of the main OPC at the L3 stage. Grey dashed line: outline of optic lobe. Scale bars are 20 µm. D) Mi9 neurons (white dashed line) are Svp+ D+ and localized in the v/dOptix and v/dDpp domains. E) Toy+ Tup+ cells, which include Dm10 neurons, are indicated by arrows in the vVsx (i), vOptix (i’), and vDpp (i’’) domains. F) Developmental origin (spatial, temporal, and Notch) and abundance of main OPC neuronal types (Table S1). TO: temporal origin, NO: Notch origin, Size: number of cells comprised in the corresponding cluster in our adult scRNA-seq atlas^11^. G) Tm9 neurons (white dashed line) are the first row of Dac+ Vvl+ neurons. Tm9v but not Tm9d is produced from the Vsx domain, in the vVsx stripe. H) Newly discovered spatial domains of the main OPC, lateral view.

We tested the specificity of the lines by checking their GFP expression pattern before the rearrangement of neuronal somas^31^, *i.e.,* either at the L3 larval stage or between 0 to 12.5% of pupal development (P0-P12.5, Methods, Fig.2B, Fig.2C). All lines labelled the expected main OPC populations and sometimes some cells outside the main OPC, which we accounted for in later analyses (Methods). Additionally, some ectopic expression in the main OPC was observed in vOptix-MC (Fig.S1C, Fig.S1D), which we accounted for in later analyses (Methods), and in pxb-MC (Fig.S1E), which we eliminated by visually selecting optic lobes that did not exhibit ectopic expression before FACS and sequencing. We used *pxb-Gal4* rather than *Vsx-Gal4*^27^ because *Vsx* is also expressed later in neurons from the v/dOptix domains (Fig.S1C, Fig.S1F).

The FACSed GFP-positive cells from each spatial domain were then subjected to scRNA-seq at the indicated stages (Fig.2B). We used our classifier^11^ to annotate the transcriptome of each cell with a confidence score. Among all the “Classes” (defined hereafter as a group of cells with the same annotation) in each dataset, we then identified the ones that were annotated with a lower confidence (Annex 1, Methods). We did not discard these cells but flagged them for later bioinformatic analyses since low-confidence annotation scores are not always an anomaly and are, for instance, expected for subtypes with similar transcriptomes (Fig.S2A). We then normalized neuronal abundances to be able to compare them between different datasets (Fig.S2B, Methods).

Although the spatial origin was very clear for most neuronal types (Fig.S2C, Annex 1), we sometimes found a low number of cells belonging to a neuronal type from a known spatial origin in datasets where they were predicted to be absent. This can be explained by experimental (*e.g.* imperfect sorting), bioinformatic (*e.g.* imperfect filtering), or biological (*e.g.* cell types with very similar transcriptomes, which are difficult to distinguish using the neural network) variation.

### Assigning a spatial origin to each optic lobe neuronal type

To identify false positives in each dataset and to validate the scRNA-seq results, we used the known spatial origins of Dm8^34^, Mi1^26,31^, Pm1/2/3^25^, T1^35^ and TEv/d^11^ neurons, and experimentally validated the spatial origins of 32 additional clusters. To do so, we identified markers expressed early in these clusters (Fig.S3A) and assessed the location of their expression by immunostaining in the main OPC cortex at late L3 stage (before cell bodies rearrange) (Fig.2D-F, Fig.S3B-J, Table S1). For instance, Mi9 expresses *svp* and *D*, and the co-expression of these markers was found in v/dOptix and in v/dDpp but not in v/dVsx at the L3 stage (Fig.2D), while Dm10 expresses *toy* and *tup,* which are only co-expressed in the ventral side of the main OPC (Fig.2E). This is consistent with the lack of Mi9 cells in our pxb dataset (Fig.S2C) and the lack of Dm10 cells in the dOptix dataset (Annex 1), thus validating our scRNA-seq results for these neurons.

We then computed the enrichment of each Class in each dataset (Fig.S2B) by dividing the normalized abundance of each Class by the normalized abundance in our previously published scRNA-seq atlas, which represents a largely unbiased sampling of optic lobe neurons from all main OPC domains. We binarized these enrichment values (Methods) into “presence” or “absence” of a Class in a dataset by setting a threshold based on our experimental validations as well as the published origins of the cell types previously mentioned (Fig.S4, Fig.S5, Annex 1). After binarization, the spatial origins determined by scRNA-seq matched the ones determined experimentally for 39 neuronal types (Fig.S4) but did not entirely match for Dm11. This neuronal type is produced in the dOptix domain (see TableS1) but was found in the pxb dataset (*i.e.*, Vsx domain) in addition to the datasets containing cells produced in the dOptix domain. However, on UMAP visualization of the pxb dataset, Dm11 is poorly separated from the transcriptionally similar Dm8 and cluster 38 (Fig.S6, TableS1), which indicates that Dm11 cells could have been misannotated by the neural network. In addition, both scRNA-seq data and experimental validations indicate that cluster 163, which we annotated as Pm1^11^, is produced in the vOptix domain. However, Pm1 was thought to be produced from vDpp^25^. One possible explanation is that the cell type corresponding to cluster 180 was previously mistaken for Pm1 (see Table S1).

Since the spatial origins determined by scRNA-seq and the *in vivo* validations were highly consistent (they matched for 39 out of 40 clusters), we generated a table assigning a spatial origin to all other main OPC clusters by considering normalized abundances, proximity to enrichment binarization thresholds, UMAP representation, annotation confidence and experimental results from our validations and from previous publications (Methods). Our assignments are explained in Table S1, while Fig.2F represents a summarized view of this table.

The intersection of a single temporal window, a single spatial origin, and a single Notch status is supposed to encode a single neuronal identity (Fig.1). However, as can be seen in Fig.2F, in many cases such a combination encodes more than one neuronal type. For instance, Notch^OFF^ neurons from the Hth window and the Vsx domain can become at least TmY15, TmY17, Pm3a, Pm3b, or Tm23. This extreme case is explained by the recent discovery of an additional mechanism to encode neuronal diversity, in which neuroblasts from the same spatial, temporal and Notch origin but born at different times produce different neuronal types^18^. Other cases could also be explained by this fourth specification mechanism. However, several cases are likely due to the existence of additional spatial domains and temporal windows. For instance, although we had previously placed Tm1/2/4/6 in the Hth/Opa temporal window^28^, because marker genes did not allow us to distinguish these highly similar cell-types in the L3 main OPC, Zhang and colleagues^36^ have shown that Tm1 and Tm4 are generated in the first window of Opa expression (Hth/Opa temporal window) while Tm2 and Tm6 are produced later, in the second wave of Opa expression (Ey/Opa temporal window). Because we lack fine temporal window resolution for most main OPC clusters, we used the broad temporal windows we had previously characterized^28^ (Fig.2F) for our analyses.

### Discovery of additional spatial subdivisions

Our scRNA-seq data suggested the existence of two additional spatial domain subdivisions, which we verified experimentally. First, according to our sequencing results, Tm9d was produced from the dOptix and dDpp domains and Tm9v from the vVsx and vOptix domains (Table S1). Such asymmetry for the dorsal and ventral subtypes of the same neuronal type was unexpected, but we indeed verified that Tm9v markers were present in a small subregion of the vVsx domain that touches the border with the vOptix domain (Fig.2G, Fig.2H, hereafter called “vVsx stripe”), and Tm9d markers were absent from the Vsx region. Second, although Dm1 and Dm12 shared markers that mapped to both sides of the border between the vOptix and vDpp domains (Fig.S3G), and both were found in the vOptix dataset, only Dm12 was also found in the dpp dataset (Annex 1). We experimentally validated^20^ that there exists a new domain that produces Dm12 but not Dm1, where *Optix* and *dpp* overlap (Dpp/Optix domain). This experimental validation of two new subdomains predicted from our scRNA-seq data (Fig.2H) further emphasizes the accuracy of our results. However, it also emphasizes that our determination of spatial origin of the optic lobe neurons could be achieved with even higher resolution by incorporating these domains. For instance, vOptix can in fact be subdivided into 4 subdomains: the anterior third producing pDm8^34^, the middle third producing yDm8^34^, and the posterior third that is split into an Optix+ Dpp-domain producing Dm1 and an Optix+ Dpp+ domain producing Dm12. These domains are partly established by *dpp* signaling, as we recently showed^20^. Notably, this means that neurons present in both the Optix dataset and the Dpp dataset must be produced from the Dpp+ Optix+ domain, but they might also be produced from the Dpp+ Optix-domain. For instance, we assigned Mi1 and Tm1/4/6 as being produced in the whole main OPC, but some evidence (see Table S1) suggests that they may not be produced in the Dpp+ Optix-domain.

### Regulation of terminal selector expression by specification factors

Our characterization of the spatial, temporal and Notch origins of each main OPC neuronal type allowed us to explore how specification mechanisms in progenitors establish the TS combinations we had previously identified in neurons^11,17^. The expression of each TS can theoretically be controlled by a single patterning mechanism, any combination of two patterning mechanisms, or all three patterning mechanisms (Fig.3A). Moreover, the expression of each individual TS can either be regulated similarly in all neuronal types or be controlled through different regulatory mechanisms across neuronal types (Fig.3B). Because neurons are produced in temporal windows that include overlapping expression of several tTFs, and from spatial origins (*e.g.*, dVsx and dOptix) that can encompass the domain of several spatial specification factors (*e.g.*, Vsx, Optix, Salm), we hereafter use “specification module” as a general term that refers to any temporal window, spatial origin, or Notch status that produces a neuronal type. To characterize how the expression of each TS is regulated, we identified the TSs expressed in all neuronal clusters that share one (Fig.S7), two (Fig.S8) or three (Fig.S9) specification modules (Methods): if the expression of a TS is activated by a certain combination of specification modules, this TS should be expressed in all neuronal types that share this combination.

**Figure 3:**
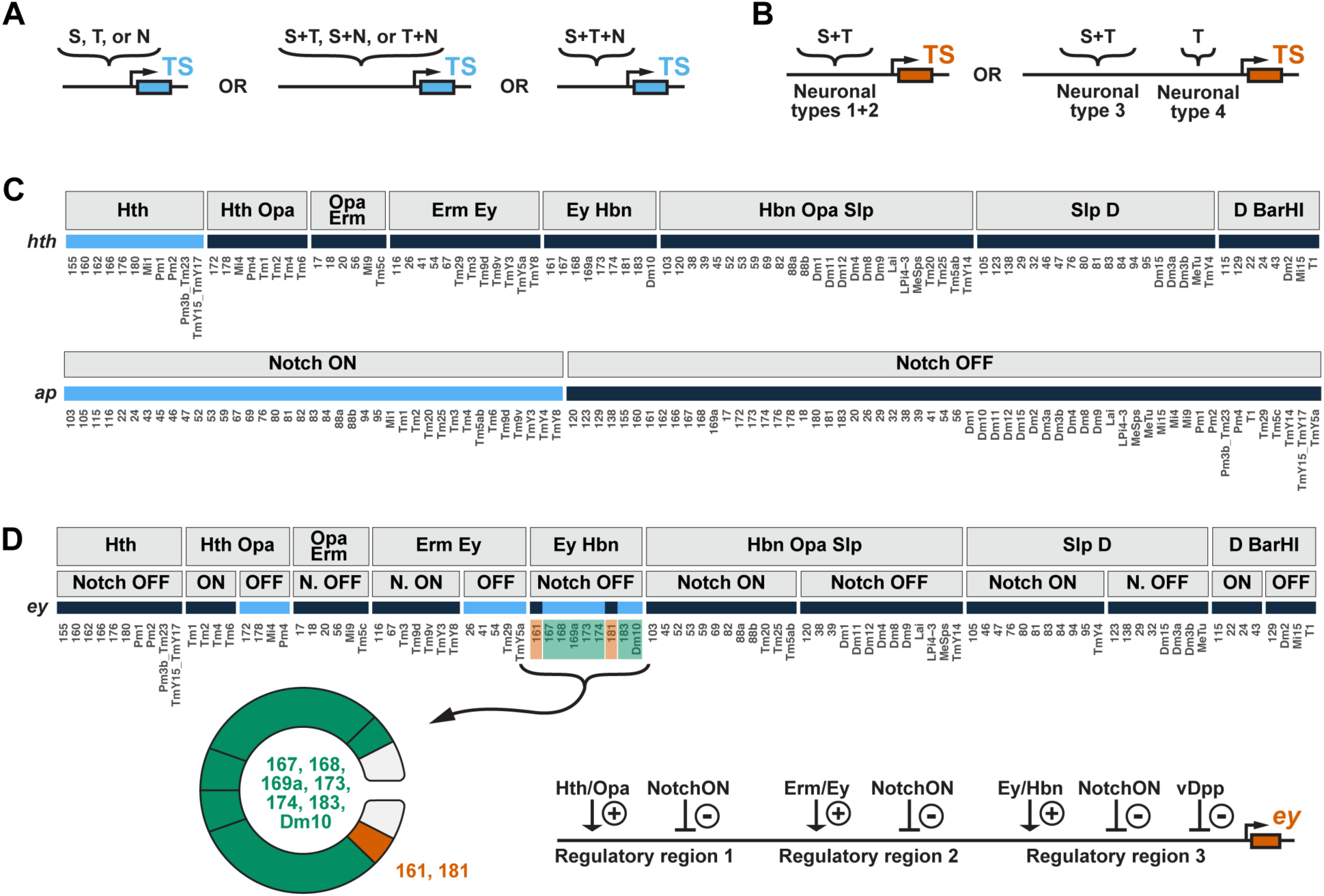
Regulation of terminal selector expression by specification factors. A) A given TS can be regulated by one, two, or three of the main OPC patterning mechanisms. N: Notch origin, S: spatial origin, T: temporal origin. B) A given TS can be regulated similarly in all neuronal types (*e.g.* neurons 1 and 2), or through different mechanisms in different neurons (*e.g.,* neurons 3 *vs.* 4). S: spatial origin, T: temporal origin. C) Only two TSs have their expression perfectly correlated with a single specification module. *hth* (top) is continuously expressed in the main OPC clusters (x-axis) from the Hth temporal window, and *ap* (bottom) is continuously expressed in Notch^ON^ main OPC clusters. Clusters are grouped either according to their temporal window (top) or their Notch status (bottom). Gene expression is represented in light blue. D) The expression of most TSs is correlated with combinations of specification modules. For instance, *ey* is expressed in all Hth/Opa Notch^OFF^ neurons, all Erm/Ey Notch^OFF^ neurons, and all Ey/Hbn Notch^OFF^ neurons that are not from the vDpp domain. A potential model of *ey* expression regulation is indicated (cluster 181 might also be the only Ey/Hbn cluster produced in both the vDpp and the dDpp domain, Fig.2F). Clusters are grouped by combinations of one temporal and one Notch specification module, and gene expression is represented in light blue. Only combinations of two specification modules containing more than 1 cluster have been plotted.

We first used these analyses to identify TSs that could be under the control of a single patterning mechanism. This was the case of *apterous* (*ap,* Fig.3C), which was expected since neurons were identified to be Notch^ON^ based on their expression of *ap*^26,28^. The only other example was *hth*, which is continuously expressed exclusively in all neurons from the Hth temporal window (Fig.3C, Fig.S7). In fact, *hth* is the only concentric gene (Fig.1D) that is expressed in all the neurons produced in a given temporal window. Concentric genes are TSs^17,28,31^ that were already known to respond to both temporal origin and Notch signaling, such as *bsh* that is expressed only in the Notch^ON^ progeny of the Hth temporal window^26^, or to both temporal and spatial origin, as indicated by their regionalized expression pattern in the main OPC^28^. Consistently, our results show that indeed all concentric genes except *hth* are only expressed in a subset of the neurons from various temporal windows, suggesting that they are subjected to additional regulation by spatial patterning or Notch signaling. Finally, we did not identify any TS continuously expressed exclusively in all neurons from any single spatial origin. TSs specific to smaller spatial domains could still exist, but we could not identify them due to the incomplete resolution of our determination of spatial origins.

The expression of most TSs was better correlated with combinations of all three patterning mechanisms, and the same TS was often regulated by distinct combinations of specification modules in different neuronal types. For instance, *ey* is expressed continuously in Notch^OFF^ neurons from both the Hth/Opa and Erm/Ey temporal windows (as a TS in neurons, not as a tTF in neuroblasts) (Fig.3D), regardless of their spatial origin, and from Notch^OFF^ neurons from the Ey/Hbn temporal window except for those produced in the vDpp region (Fig.2F). Therefore, *ey* expression seems to be controlled by at least three different combinations of specification modules (one potential scenario is shown in Fig.3D).

Similar to what was presented above for *ey*, the tables in Fig.S7, Fig.S8, and Fig.S9 can be used to form hypotheses on the specification modules controlling the expression of other TSs. For each TS, it is important to consider the largest group of neurons that share both the expression of the TS and a given combination of specification modules. For instance, *ap* is expressed in all the six dOptix clusters (Fig.S7A): one could therefore hypothesize that *ap* is turned on by the dOptix specification module. However, all thirty-nine Notch^ON^ clusters, regardless of spatial origin, express *ap* (Fig.3C). This indicates that *ap* expression is not controlled by spatial patterning and is instead controlled by Notch signaling, as was previously shown^25^.

Our analyses also highlight the interdependency between TS expression and the combinations of specification modules that produce neuronal types. For instance, all dOptix-only clusters are either from the Hbn/Opa/Slp or the Slp/D temporal windows (Fig. S8B). Neurons from other temporal windows are therefore never produced exclusively from the dOptix domain: they must either be produced from larger regions that encompass dOptix (for instance Tm3 is from the Erm/Ey temporal window and is produced across the Optix and Vsx domains, Fig.2F), or be produced exclusively from the dOptix domains but be culled. This also implies that no TS is under the control of dOptix associated with temporal windows other than Hbn/Opa/Slp or Slp/D (since no neuronal type is produced from these combinations of specification modules).

### Regulation of the presence of neuronal features by the terminal selectors

Next, we explored the regulation of gene expression by the TSs, first by studying the regulation of the cholinergic marker *VAChT*, the glutamatergic marker *VGlut,* and the GABAergic marker *VGAT* in main OPC neurons. *VAChT* is expressed in 37 clusters (Fig.4A): although it could be under the control of different TS combinations in each of these neuronal types (Fig.4B), this would require 37 regulatory mechanisms. A more parsimonious possibility is that it is under exclusive control of one or more TSs that are present in all these neuronal types (Fig.4C). This is almost the case of the TS *ap*^28^, an activator of cholinergic identity in the optic lobe^12^, which is expressed in 34 out of 37 *VAChT*+ clusters (Fig.4A). The three remaining *VAChT*+ clusters that do not express *ap* are the Notch^OFF^ neurons of the D/BarH1 temporal window. Moreover, the four Notch^ON^ clusters from the same D/BarH1 temporal window are the only *ap+* neurons that do not express *VAChT* (Fig.4A). This suggests that *ap* is a global activator of cholinergic phenotype, except in neurons from the D/BarH1 temporal window where the TSs common to all Notch^ON^ neurons could prevent the activation of *VAChT* expression by *ap*, and the TSs common to all Notch^OFF^ neurons could activate the expression of *VAChT* in the absence of *ap*. Therefore, 3 regulatory mechanisms instead of 37 would be sufficient to explain *VAChT* expression.

**Figure 4:**
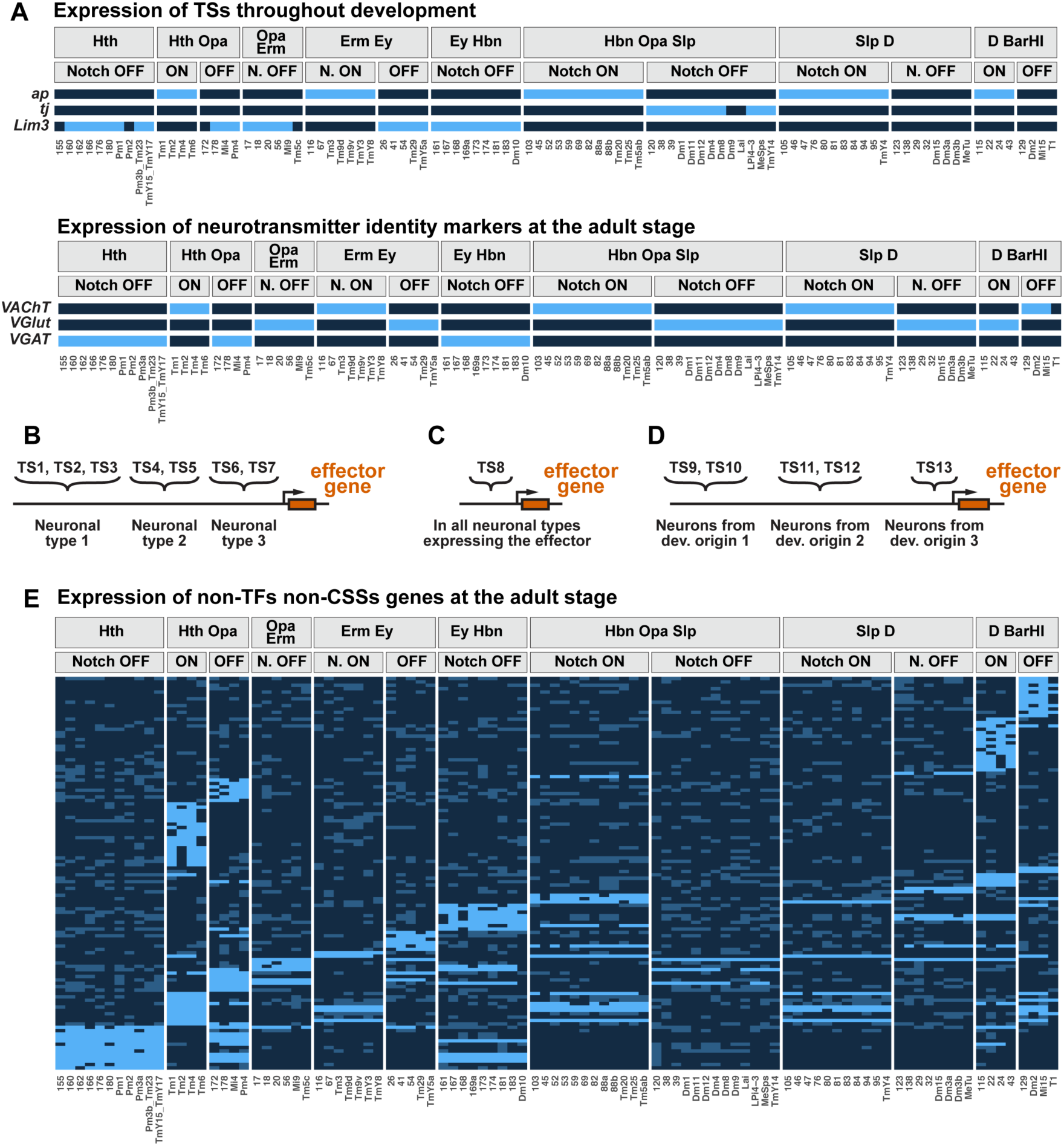
Regulation of effector gene expression by terminal selectors. A) Expression of the indicated genes (y-axis) in main OPC clusters (x-axis) grouped by combinations of one temporal and one Notch specification module. Light blue indicates either the clusters in which the TSs are continuously expressed (top), or the clusters in which the markers of neurotransmitter identity are expressed at the adult stage (bottom). B-D) The expression of a given effector gene can be controlled by different TSs in each neuronal type (B), by the same TS in all neuronal types (C), or by the same TSs in all neuronal types sharing a given developmental origin (*i.e.*, any combination of spatial, temporal and Notch origin) (D). E) Expression of non-TF non-CSS genes (y-axis) in main OPC clusters (x-axis) grouped by combinations of one temporal and one Notch specification module. The names of the genes are indicated on the plot in Annex 2. Light blue: more than 75% of clusters from a given combination of specification modules express the gene at the adult stage, Medium blue: fewer than 75% of clusters from a given combination of two specification modules express the gene at the adult stage, Dark blue: gene not expressed (Methods). A, E) Only combinations of two specification modules containing more than 1 cluster have been plotted.

Thus, *VAChT* expression can be explained by combining temporal and Notch patterning, and it is therefore likely to be controlled by the expression of the TSs specific to the corresponding combinations of temporal and Notch origin (Fig.S8A). Similar observations can be made with the TS *tj*, a regulator of glutamatergic identity^12^ marked by *VGlut*, and the TS *Lim3*, a regulator of GABAergic identity^12^ marked by *VGAT* (Fig.4A). Therefore, neurotransmitter identity is correlated with combinations of temporal windows and Notch status in the main OPC^28^, and therefore with the TSs specific to each of these combinations of origins (Fig.S8A).

This suggests that the expression of effector genes can be regulated by the TSs corresponding to a given combination of specification modules (*i.e.*, a given developmental origin, Fig.4D). To test whether this mode of regulation could be generalized to other genes, and whether it could be extended to morphological features, we first surveyed the Virtual Fly Brain website as well as more than 40 publications to build a table (Table S2, Methods) summarizing about 600 morphological features for the more than 70 neuronal types for which we have identified the corresponding clusters in our scRNA-seq atlas. This table can be amended with newly identified clusters, or new characterizations of neuronal morphologies^9,10^, and used as the basis for further high-throughput studies of the regulation of neuronal morphology. This table, as well as binarized gene expression data, allowed us to identify genes and morphological features present in all neuronal types sharing any combination of one to three specification modules at each developmental stage, consistent with their regulation by TSs associated with the corresponding developmental origin (Fig.4D). All plots are provided in Annex 2 for gene expression (for readability, we split the genes into TFs, Cell Surface and Secreted (CSS) proteins, and non-TF/non-CSS genes) and in Fig.S10 and Fig.S11 for morphological features. Fig.4E presents the example of non-CSS and of non-TF genes at the adult stage in neurons sharing the same combination of temporal and Notch origin, to illustrate that many genes follow the same expression pattern as the markers of neurotransmitter identity and therefore may be under the control of the same TSs.

Our results show that neurons sharing combinations of specification modules share the expression of some TSs, and some modules of neuronal features (effector genes and morphological features). These TSs are therefore good candidates to regulate their associated module of neuronal features, which was for instance validated^12^ for *ap*, *tj* and *Lim3* in the case of neurotransmitter identity (Fig.4A). Although validation of these correlations is out of the scope of this paper, this can serve as a basis for future studies. These results also suggest that some TSs can regulate the same targets in all neurons sharing combinations of specification modules, regardless of the other TSs expressed by these neurons. This and other results^17^ suggest that the targets of the TSs expressed in a given neuronal type do not always overlap. This is consistent with a model whereby it is not required that all TSs be involved in the regulation of all terminal effectors in a given neuron, but rather, any effector gene can evolve to be regulated by any combination of available TSs in that neuron. Because the expression of many differentially expressed genes was not associated with any combination of specification modules, our work could not address their regulation. Further studies will be required to understand how the expression of these genes is regulated by TS expression.

### Specification modules and terminal selectors encode similar information

Our results so far are consistent with the TS model whereby the information carried by the patterning mechanisms determining each neuronal identity are propagated and maintained by a TS code that regulates specific neuronal features. According to this model, TS expression in neurons must be sufficient to fully encode the information inherited from the specification modules in the progenitors. We tested this model in a high-throughput manner using machine learning. Intuitively, if several neuronal types share a common feature (*e.g.* targeting a given medulla layer or expressing a specific neurotransmitter transporter), and all specifically express a TF, this TF is a good candidate regulator of this feature. Its expression could therefore be used to predict the presence of the feature in other neuronal types. To identify such correlations for all neuronal features we used the random forest machine learning algorithm^37^ (Fig.5A). For each neuronal feature at any given developmental stage from P15 to adult, we trained a random forest model on a randomly sampled subset of clusters (“training set”). This established a model that correlates the expression of potential regulators with the presence or absence of a given neuronal feature. These potential regulators, called predictors, were either TSs, TFs, CSSs, or the specification modules, and the neuronal features predicted were either the expression of a gene or the presence of a morphological feature (Table S2). To assess the quality of each model, we used them to predict the expression of the feature they were trained to predict in a “test set”, *i.e.,* neuronal clusters not used to train the model and for which we already knew the real status of the feature. The similarity between predicted and actual status of the feature was then evaluated by using the Matthews Correlation Coefficient (MCC, a value of 1 denotes perfect correlation and -1 perfect anti-correlation, see Methods).

**Figure 5:**
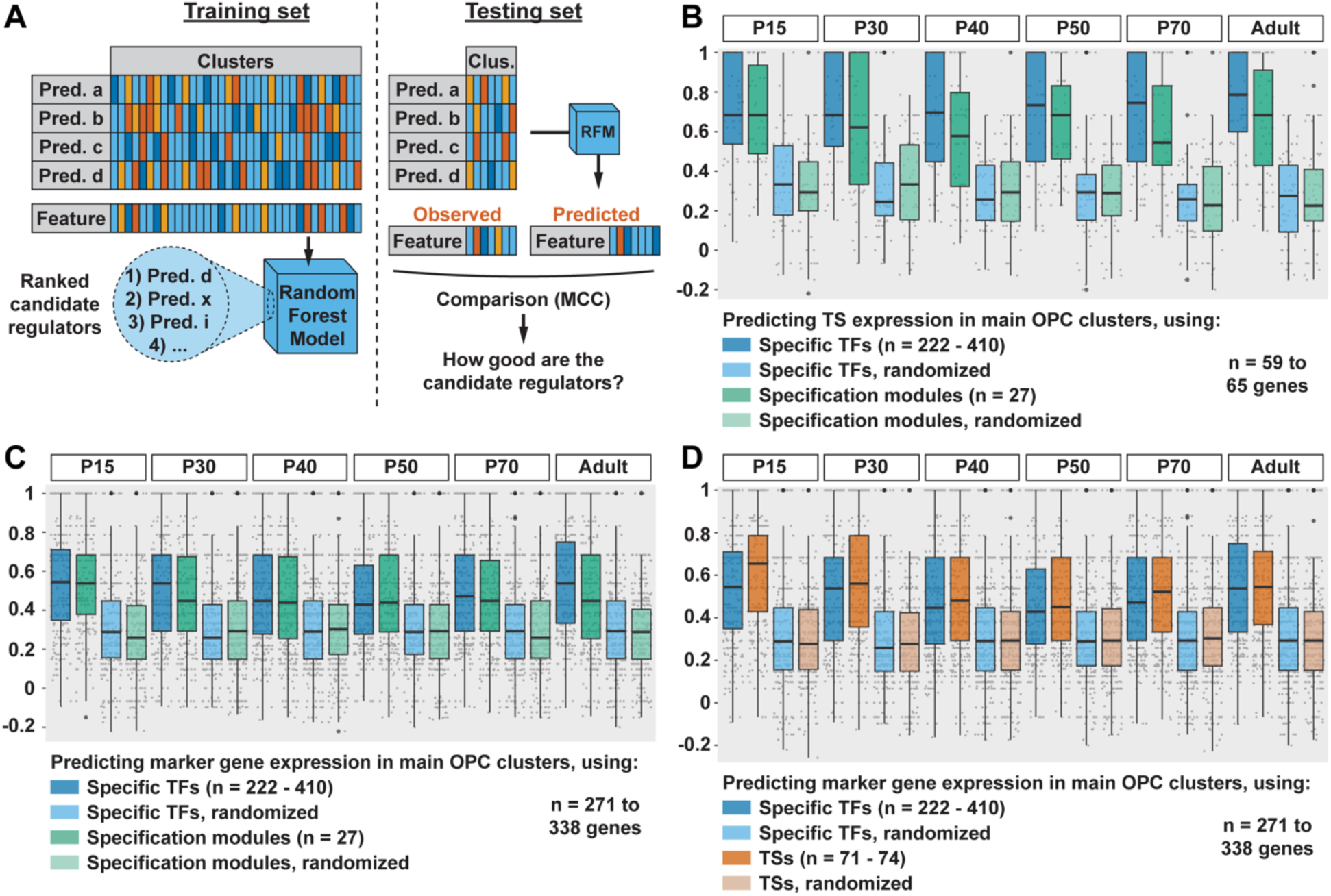
Specification modules and terminal selectors encode similar information. A) Machine learning approach used to find candidate regulators (TFs, TSs, CSSs, or specification modules) for neuronal features (gene expression or morphologies). Clus. = clusters, Pred. = predictor, RFM = Random Forest Model, MCC = Matthews Correlation Coefficient. B-D) Matthews Correlation Coefficient between the observed expression of TSs (B) or marker genes (C, D), and their expression predicted using either specific TFs (*i.e.*, TFs differentially expressed between neuronal types, B-D), specification modules (B, C), or TSs (D). Each gray dot represents the MCC value for a given gene, and the values are summarized as boxplots that display the first, second and third quartiles. Whiskers extend from the box to the highest or lowest values in the 1.5 interquartile range, and outlying data points are represented by large black dots.

We first built models using developmental origin to predict the expression of TSs at all stages by using the specification modules as predictors. We obtained high predictivity (median MCC around 0.6, Fig.5B) and, as expected, low predictivity (median MCC around 0.3, Fig.5B) when building the models using randomized specification modules. Considering the imperfect resolution of our determination of both temporal and spatial origins, this suggests that TS expression in neurons is highly correlated with specification modules, consistent with the previous sections. However, developmental origin performed worse when predicting the expression of TFs other than TSs (median MCC around 0.3, Fig.S12A), suggesting that the expression of these TFs is not directly controlled by the temporal, spatial and Notch related specification mechanisms.

Second, we used either specification modules or TS expression to predict the expression of marker genes (*i.e.,* differentially expressed genes that are therefore the most representative of each neuronal identity). We compared the results to those obtained using all differentially expressed TFs as predictors of marker gene expression. Both TS expression and developmental origin performed similarly to using all differentially expressed TFs to predict the expression of marker genes, and sometimes performed even better (Fig.5C and D). As expected, when using spatial, temporal, or Notch origin independently, the predictivity was lower than when using all three combined (Fig.S12B). Despite their much lower number, specification modules (n=27) and TSs (n ∼ 75 in the main OPC) are thus sufficient to encode the information contained in all differentially expressed TFs (222<n<410 depending on developmental stage). Similar results were obtained when predicting morphological features (Fig.S12C, D), although only 12-60 features could be tested (see Methods), compared to hundreds of marker genes. These results show that developmental origin, TSs, and all differentially expressed TFs, can predict neuronal identity with a similar power. When predicting the expression of genes that are neither marker genes nor pan-neuronal (*i.e.*, expressed in many but not all neurons), however, the performance was slightly lower for TSs, and much lower for developmental origin (Fig.S12E, F). This could be because the regulation of their expression is more complex than the one of marker genes: for instance, more broadly expressed genes may be more likely to undergo phenotypic convergence^12^ (*i.e.*, to be regulated differently in different neuronal types) because they are expressed in neurons from a wider variety of developmental origins.

In addition to using TF expression to predict gene expression or the presence of morphological features (Fig.S13A, B), we also produced models predicting the presence of morphological features from the expression of CSSs (Fig.S13C). Each trained random forest model provides a ranking of the predictors according to how important they are for the performance of the model. If a model performs well, it has identified correlations present both in the training and the test set, and therefore its best ranked predictors are candidate regulators of the feature of interest. Although it is out of the scope of this study, this can be tested experimentally. For instance, using all TFs to predict the expression of neurotransmitter-related genes we identify *Lim3* as the best candidate regulator of *VGAT* and *Gad1*, and *ap* as best candidate regulator of *VAChT* and *Cha* at the adult stage. This was expected since when knocking down *Lim3* and *ap* in the adult brain, the levels of *Gad1* (maker of GABAergic identity, like *VGAT*) and *Cha* (maker of cholinergic identity, like *VAChT*) are downregulated^12^. The known regulator of glutamatergic identity *tj* is ranked low in the list of candidate regulators of *VGlut*, but this is expected because *tj* is only expressed in a subset of glutamatergic neurons (Fig.4A). Instead, the five best candidate regulators are *fd59A*, *tup*, *toy*, *Ets65A*, and *kn*, whose role could be tested experimentally. We provide in Annex 3, for all neuronal features, tables listing their best candidate regulators which can be used by the community to explore in more detail the regulation of given features (see guidance on how to use and interpret these tables in Methods).

### Specification modules may not be remembered epigenetically after terminal selector combinations are established

Because the expression of the spatial specification factors is not maintained in neuroblasts, the information they carry must be propagated to differentiating and adult neurons by epigenetic mechanisms or by downstream TFs. In the *Drosophila* ventral nerve cord, the TF *hunchback* regulates different targets in neuronal types from different spatial origins because these neuronal types have different chromatin accessibility^38^. This led to the hypothesis that, in the optic lobe, the information encoded by the spatial factors could be maintained epigenetically in neurons, most likely by differential chromatin accessibility between neurons from different spatial domains^22,39^. The discovery of the TS combinations implies that such an epigenetic mechanism would only be necessary to carry the spatial information from neuroepithelium to young neurons, in which the TS combinations are already established. Nevertheless, we tested the possibility that TF (including TSs) targets might be affected by the spatial origin of the neuronal types even after the establishment of TS combinations, *i.e.*, in developing or adult neurons. If TSs were not sufficient to encode all differential gene expression between neurons from different spatial domains, using random forest to predict gene expression with both developmental origin and TS expression together would be expected to have higher predictive power than TS expression alone. However, using specification modules as predictors in addition to TF or TS expression did not lead to increased predictivity of our random forest models for marker gene expression (Fig.S14A, B). This suggests that either there are no epigenetic mechanisms “remembering” specification modules in developing or adult neurons (past the early specification of neurons and the early expression of TSs) in addition to the TS combinations, or that if epigenetic mechanisms do exist, they encode information redundant with the expression of TSs. This would be similar to what happens in the *Drosophila* ventral nerve cord where a spatial TF regulating chromatin accessibility, *gooseberry*, remains expressed in neurons^38^.

## Discussion

### Specification factors can encode an excess of neuronal types

This work extends our comprehensive description of the spatial, temporal, and Notch origins of all neuronal types in the main OPC. Previously, it was thought that 10 temporal windows, 2 Notch statuses, and 6 spatial domains (v/dVsx, v/dOptix, v/dDpp) could specify the identity of up to 120 potential neuronal types in the main OPC^25,28^ (10x2x6). This was a number sufficient to explain the specification of the ∼100 neuronal types that were thought to exist in the medulla^11,14,15^. However, we have described additional spatial domains in this study and others^20,34^, and our previous determination of the temporal origin of main OPC neurons was conservative due to the use of only a subset of marker genes, which were sometimes insufficient to separate broad temporal windows into narrower ones^28^. Lastly, Arain et al.^18^ recently discovered a fourth axis of neuronal specification, in which progenitors from the same spatial domain and in the same temporal window, but produced at different time points during the progression of the neurogenic wave that generates neuroblasts during larval development, generate different Ap-(Notch^OFF^) neuronal progeny^18^.

Together, these results suggest that the specification factors could potentially encode hundreds of neuronal types, *i.e.*, considerably more than the number of neuronal types that are actually produced in the main OPC, even though numerous additional neuronal types have recently been discovered from connectomics studies^9,10^. On the other hand, some neuronal types could be culled during development by cell death (this is for instance the case of Notch^OFF^ neurons of the Hth temporal window in the Optix domain^25^). Furthermore, many neuronal types are produced across several spatial domains (e.g., Mi9 is produced in the vDpp, vOptix, dDpp and dOptix domains), which reduces the diversity of neuronal types that can be encoded by spatial factors. Moreover, our work highlights that neuronal types are sometimes produced across spatial domains, and sometimes not, and provides testable hypotheses for how neuronal type production is regulated. For instance, Mi1 is produced in both Optix and Vsx, while Mi9 is produced in Optix but not Vsx (Fig.3F). This could be explained if the expression of all Mi1-specific TSs was activated (or not repressed) by both Optix and Vsx, while the expression of at least one of the Mi9-specific TS was either activated by Optix or repressed by Vsx. Similar reasoning can be applied to explain the spatial origin of any given neuronal type and can be tested experimentally.

### Regulation of neuronal abundance in the main OPC

We previously showed that the Notch^ON^ progeny of the Hth temporal window produces Mi1, a synperiodic neuron produced from the whole main OPC, while the Notch^OFF^ progeny produced several infraperiodic neuronal types from restricted spatial domains^18,25^ and we suggested that Notch^ON^ neurons ignore spatial patterning while Notch^OFF^ neurons do respond to spatial information. This appears to be correct for the early temporal windows that generate Mi1 in the Hth window, Tm1 and Tm4 in the first Hth/Opa window and Tm2 and Tm6 in the second Ey/Opa window from the Vsx, Optix and Dpp domains. However, at later temporal windows, most Notch^ON^ neurons are produced in restricted parts of the main OPC and most are infraperiodic (Fig.2F). On the other hand, a large majority of Notch^OFF^ neuronal types are produced from restricted spatial domains and are infraperiodic, although some (such as T1, Dm3, Dm2 and Tm5c) appear to be generated by the entire main OPC and are synperiodic. In addition, Notch^ON^ neurons tend to be more abundant and to come from less restricted domains, especially in the early temporal windows (19/39 Notch^ON^ from all windows have more than 500 neurons in our experimental adult clusters) while Notch^OFF^ neurons are less abundant (only 19/65 have more than 500 neurons in the clusters) (Fig.2F).

Therefore, neither the Notch status nor the size of the spatial domains correlate perfectly with the production of synperiodic neurons. For Dm neurons, the size of the spatial domain correlates with neuronal abundance^20^. However, some abundant neurons come from small domains (see Methods for neuronal type abundances). For instance, ∼800 Mi1 are produced from the entire main OPC neuroepithelium while 550-800 Dm8^9,10,20,34^ come from a much smaller vOptix domain. This could be explained if the neuroblasts produce Dm8 for a longer time-period than they produce Mi1^20^.

### Establishment of terminal selector combinations

We had previously identified combinations of TSs specific to each optic lobe neuronal type^11^, and showed that modifying these combinations could convert the identity of one neuron into another^17^. However, how these combinations are established, and how they control downstream neuronal features, is unknown. Our work suggests that the expression of a given TS can be controlled through multiple regulatory mechanisms, each activated by a different combination of one to three specification modules. This entails that the TS combination specific to a given neuronal type is established by integrating different subsets of the patterning information for different TSs (Fig.6). It is likely that improving our characterization of the developmental origins of neurons will improve the correlations we identified, as these results were obtained despite the imperfect resolution of our determination of the temporal and spatial origins, and the lack of characterization of a fourth axis of neuronal specification^18^.

**Figure 6:**
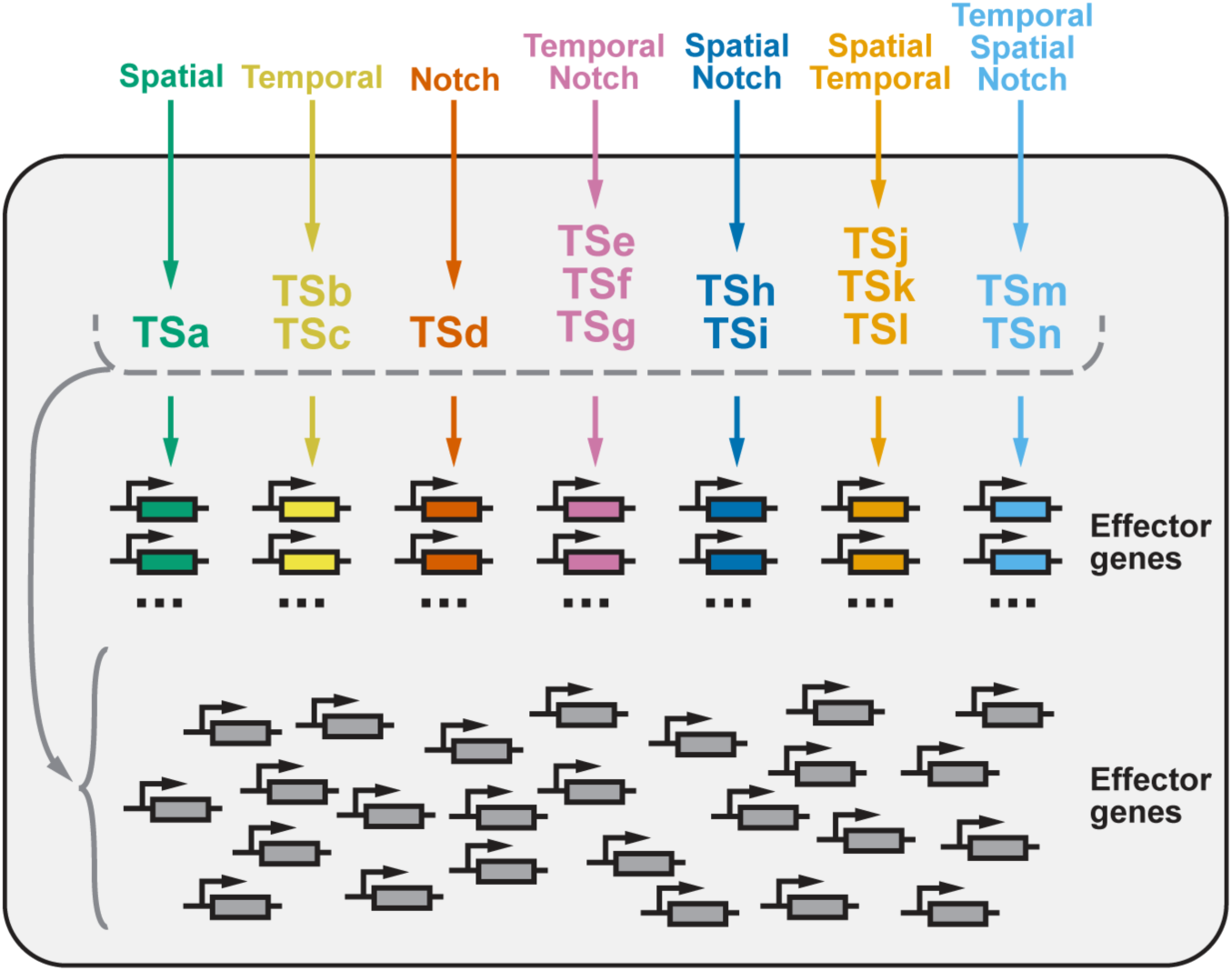
Establishment and maintenance of neuronal features in the *Drosophila* optic lobe. Each neuronal type (grey box) expresses a unique combination of TSs (TSa-n, top part of the box). The expression of each of these TSs is established either by one, two, or three specification modules (spatial origin, temporal window, Notch signaling). Moreover, some effector genes (in colors, middle part of the box) are regulated by the TSs corresponding to a given combination of specification modules (ex: a given Notch status activates the expression of TSd, which activates the expression of the red effector genes, in all neurons sharing this Notch status). The expression of each effector gene represented in grey is not correlated with any combination of specification modules, and can be regulated by any combination of the TSs expressed in this neuronal type.

Although our work provides correlations between combinations of specification modules and TS expression, which could be used as a basis to identify motifs bound by specification factors in the enhancers of TSs, further work is required to validate these correlations and characterize the corresponding molecular mechanisms. Notably, in some cases the expression of a TS could be regulated indirectly by a specification factor, and some TSs have also been shown to cross-regulate each other^17,40^. In addition, because spatial factors are expressed in the neuroepithelium but not in the neuroblasts, temporal TFs are expressed in neuroblasts but rarely maintained in neurons, and Notch effectors are only transiently expressed in newborn neurons, it will be of interest to understand when exactly the TS combinations are established. This could be an iterative process, in which for instance epigenetic marks (*e.g.*, chromatin accessibility) are established in the neuroepithelium by the spatial factors, these epigenetic marks interact with tTFs and produce preliminary TS combinations in neuroblasts and/or ganglion mother cells, which are then finalized by Notch signaling in newborn neurons. It could also happen in a single step in newborn neurons, provided that spatial and temporal origins are remembered up to this point, for instance by the expression of intermediate factors or by epigenetic mechanisms.

### Implications for the evolution of neuronal diversity

Our results have implications for neuronal type evolution. New neurons could appear following the emergence of a new tTF (producing a new temporal window affecting all spatial domains), or of a new spatial specification factor (producing a new spatial domain affecting all temporal windows). However, these large-scale modifications of neuronal specification could have deleterious effects, either by leading to the appearance of many new neuronal types at once, or by affecting existing specification mechanisms. For instance, the expression of a new tTF would likely overlap with the expression of existing tTFs, therefore affecting existing temporal windows in addition to creating a new one.

On the other hand, because the specification factors can encode more neuronal types than the ones produced in the optic lobe, single neuronal types could emerge either by rescuing a type previously eliminated by cell death, or for instance by splitting a spatial domain within a single temporal window (for instance, by producing one neuronal type from vOptix and one from vVsx, instead of only one from vOptix/vVsx). Because any effector gene can evolve to be regulated by any combination of available TS, most of the genes expressed in a newly evolved neuronal type cannot be predicted. Nevertheless, according to our model (Fig.6), a newly produced neuronal type from a given temporal window and of a given Notch status should present a pre-determined module of molecular and morphological features associated with its temporal+Notch origin, such as the expression of *VAChT* in all Notch^ON^ neurons that are not produced in the BarH1 temporal window (Fig.4A). In addition, such a neuronal type would also present modules of features associated with any combination of its specific temporal, spatial and Notch origins. These pre-determined modules of features could bias newly evolved neurons to integrate into given circuits or to assume given functions: similar environments could lead to the selection of new neuronal types with similar functions, and therefore produced according to similar specification mechanisms. Such reasoning could be tested experimentally.

## Supporting information

Annex 1

Annex 2

Annex 3

## Acknowledgments

We thank all members of the Desplan and Konstantinides laboratories for helpful feedback while working on the manuscript, and Oliver Hobert for critical reading of the manuscript. The Desplan laboratory was supported by grants from the National Eye Institute R01EY017916 and R01EY13010, and by Tamkeen under the NYU Abu Dhabi Research Institute Award to the NYUAD Center for Genomics and Systems Biology (ADHPG-CGSB). Research in the Konstantinides lab is supported by the European Research Council (ERC) under the European Union’s Horizon 2020 research and innovation program (grant agreement No. 949500). F.S. was supported by New York University (MacCracken Fellowship and Dean’s Dissertation Ph.D. Fellowship). I.H. was supported by a Human Frontier Science Program Postdoctoral Fellowship LT000757/2017-L and by a senior postdoctoral fellowship from the Kimmel Center for Stem Cell Biology. Y.-C.C. was supported by New York University (MacCracken Fellowship), by a NYSTEM institutional training grant (Contract #C322560GG), and by a Scholarship to Study Abroad from the Ministry of Education, Taiwan. J.M. was supported by NIH fellowships from the National Eye Institute and the BRAIN initiative (F32EY028012 and K99EY032269, respectively). P.V. was supported the Vision Science Research Program (University of Toronto: Ophthalmology and Vision Sciences and UHN), the Ontario Graduate Scholarship, and the Queen Elizabeth II/Pfizer Graduate Scholarship in Science and Technology. T.E. was supported by an NSERC Discovery Grant (RGPIN2015-06457). M.N.O. has been supported by K99/R00NS125117 from the National Institute of Neurological Disorders and Stroke, and Stowers Institute for Medical Research. This work was supported in part through the NYU IT High Performance Computing resources, services, and staff expertise.

## Author contributions

F.S. and C.D. conceived the project and wrote the manuscript. F.S., I.H., Y.-C.C., J.M, P.V, T.E, M.N.O., N.K. and C.D. interpreted the results. F.S., Y.-C.C., J.M, P.V. and T.E. built and tested the spatial origin lines. Y.-C.C. performed the re-analysis of the connectomics data for the medulla neuropil. C.N.-D. contributed to the building of *Drosophila* lines and to the writing of the code for scRNA-seq data analysis. I.H. experimentally described the spatial domain of origin of many medulla neurons and performed the experimental validation of the spatial origins from scRNA-seq data. R. N. E.-D. and K. K. produced the *wg* dataset. F.S. produced all other scRNA-seq data and performed all other analyses.

## Data availability

All raw and processed data are available in GEO (GSE254562). The scripts used to process and visualize the data will be made available upon publication of the manuscript.

## Supplementary figures

**Supplementary figure 1:**
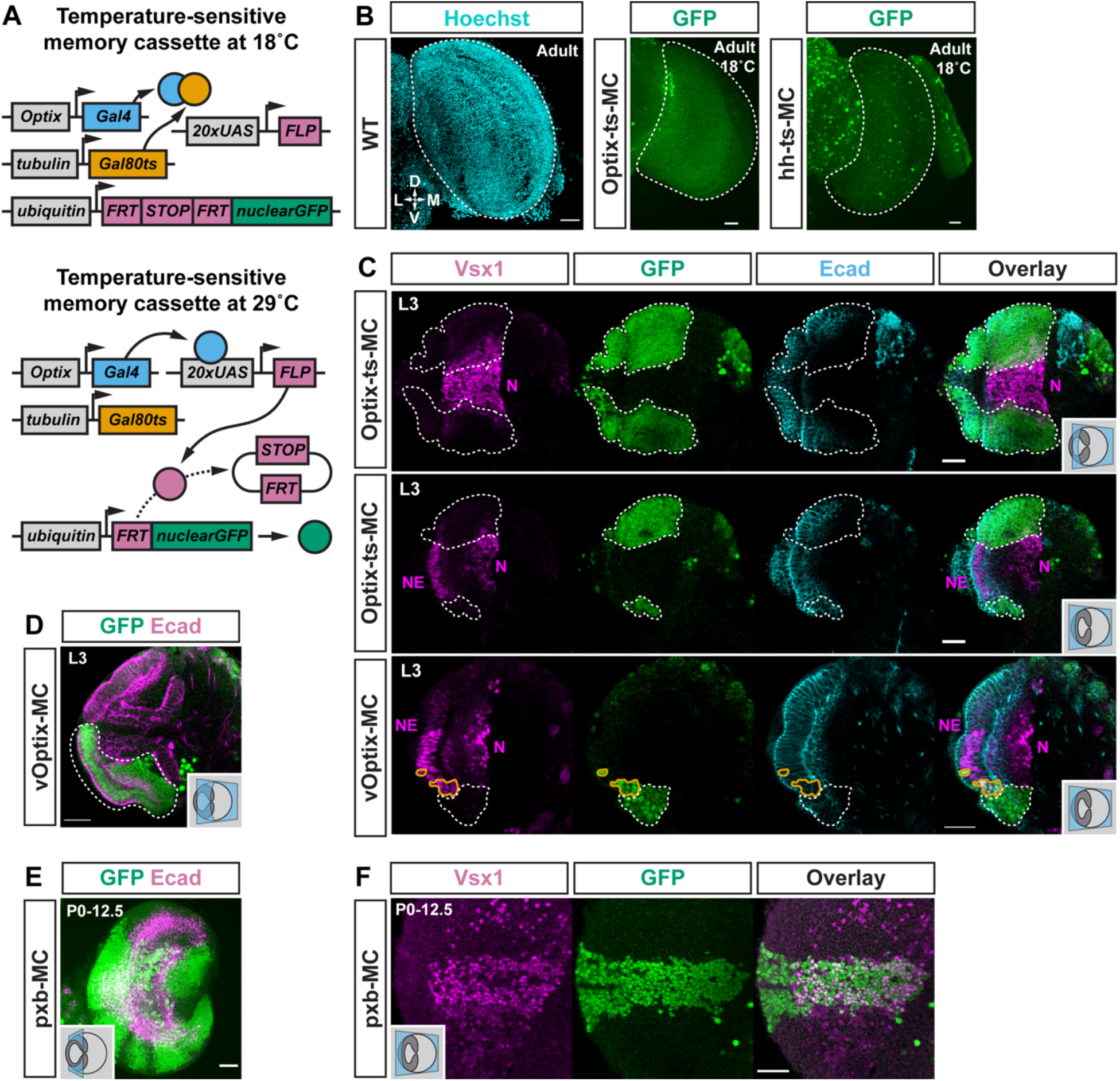
Characterization of the lines used to identify the spatial origin of main OPC neurons by scRNA-seq. A) Schematic describing how a temperature-sensitive (ts) memory cassette (MC) can be used to label all progeny of the cells expressing Gal4 (here driven by *Optix-Gal4*) when placed at the permissive temperature of 29°C. Memory cassettes that are not temperature-sensitive lack the *tub-Gal80^ts^* construct. In this case, it would label any cell that ever expressed *Optix-Gal4*, at any temperature, as well as its progeny. B) Hoechst labelling of an adult optic lobe, and expression pattern of Optix-ts-MC and hh-ts-MC, in adult flies grown exclusively at 18°C. All images are maximum intensity z-projections of 15 slices spanning the whole optic lobe, showing that the number of cells labelled in Optix-ts-MC and hh-ts-MC is negligible compared to the number of optic lobe nuclei, and that the Gal80^ts^ efficiently prevents activation of the MC. D: dorsal, L: lateral, M: medial, V: ventral. Dashed lines: optic lobe, with the lamina excluded. C) Top: Optix-ts-MC third-instar larval expression pattern showing that some neurons from the Optix domain express Vsx1. Middle: additional view from the same brain showing that the Vsx domain neuroepithelial cells (high Vsx1+) are not labelled by the Optix-ts-MC line. Bottom: the vOptix-MC line is ectopically expressed in neuroepithelial cells from the vVsx stripe (circled in orange). N: neurons, NE: neuroepithelium. White dashed line: GFP expressed in the main OPC. For orientation, see schematics and Fig.1C. D) In about 25% of the optic lobes, vOptix-MC is ectopically expressed in the ventral tip of the OPC (which is not part of the Optix domain). Labelled neurons outside of the dashed lines are not part of the optic lobe. For orientation, see schematic and Fig.1C. E) In some optic lobes, pxb-MC labels parts of the main OPC that were not produced in the Vsx domain (this shows an extreme example, at early pupal stage). We therefore checked the pattern of GFP expression of each optic lobe by epifluorescence microscopy before sorting and sequencing. However, due to the low resolution of this technique, it is conceivable that small amounts of false positive cells were not detected. For orientation, see schematic and Fig.1C. F) pxb-MC expression pattern, showing that pxb-MC labels a band of Vsx1+ cells that corresponds to the Vsx domain at early pupal stage, and that pxb-MC does not label the Vsx1+ cells scattered in the Optix domain that can also be seen on Fig.S1C. For orientation, see schematic and Fig.1C. (C-F) The immunostainings were done following the protocol summarized on Fig.2B. Unless indicated, all scale bars are 20 µm.

**Supplementary figure 2:**
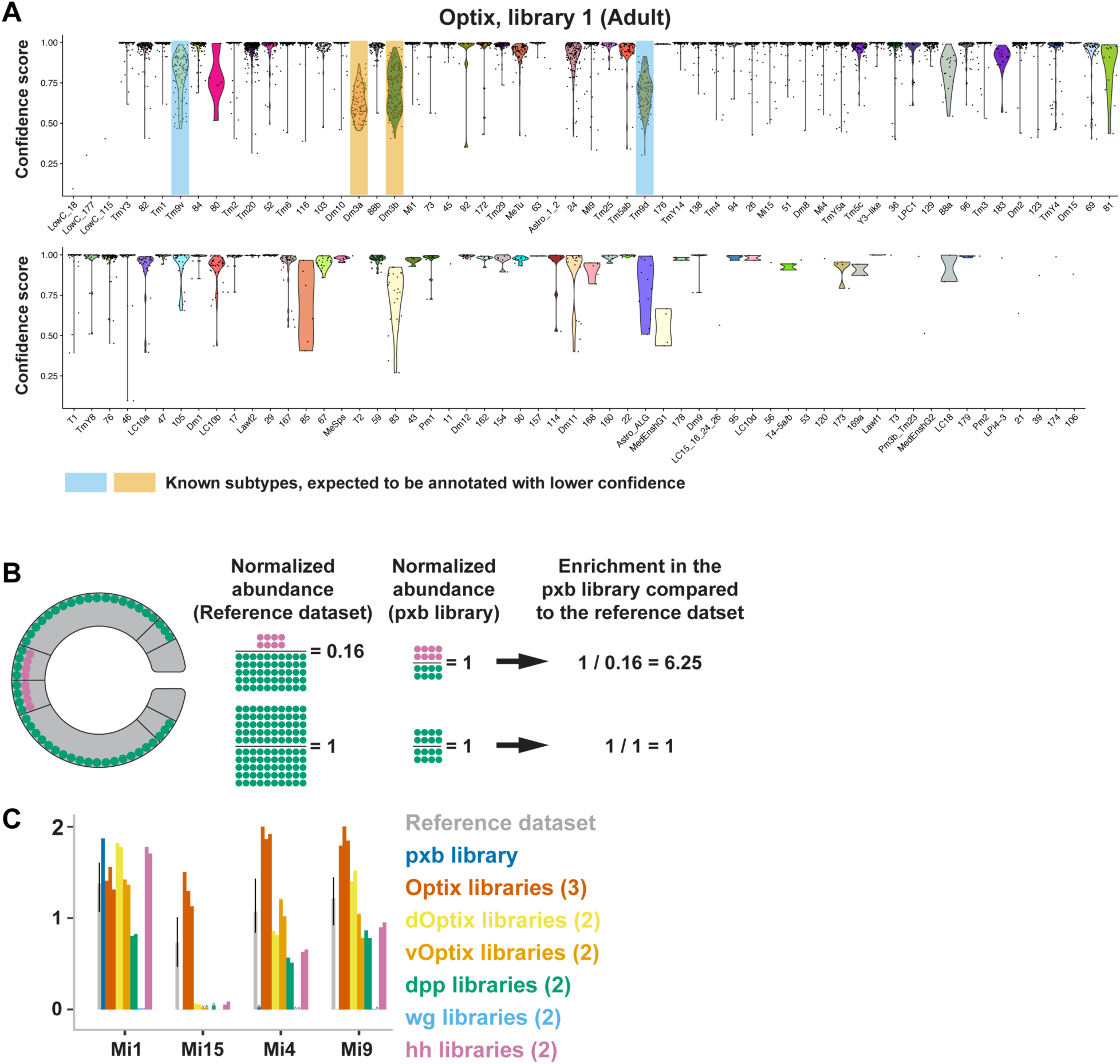
Confidence in cell annotations and calculation of a Class enrichment score in the scRNA-seq libraries. A) Confidence in the annotation of each cell, grouped by Class, of one of the Optix libraries. Each dot is a cell, and the violin plots show their distribution. This shows that although annotation confidence is high for most Classes, it is (as expected) lower for Classes corresponding to transcriptionally similar neurons (Dm3a/b, Tm9v/d). Classes with more than 80% of the cells with a score below 0.5 are prefixed with “LowC”, *i.e.,* “Low Confidence”. B) Schematic explaining the rationale of abundance normalization and enrichment scores. The normalized abundance of a neuron exclusively produced in the Pxb domain is higher in a dataset produced from the Pxb domain (even though the number of cells, here 8 pink cells, is the same across both datasets). A reference dataset is a dataset containing neurons randomly sampled from all main OPC domains. C) Normalized abundance of the indicated Medulla intrinsic (Mi) neurons in the different libraries.

**Supplementary figure 3:**
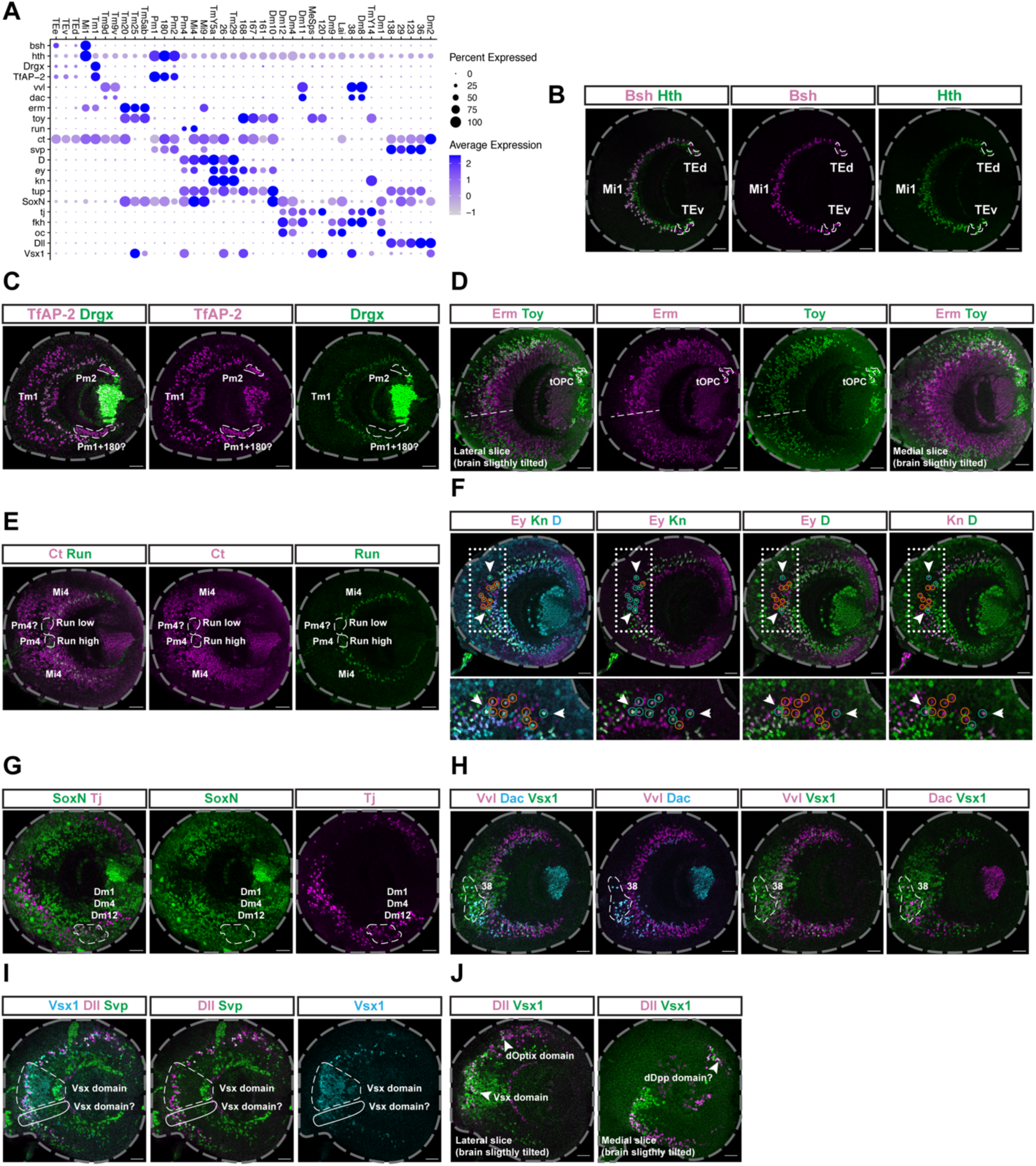
Experimental validation of the spatial origin of main OPC neurons. This figure shows the domain of expression of various TFs, used as markers to confirm spatial origins obtained from scRNA-seq data. The markers used for each cell type, and how their expression was used to assess spatial origin, is detailed in Table S1. A) Expression of the indicated TFs (y-axis) in the indicated neuronal clusters (x-axis), at P15. For each neuronal cluster, their markers can be expressed in a main OPC domain larger than their spatial origin, because most of the markers are expressed in several neuronal clusters (some not shown in this panel). B-J) Expression pattern of the indicated markers in the main OPC at the L3 stage. Grey dashed line: outline of the optic lobe, white dashed lines: localization of the labelled neurons (see Table S1), unless otherwise indicated. Scale bars are 20 µm. D) Dashed line: estimation of the ventral border of the Vsx domain. F) Cells in the Vsx domain are circled in green when they co-express the indicated markers, and circled in red when they do not. This shows that no cell expresses *ey*, *kn* and *D* in the Vsx domain, except maybe cells indicated by an arrowhead (these are uncertain because they are located at the limit of the Vsx domain). The white dotted lines correspond to the insets placed at the bottom, which are rotated 90 degrees clockwise to the original image. I) The dashed white line indicates cells in the Vsx domain, while the solid white line indicates cells that might not be part of the Vsx domain. J) The arrowhead indicates Dll+ Vsx1+ cells in the Vsx, dOptix and what is likely the dDpp domains.

**Supplementary figure 4:**
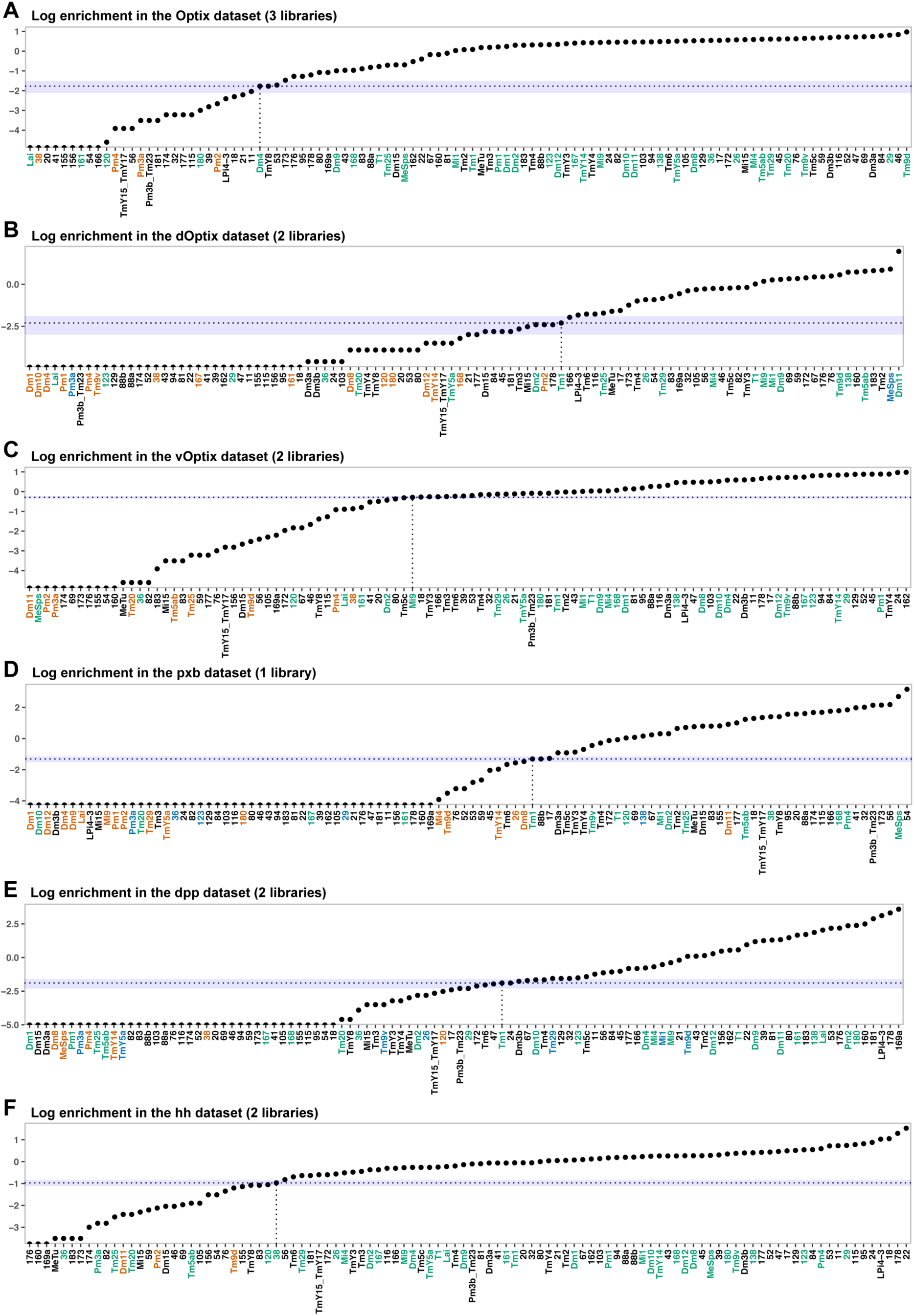
Thresholds used to binarize the enrichment scores of main OPC neurons. A-F) Log-enrichment scores of the indicated Classes (x-axis) in each dataset. Enrichment scores were calculated by dividing the normalized abundances in the indicated datasets, by the normalized abundances in the Reference dataset. Vertical dotted lines: enrichment of the Class used as a threshold (represented by horizontal dotted lines) to binarize the enrichment scores. Any class above this dotted line was considered as produced from the main OPC spatial domain corresponding to the dataset. Blue horizontal shaded regions: enrichment values close to the binarization threshold (Methods). The name of each neuronal Class is colored according to our validation experiments (Table S1). Red: neuronal types that should not be produced from the main OPC spatial domain corresponding to the indicated dataset, Green: neuronal types that could be produced from the main OPC spatial domain corresponding to the indicated dataset, Blue: neuronal types for which the presence in the corresponding main OPC domains was unclear.

**Supplementary figure 5:**
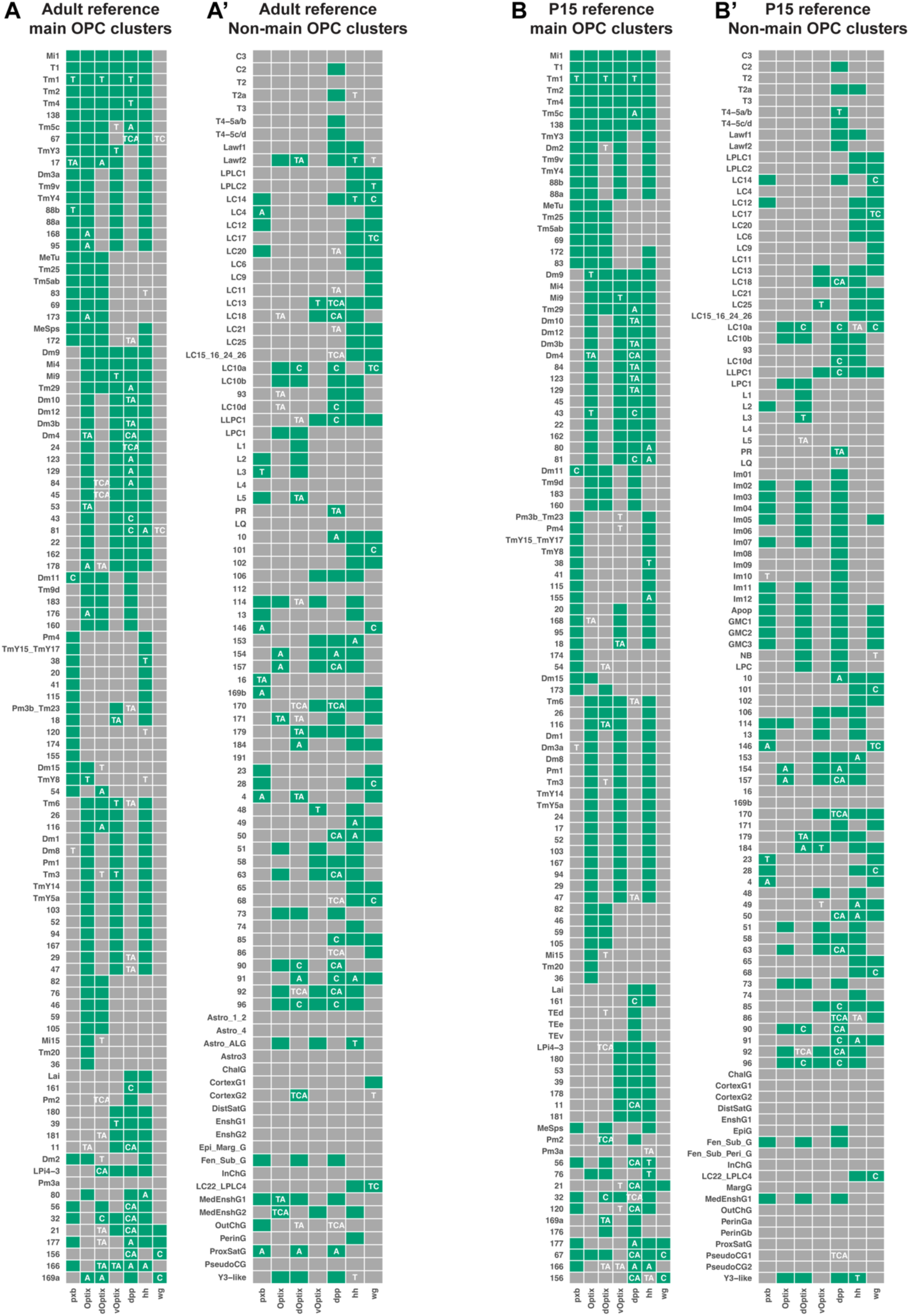
Binarized enrichment scores of the Classes in each dataset, and corresponding quality control metrics. A-B’) Green squares: Classes (y-axis) considered as enriched in the corresponding dataset (x-axis) according to the binarization thresholds indicated in Fig.S4 (adult reference dataset) or Annex 1 (P15 reference dataset). P15 was used as reference dataset in addition to the adult stage because TE neurons die before adulthood. Grey squares: Classes considered as depleted in the corresponding dataset. A = at least one of the libraries of the dataset had low abundance, *i.e.,* contained fewer than 3 cells of this Class, C = at least 80% of the cells of the Class were annotated with a confidence score strictly below 0.5, T = the enrichment of the Class was close to the binarization threshold (horizontal blue shaded region in Fig.S4). Abbreviations used as Class names are explained in Table S1 (the Class names ending with a capital G are glial cells).

**Supplementary figure 6:**
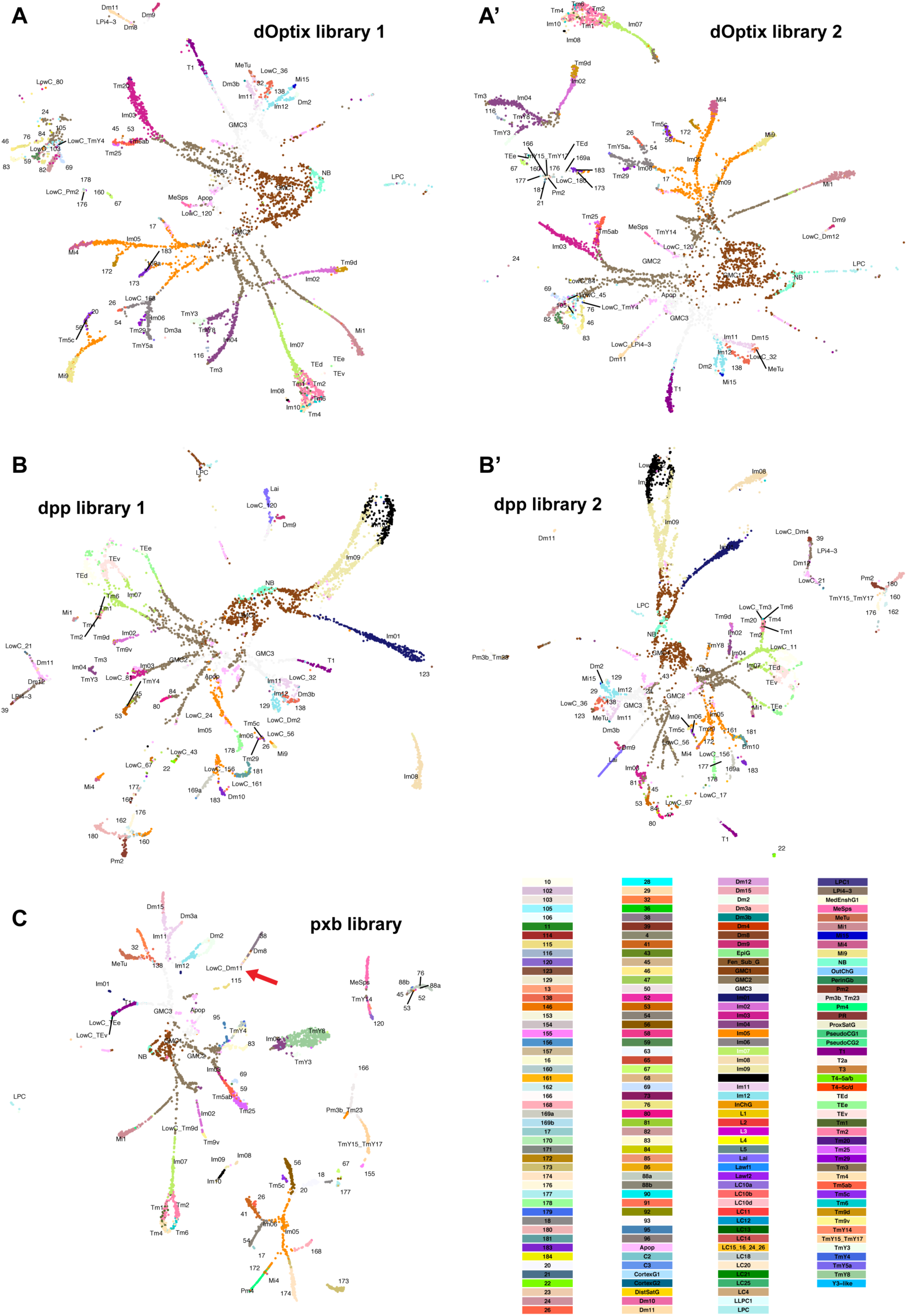
UMAP visualizations of the libraries obtained at early pupal stages. A-C) UMAP plots obtained using 150 principal components. The cells are labeled and colored according to the key presented on the bottom right. C) Red arrow highlights Dm11 (see main text).

**Supplementary figure 7:**
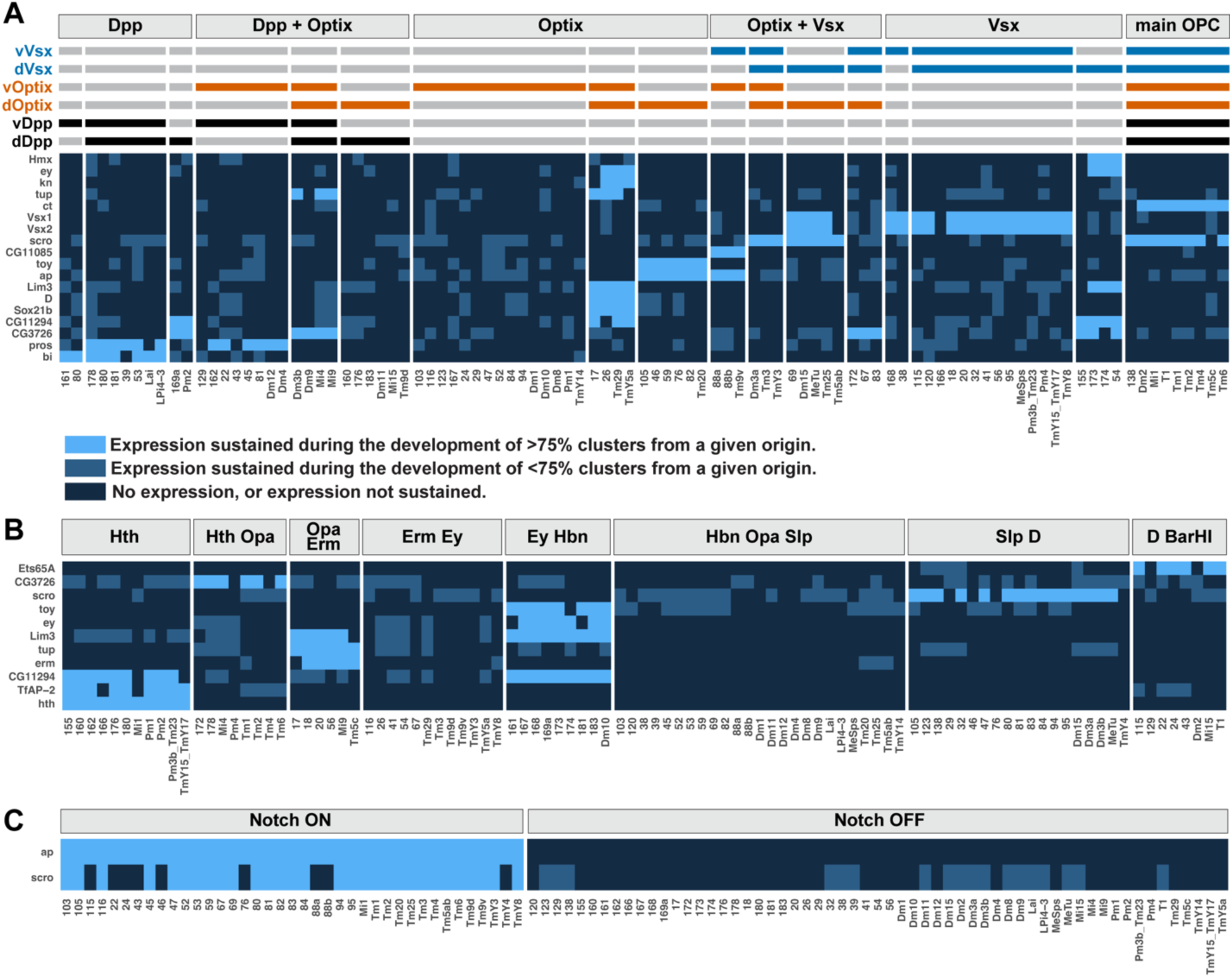
Terminal selectors continuously expressed in all neurons sharing a specification module. A-C) Terminal selectors (y-axis) continuously expressed in more than 75% of main OPC clusters (x-axis) from at least one spatial, temporal or Notch specification module. We used a threshold of 75% of clusters expressing continuously a TS instead of 100% to account for the imperfect resolution of our determination of developmental origins. Light blue: more than 75% of clusters from a given specification module express the gene throughout development, Medium blue: fewer than 75% of clusters from a given specification module express the gene throughout development, Dark blue: gene not expressed throughout development (Methods).

**Supplementary figure 8:**
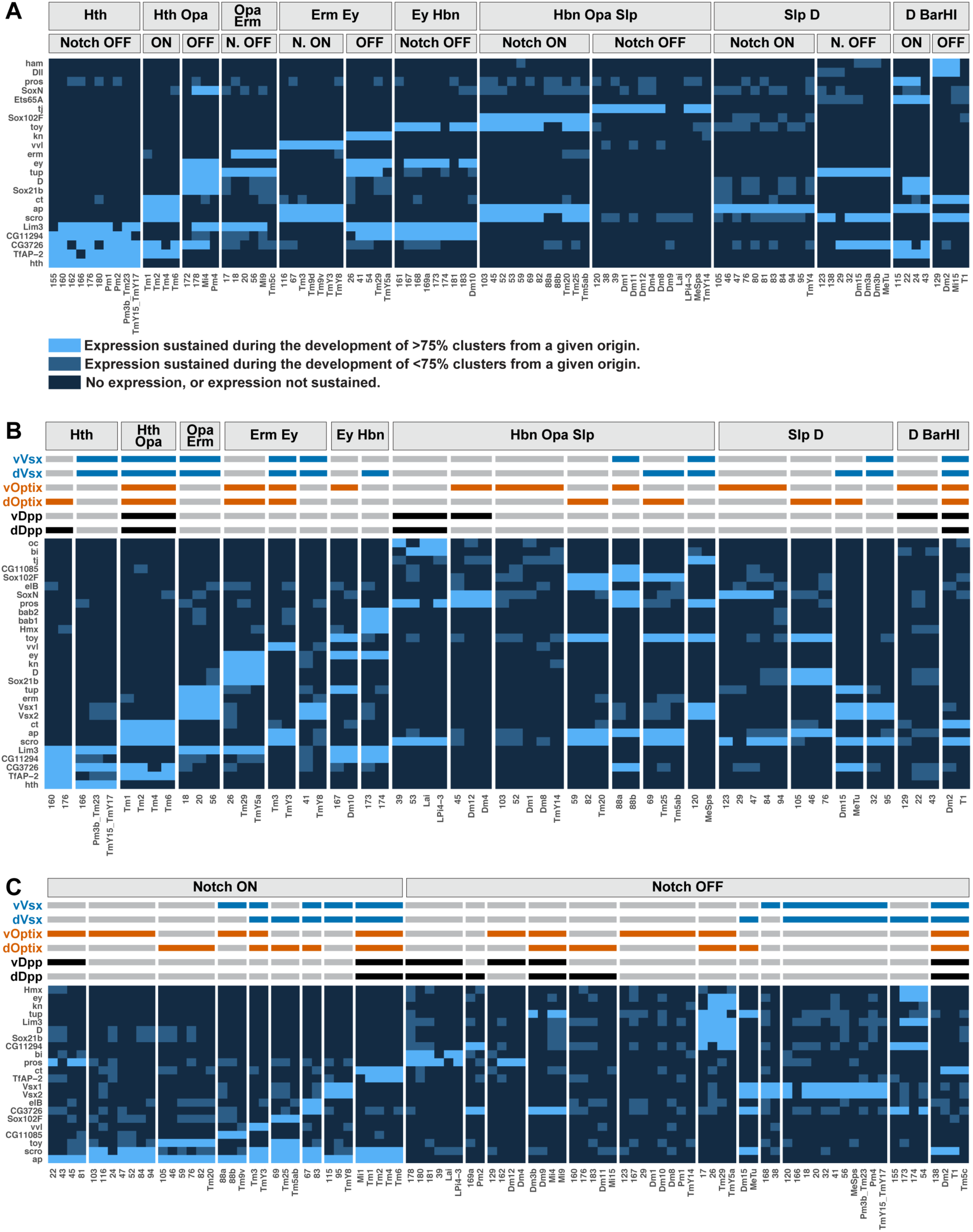
Terminal selectors continuously expressed in all neurons sharing a combination of two specification modules. A-C) Terminal selectors (y-axis) continuously expressed in more than 75% of main OPC clusters (x-axis) from at least one combination of two specification modules. We used a threshold of 75% of clusters expressing continuously a TS instead of 100% to account for the imperfect resolution of our determination of developmental origins. Only combinations of two specification modules containing more than 1 cluster have been plotted. Colors as in Fig.S7.

**Supplementary figure 9:**
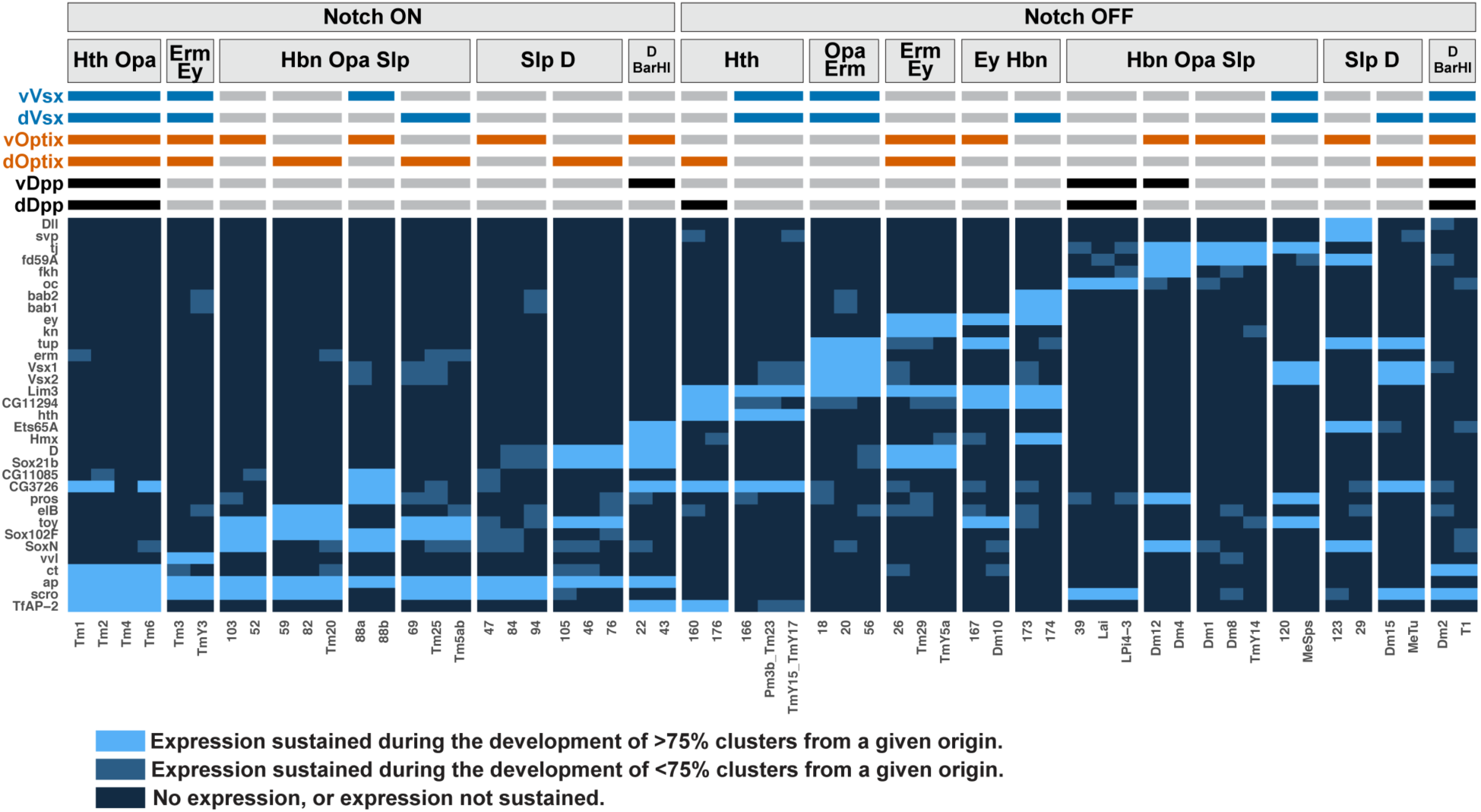
Terminal selectors continuously expressed in all neurons sharing a combination of three specification modules. Terminal selectors (y-axis) continuously expressed in more than 75% of main OPC clusters (x-axis) from at least one combination of spatial, temporal and Notch origins. We used a threshold of 75% of clusters expressing continuously a TS instead of 100% to account for the imperfect resolution of our determination of developmental origins. Only combinations of three specification modules containing more than 1 cluster have been plotted. Colors as in Fig.S7.

**Supplementary figure 10:**
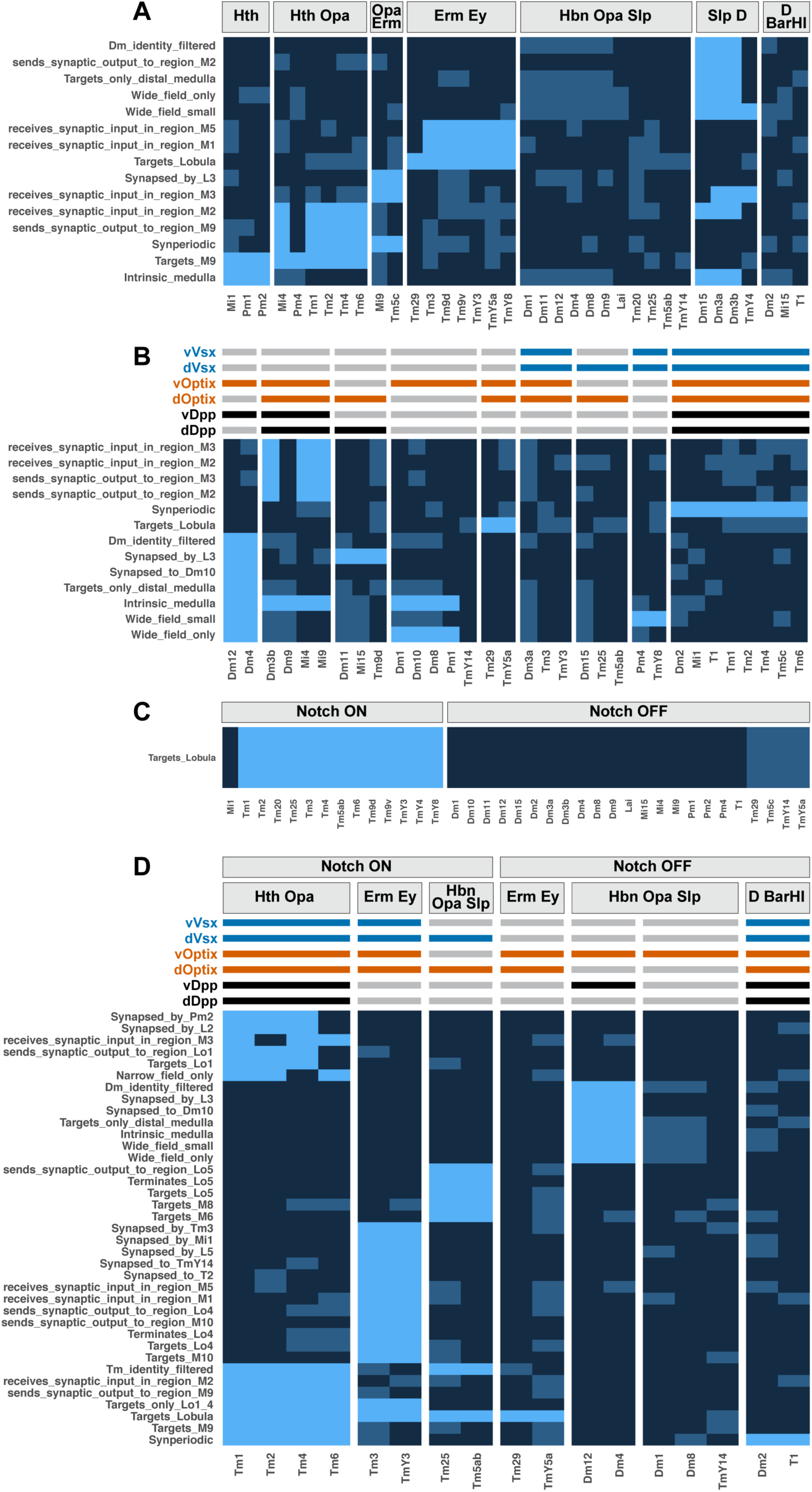
Adult morphological features present in all neurons sharing various combinations of specification modules. Adult morphological features (y-axis) present in more than 75% of main OPC clusters (x-axis) from at least one of the following: Temporal (A), Spatial (B) or Notch (C) origin, or from at least one combination of all three developmental origins (D). We used a threshold of 75% of clusters presenting a morphological feature instead of 100% to account for the imperfect resolution of our determination of developmental origins. The names of the morphological features are explained in Table S2. Only combinations of specification modules containing more than 1 cluster have been plotted. Light blue: more than 75% of clusters from a given origin present the morphological feature. Medium blue: fewer than 75% of clusters from a given origin present the morphological feature. Dark blue: lack of the morphological feature. These results could change when more clusters are annotated.

**Supplementary figure 11:**
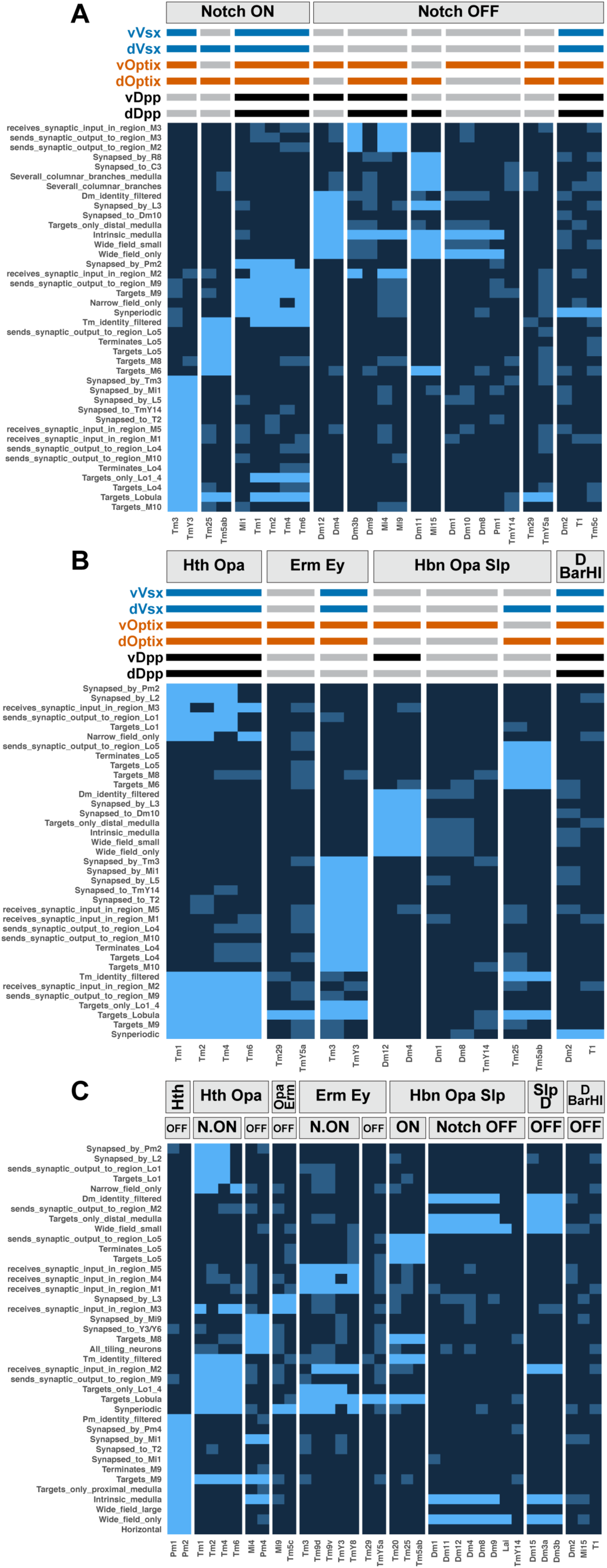
Adult morphological features present in all neurons sharing combinations of two specification modules. Adult morphological features (y-axis) present in more than 75% of main OPC clusters (x-axis) from at least one combination of two developmental origins. We used a threshold of 75% of clusters presenting a morphological feature instead of 100% to account for the imperfect resolution of our determination of developmental origins. The names of the morphological features are explained in Table S2. Only combinations of specification modules containing more than 1 cluster have been plotted. Colors as in Fig.S10. These results could change when more clusters are annotated.

**Supplementary figure 12:**
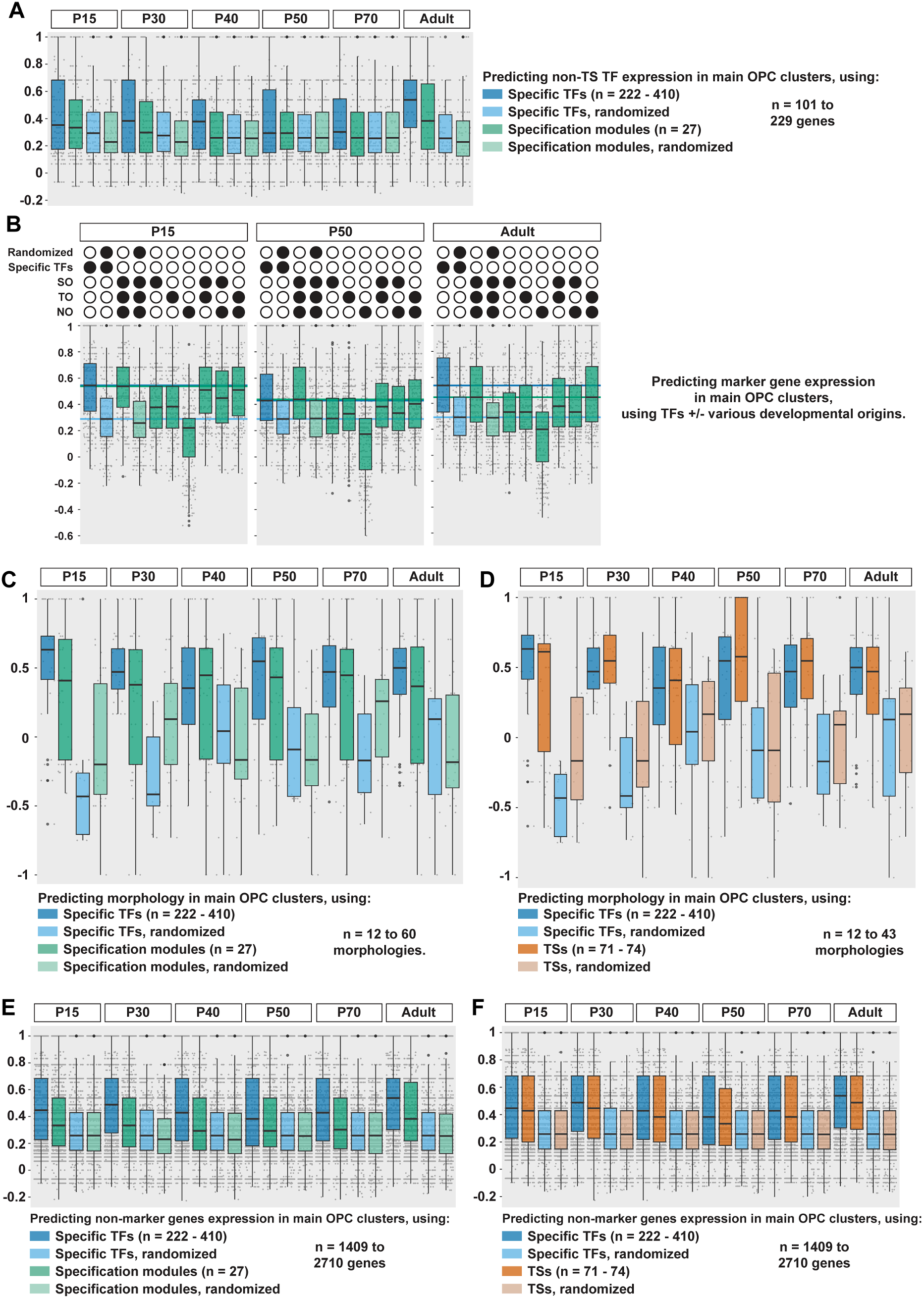
Specification modules and terminal selectors encode similar information. A-F) Matthews Correlation Coefficient between observed and predicted presence of a neuronal feature. The type of neuronal feature predicted and the type of predictor used are indicated on each plot. The specific TFs are the TFs differentially expressed between neuronal types. See Fig.5 for boxplots legend. NO = Notch Origin, SO = Spatial Origin, TO = Temporal Origin. B) Dark blue line: median of the MCC obtained when using specific TF expression as predictors, Light blue line: median of the MCC obtained when using randomized specific TFs expression as predictors, Green line: median of the MCC obtained when using developmental origin (SO+TO+NO) as predictors.

**Supplementary figure 13:**
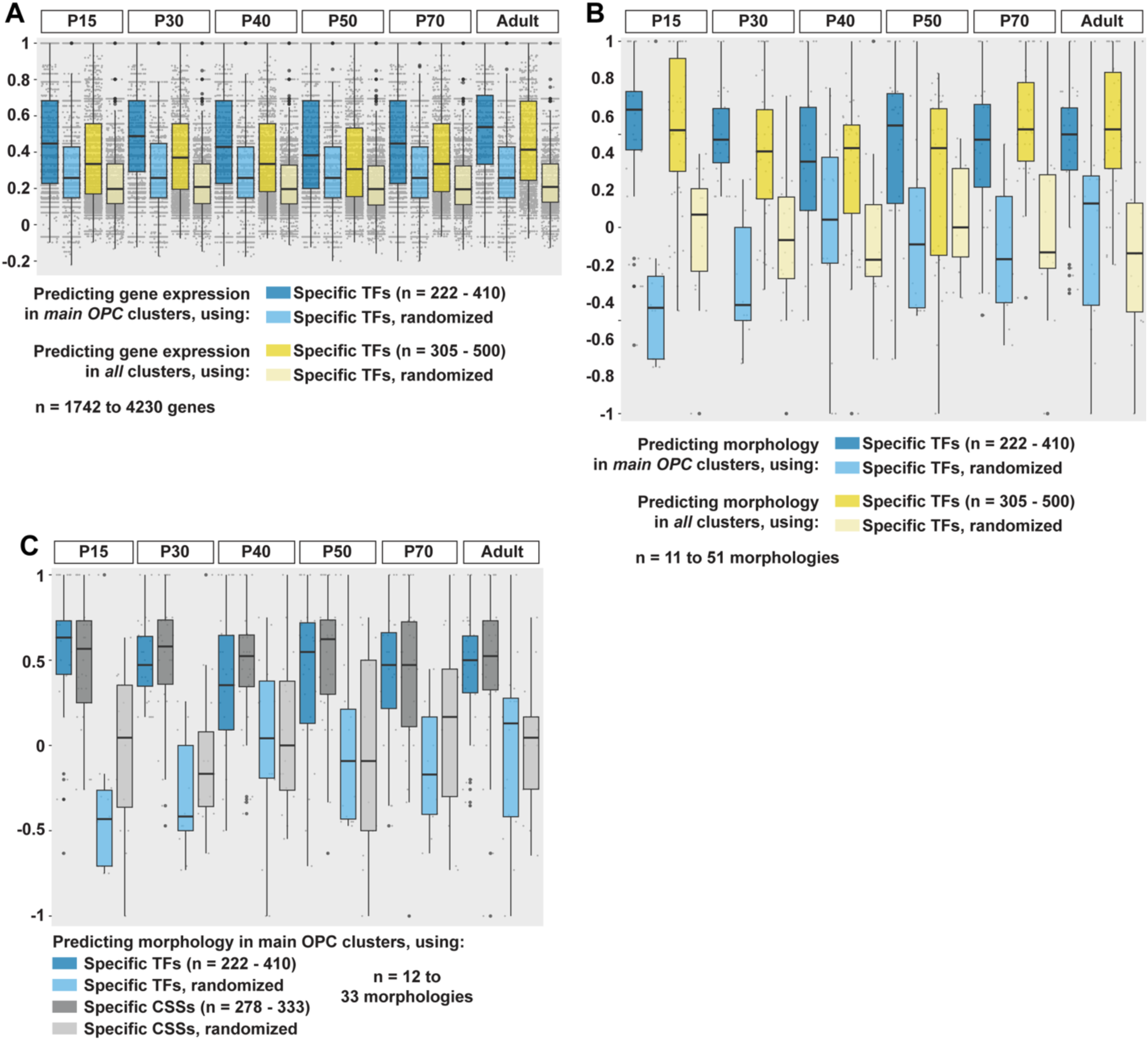
Prediction of neuronal features using TF expression or CSS expression, in main OPC or in all neuronal clusters. A-C) Matthews Correlation Coefficient between observed and predicted presence of a neuronal feature. The type of neuronal feature predicted and the type of predictor used are indicated on each plot. The specific TFs are the TFs differentially expressed between neuronal types. See Fig.5 for boxplots legend.

**Supplementary figure 14:**
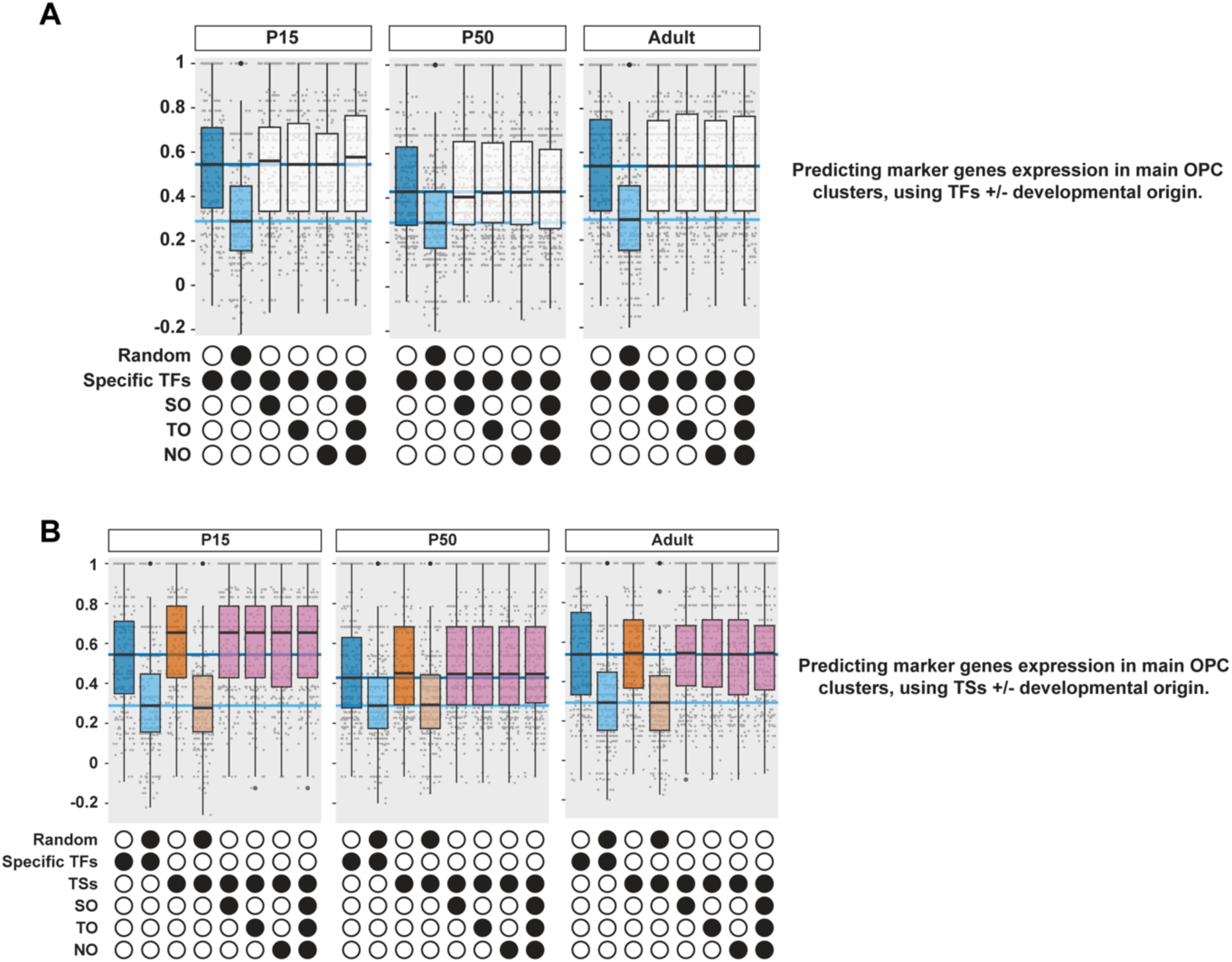
Specification modules are not remembered epigenetically after TS combinations are established. A, B) Matthews Correlation Coefficient between observed and predicted presence of a neuronal feature. The type of neuronal feature predicted and the type of predictor used are indicated on each plot. The specific TFs are the TFs differentially expressed between neuronal types. See Fig.5 for boxplots legend. SO = Spatial Origin, TO = Temporal Origin, NO = Notch Origin. Dark blue line: median of the MCC obtained when using specific TFs expression as predictors, Light blue line: median of the MCC obtained when using randomized specific TFs expression as predictors.

## Methods

### Immunohistochemistry

Whole brains were dissected in Schneider’s media (Sigma S0146), transferred to Schneider’s media on ice for no more than 30 min, and fixed using 4% paraformaldehyde (Electron Microscopy Sciences 15710) diluted in PBS 1X, for 20 min at room temperature. They were then rinsed 3 times in 0.5% Triton™ X-100 (Sigma) diluted in PBS 1X (PBTx), washed for 30 min in PBTx, and incubated at least 30 min in PBTx with 5% horse (ThermoFisher 26050070) or goat (ThermoFisher 16210064) serum (PBTx-block) at room temperature. They were then incubated at 4°C for 1-2 overnights in primary antibodies diluted in PBTx-block. They were then rinsed 3 times in PBTx, incubated two times for 30 min in PBTx, and incubated at 4°C overnight in secondary antibodies diluted in PBTx-block. Lastly, they were rinsed 3 times in PBTx, incubated two times for 30 min in PBTx, and mounted in SlowFade™ (ThermoFisher S36936) before imaging with a Leica SP8 confocal microscope using a 63x glycerol objective. Images were processed in Fiji and Adobe Illustrator.

The following primary antibodies were used: mouse anti-Svp (1:20) (DSHB), mouse anti-tup (1:100) (DSHB), mouse anti-Dac 2-3 (1:40) (DSHB), mouse anti-Cut (1:10) (DSHB), rabbit anti-Dichaete (1:500) (modENCODE), rabbit anti-toy (1:500)^9^, rabbit anti-Bsh (1:1800)^9^, rat anti-vvl (1:50)^32^, guinea pig anti-Vsx1 (1:100)^15^, guinea pig anti-Hth (1:500) (gift from Richard Mann), guinea pig anti-TfAP-2 (1:100)^32^, rat anti-drgx (1:250)^15^, rat anti-erm (1:100)^32^, guinea pig anti-run (1:500)^32^, rat anti-Ey (1:100) (this study), guinea pig anti-Kn (1:100)^32^, rabbit anti-SoxN (1:250)^15^, guinea pig anti-Tj (1:250) (gift from Dorothea Godt^41^), rabbit anti-Dll (1:100)^32^, chicken anti-GFP (1:200) (Millipore Sigma), rat anti-Ecad (1:200) (DSHB).

The following secondary antibodies, all from donkey, were used: anti-rat DyLight 405 (1:100), anti-rabbit DyLight 405 (1:100), anti-guinea pig DyLight 405 (1:100), anti-mouse Alexa Fluor 488 (1:400), anti-mouse Alexa Fluor 555 (1:400), anti-guinea pig Cy3 (1:400), anti-rabbit Alexa Fluor 555 (1:400), anti-rat Alexa Fluor 647 (1:200), anti-rabbit Alexa Fluor 647 (1:200), and anti-guinea pig Alexa Fluor 647 (1:200).

### Antibody generation

A rat polyclonal antibody against Ey was generated by Genscript using the following epitope (https://www.genscript.com/):

MFTLQPTPTAIGTVVPPWSAGTLIERLPSLEDMAHKDNVIAMRNLPCLGTAGGSGLGGIAGKPSPTMEAVEASTASHPHSTSSYFATTYYHLTDDECHSGVNQLGGVFVGGRPLPDSTRQKIVELAHSGARPCDISRILQVSNGCVSKILGRYYETGSIRPRAIGGSKPRVATAEVVSKISQYKRECPSIFAWEIRDRLLQENVCTNDNIPSVSSINRVLRNLAAQKEQQSTGSGSSSTSAGNSISAKVSVSIGGNVSNVASGSRGTLSSSTDLMQTATPLNSSESGGASNSGEGSEQEAIYEKLRLLNTQHAAGPGPLEPARAAPLVGQSPNHLGTRSSHPQLVHGNHQALQQHQQQSWPPRHYSGSWYPTSLSEIPISSAPNIASVTAYASGPSLAHSLSPPNDIESLASIGHQRNCPVATEDIHLKKELDGHQSDETGSGEGENSNGGASNIGNTEDDQARLILKRKLQRNRTSFTNDQIDSLEKEFERTHYPDVFA

### Determination of the spatial origin of main OPC clusters

#### Obtaining the animals for scRNA-seq and testing GFP expression

The genotypes of the *Drosophila* lines used were:

- Optix-ts-MC line: *20xUAS-FlpG5::PEST; Optix-Gal4,tub-Gal80ts/CyO; ubi>STOP>Stinger/TM6*
- vOptix-MC line: *w*,disco-T2A-VP16AD; optix-T2A-Gal4DBD/+; UAS-Flp.D/UAS-Flp.Exel,UAS-RedStinger,Ubi>Stop>Stinger(G-Trace)*
- dOptix>Gal4: *w1118; salr-T2A-VP16AD,Optix-T2A-Gal4DBD/CyO*
- hh-ts-MC line: *20xUAS-FlpG5::PEST; ubi>STOP>Stinger/CyO; hh-Gal4,tub-Gal80ts/TM6B,Tb*
- pxb-MC line: *20xUAS-FlpG5::PEST; ubi>STOP>Stinger/CyO-nlsGFP; pxb-T2A-Gal4/TM6B,Tb*
- dpp>GFP line: *w; (UAS-Stinger; dpp-Gal4)/[SM6a:TM6B, Tb, actGFP]*

For the Optix dataset, Canton S virgins crossed with Optix-ts-MC males were allowed to lay eggs for 2 hours at 18°C. The eggs were then placed at 29°C for 2 days and 9 hours (late L2 stage), then back at 18°C for the remainder of their development. GFP expression was tested in L3 wandering larvae and sequencing was performed in adults (Fig.2B). We produced 3 libraries from the same cell suspension of optic lobes from 5 female flies that were a few hours old.

For the vOptix dataset, vOptix-MC flies were kept at 25°C. GFP expression was tested in L3 wandering larvae and sequencing performed in adults (Fig.2B). We produced 2 libraries from the same cell suspension of the optic lobes of 15 female flies that were more than two-week-old.

For the dOptix dataset, *w;; UAS-Stinger* virgins were crossed with dOptix>Gal4 males and kept at 25°C. GFP expression was tested, and sequencing was performed, in P0-P12.5 pupae (Fig.2B). We produced 2 libraries from the same cell suspension of 24 optic lobes.

For the hh dataset, Canton S virgins crossed with hh-ts-MC males were allowed to lay eggs for 2 hours at 18°C. The eggs were then placed at 29°C for 2 days and 3 to 9 hours (mid to late L2 stage), then back at 18°C for the reminder of their development. GFP expression was tested in L3 wandering larvae and sequencing was performed in adults (Fig.2B). We produced simultaneously 2 libraries from independently produced cell suspensions, each from the optic lobes of 7 two-week-old female flies.

For the pxb dataset, Canton S virgins were crossed with pxb-MC males and kept at 25°C. GFP expression was tested, and sequencing was performed, in P0-P12.5 pupae (Fig.2B). We produced 1 library from 8 optic lobes.

For the dpp dataset, Canton S virgins were crossed with dpp>GFP males and kept at 25°C. GFP expression was tested, and sequencing was performed, in P0-P12.5 pupae (Fig.2B). We produced simultaneously 2 libraries from independently produced cell suspensions, each from 12 optic lobes of young pupae.

All animals used for scRNA-seq were selected for the lack of balancer chromosomes. P0-P12.5 animals were obtained by selecting pupae in which the head had not yet detached from the pupal case^42^.

For the wg dataset, the sequencing of the wg-Gal4 line is described in El-Danaf et al. 2024^21^.

#### Sample preparation and single-cell mRNA-sequencing

We followed a similar cell dissociation protocol to generate all spatial domain datasets. Whole brains were extracted in ice-cold Schneider’s media. The brains were then transferred into DPBS 1X without calcium and magnesium (Corning/Fisher 21-031-CV). We used an epifluorescence microscope to remove brains with ectopic main OPC GFP expression (see main text), and placed the brains back in Schneider’s media. The brains were then incubated at 25°C in Schneider’s media with 2 mg/mL Collagenase (Sigma C0130) and 2 mg/mL Dispase (Sigma D4693-1G), for 1 hr 30 min (adult brains) or for 30 min (pupal brains), then washed 3 times with ice-cold Schneider’s media to stop the dissociation. Optic lobes were then separated from the central brain, transferred to ice-cold DPBS + 0.04% BSA (ThermoFisher AM2618), and washed twice with ice-cold DPBS + 0.04% BSA. For pupal brains, the separation between central brain and optic lobes can be seen as a difference in texture: the optic lobe forms a smoother and cohesive structure that can be removed using a fine tungsten needle. Optic lobes and 150 μL of DPBS + 0.04% BSA were then transferred to a Eppendorf™ DNA LoBind™ Tubes (Fisher Scientific 13-698-791), and dissociated by pipetting vigorously 50 times, resting the cells for 1 minute on ice, and pipetting 50 times more. The cell suspension was then checked under a dissecting microscope, and we pipetted up to 50 more times if the size of remaining clumps was too big. We did not try to dissociate small clumps since they are most likely neuropils, not somas, and indeed we empirically noticed a better quality of the data obtained after lowering pipetting repetitions. The suspension was then passed through a 20 µm strainer (pluriSelect 43-10020-40), and the filter was washed with 400 μL of DPBS + 0.04% BSA which we also collected (pipetting from the opposite side may be necessary to collect the entire cell suspension).

Stinger(nuclear GFP)-positive cells were sorted with a FACSAria II using FACSDiva 8.0.2 to set the gates shown in Annex 1, and we loaded the single-cell sequencing chip with 43.5 μL of the cell suspension provided by the FACS, undiluted, mixed with the master mix. Moreover, we used a tip coated with 70 μL of DPBS + 0.04% BSA and 0.02% Tween-20 (Bio-Rad 1662404) to sample the cell suspension, and kept the same tip to load the chip, which we empirically noticed drastically reduces the loss of cells. All steps of single-cell sequencing were performed following the Chromium Next GEM Single Cell 3ʹ Reagent Kits v3.1 (Dual Index) instructions.

Sequencing was performed using Illumina NextSeq 500 or Illumina NovaSeq 6000 at the Genomics Core at the Center for Genomics and Systems Biology at New York University.

#### Supervised annotation of the scRNA-seq datasets

The libraries were analyzed using Seurat 4.3.0. For each library, a Seurat Object was created with all genes expressed at least in 3 cells, and all cells expressing at least 200 genes. The objects were then filtered by keeping all cells below a certain percentage of mitochondrial genes, below a certain number of UMIs, and above a certain number of genes, chosen according to the distribution of these parameters (Annex 1). The thresholds chosen were identical for all libraries acquired at a given day with flies of a given genotype, but were different otherwise. For the Optix libraries the thresholds were 7/17000/800 (percent of mitochondrial genes, number of UMIs, number of genes), for the vOptix libraries 10/10000/500, for the dOptix libraries 5/20000/1000, for the hh libraries 10/20000/700, for the dpp libraries 5/20000/900, for the pxb library 5/30000/1300, for the wg libraries 10/25000/1200. After filtering, we obtained an Optix dataset of 19,036 cells (3 libraries), a hh dataset of 9,182 cells (2 libraries), a vOptix dataset of 10,596 cells (2 libraries), a dOptix dataset of 12,302 cells (2 libraries), a pxb dataset of 4,988 cells (1 library), a dpp dataset of 12,862 cells (2 libraries), and a wg dataset of 12,974 cells (2 libraries). For each library we then ran NormalizeData, FindVariableFeatures, and ScaleData with default parameters, as well as RunPCA, RunTSNE, and RunUMAP with default parameters and a dimensionality of 150. Dimensionality reductions were run purely for visualization purposes; they were not used to perform any clustering. Instead, we used the normalized expression of marker genes and our neural network classifier^11^ to assign each cell of each library from this study to its corresponding cluster in our previously published scRNA-seq atlas^11^ (metadata fields “NN_Cluster_Number”), and to give this assignment a confidence score between 0 and 1 (metadata field “Confidence_NN_Cluster_Number”). Notably, since we know which clusters are produced in the main OPC^28^, this allowed us to identify non-main OPC cells present in our datasets, including the ones due to expression of our reporters outside of the main OPC. For all figures, we replaced cluster numbers by cluster annotations (metadata fields “Annotation”) using Table S3. Lastly, if at least 80% of the cells of a given Class were annotated with a confidence score strictly below 0.5, the Class was flagged as “low confidence” by assigning it the value “1” in the metadata field “cells_flagged”, and its annotation was changed from “X” to “LowC_X”.

#### Binarization of the spatial origin of each main OPC cluster

We then normalized the abundance of each neuronal Class in each dataset. Indeed, a neuron produced exclusively in the Vsx domain is expected to be relatively more abundant in a library produced from sorted pxb cells compared to our published atlas, which contains neurons produced in all domains. However, a neuron produced in each domain equally should have a similar abundance in all datasets (Figure S2B). To allow comparisons of Class abundances between libraries, we used the abundances of neurons produced in the whole main OPC, since they represent a constant between main OPC domains. In each library, we therefore divided the abundance of each Class by the abundance of all the Classes corresponding to neuronal clusters produced in all the main OPC. We also normalized the abundance of each Class in our single cell atlas but averaged the normalized abundance across all libraries to simplify the comparisons. We performed a first round of normalization using Mi1, Tm1 and T1. We then used the results of this round of normalization to determine that T1, Mi1, Tm1, Tm2, Tm4, and cluster 138 were produced in all main OPC domains and therefore used them for a second round of normalization. Because this second round of normalization did not show additional neurons produced in the whole main OPC, we did not perform further rounds of normalization. However, these clusters are not produced in the wg domain, so they could not be used to normalize wg libraries. We therefore only normalized this dataset by dividing the abundance of each Class by the abundance of its most abundant cluster.

However, normalized abundances are not enough to decide whether a neuronal type is produced in a given domain, because rare cell types would still represent only a small part of any dataset (for instance cluster 38 is produced in the vVsx domain, which represents only half of the Vsx domain and thus the pxb dataset). We therefore computed an enrichment score for each Class in each spatial domain (Fig.S2B). To do so, we divided the average normalized abundance of a Class in all libraries of a spatial domain dataset by its average normalized abundance in our single cell atlas. Theoretically, any score above 1 should indicate an enrichment of the Class in a main OPC domain compared to the whole main OPC, and therefore indicate that this cluster is produced in this domain. However, for a given Class, the minimal and maximal normalized abundances across libraries of a same main OPC domain are usually variable (see Annex 1), probably due to technical reasons (dissociation, FACS…), and therefore enrichment scores could not be analyzed in such a straightforward manner.

Therefore, we then binarized these “enrichment scores” by choosing thresholds based on our experimental validations of spatial origin (Fig.S4). For the Optix dataset, we chose Dm4 as a threshold because this is the least abundant neuronal type that we ascertained to come from this domain. For the dOptix dataset, we chose Tm1 as a threshold instead of TmY5a or Tm20 (that we know are produced in the dOptix domain), because some of these might have been annotated as immature neurons (with which they form a trajectory on UMAP visualization, Fig.S6). For the vOptix dataset, we chose Mi9 as a threshold instead of cluster 161, because cluster 161 is a very small cluster that is unlikely to be produced in all of the vVsx, vOptix and vDpp domains (which is where we found the expression of the markers for cluster 161, 167, 168, and Dm10, Table S1). For the pxb and dpp datasets, we chose Tm1 as a threshold because out of all the neuronal types we experimentally validated as coming from these domains, Tm1 was the least abundant in both the pxb and dpp datasets. For the hh dataset, we chose cluster 38 as a threshold because we know it is produced exclusively in the ventral OPC. We did not choose Tm5ab because 1) this cluster contains two cell types, and 2) their markers (shared with other clusters) are found in the d/vVsx, dOptix and dDpp: the ventral main OPC represent only a small part of their potential domain of production.

We performed all these analyses using both our scRNA-seq atlas at the P15 and adult stages as a reference, because some clusters (for instance the TE cells) are only found at one of the two stages. However, we favored results obtained with the adult dataset, because it contains more cells, and at this stage their identities are more defined (there are no more immature neurons). Moreover, we performed these analyses for both main OPC and non-main OPC clusters but displayed these clusters separately in our plots. The presence of non-main OPC neurons in our datasets can mean that 1) the neuron is produced outside the main OPC but still expressed our reporters, 2) the neuron is in fact produced in the main OPC and was mistakenly assigned as non-main OPC neurons^28^.

#### Final assignment of a spatial origin to each main OPC cluster

We primarily used binarized enrichment to assign an origin to each Class. However, we deemed the binarized enrichment of a Class in a given dataset as less trustworthy if a class was flagged as being annotated with low confidence, and/or there were 3 cells or fewer of this class found in the dataset, and/or its enrichment value was +/- 0.05 compared to the binarization threshold (Fig.S5). In this case, we also considered additional parameters (see Table S1), such as its localization on the UMAP (Fig.S6). For instance, if “ClassA” contains only a few cells annotated with low confidence, and its cells are mixed on the UMAP with cells from “ClassB” that are annotated with high confidence, it is likely that ClassA cells are in fact ClassB cells misannotated by the neural network. We also accounted for the expression of vOptix-MC in the vVsx stripe and in the tips of the OPC: if a neuronal type is produced only in the Dpp domain, for instance, it should be absent in the Optix dataset but present in the vOptix dataset. Similar reasoning applies to a neuron produced in vVsx. Our choices are explained in Table S1. Lastly, it should be noted that in the absence of dVsx and dDpp datasets, assignment to these domains could only be extrapolated. For instance, if a Class was found in the dpp and the hh datasets, it is clearly produced in vDpp, and maybe in dDpp. However, if the Class was also found in the vOptix and dOptix datasets, then the corresponding neuron is very likely to be also produced in the dDpp domain. Such determinations are also explained in Table S1.

#### Identifying clusters corresponding to synperiodic neurons

Synperiodic neurons are the most abundant neurons in the optic lobe: they are present in ∼800 copies, *i.e.*, one per column. The clusters corresponding to synperiodic neurons should therefore be the largest ones in our scRNA-seq atlas^11^. Based on electron microscopy data^10^, among the clusters containing more than 1,000 cells in our adult scRNA-seq dataset, most of them clearly correspond to synperiodic neurons: Tm3 (1037 cells. Tm3 was previously described as ultra-periodic, *i.e.*, being present in more than one copy per column^7^), T1 (892 cells), Tm1 (890 cells), Mi4 (889 cells), Mi9 (889 cells), Mi1 (887 cells), Tm9 (887 cells), Tm2 (883 cells), Tm20 (876 cells), Tm4 (833 cells), Tm6 (770 cells), Dm2 (726 cells). A few correspond to neurons of intermediate abundance: TmY5a (678 cells), Mi15 (582 cells), Dm8 (550 cells), TmY3 (409 cells), Dm10 (311 cells). The status of Dm3 is unclear, as our dataset contains only two corresponding clusters, Dm3a and Dm3b, although three abundant Dm3 subtypes have now been described (Dm3a, 601 cells, Dm3b, 571 cells, Dm3c, 401 cells). The list of neurons with quantified abundance did not include TmY8 and Tm25^10^, and two of our clusters containing more than 1,000 cells are unannotated (clusters 82 and 36).

### Producing the table of morphological features

To produce the table, we surveyed the Virtual Fly Brain website (https://v2.virtualflybrain.org/) as well as 40 publications (Table S2). The definition of each feature is indicated in Table S2, and the confidence in the values are color coded as explained in the “legend” tab. One important characteristic of the table is that, for each feature, we distinguished between known absence in a cluster, or whether its presence was simply not evaluated, which we denoted by “NA” (in which case the cluster was removed when building machine learning models for this feature). Moreover, the control of neuronal morphology by TF or CSS expression is likely to be context specific. For instance, a CSS might be important to target lobula layer 4 in some neurons but have a completely different role in a neuron that does not enter the lobula. Therefore, some of the molecular features in Table S2 are duplicated: in one case they are evaluated in all neuronal types, and in a second case (identified by adding “filtered” to the name of the feature) neurons that cannot express the feature (ex: targeting of lobula layer 4 for non-lobula neurons) are flagged with “NA”, for “non-applicable”, which also allowed us to remove them when building random forest models for these features.

Among the morphological features, layer targeting was one of the least straightforward to assess because the available information was often conflicted between different sources. This can be due to differences in layer definitions between different authors (for instance differences in the definition of the 4^th^ medulla layer^1,43^), or different imaging techniques (for instance, Takemura and colleagues^7^ note that “In some cases […], the cells reconstructed from EM appear to have fewer processes than the light microscopy images. This relative sparsening results from difficulties in connecting the fine processes of the arborizations to the main body of the neuron during EM reconstruction.”). Moreover, the binarization of layer targeting data was difficult due to lack of defined threshold and inter-individual variation. For instance, Takemura and colleagues^7^ present two reconstructed Mi4 cells: one terminates in M9 while contacting the M10 layer border, and the other terminates in M10. One shows clear branching in M9, the other does not. And both show one arborization in M7, but these are much smaller than arborizations in other layers, and Mi4 is usually not reported to arborize in M7. We did our best to indicate low confidence in our assessment of targeting determinations by a color code (Table S2), and generally chose a non-conservative approach in which for any each neuronal type we indicated all layers in which it was shown to produce significant arborizations, even if only one report of such instance was known. In the future, using synapse coordinates instead of layer targeting would be a more rigorous approach.

For connectivity in the lamina, we used Table S2 of the study by Rivera-Alba and colleagues^5^, in which we removed all synapses that were considered as uncertain and grouped the cell types L4, L4+x and L4+y. We then converted the number of presynapses from cell type A to cell type B into a proportion of all presynapses from cell type A (number of presynapses made by A to B / total number of presynapses made by A). We did the same for the postsynapses (number of postsynapses made by A from B / total number of postsynapses made by A). For connectivity in the medulla, we utilized synaptic connections between neurons from the Fib25 dataset from neuPrint (GitHub Repository: connectome-neuprint/neuPrint; commit hash: 69fbda4). Briefly, we retrieved all annotated neurons that have any synapses, and we matched the annotated pre- and post-synapses with a confidence score to produce a table called “raw_connection_by_cell.csv” that contains all occurrences of a synapse between a given couple of pre and postsynaptic neurons. We kept only synapses with a perfect confidence score, and duplicated the results obtained for Tm9 into Tm9v and Tm9d (which assumes they connect to the same cell types). We then produced “Table_presynapses_to.csv” that contains the proportion of presynapses that a given cell type (ex: L2, column) makes with any other cell type (ex: Mi1, row), and “Table_postsynapses_by.csv” that contains the proportion of postsynapses that a given cell type (ex: L2, column) makes with any other cell type of the study (ex: Mi1, row). The code used for this re-analysis will be made available upon publication of the manuscript. Both for connectivity in the lamina and the medulla, we also produced “Synapsed_to_A” and “Synapsed_by_A” features by binarizing the proportions of synapses by neuronal type A, or to neuronal type A, with a threshold of 5%.

In addition to features related to the shape and the connectivity of individual neuronal types, we also added a few “identity” features. For instance, the feature “Dm_identity_filtered” is true for all clusters corresponding to a Dm neuronal type (Dm1, Dm2, etc.), false for all clusters corresponding to non-Dm neuronal types, and “NA” (*i.e.*, not used in machine learning) if the cluster is not annotated. Therefore, this feature was used to find good predictors of “being a Dm”. Similar identity features were done for other groups of neuronal types (Tms, Mis, etc.).

### Production and evaluation of the random forest models

#### Selection of the clusters used

For each stage, we performed the analyses using either “All clusters” or “main OPC clusters”. By “All clusters”, we mean all neuronal clusters (as indicated in Table S3), except a) clusters containing less than 5 cells, b) clusters that could contain more than 1 cell type (heterogeneous clusters indicated in Table S3), c) clusters containing features of low quality cells, multiplets or central brain neurons (38, 85, 112, 102 and 120, see Table S3), and d) clusters containing TE cells (220, 223, 224, and 233). We removed TE neurons because they are very different from other optic lobe neurons^11^, and might present unique regulatory relationships that would complicate model building. The removed heterogenous clusters are based on previous work^11^, and on results obtained by grouping our P50 optic lobe atlas^11^ with another optic lobe atlas^14^, produced at P48: this increase in cell number allowed splitting some of our previously published clusters. By “main OPC clusters”, we mean the subset of “All clusters” that were identified as being produced in the main OPC^28^ and for which we were able to assign a spatial, temporal and Notch origin (see Table S1).

#### Production of the gene expression tables

We filtered the scRNA-seq datasets of the optic lobe produced previously^11^ at P15, P30, P40, P50, P70 and adult stage to keep only “All clusters” or “main OPC clusters” as detailed above, and to keep only the subset of genes that we considered as unambiguously expressed, *i.e.,* with mRNA detected in at least 10% of the cells for at least 1 neuronal cluster. We then used the AverageExpression function of Seurat_4.3.0 to produce tables containing the log-normalized average gene expression of each cluster. We used averaged gene expression for our analyses rather than single-cell gene expression to mitigate the effect of dropouts (*i.e.,* false negative gene expression due to randomness in the capture of mRNA). We also produced additional tables in which, for each gene, we shuffled the values between clusters to produce log-normalized randomized average gene expression.

#### Binarization of gene expression

Several steps of our analyzes require binarization of gene expression. To do so, we used our previously published mixture modeling tables, which quantify the probability of expression for each gene in each cluster of our scRNA-seq atlas^11^. We binarized these expression probabilities using a threshold of 0.5, and we refer to these binarized tables as “MM0.5” hereafter.

#### Selection of the predictors

To select the predictor genes, we further filtered the log-normalized average gene expression tables for TFs, CSSs or TSs. For TFs, we used a list containing “TFs with characterized binding domains, computationally predicted (putative) TFs, chromatin-related proteins and transcriptional machinery components”^44^. For CSSs, we used a list of “*Drosophila* CSS proteins potentially involved in cell recognition” obtained by performing “BLAST searches with extracellular […] domain sequences from a variety of species and collated published information”^45^. Lastly, for TSs we used the list we previously published^17^. Moreover, for each list at each stage, we kept only the genes expressed at least once, and in fewer than 75% of either “All clusters” or “main OPC clusters”, according to MM0.5. Indeed, since our goal was to predict neuronal features only present in a subset of the clusters (differentially expressed genes, non-pan-neuronal morphologies) and to find candidate regulators for them, it made sense to focus on differentially expressed predictors. Moreover, pan-neuronal predictors will always be well correlated with any feature that is present is many neurons, even though they are not involved in its regulation. We produced the tables of predictors using either regular or randomized gene expression.

The developmental origins used as predictors are found in Table S1 for temporal and Notch origins. However, some clusters can be produced in more than one spatial domain. Therefore, for each cluster, we used Table S1 to assign a unique origin among vDpp, vdDpp, dDpp, vDpp.vOptix, vdDpp.vdOptix, dDpp.dOptix, vOptix, vdOptix, dOptix, vOptix.vVsx, vdOptix.vdVsx, dOptix.dVsx, vVsx, vdVsx, dVsx, vVsx.dVsx.dOptix, vOptix.vVsx.dVsx and whole_OPC. We also removed the few main OPC clusters for which a spatial origin could not be assigned. We shuffled which clusters are produced in each spatial, temporal and Notch origin to produce randomized tables of developmental origins.

Importanly, although vDpp and dDpp contain clusters produced exclusively in these domains, because of the lack of dataset for neurons from the dDpp domain in most cases we could not identify neuronal types produced in the vDpp but not dDpp. Therefore, some neurons in vdDpp might be in fact produced exclusively in the vDpp domain. Such reasoning also applies to the vVsx, dVsx and vdVsx origins.

#### Production of the training and testing sets for molecular features

To choose which clusters and genes to use for random forest predictions, we further filtered the log-normalized average gene expression tables, in “All clusters” and “main OPC clusters independently. Both training and testing sets should contain examples of positive and negative regulation of the feature of interest, and therefore all genes used to produce a random forest model should be expressed in at least 2 clusters. In addition, genes expressed in only 1 cluster are perfectly correlated with the combination of TSs specific to this cluster^11,17^, which are their best candidate regulators. Thus, we removed all genes expressed in 0 or 1 clusters according to MM0.5. We also removed all genes we considered pan-neuronal, *i.e.,* expressed in more than 75% of the clusters according to MM0.5, because these could be regulated by any pan-neuronal TFs and a random forest model would therefore not be informative of their regulation. For each gene, we then produced a training set containing a randomly chosen subset of 80% of the clusters and a test set containing the remaining 20%, making sure that both the training and testing sets contained at least one cluster expressing the gene and one not expressing the gene, according to MM0.5.

#### Production of the training and testing sets for morphological features

For molecular features, we also treated independently “All clusters” and “main OPC clusters”. We chose only the features evaluated (either present or absent) in more than 10 neuronal clusters. Similarly to genes, we also selected only the morphological features present in at least 2 clusters. Moreover, we selected only qualitative features, because the models produced for quantitative features would have to be evaluated in a different way. For each morphological feature we then produced a training set containing a randomly chosen subset of 80% of the clusters and a test set containing the remaining 20%. We made sure that both the training and testing sets contained at least one cluster presenting the feature and one not presenting the feature. The test set therefore contain at least 2 clusters, and the train set at least 8 clusters.

#### Production of the models

For each stage and each feature (molecular or morphological), we used the expression of different predictors (TFs, CSSs, TSs, TFs that are not TSs, developmental origins), in “All clusters” or “main OPC clusters”, with regular or randomized expression, to train a random forest model in the corresponding training set. If the feature to predict was part of the list of predictors, the feature was first removed from the predictors. We used the function randomForest with default parameters except for the number of trees that we set at 1000, using regression to predict gene expression or classification to predict morphological features (therefore we only built models for the morphologies that were assessed qualitatively and not quantitively), using the R package randomForest 4.6-14.

#### Evaluation of the models

To evaluate the quality of the model, we then computed either Pearson correlation (for molecular features) or MCC (for all features) between predicted values and real values, that we call observed values. We used the MCC because it gives the same importance to all 4 categories of the confusion matrix (true and false positives and negatives), which is well suited for unbalanced datasets^46^ such as differentially expressed genes or morphological features. Indeed, since we are interested in features present only in a subset of neuronal types, other metrics such as Pearson correlation would be disproportionately affected by the ability of the model to predict the absence of the feature (*i.e.,* the majority of the clusters).

However, MCC requires binarized predicted and observed values. For morphological features, both the observed and predicted values were already binarized in the case of qualitative features. For molecular features, we used MM0.5 as observed values, and we binarized the predicted values. The predicted values range between a maximal and a minimal value, different for each gene, with higher values in clusters where the gene is most likely to be expressed according to the model. However, it is unclear which of these values correspond to expression and non-expression according to the model. Therefore, to binarize the predictions, we scaled them between 0 and 1, and then binarized them using all thresholds between 0 and 1 with an increment of 0.01 (0, 0.01, 0.02, etc.): a threshold of 0 would correspond to a gene always expressed in the test set, a threshold of 1 to a gene never expressed, both impossible cases based on how we produced the test sets. If the model made meaningful predictions, and intermediate threshold will give better results. We therefore used the threshold giving the best fit between predicted and observed values, *i.e.*, the one giving the highest possible MCC. MCC ranges between 1 (perfect correlation) and -1 (perfect anticorrelation), with 0 denoting lack of correlation. However, the MCC values we obtained are slightly inflated because we choose the best possible MCC for each feature: our results obtained with regular predictors should therefore be compared to the results obtained with randomized predictors, and not to an MCC of 0.

#### Result tables and their interpretation

For molecular features, for each stage we produced several tables corresponding to different combinations of type of predictors, type of expression (regular or random), and clusters used (“All” or “main OPC”). These table contain the following columns: 1) “Features”: the gene for which the model was built, 2) “Top_predictors”: a ranking of their best 30 predictors (less if there are less predictors), which are the most likely candidate regulators of this feature, 3) “%IncMSE”: the percent increase in mean square error of the model when the values of the predictor are shuffled 4) “RMSE_TestSet”: The square root of the mean squared error between observed and predicted values, 5) “Cor_TestPreds_TestObs”: Pearson correlation between observed and predicted values, 6) “Best_MCC_TestSetBin_TestSetMM”: best MCC obtained between observed and predicted values, and 7) “Best_MCC_Threshold”: the threshold giving the best MCC for this model. For morphological features, we produced similar tables, with the following columns: 1) “Features”, 2) “Top_predictors”, 3) “%IncMSE”, 4) “OOBE”: Out Of Bag estimate of error rate, 5) “MCC_TestObs_TestPreds”.

For any feature and each type of predictors (TFs, CSSs, developmental origin…), high MCC values indicate that the random forest algorithm was able to identify a correlation between the predictors and the feature that is valid both in the train and the test sets. Different train set would lead to different models, and different test set would lead to different MCC values for a given model. Moreover, random forest functions by building a forest of many decision trees built from a randomly sampled subset of the data: the algorithm will produce slightly different models, and therefore obtain a slightly different MCC value, each time it is run. By chance, sometimes this MCC value will be a particularly high or low outlier. Lastly, sometimes correlation between predictors and a feature can be due to chance and not be the result of a molecular mechanism, which explains why randomized predictors sometimes yield models with high MCC values. To mitigate these caveats, in this work for each feature we used a randomly chosen train and test set, and we only drew conclusions by using all the MCC values obtained for the hundreds of molecular and morphological features tested. Because the status of many morphological features could be evaluated only in few clusters, in several cases the train and test set were particularly small (in the worst cases, 8 and 2 clusters respectively). This strongly increased the chance that predicted status of these molecular features were the same in each cluster of the test set, and in such cases MCC is not defined. Because of this, models could be produced for only a small number of morphological features, which limits the strength of the conclusions drawn from morphological features.

For anyone interested in using the results of our random forest approach to experimentally find regulators of a given feature, it should be noted that all features present in our result tables will have corresponding ranked predictors, but only those associated to a random forest algorithm with good MCC values should be considered as candidate regulators of the feature. Moreover, the models produced using only main OPC clusters performed, on average, better than models produced using all neuronal clusters (*i.e.,* including main OPC clusters, but also tOPC, inner proliferation center, lamina, central brain… Fig.S13), which suggests that the regulation of neuronal features is different in neurons produced from different neurogenic domains. Therefore, it would be advisable to use the models produced using all neuronal clusters only when necessary, *i.e.,* to study regulations in non-main OPC clusters. Lastly, because random forest can only identify correlations, additional ways to reduce the number of candidate regulators for a given feature should be considered, for instance by plotting the expression of the features and of the ranked predictors, or taking into account other published data (presence of TF binding sites near the promoter, ChIP-seq data even if it was in other tissues or developmental stages, literature search…).

### Production of the heatmaps of the presence of neuronal features

To produce heatmaps of the TSs expressed during all of development in all clusters sharing a combination of specification modules, we only used the TSs expressed in less than 50% of the main OPC neuronal clusters according to MM0.5. We considered that a TS was expressed in a given cluster during all its development if it was expressed in at least 5 out of the 6 developmental stages of our scRNA-seq atlas according to MM0.5. We did not require for 6 out of 6 stages to account for imperfection in the modeling of gene expression by Mixture Modeling.

To produce heatmaps of genes expressed at a given stage in all clusters sharing a combination of specification modules, we only used genes expressed in less than 50% of the main OPC neuronal clusters according to MM0.5.

To produce heatmaps of morphological features present in all clusters sharing a combination of specification modules, if the presence of a feature in a neuronal type was unsure, we considered it absent (*i.e.*, all “NA” values were replaced by a value of 0). Indeed, we chose to be conservative because the purpose of these heatmaps is to find features present in all neuronal types sharing a common origin.

In any case, we considered that a feature (gene, TSs, morphological feature) was present in all clusters from a given origin when 75% of the clusters from this origin presented the feature, instead of 100%, to account for the imperfect resolution of our determination of developmental origins. For all the heatmaps, the list of TFs, CSSs or TSs used where the same as for the random forest analyses.

