## Supplementary material for "Establishment of terminal selector combinations in optic lobe neurons": Annex 1

### Gates used to sort the scRNA-seq libraries

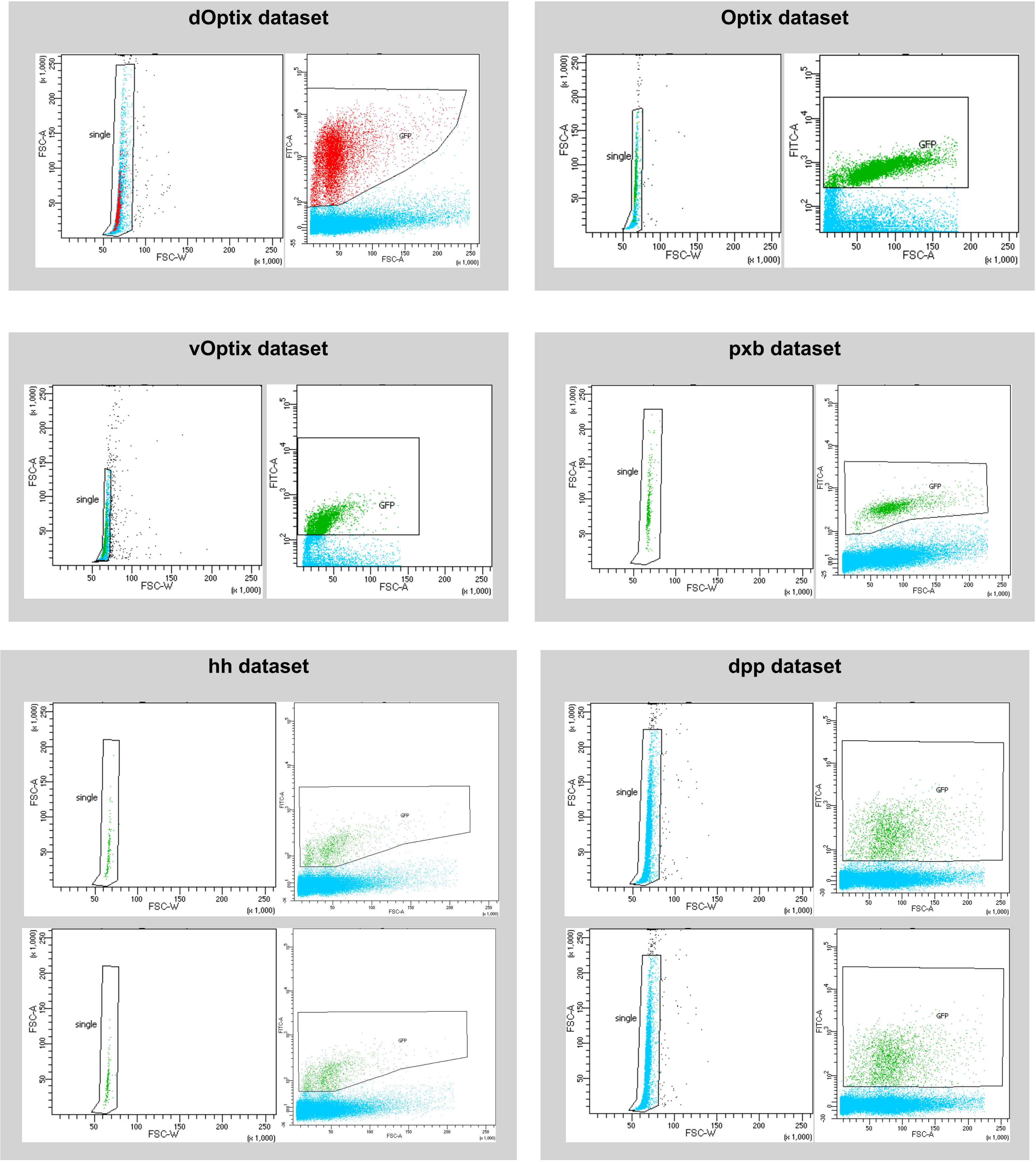

**Thresholds used to filter cells according to their number of genes (Gene nb), number of UMIs (RNA nb) and percentage of mitochondrial genes (Mito %)**

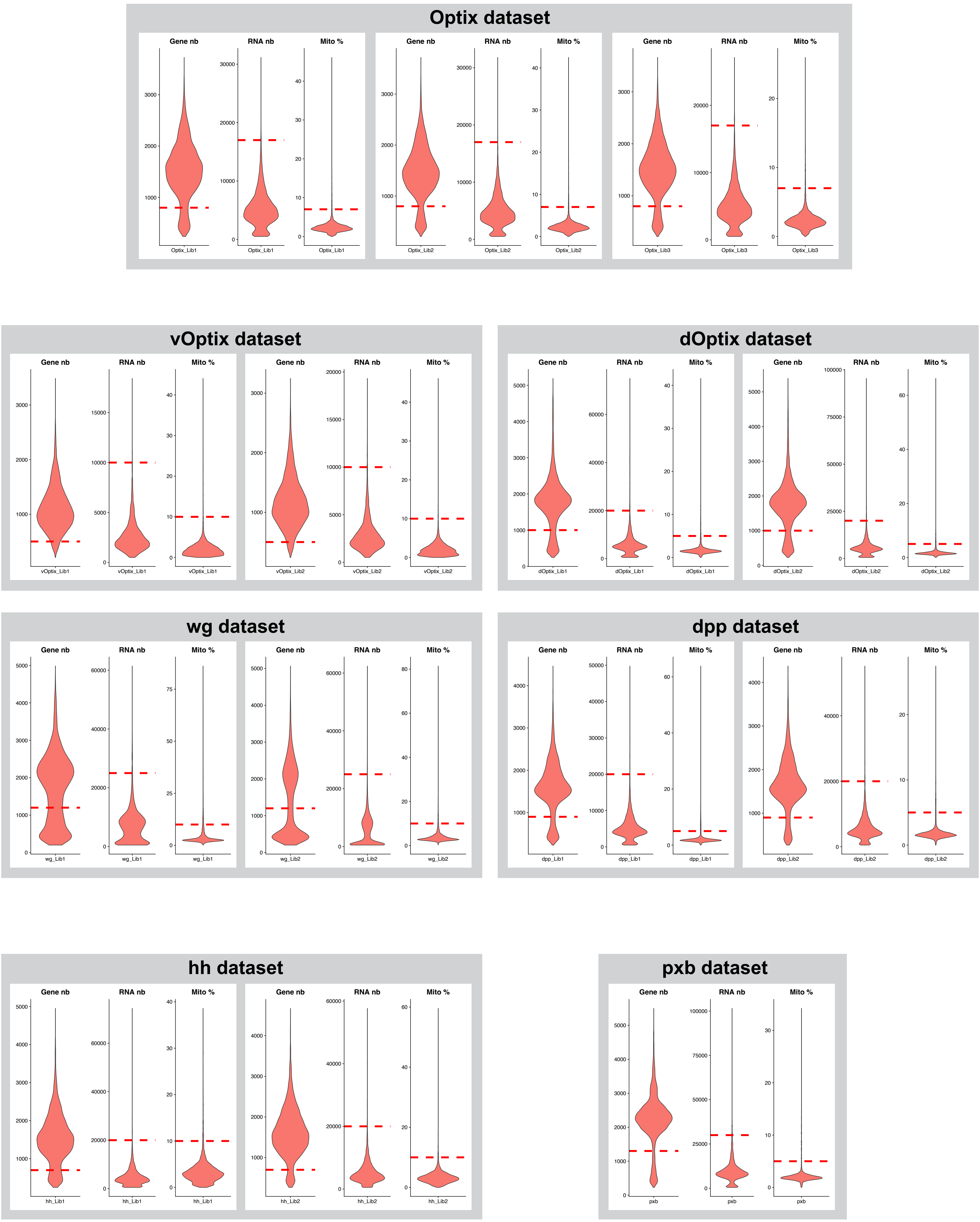

### Confidence in the annotation of each cell, grouped by class.

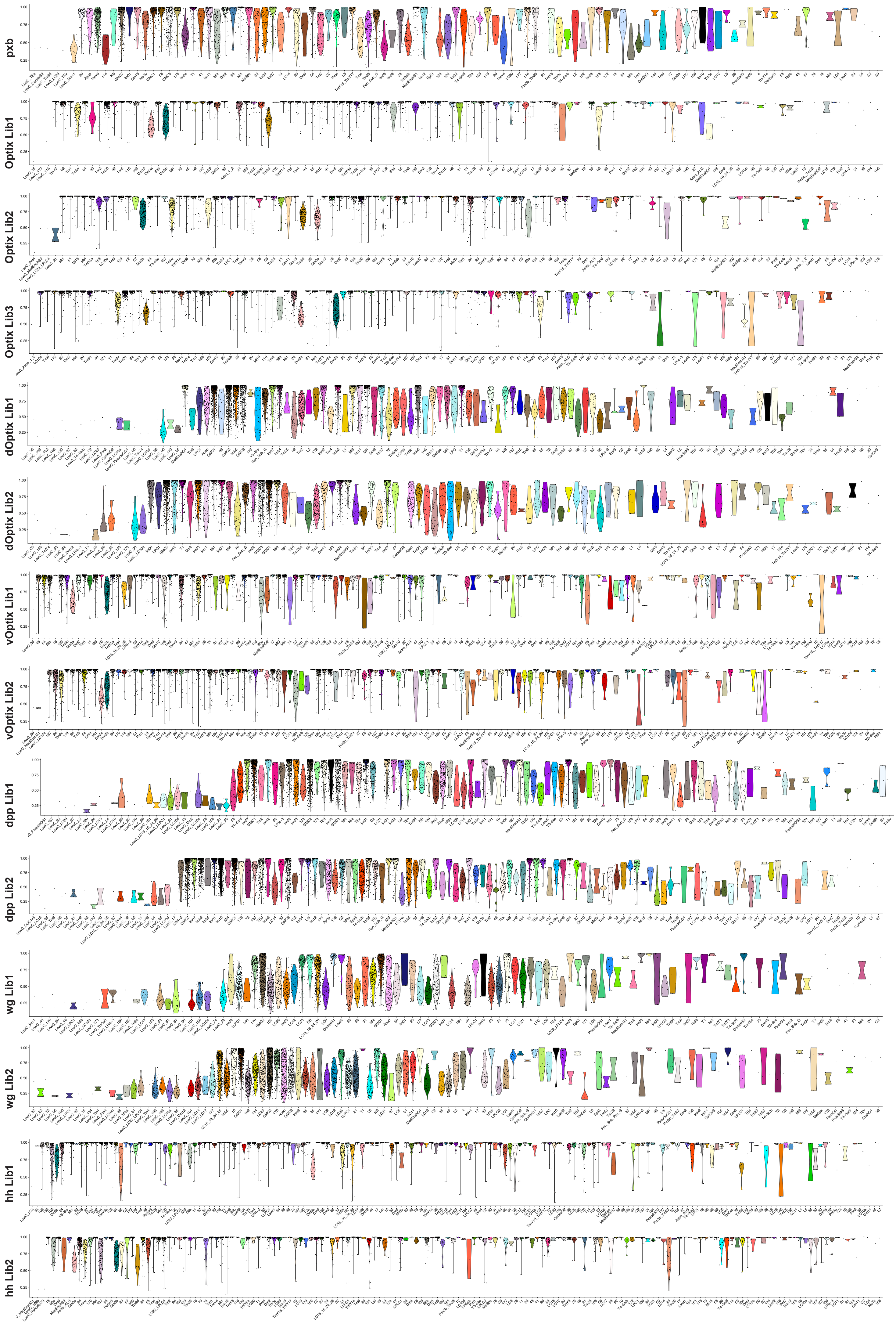

**Annotation of the different libraries by the neural network classifier (tSNEs with colored cells) and classes flagged as annotated with low confidence (grey tSNEs with red cells)**

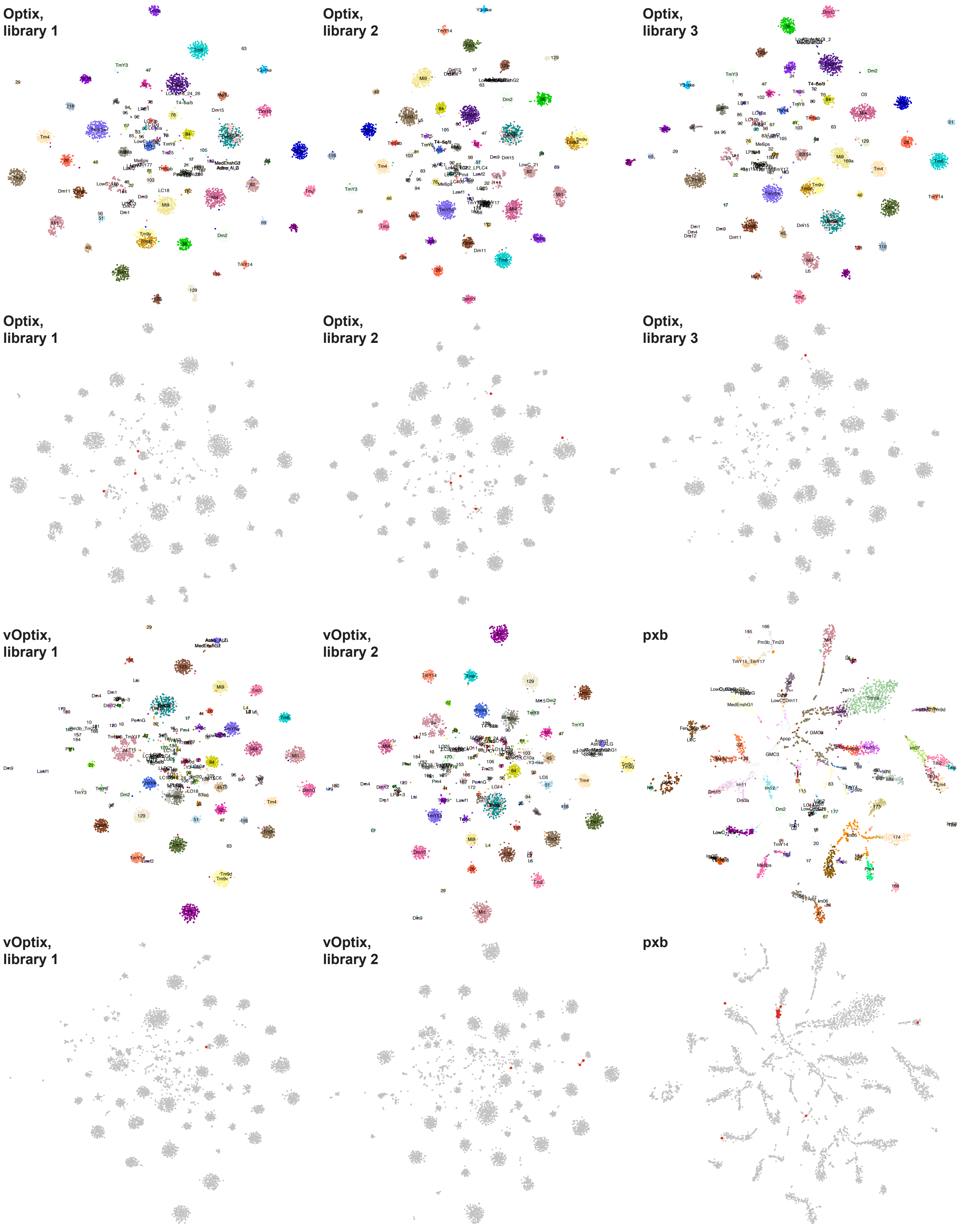

**Annotation of the different libraries by the neural network classifier (tSNEs with colored cells) and cells flagged as annotated with low confidence (grey tSNEs with red cells)**

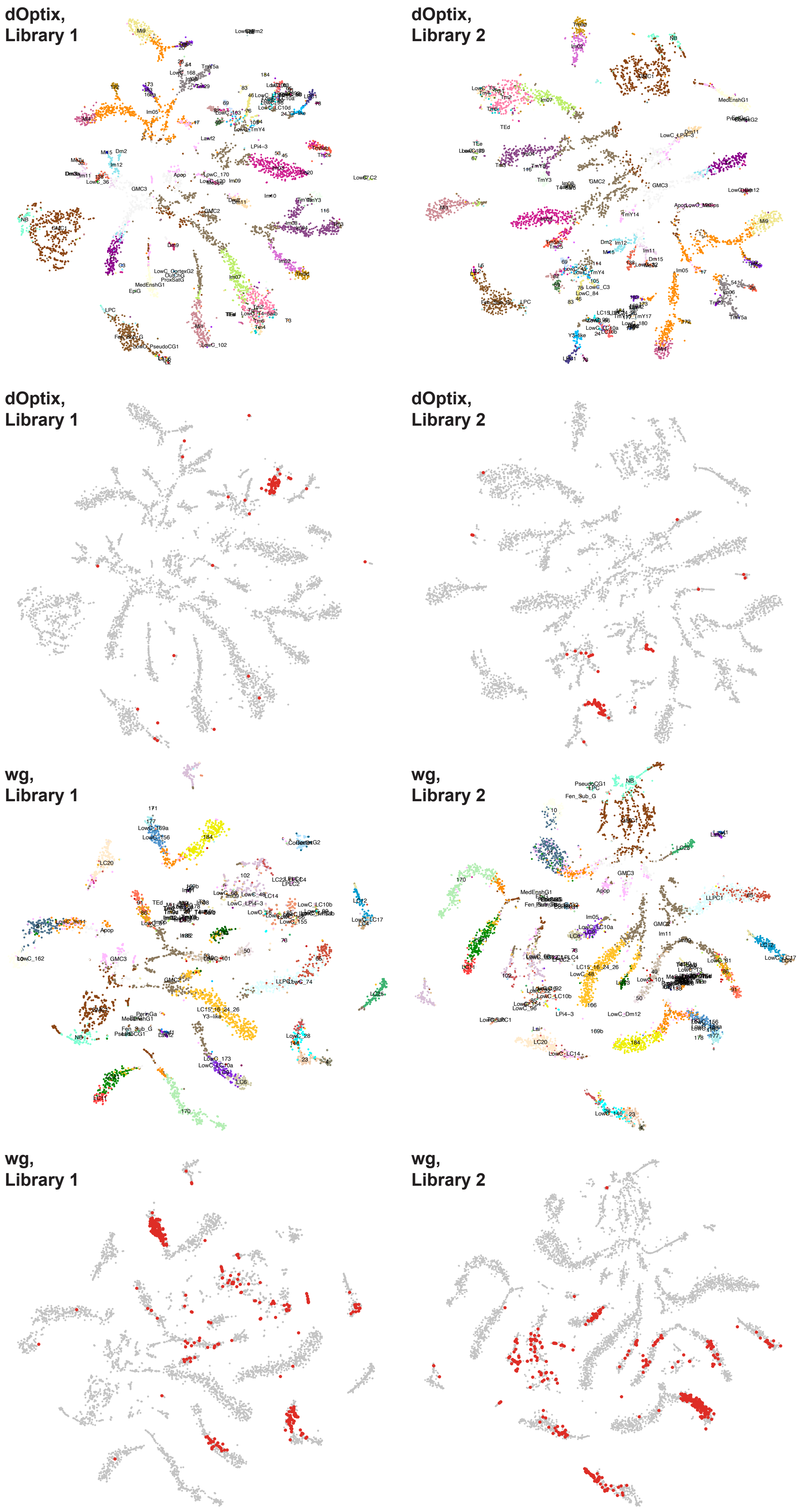

**Annotation of the different libraries by the neural network classifier (tSNEs with colored cells) and cells flagged as annotated with low confidence (grey tSNEs with red cells)**

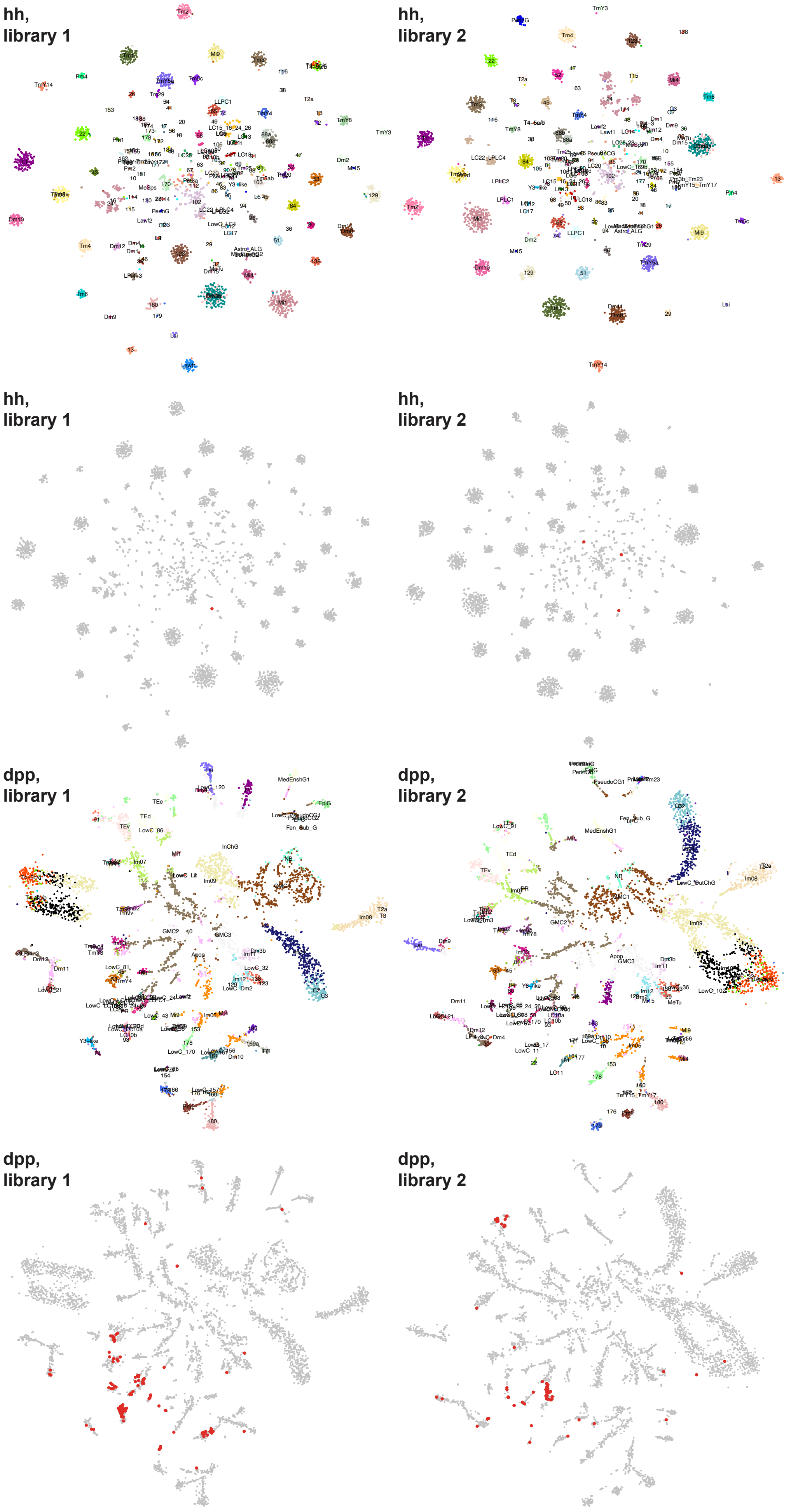

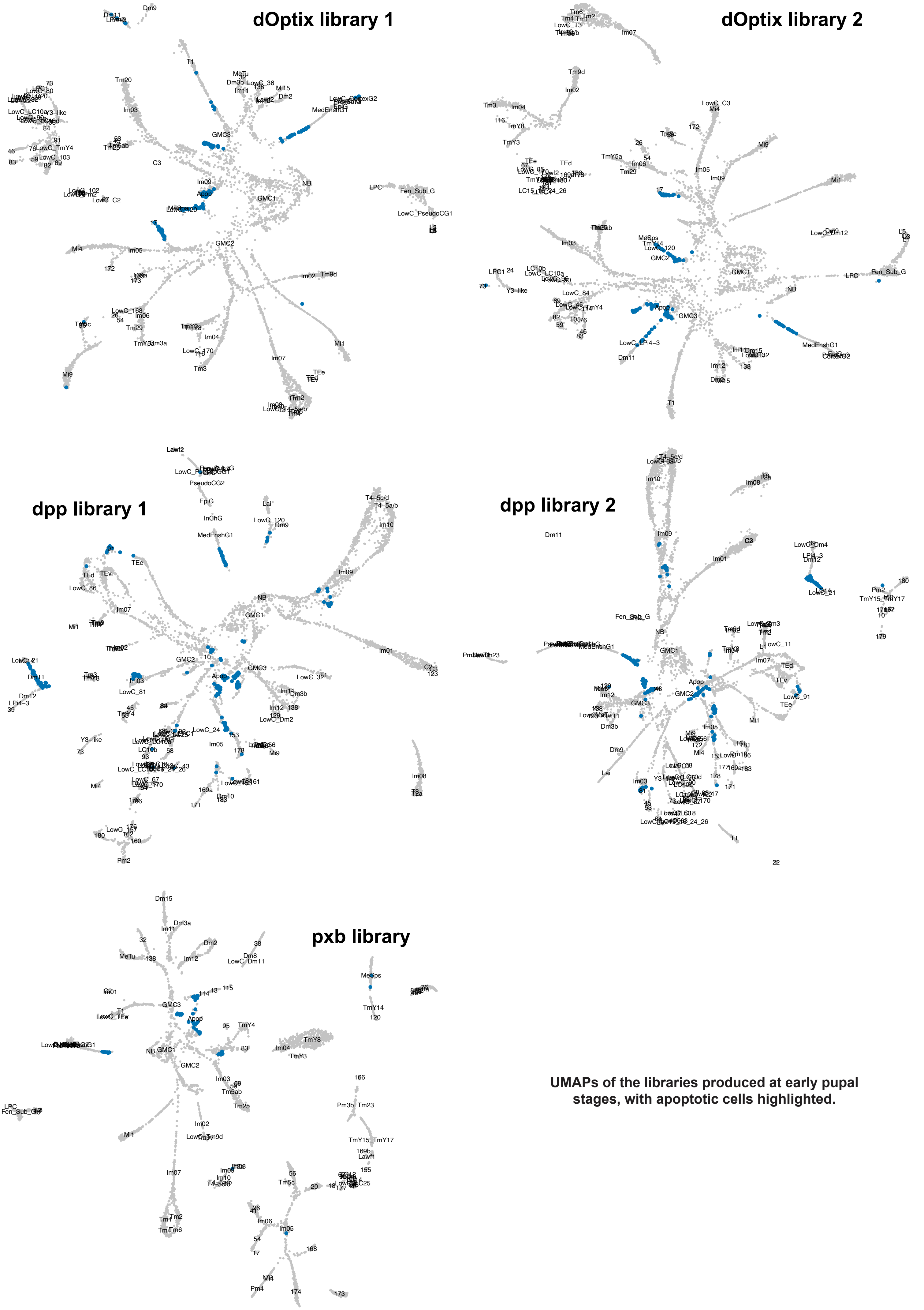


**UMAPs of the libraries produced at early pupal stages, with apoptotic cells highlighted.**

### Abundance of main OPC clusters in each dataset (reference dataset: Adult, clusters used for normalization: T1, Mi1, Tm1, Tm2, Tm4, 138)

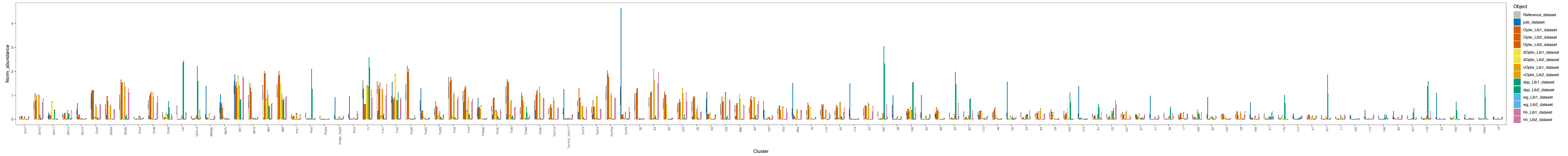


### Abundance of non-main OPC clusters in each dataset (reference dataset: Adult, clusters used for normalization: T1, Mi1, Tm1, Tm2, Tm4, 138)

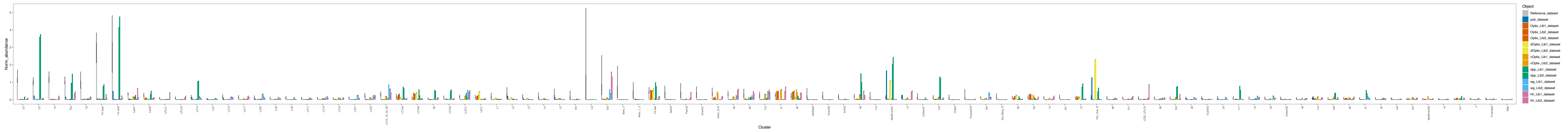


### Abundance of main OPC clusters in each dataset (reference dataset: P15, clusters used for normalization: T1, Mi1, Tm1, Tm2, Tm4, 138)

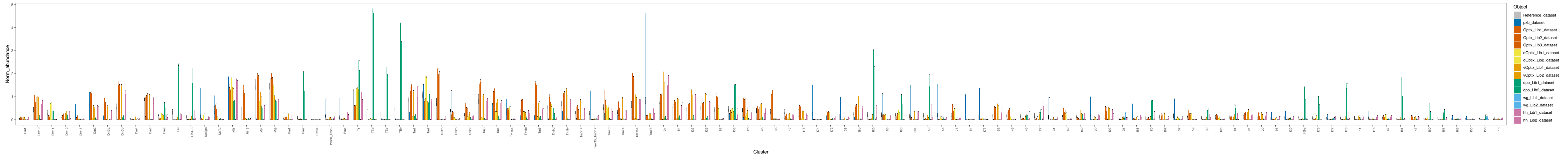


### Abundance of non-main OPC clusters in each dataset (reference dataset: P15, clusters used for normalization: T1, Mi1, Tm1, Tm2, Tm4, 138)

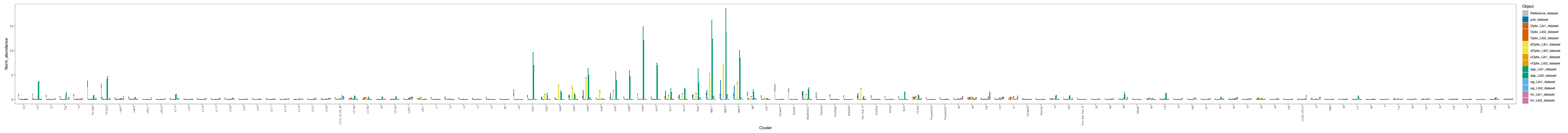

### Binarization thresholds (reference dataset: Adult, clusters used for normalization: T1, Mi1, Tm1, Tm2, Tm4, 138)

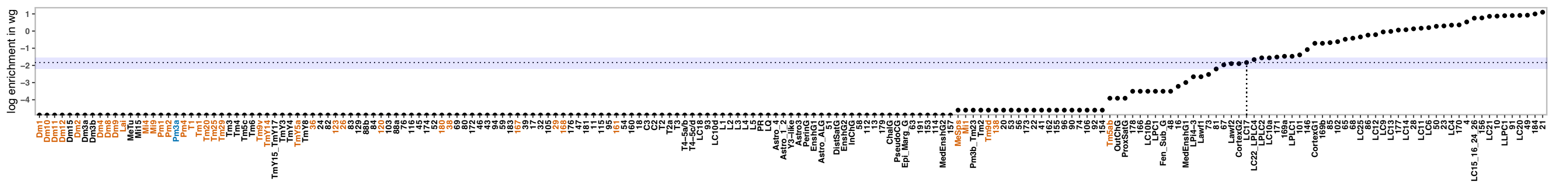


### Binarization thresholds (reference dataset: P15, clusters used for normalization: T1, Mi1, Tm1, Tm2, Tm4, 138)

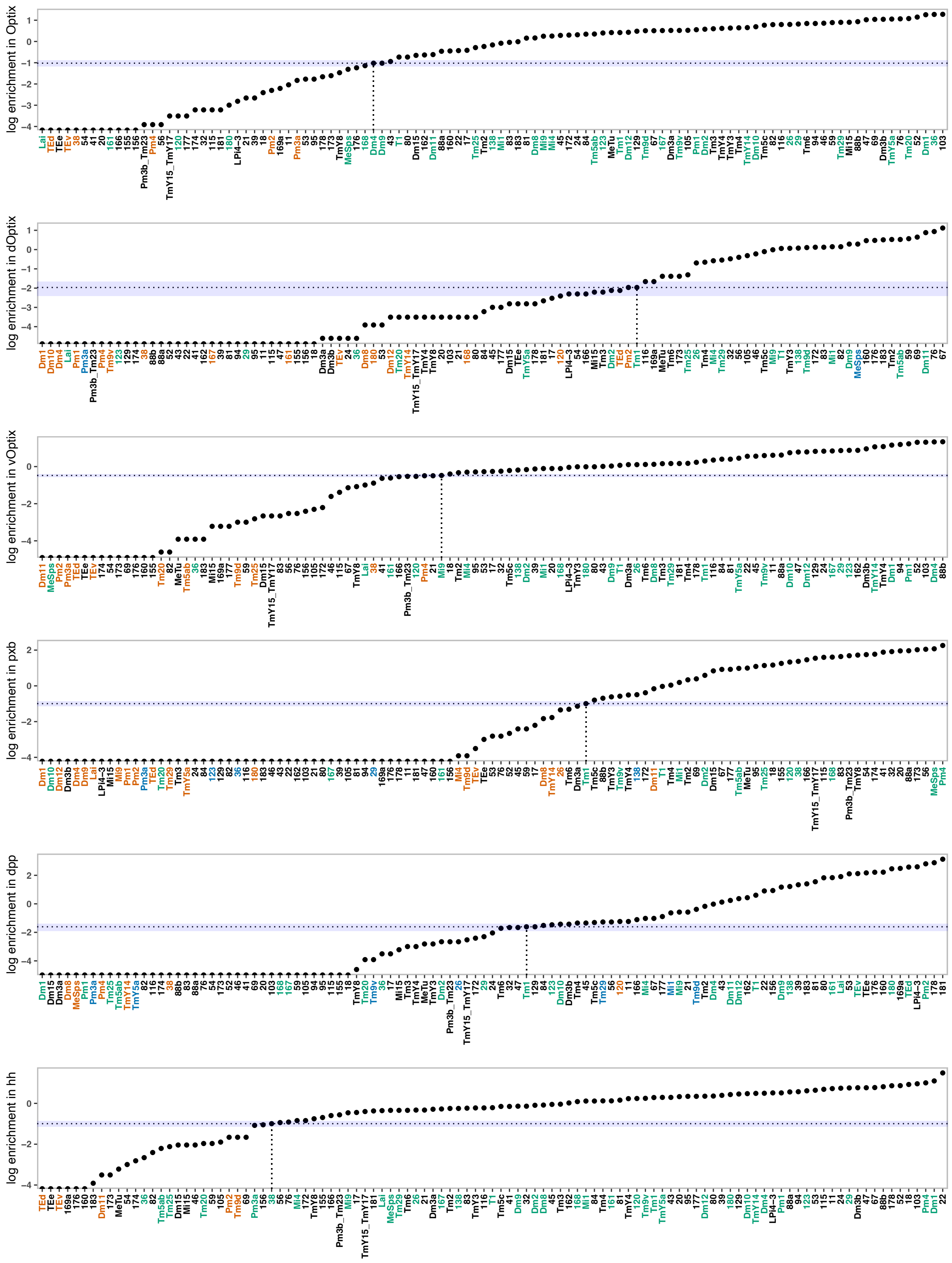


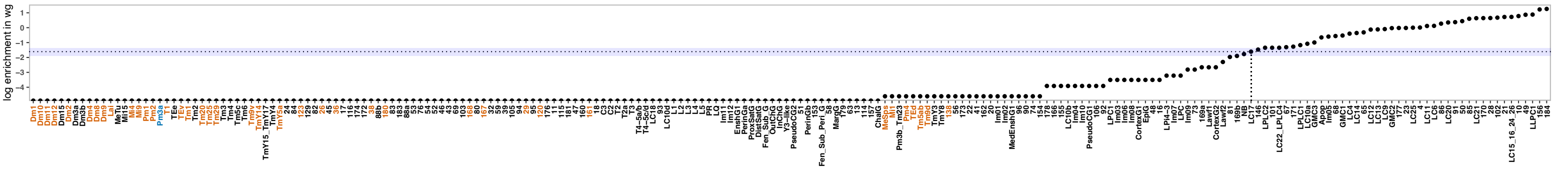
