## Supplementary material for "Establishment of terminal selector combinations in optic lobe neurons": Annex 2: Read_Me.txt

Please use the legend of Fig4 to read these plots, and not the scale on the side of these heatmaps.

The name of the files include values of the parameters used to produce the files: see Methods.

CSSs = Cell Surface and Secreted molecules
TSCs = Terminal Selector candidates
NO, TO, SO = Notch, Temporal and Spatial origin
