## Supplementary figures and images for "Establishment of terminal selector combinations in optic lobe neurons"

### Adult_NO_non_TFs_CSSs_0.75ClusterFreq_0.5BinThreshold_0.5SpecificityThreshold_Simplified.pdf

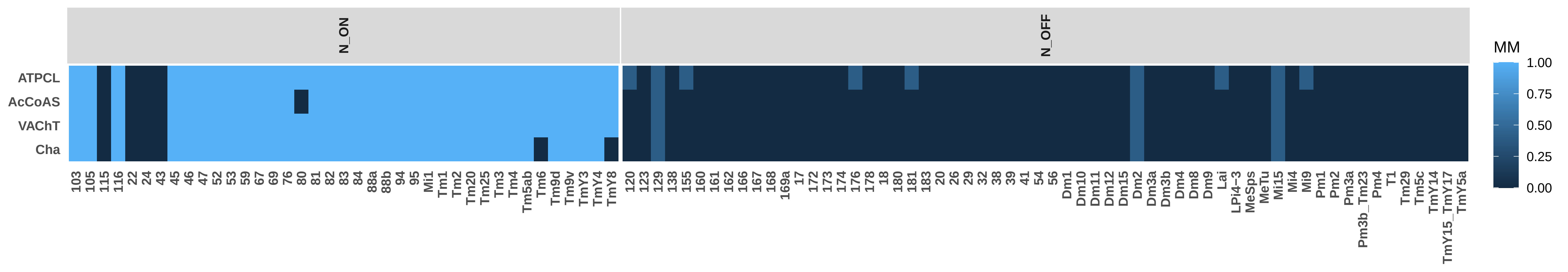

### Adult_NO_SO_CSSs_0.75ClusterFreq_0.5BinThreshold_0.5SpecificityThreshold_Simplified.pdf

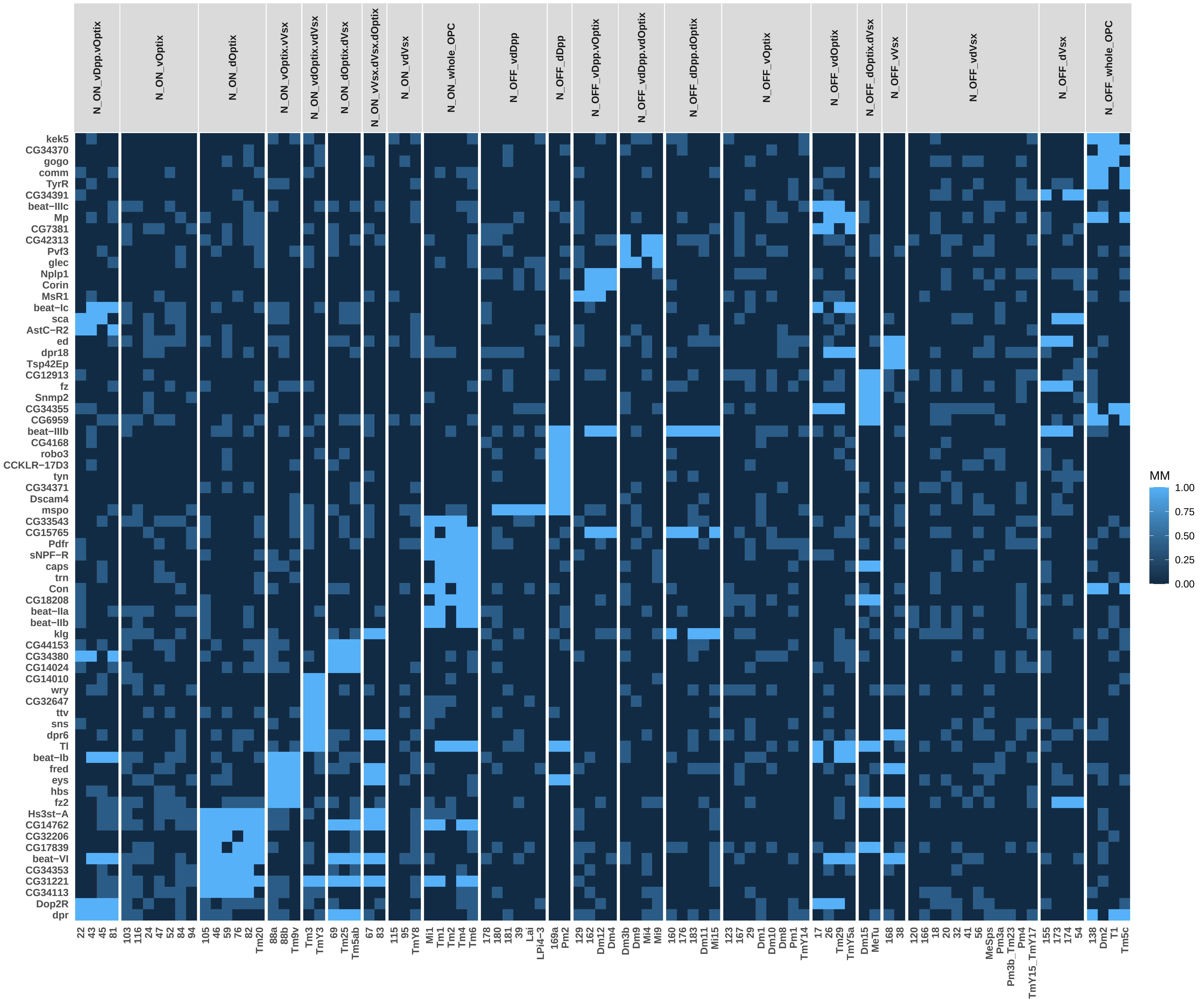

### Adult_NO_SO_non_TFs_CSSs_0.75ClusterFreq_0.5BinThreshold_0.5SpecificityThreshold_Simplified.pdf

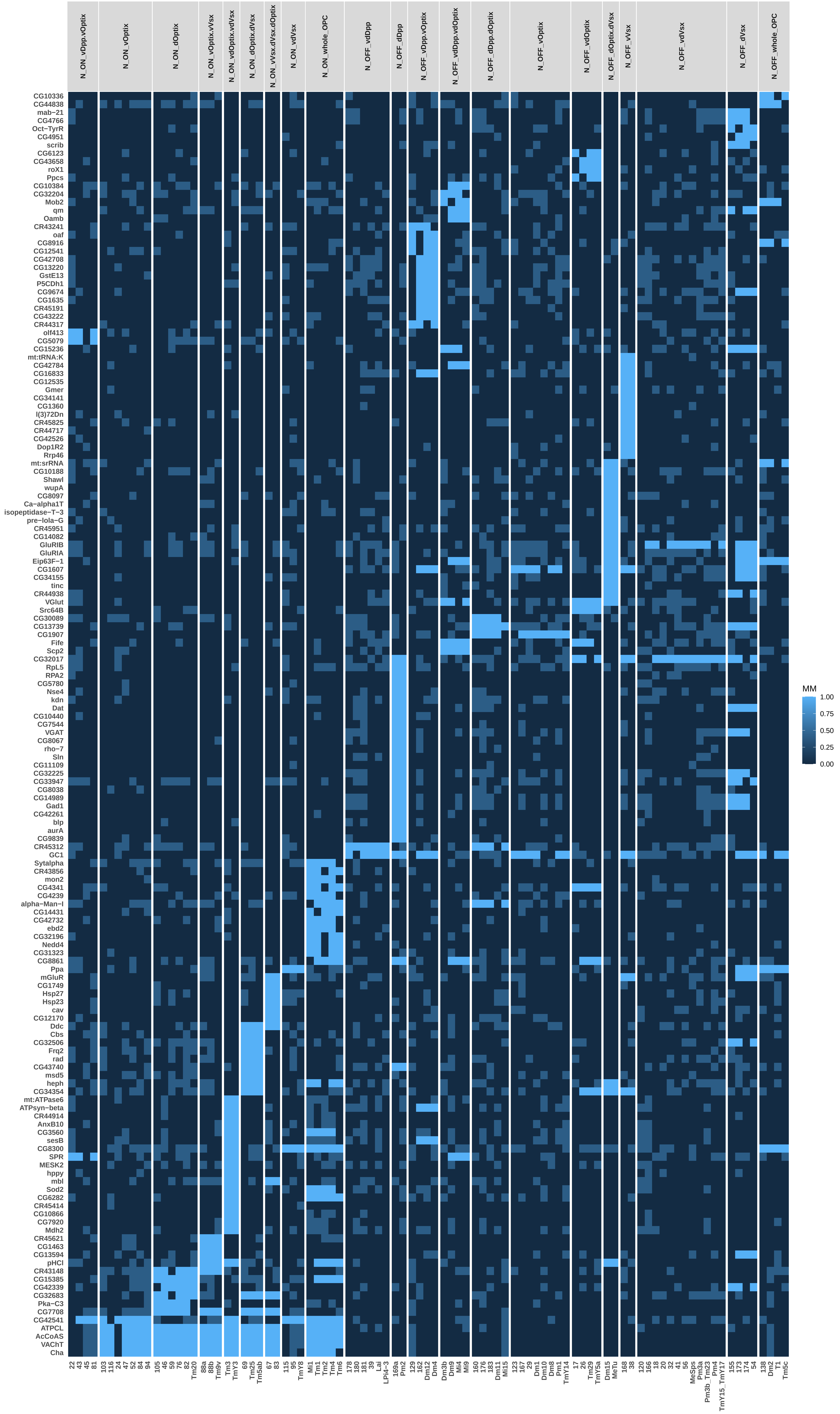

### Adult_NO_TO_SO_CSSs_0.75ClusterFreq_0.5BinThreshold_0.5SpecificityThreshold_Simplified.pdf

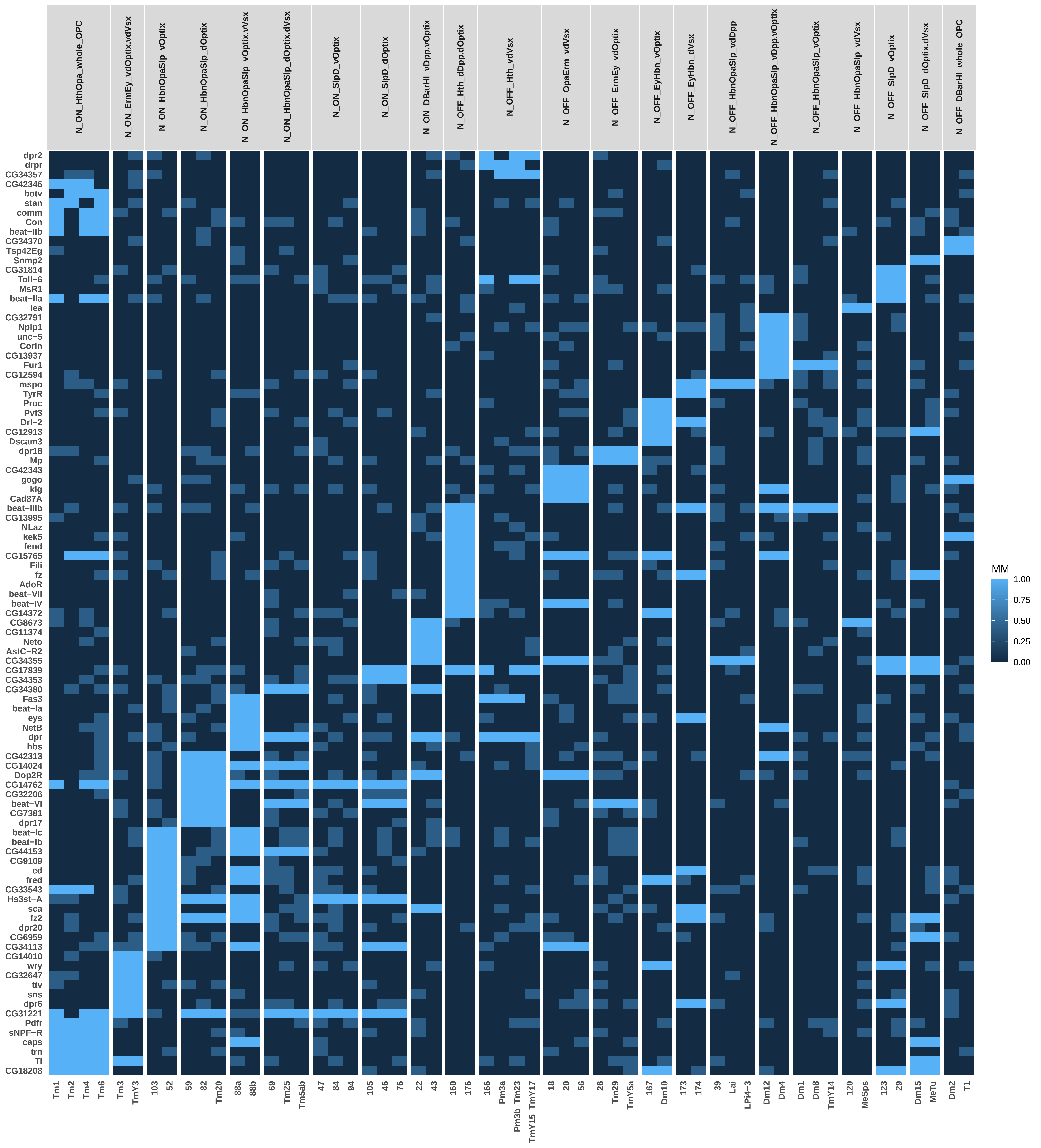

### Adult_NO_TO_SO_non_TFs_CSSs_0.75ClusterFreq_0.5BinThreshold_0.5SpecificityThreshold_Simplified.pdf

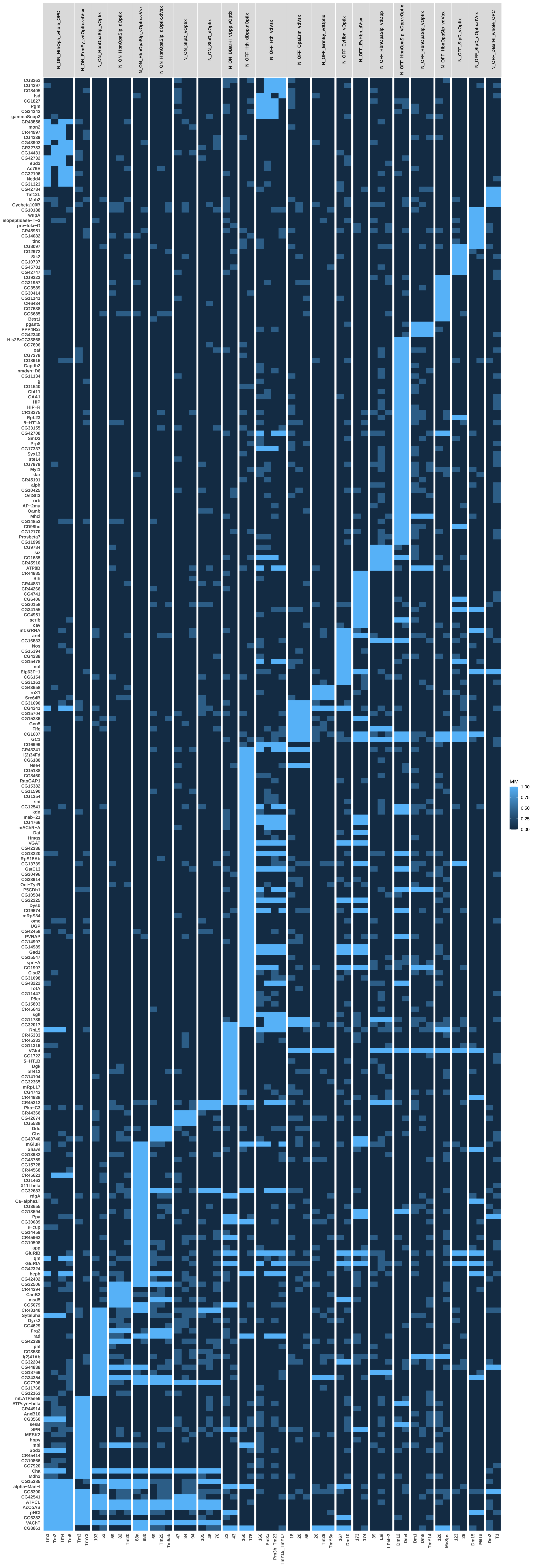

### Adult_SO_CSSs_0.75ClusterFreq_0.5BinThreshold_0.5SpecificityThreshold_Simplified.pdf

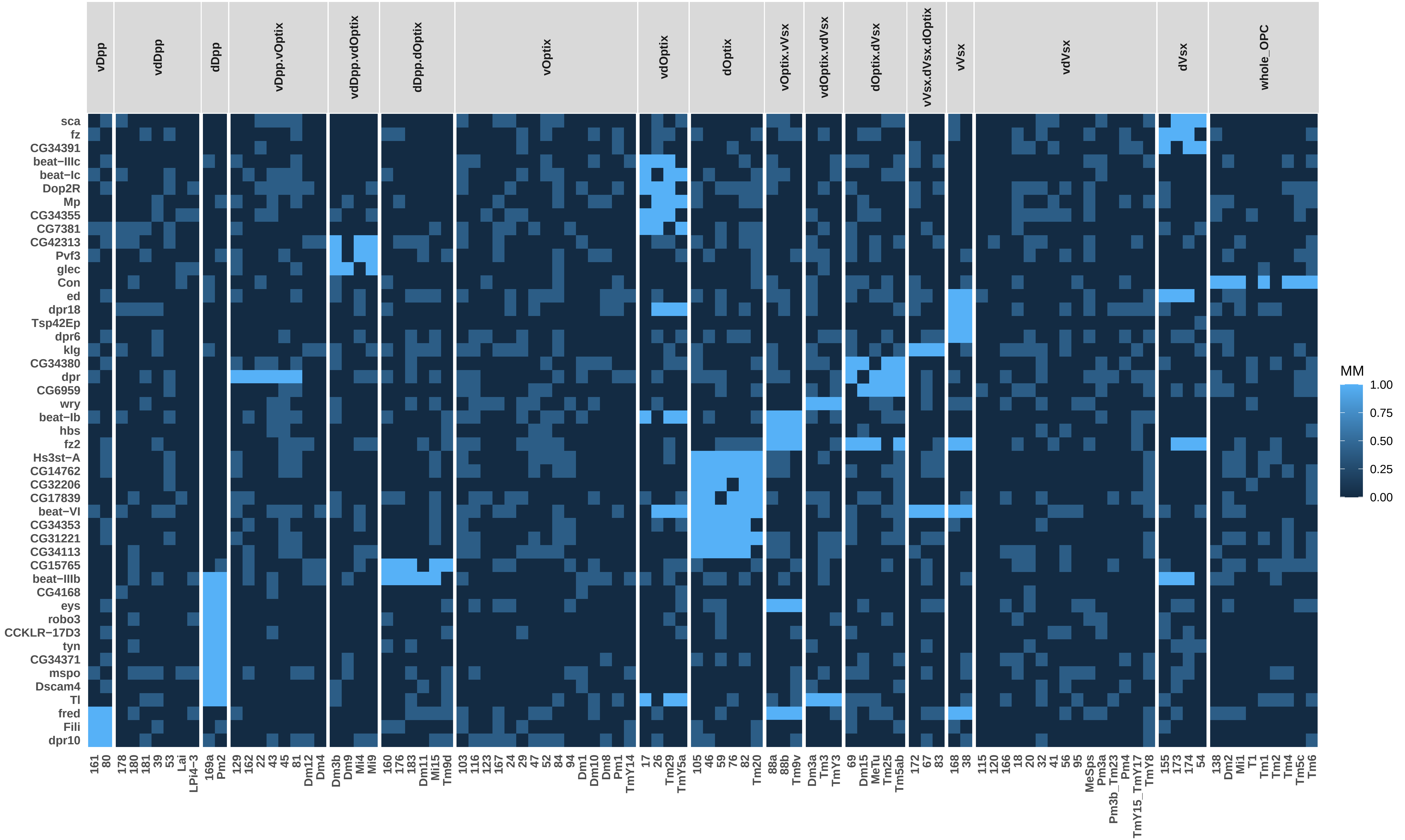

### Adult_SO_non_TFs_CSSs_0.75ClusterFreq_0.5BinThreshold_0.5SpecificityThreshold_Simplified.pdf

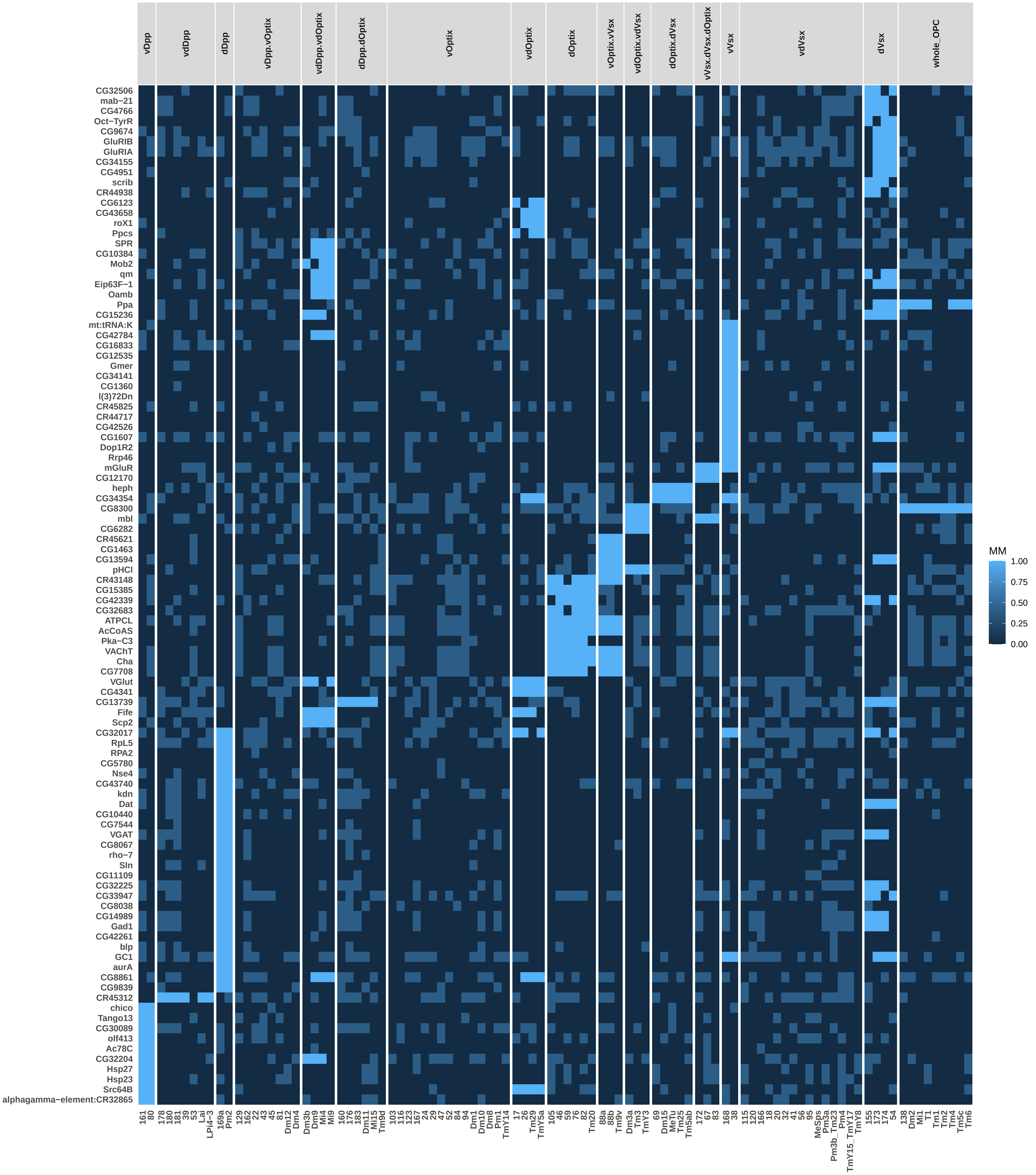

### Adult_TO_CSSs_0.75ClusterFreq_0.5BinThreshold_0.5SpecificityThreshold_Simplified.pdf

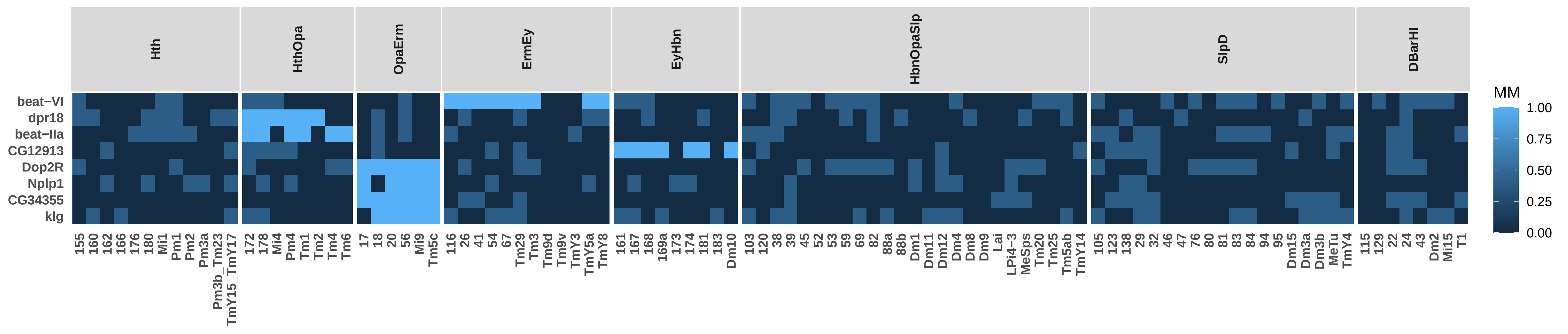

### Adult_TO_NO_CSSs_0.75ClusterFreq_0.5BinThreshold_0.5SpecificityThreshold_Simplified.pdf

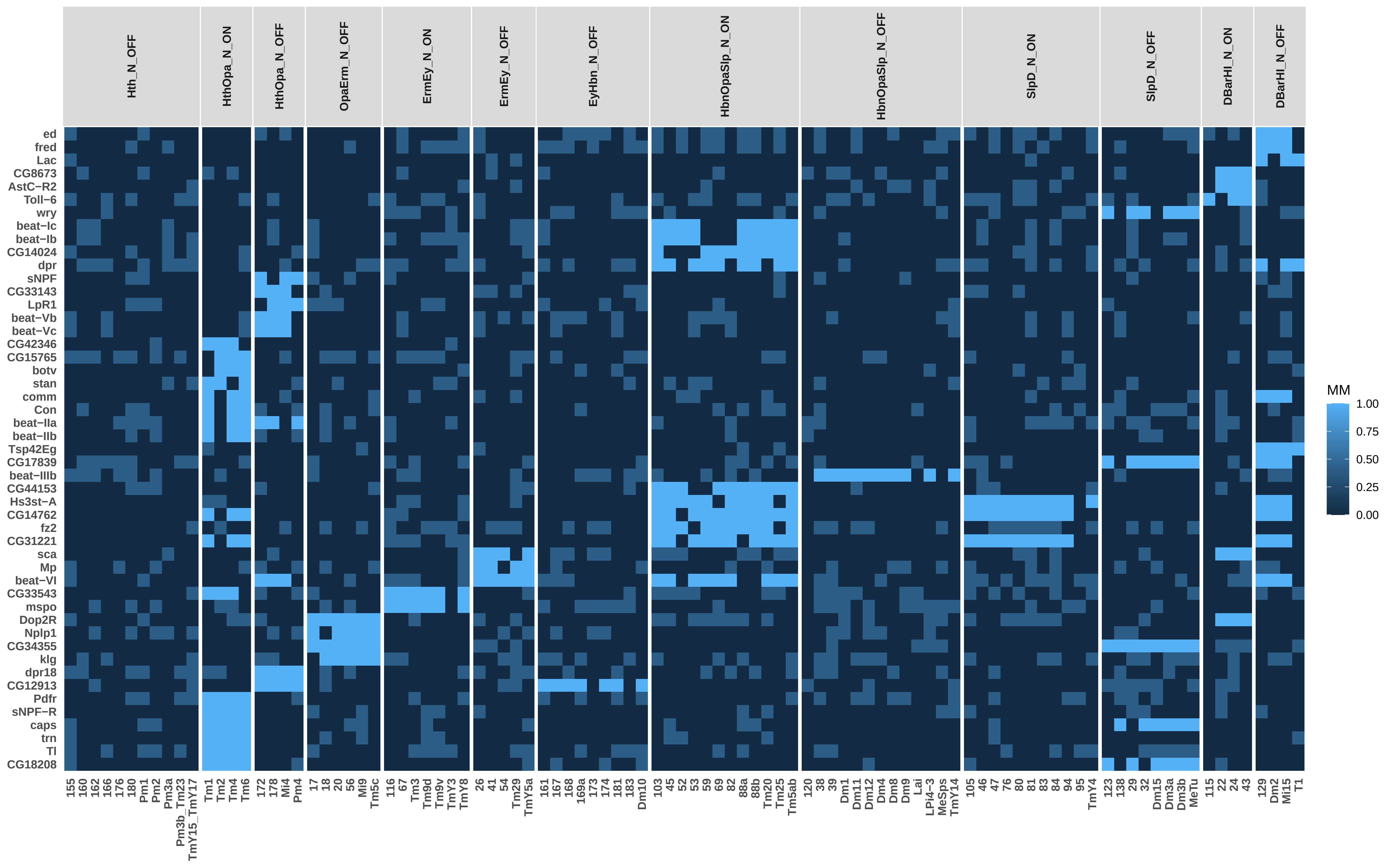

### Adult_TO_NO_non_TFs_CSSs_0.75ClusterFreq_0.5BinThreshold_0.5SpecificityThreshold_Simplified.pdf

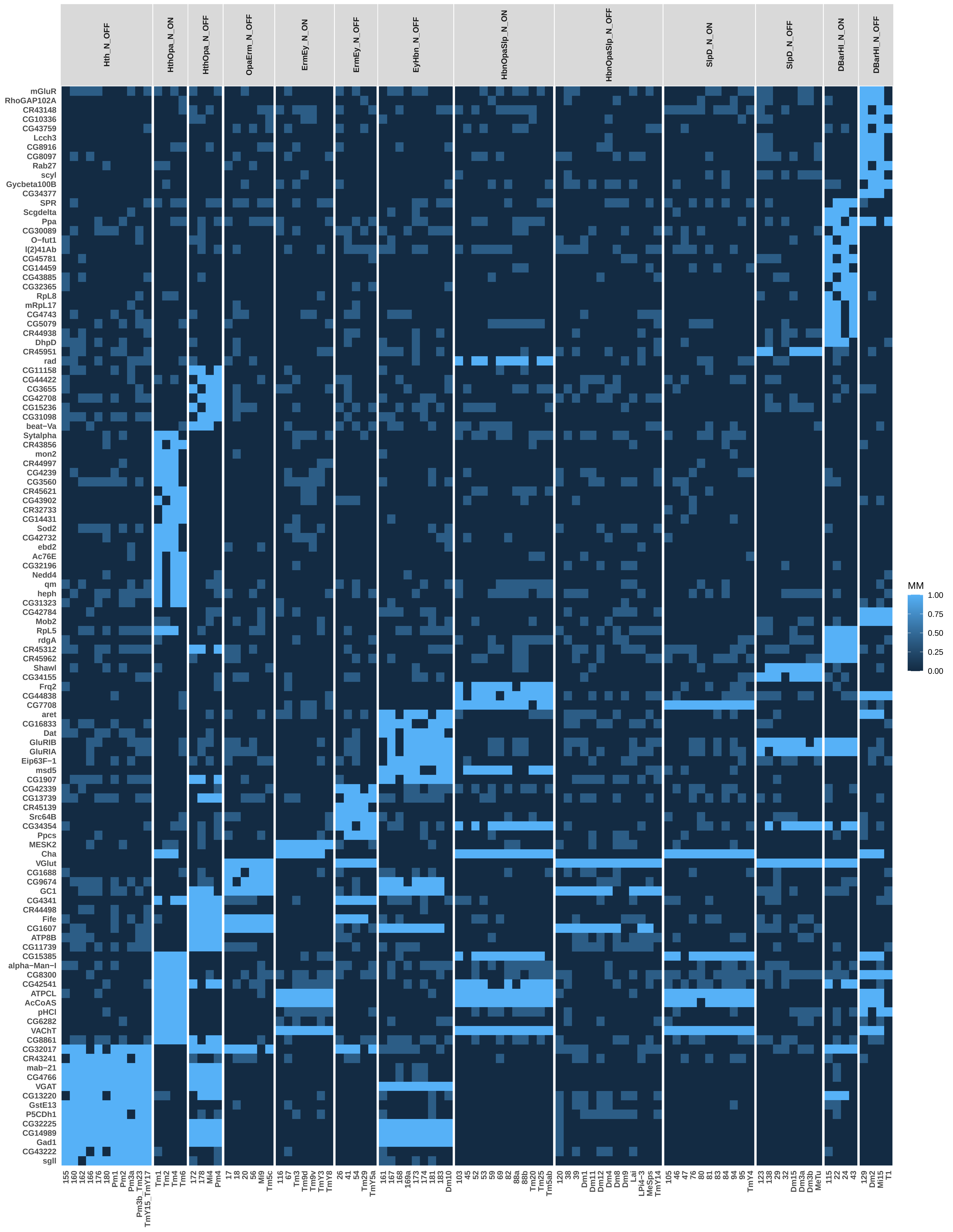

### Adult_TO_non_TFs_CSSs_0.75ClusterFreq_0.5BinThreshold_0.5SpecificityThreshold_Simplified.pdf

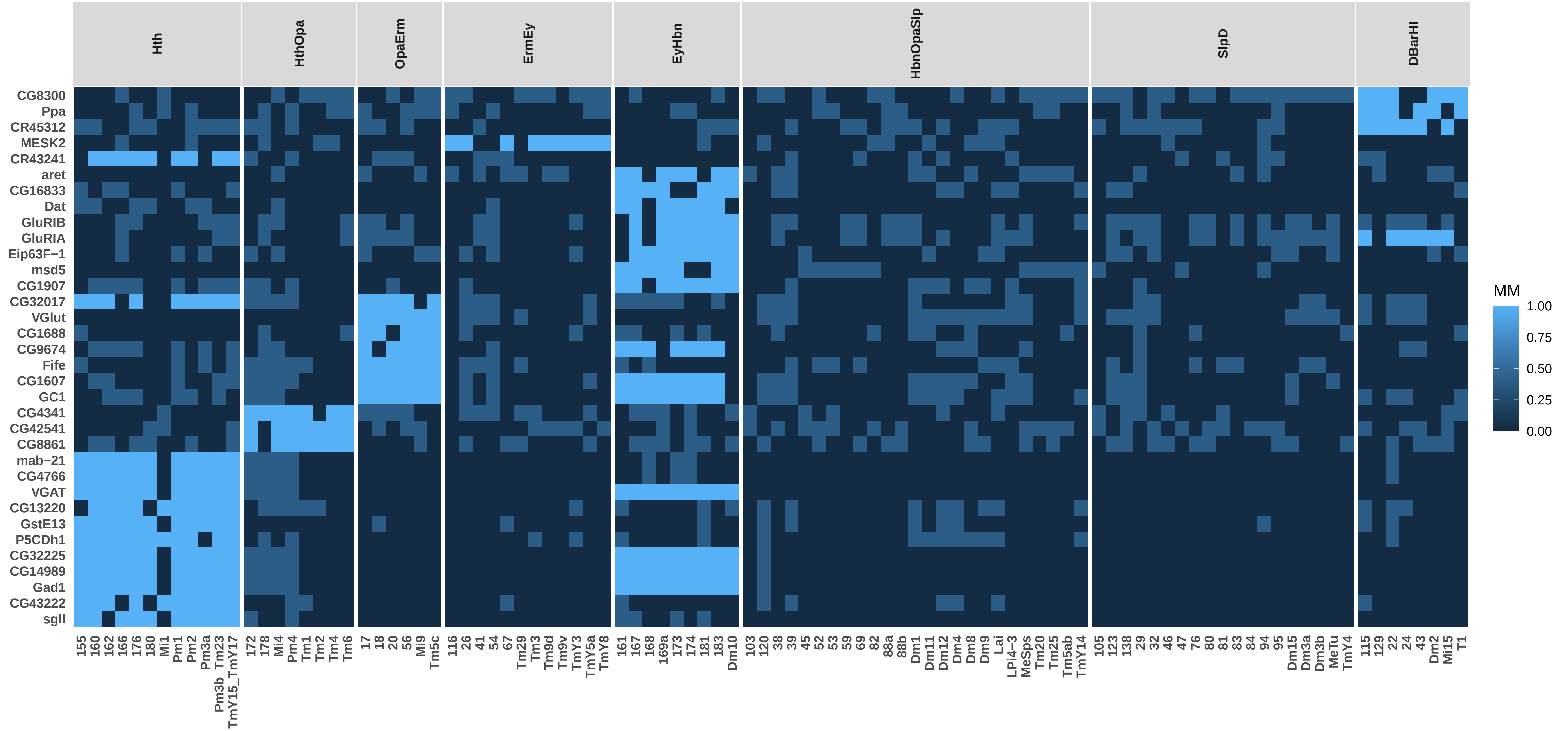

### Adult_TO_SO_CSSs_0.75ClusterFreq_0.5BinThreshold_0.5SpecificityThreshold_Simplified.pdf

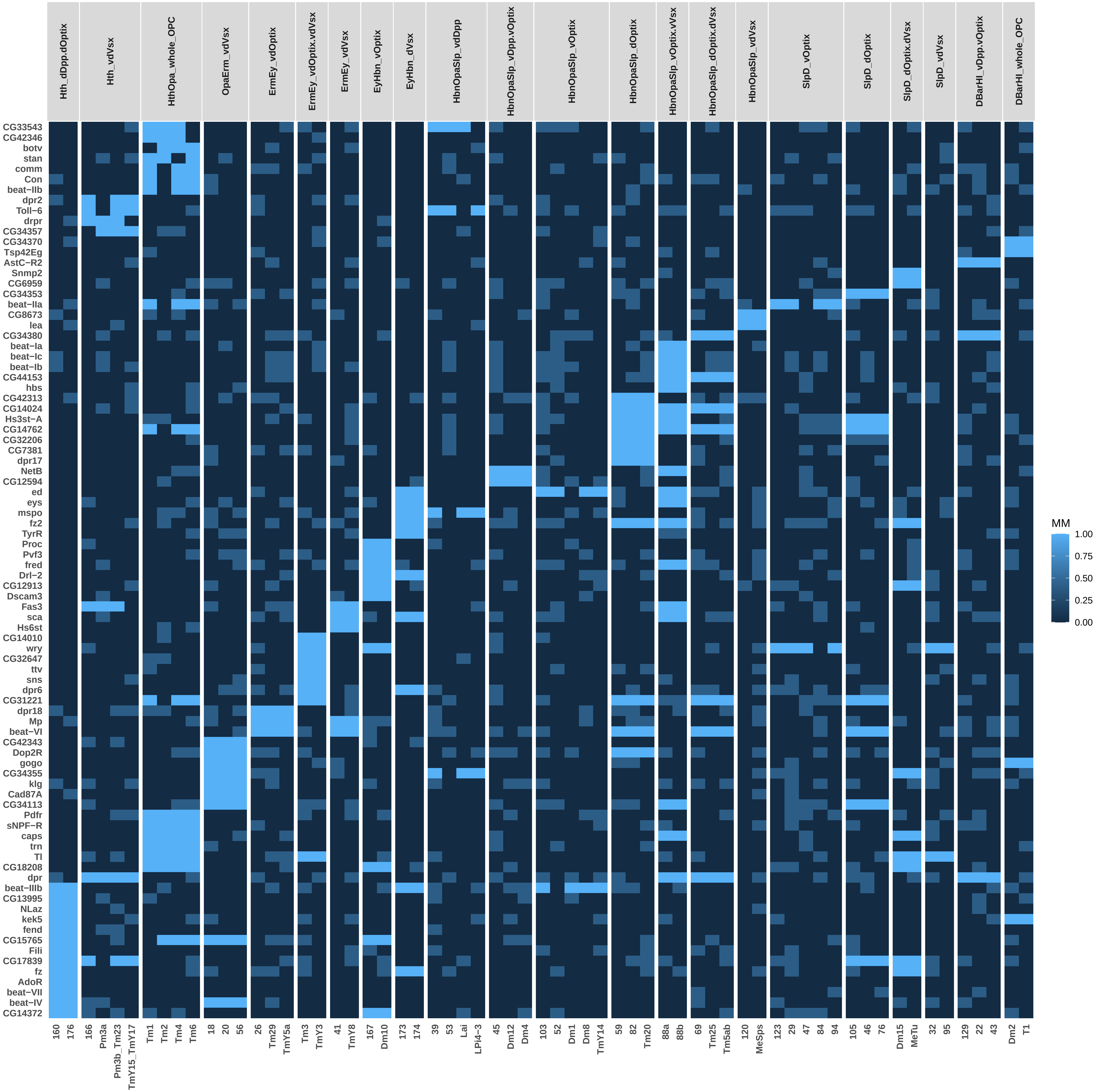

### Adult_TO_SO_non_TFs_CSSs_0.75ClusterFreq_0.5BinThreshold_0.5SpecificityThreshold_Simplified.pdf

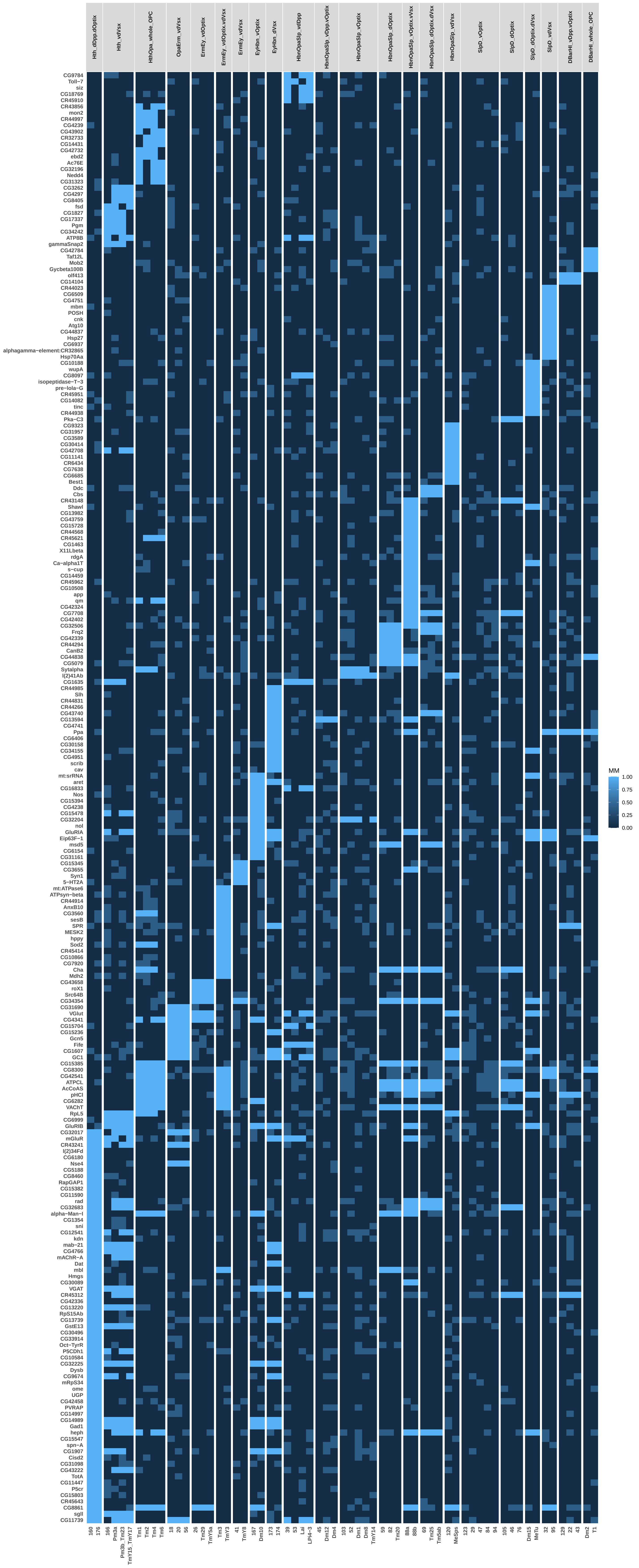

### P15_NO_CSSs_0.75ClusterFreq_0.5BinThreshold_0.5SpecificityThreshold_Simplified.pdf

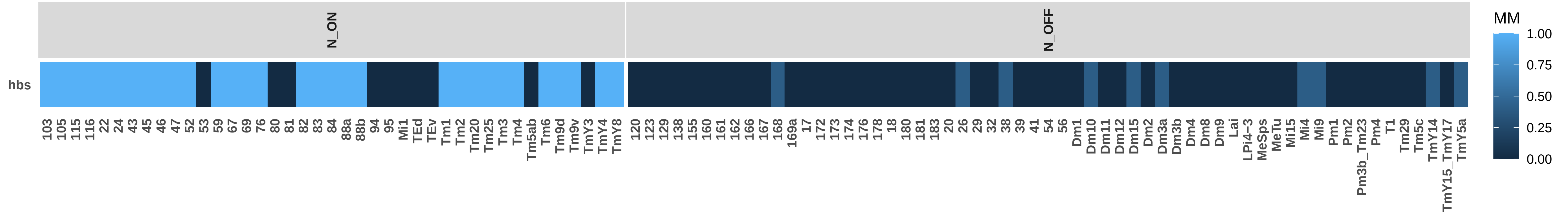

### P15_NO_non_TFs_CSSs_0.75ClusterFreq_0.5BinThreshold_0.5SpecificityThreshold_Simplified.pdf

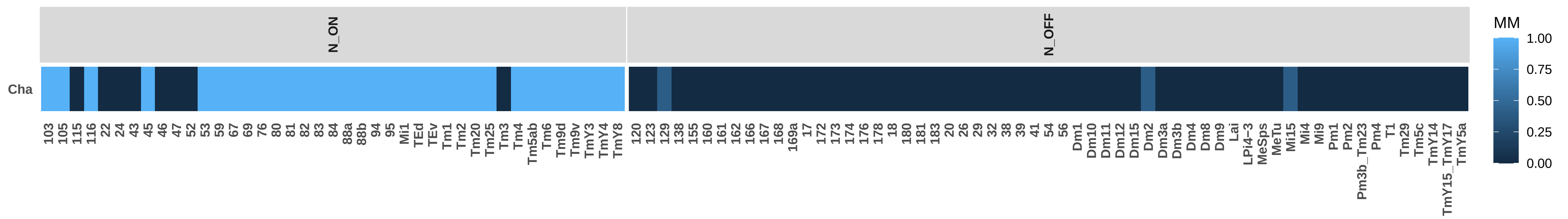

### P15_NO_SO_CSSs_0.75ClusterFreq_0.5BinThreshold_0.5SpecificityThreshold_Simplified.pdf

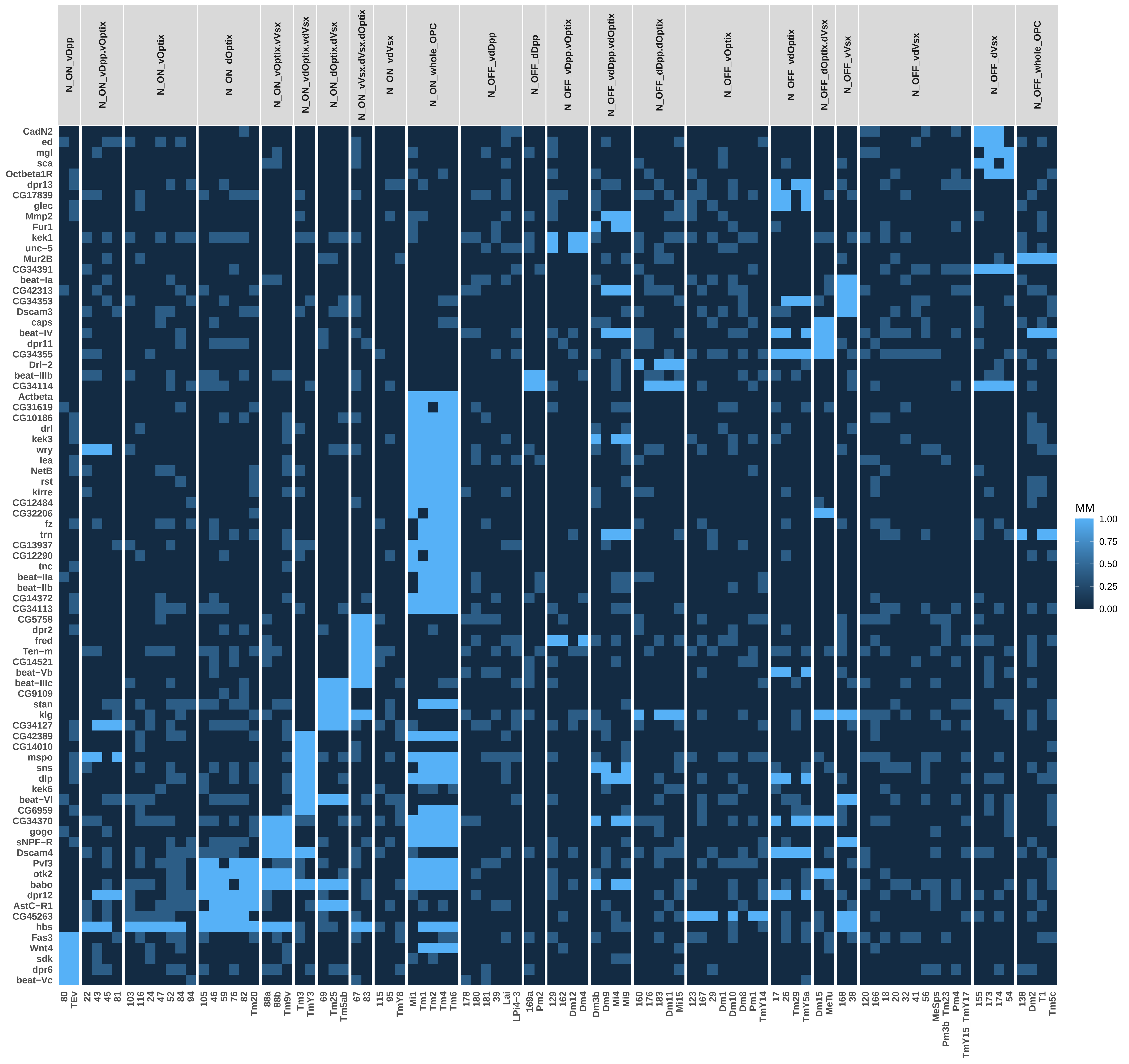

### P15_NO_SO_non_TFs_CSSs_0.75ClusterFreq_0.5BinThreshold_0.5SpecificityThreshold_Simplified.pdf

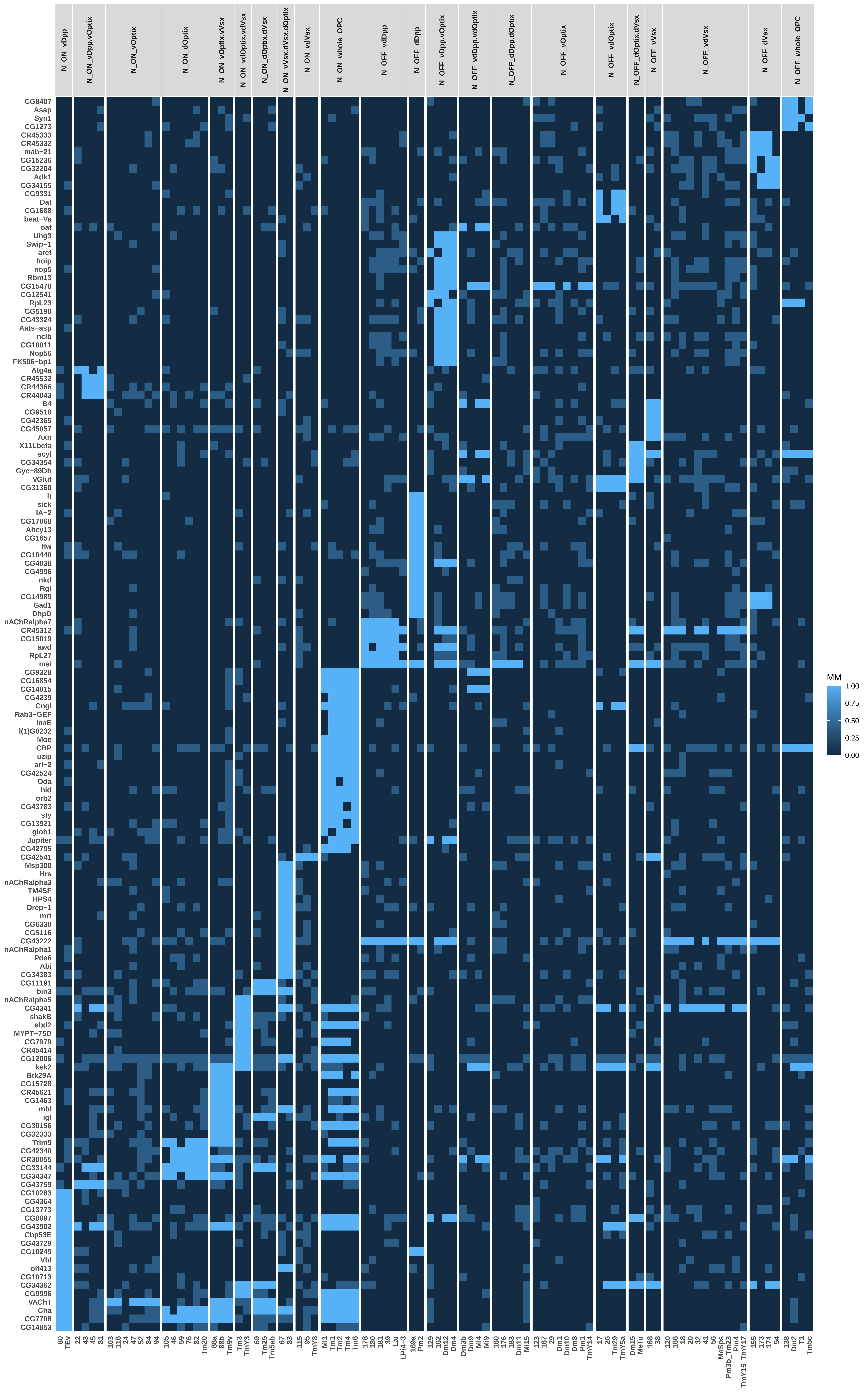

### P15_NO_TO_SO_CSSs_0.75ClusterFreq_0.5BinThreshold_0.5SpecificityThreshold_Simplified.pdf

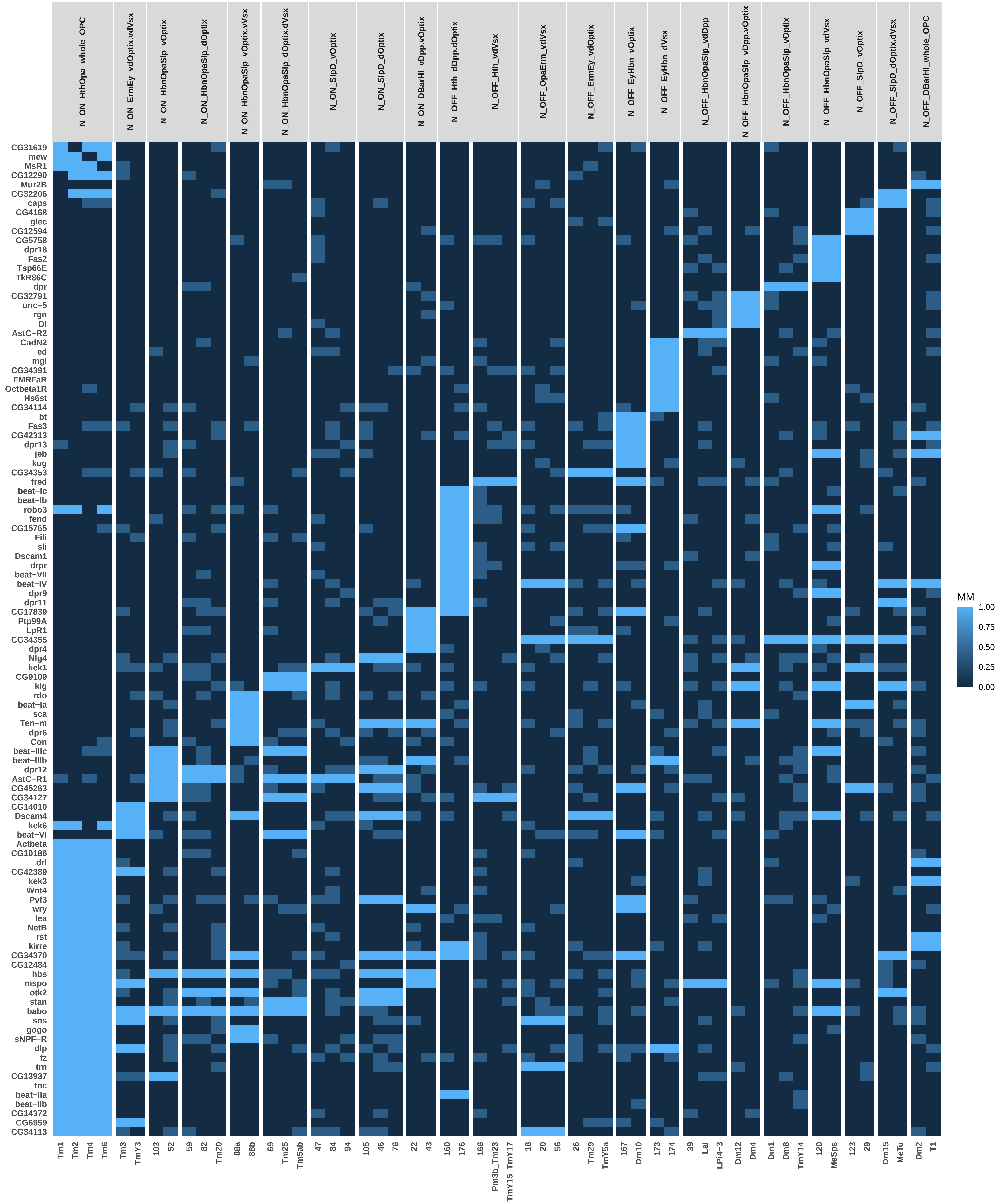

### P15_NO_TO_SO_non_TFs_CSSs_0.75ClusterFreq_0.5BinThreshold_0.5SpecificityThreshold_Simplified.pdf

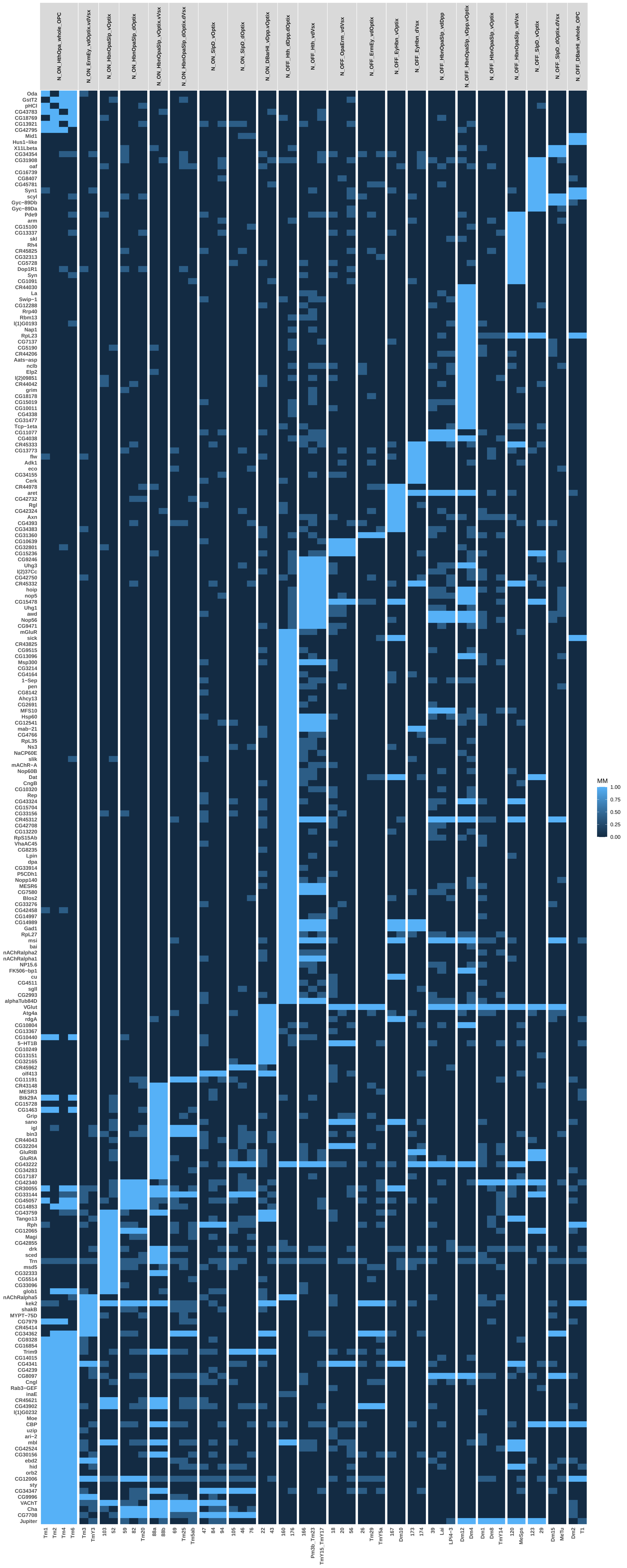

### P15_SO_CSSs_0.75ClusterFreq_0.5BinThreshold_0.5SpecificityThreshold_Simplified.pdf

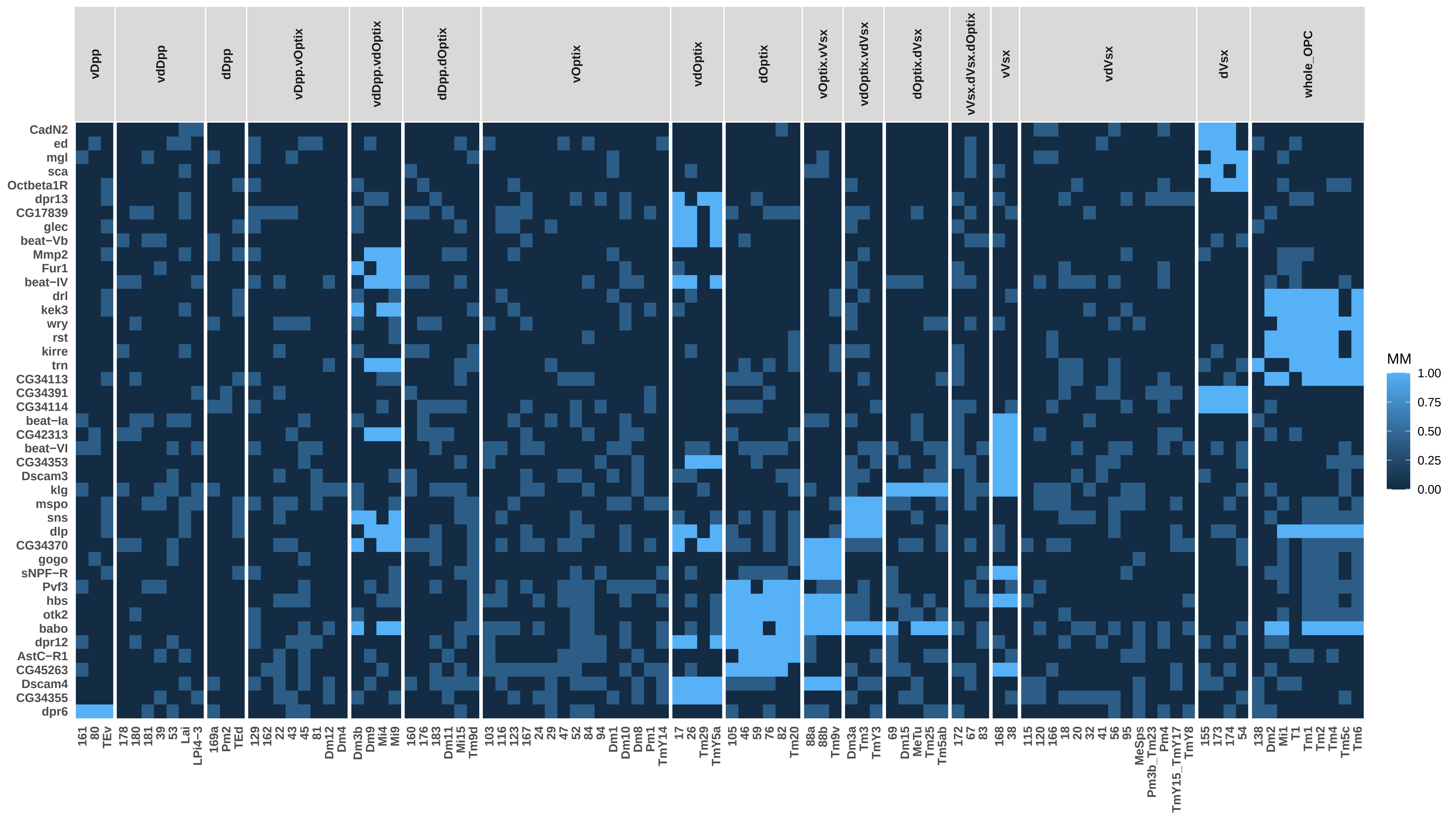

### P15_SO_non_TFs_CSSs_0.75ClusterFreq_0.5BinThreshold_0.5SpecificityThreshold_Simplified.pdf

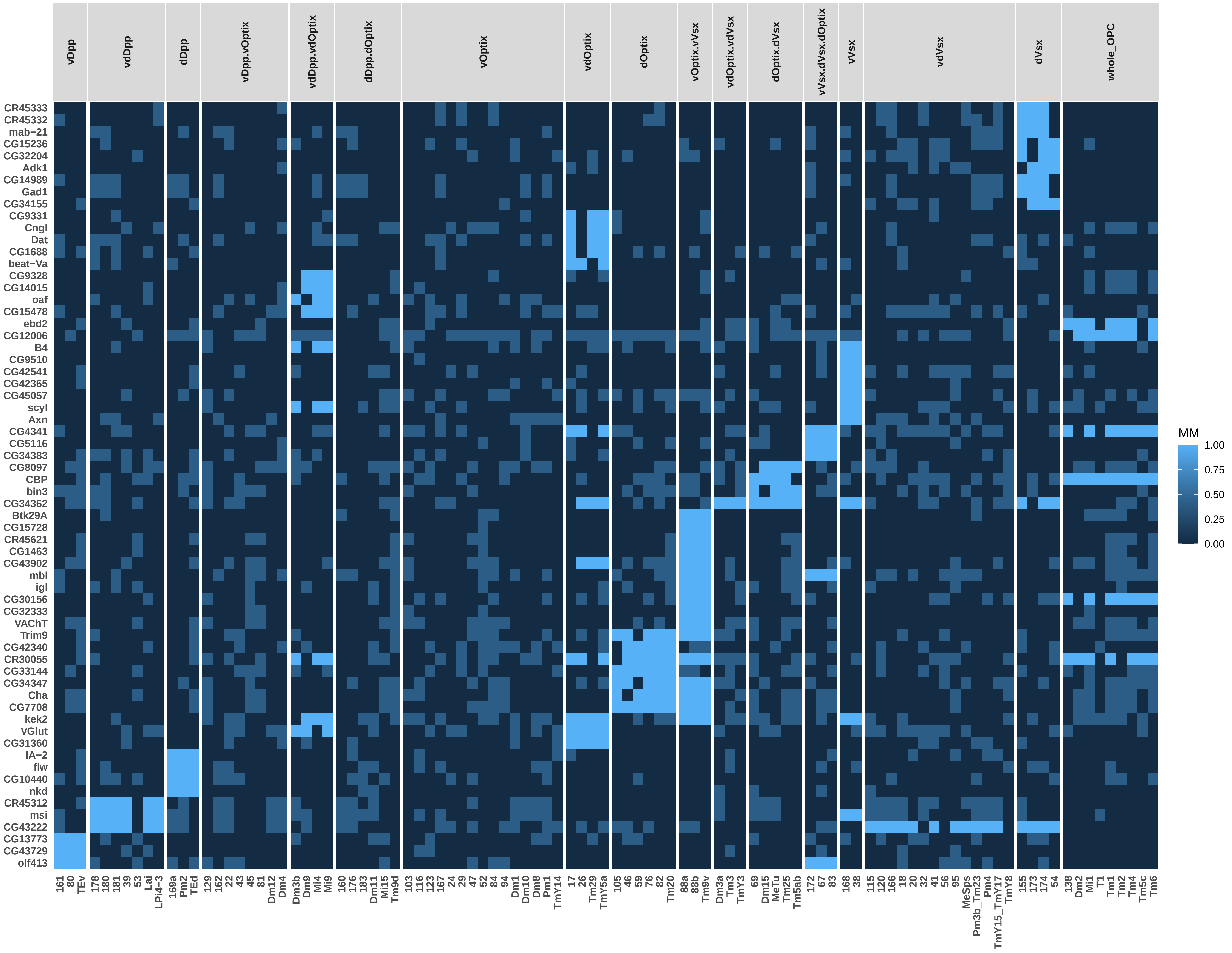

### P15_TO_CSSs_0.75ClusterFreq_0.5BinThreshold_0.5SpecificityThreshold_Simplified.pdf

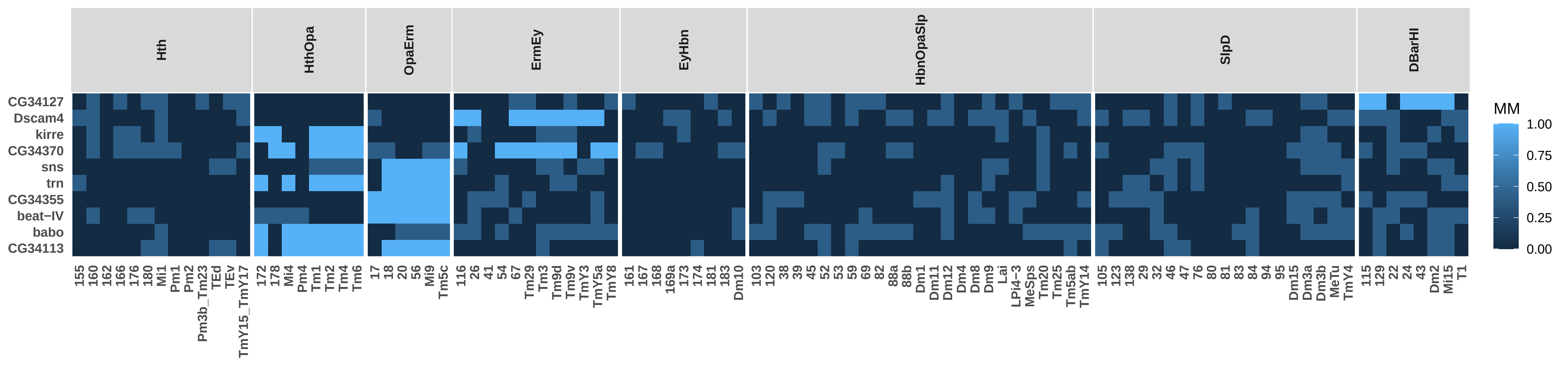

### P15_TO_NO_CSSs_0.75ClusterFreq_0.5BinThreshold_0.5SpecificityThreshold_Simplified.pdf

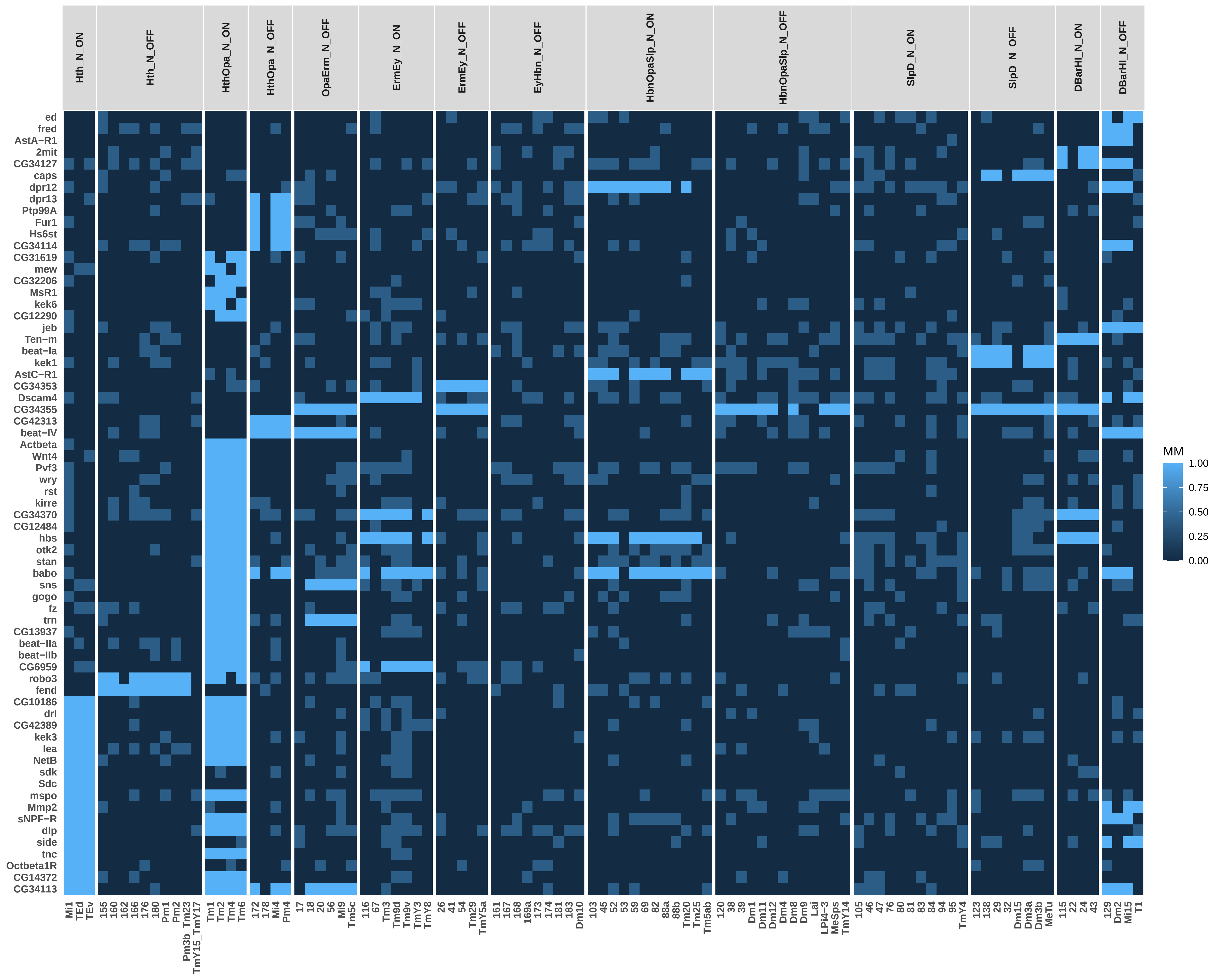

### P15_TO_NO_non_TFs_CSSs_0.75ClusterFreq_0.5BinThreshold_0.5SpecificityThreshold_Simplified.pdf

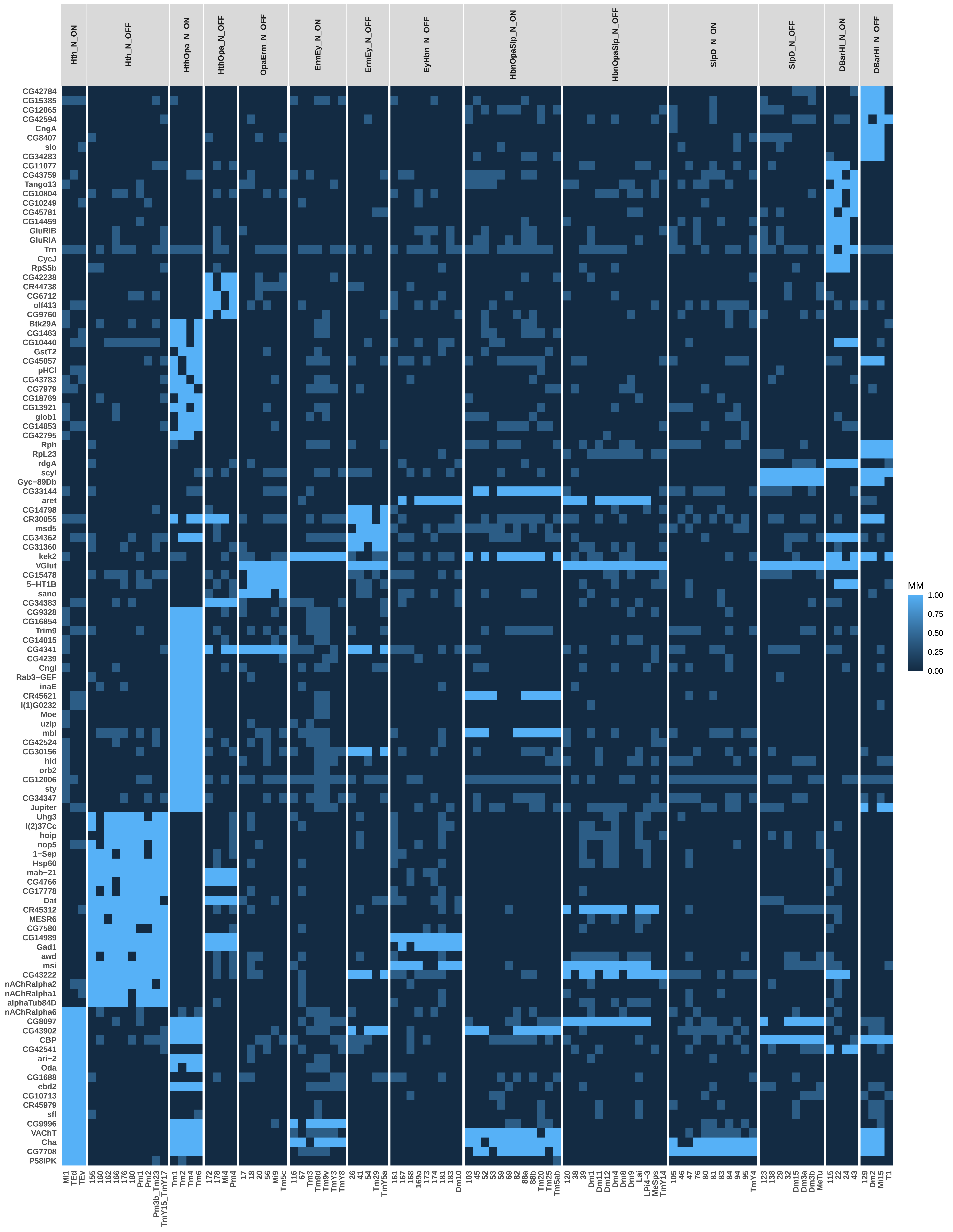

### P15_TO_non_TFs_CSSs_0.75ClusterFreq_0.5BinThreshold_0.5SpecificityThreshold_Simplified.pdf

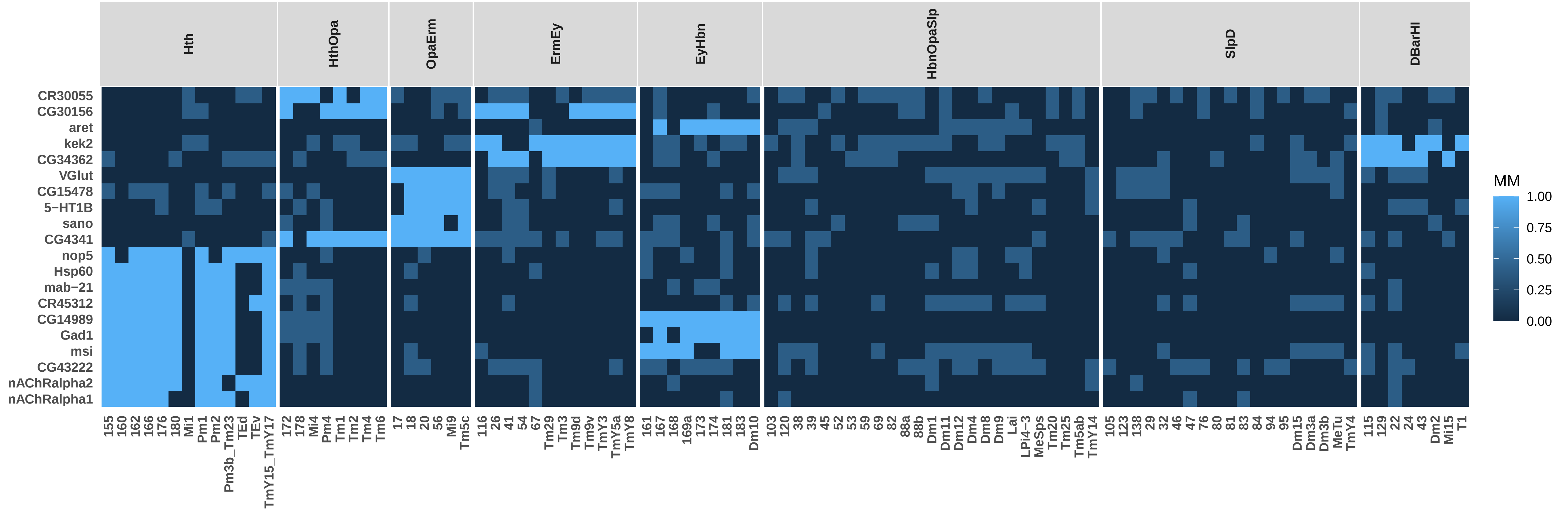

### P15_TO_SO_CSSs_0.75ClusterFreq_0.5BinThreshold_0.5SpecificityThreshold_Simplified.pdf

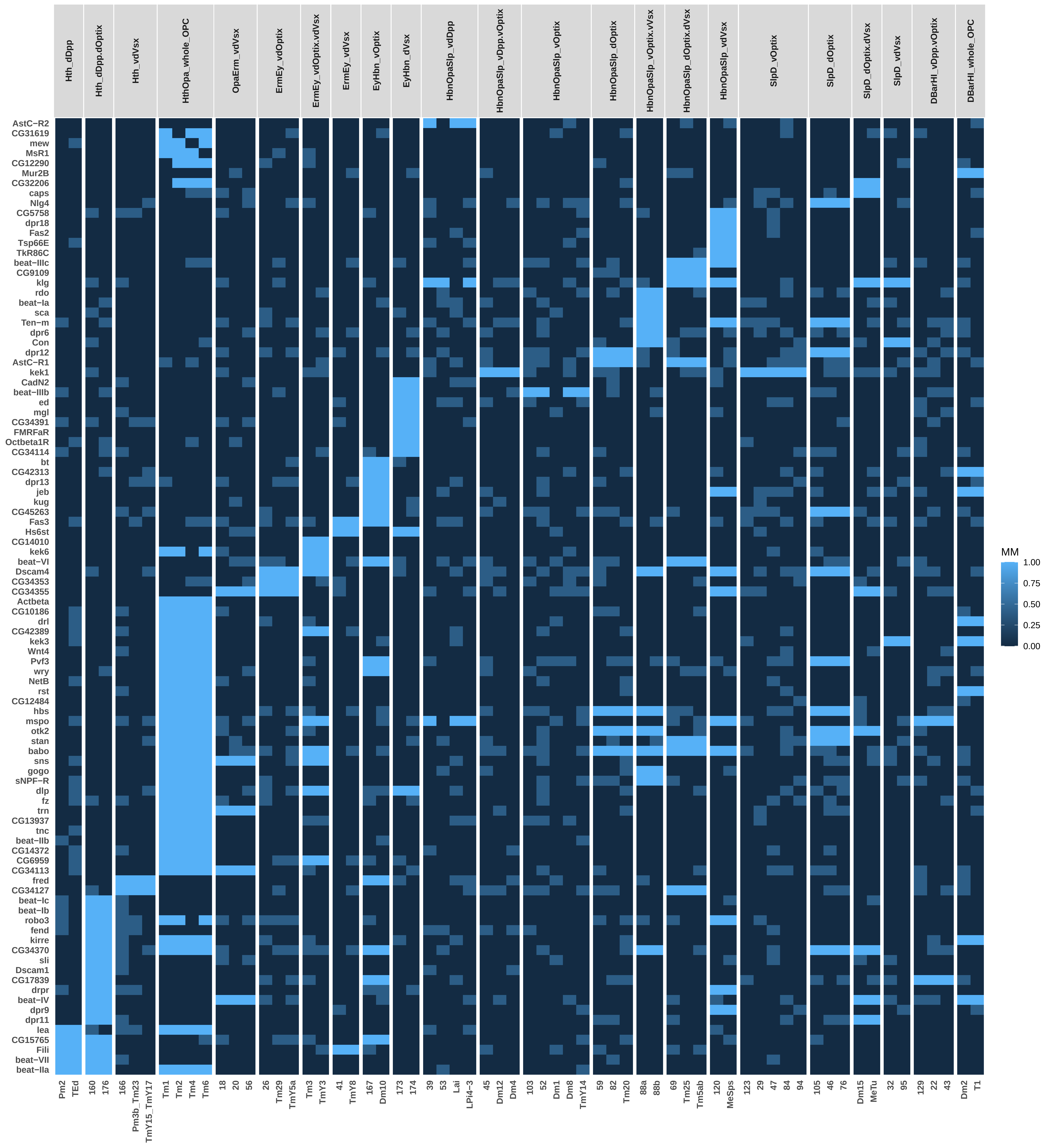

### P15_TO_SO_non_TFs_CSSs_0.75ClusterFreq_0.5BinThreshold_0.5SpecificityThreshold_Simplified.pdf

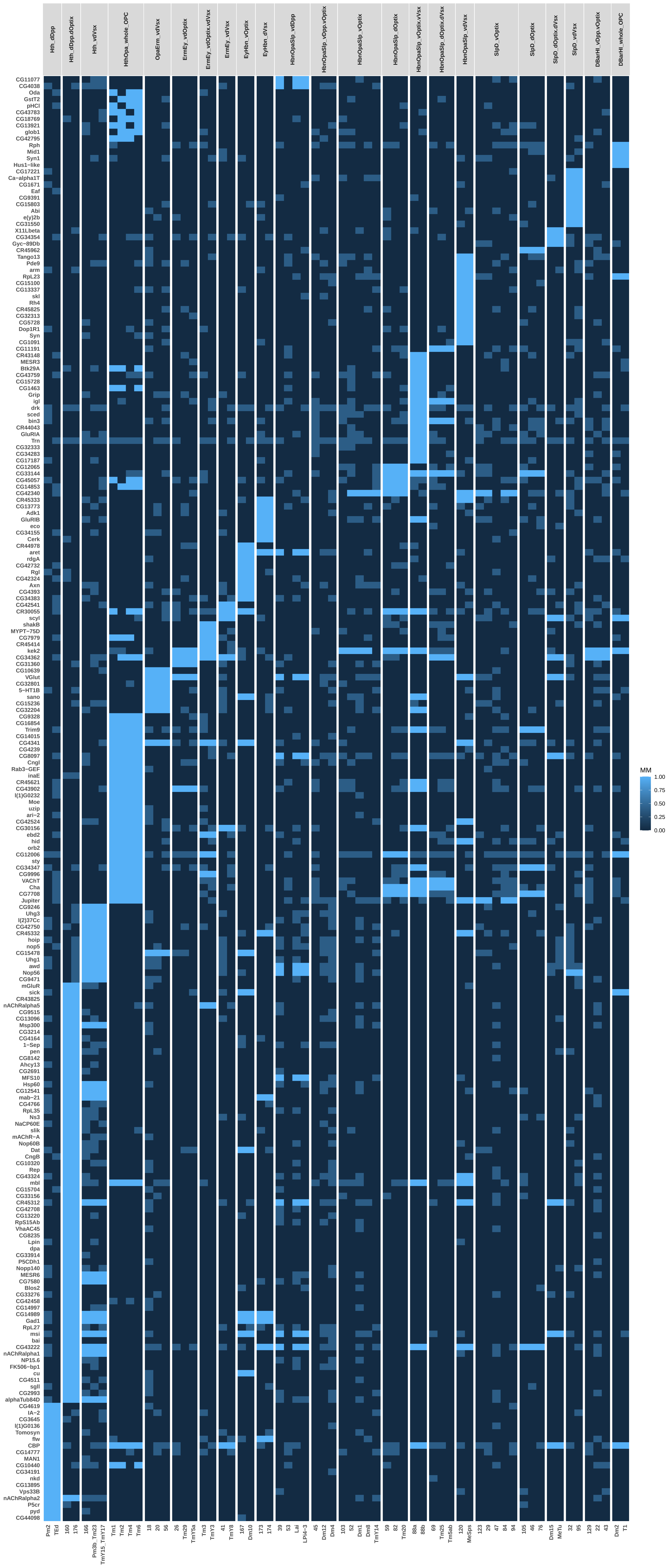

### P30_NO_non_TFs_CSSs_0.75ClusterFreq_0.5BinThreshold_0.5SpecificityThreshold_Simplified.pdf

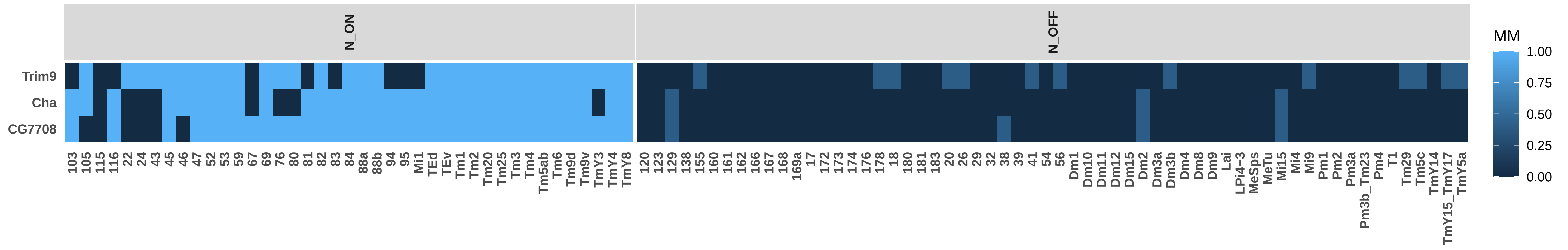

### P30_NO_SO_CSSs_0.75ClusterFreq_0.5BinThreshold_0.5SpecificityThreshold_Simplified.pdf

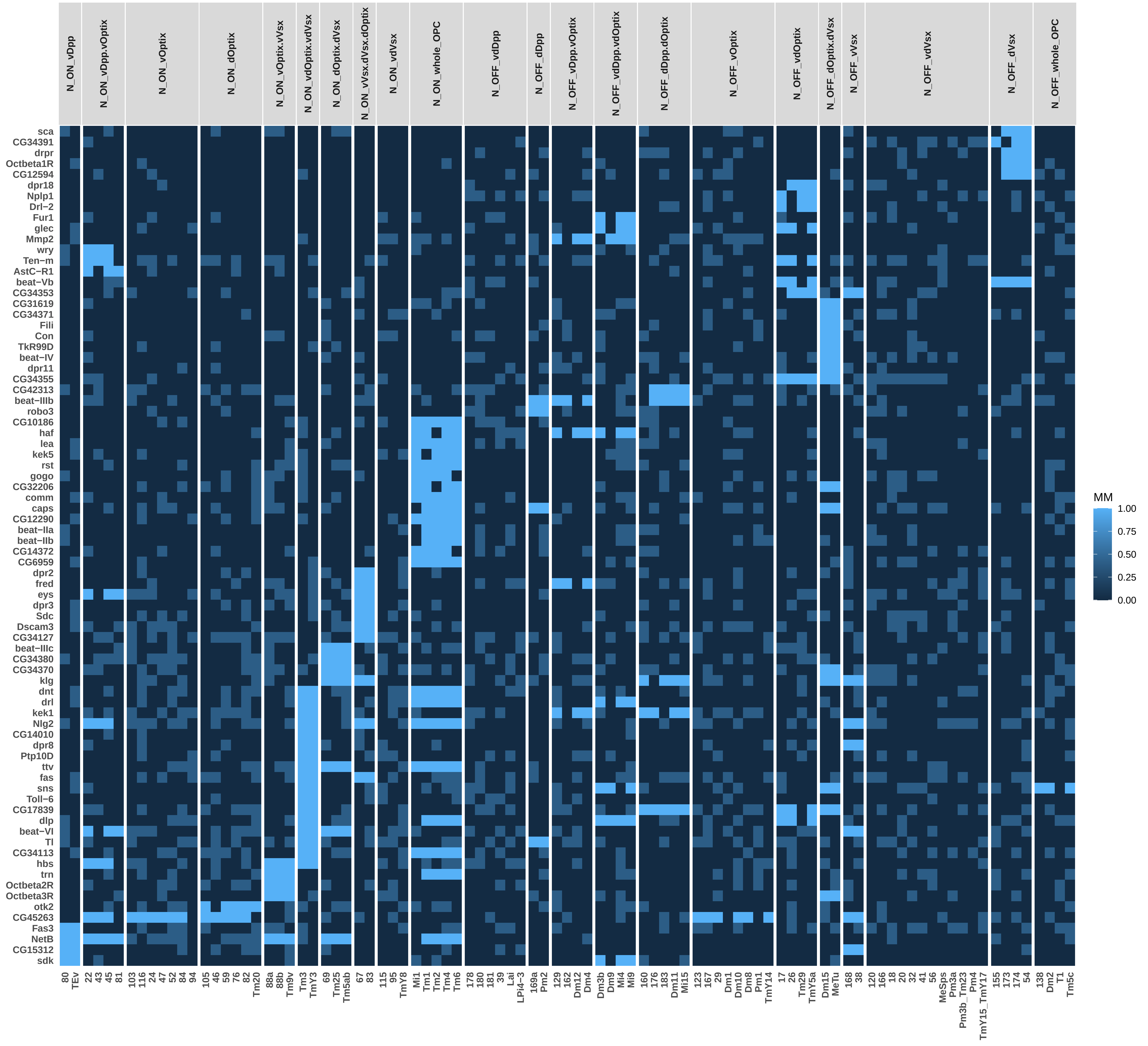

### P30_NO_SO_non_TFs_CSSs_0.75ClusterFreq_0.5BinThreshold_0.5SpecificityThreshold_Simplified.pdf

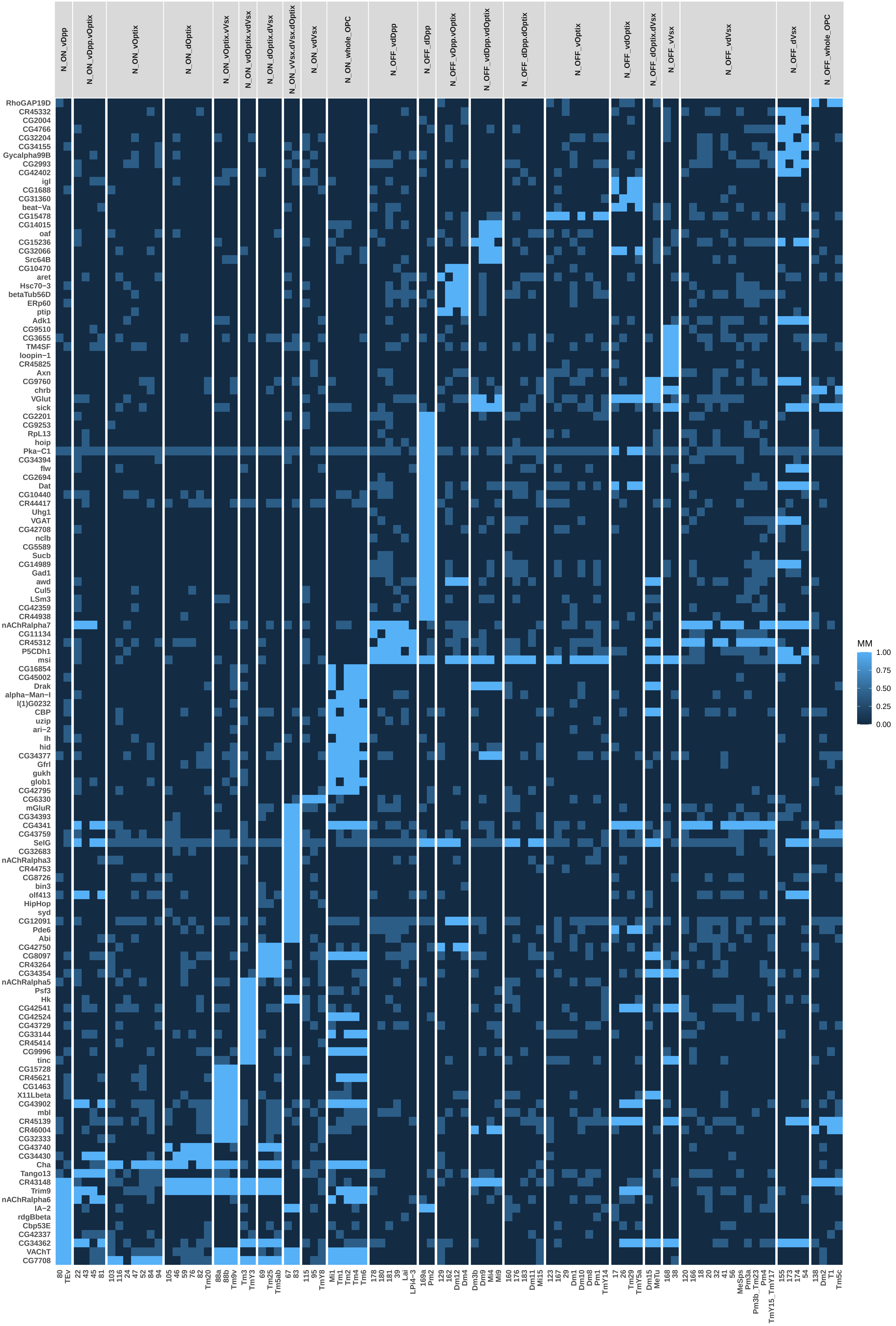

### P30_NO_TO_SO_CSSs_0.75ClusterFreq_0.5BinThreshold_0.5SpecificityThreshold_Simplified.pdf

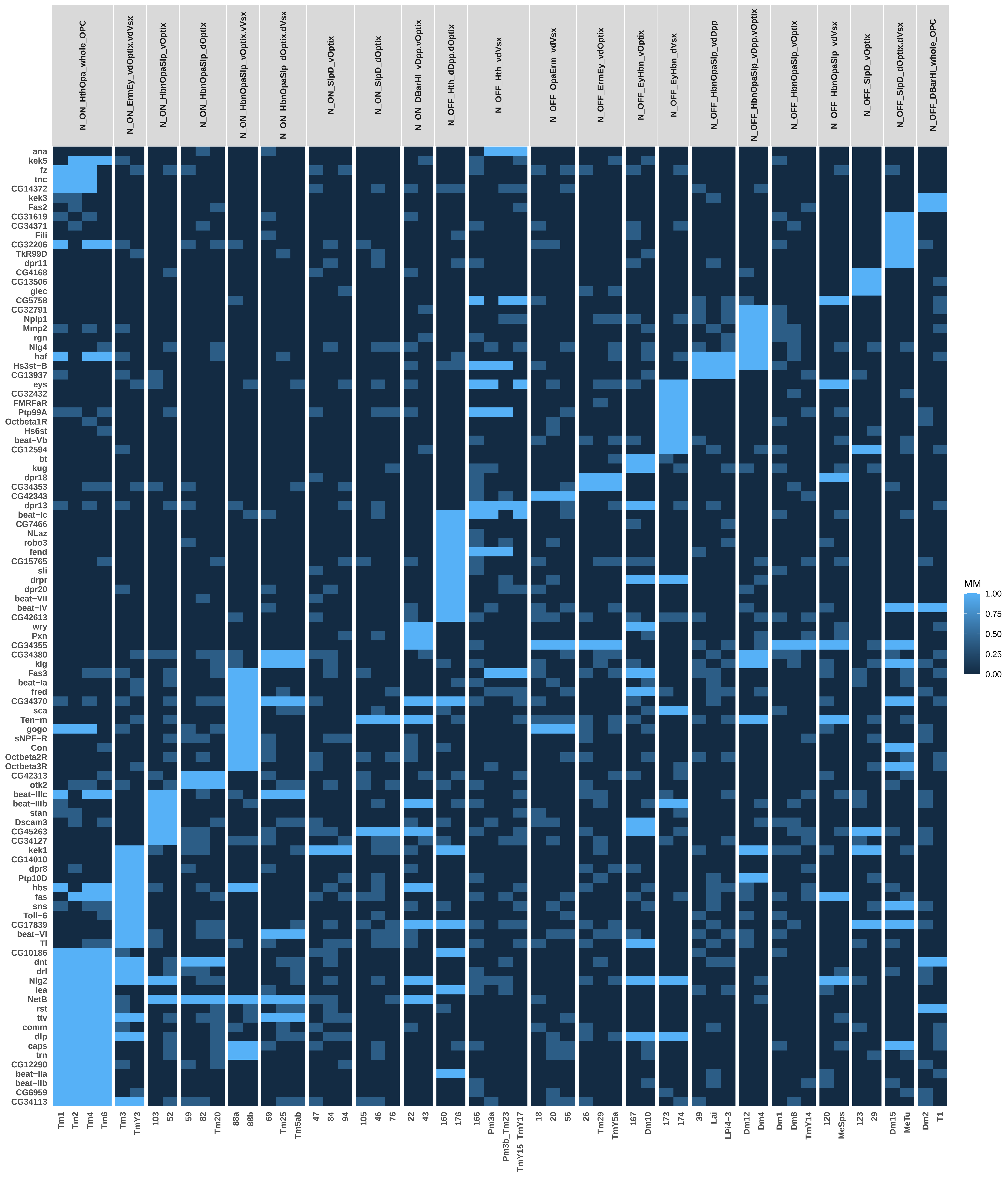

### P30_NO_TO_SO_non_TFs_CSSs_0.75ClusterFreq_0.5BinThreshold_0.5SpecificityThreshold_Simplified.pdf

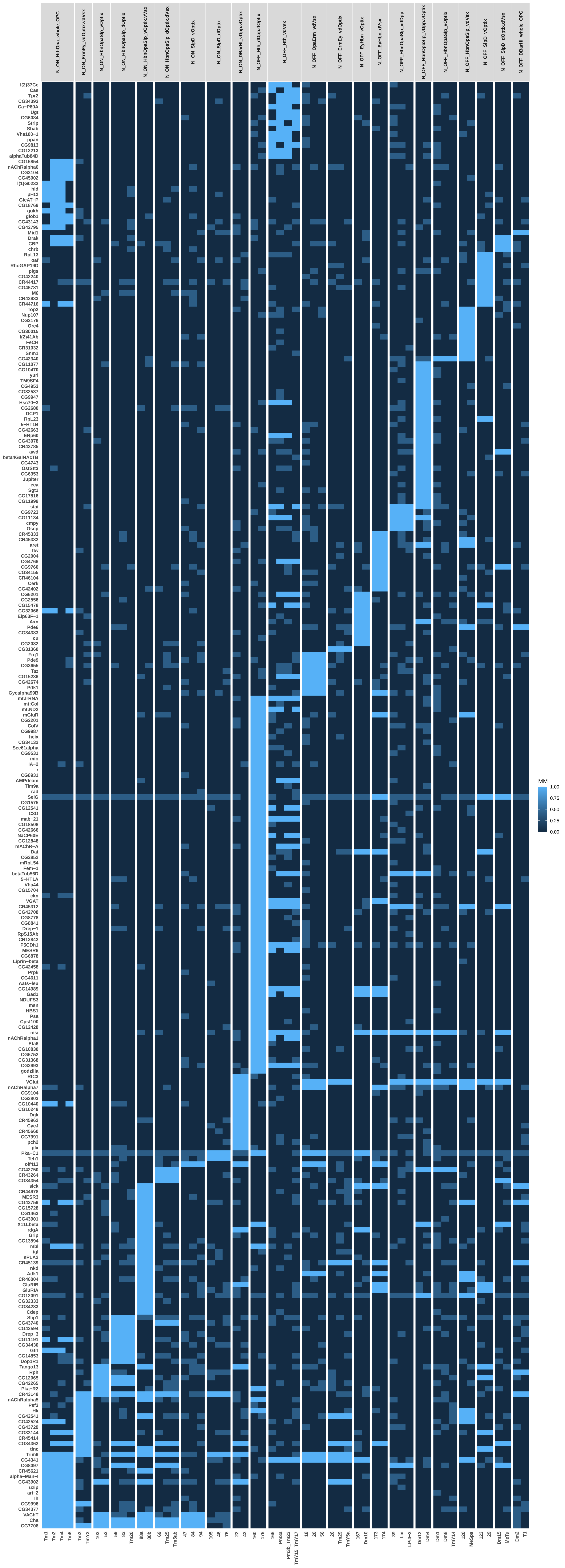

### P30_SO_CSSs_0.75ClusterFreq_0.5BinThreshold_0.5SpecificityThreshold_Simplified.pdf

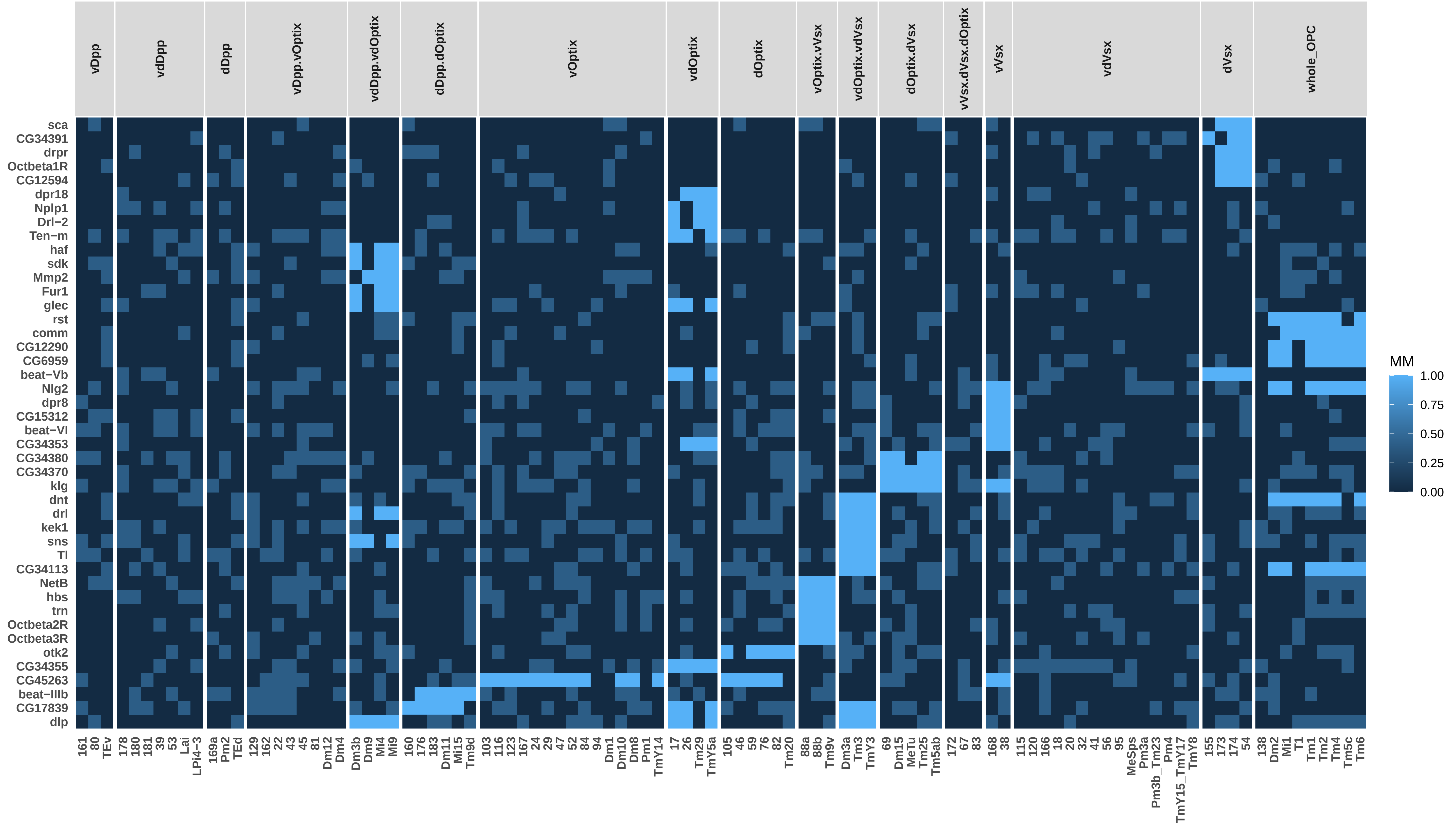

### P30_SO_non_TFs_CSSs_0.75ClusterFreq_0.5BinThreshold_0.5SpecificityThreshold_Simplified.pdf

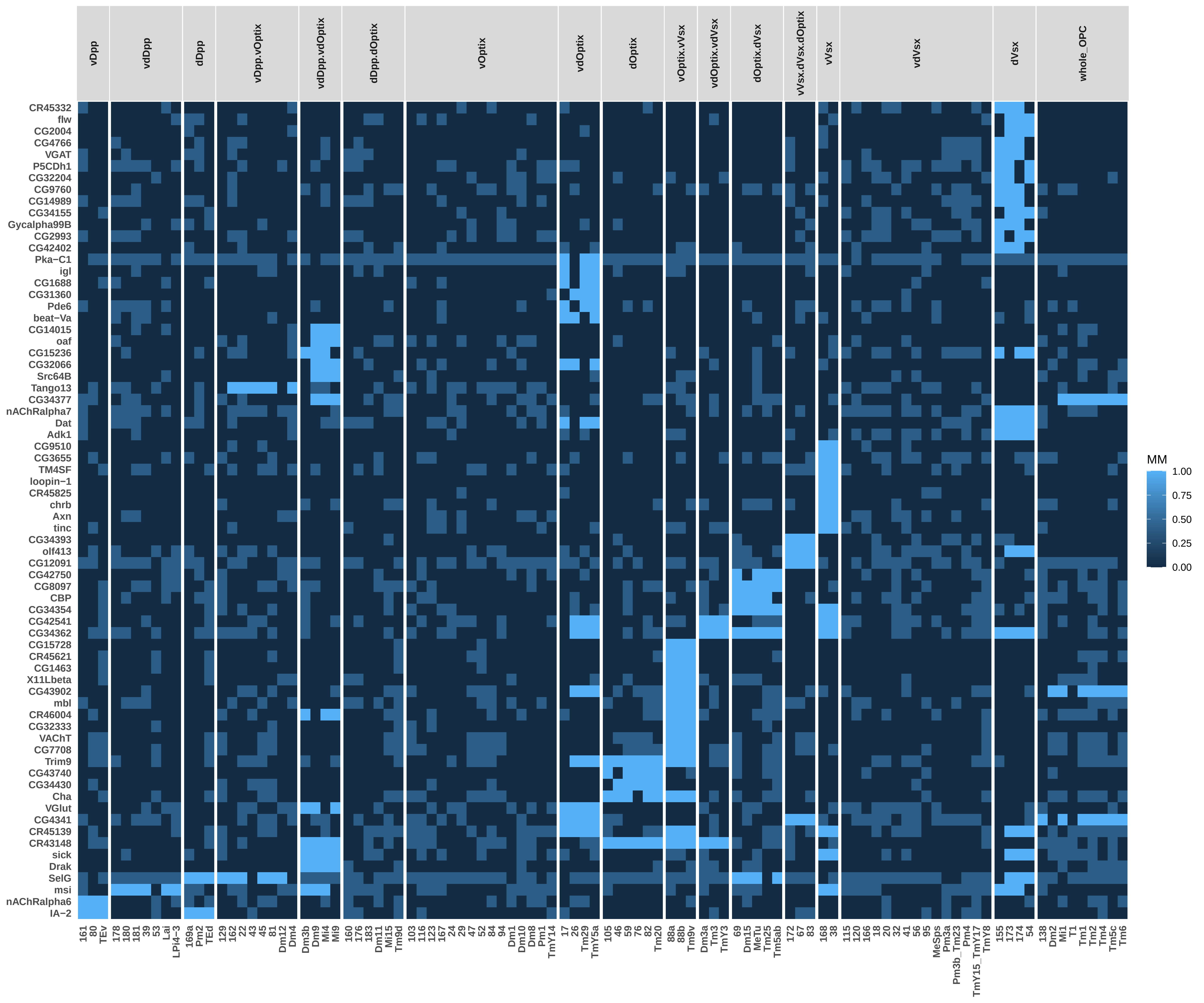

### P30_TO_CSSs_0.75ClusterFreq_0.5BinThreshold_0.5SpecificityThreshold_Simplified.pdf

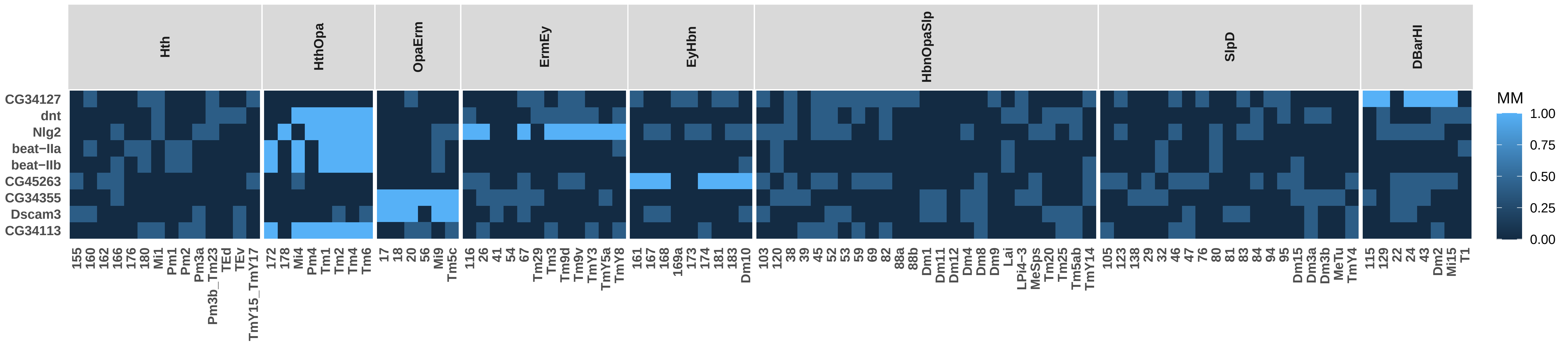

### P30_TO_NO_CSSs_0.75ClusterFreq_0.5BinThreshold_0.5SpecificityThreshold_Simplified.pdf

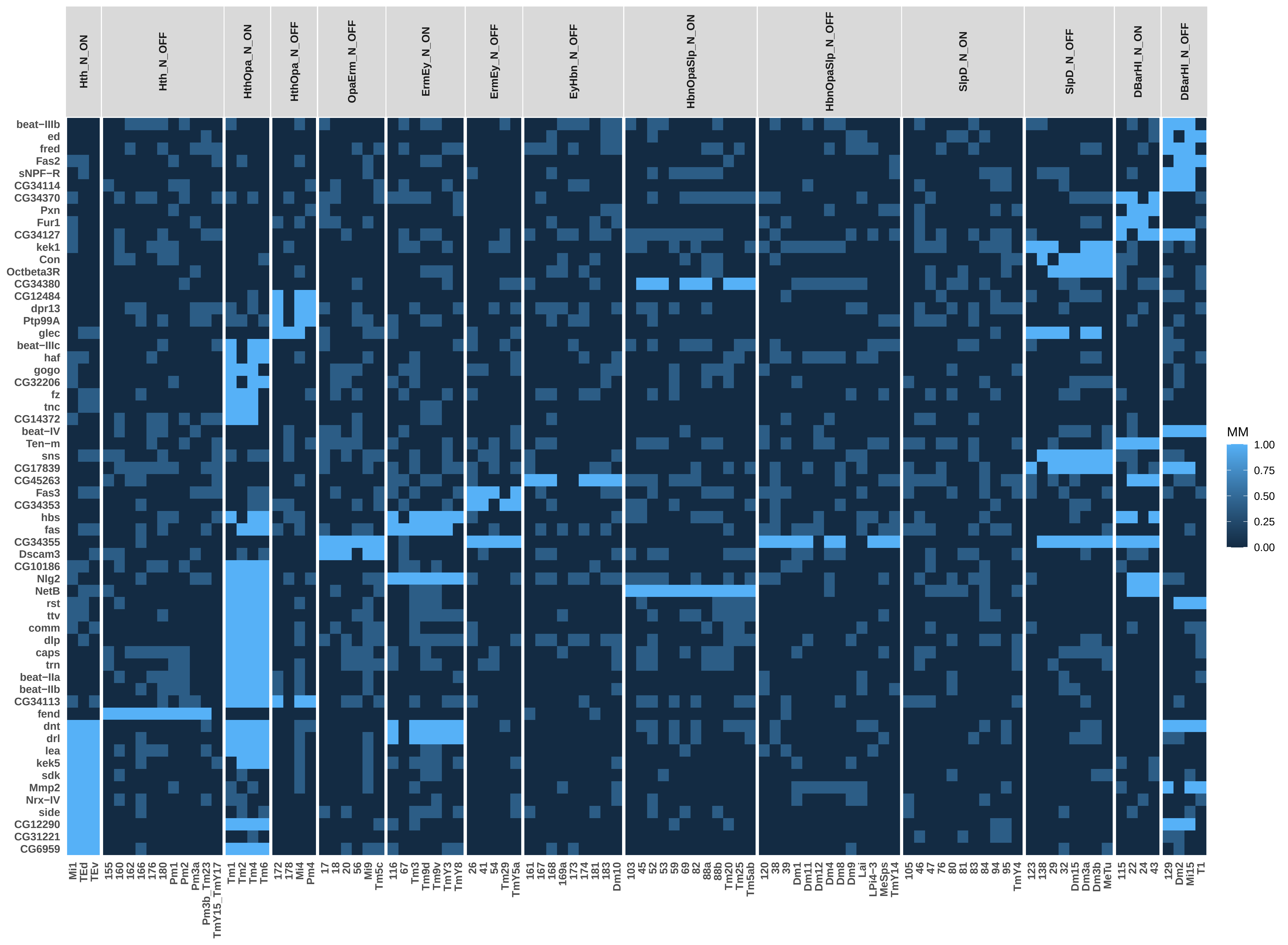

### P30_TO_NO_non_TFs_CSSs_0.75ClusterFreq_0.5BinThreshold_0.5SpecificityThreshold_Simplified.pdf

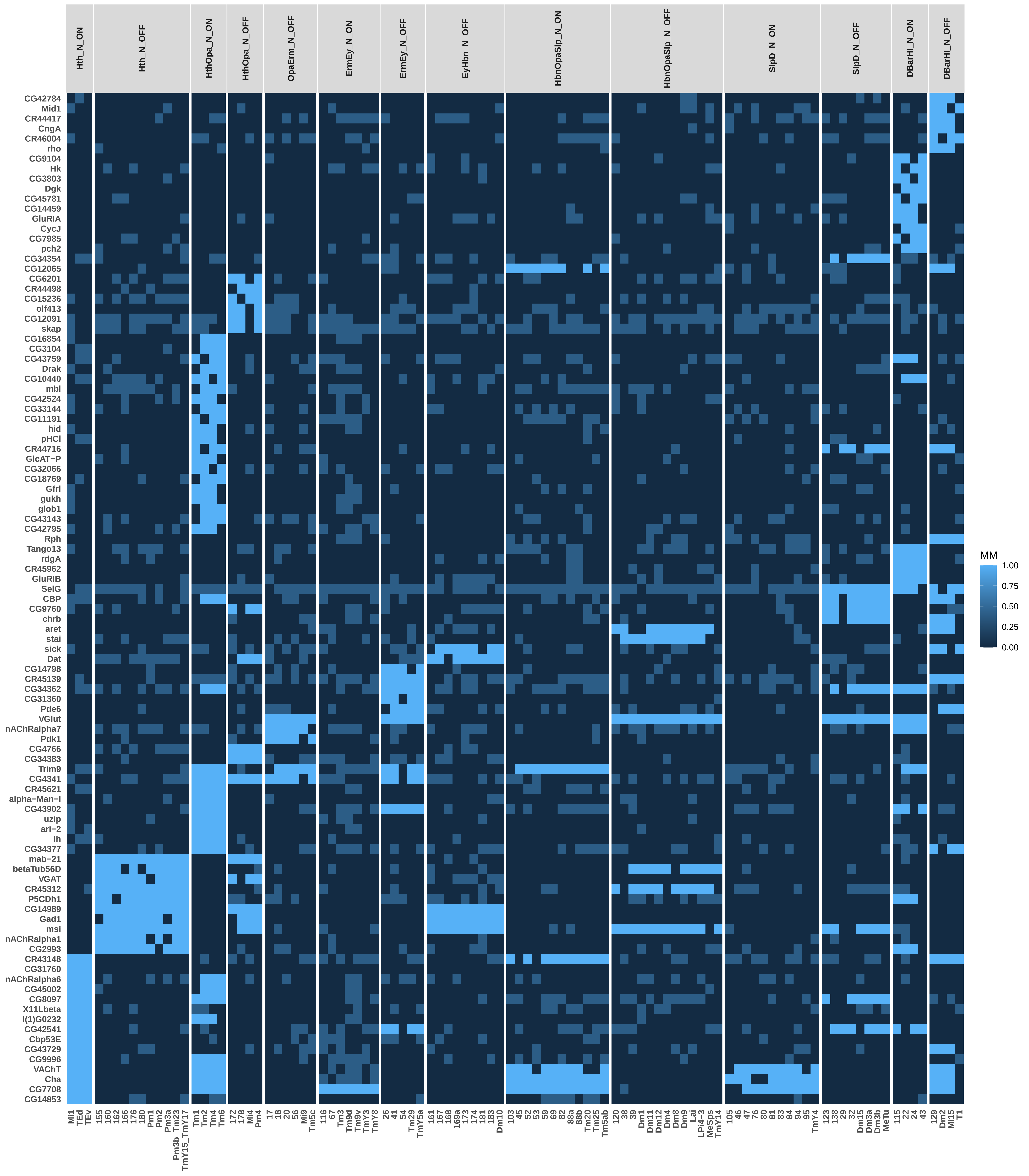

### P30_TO_non_TFs_CSSs_0.75ClusterFreq_0.5BinThreshold_0.5SpecificityThreshold_Simplified.pdf

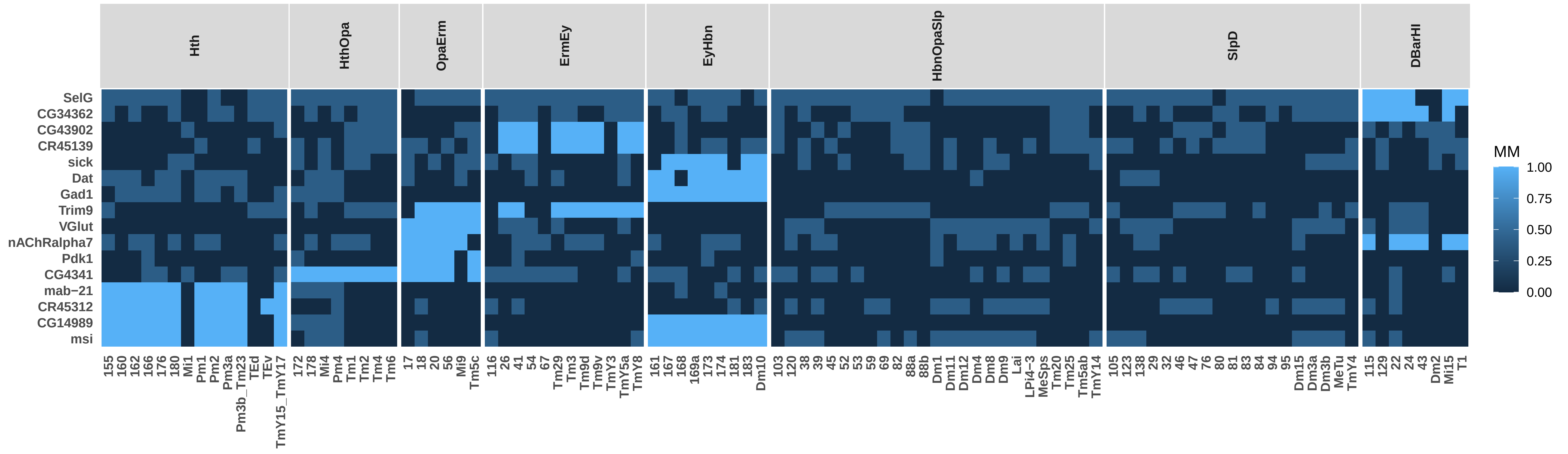

### P30_TO_SO_CSSs_0.75ClusterFreq_0.5BinThreshold_0.5SpecificityThreshold_Simplified.pdf

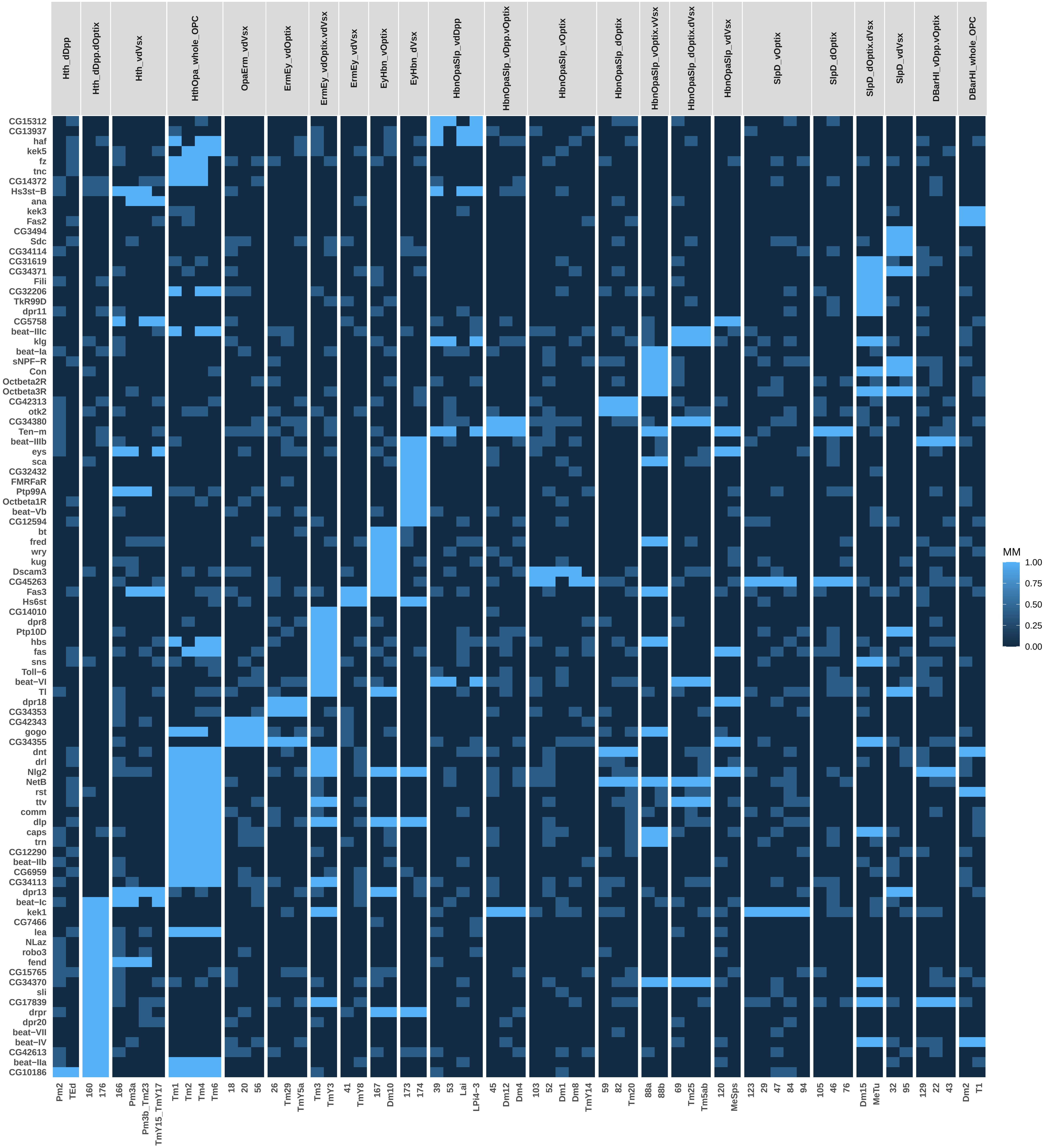

### P30_TO_SO_non_TFs_CSSs_0.75ClusterFreq_0.5BinThreshold_0.5SpecificityThreshold_Simplified.pdf

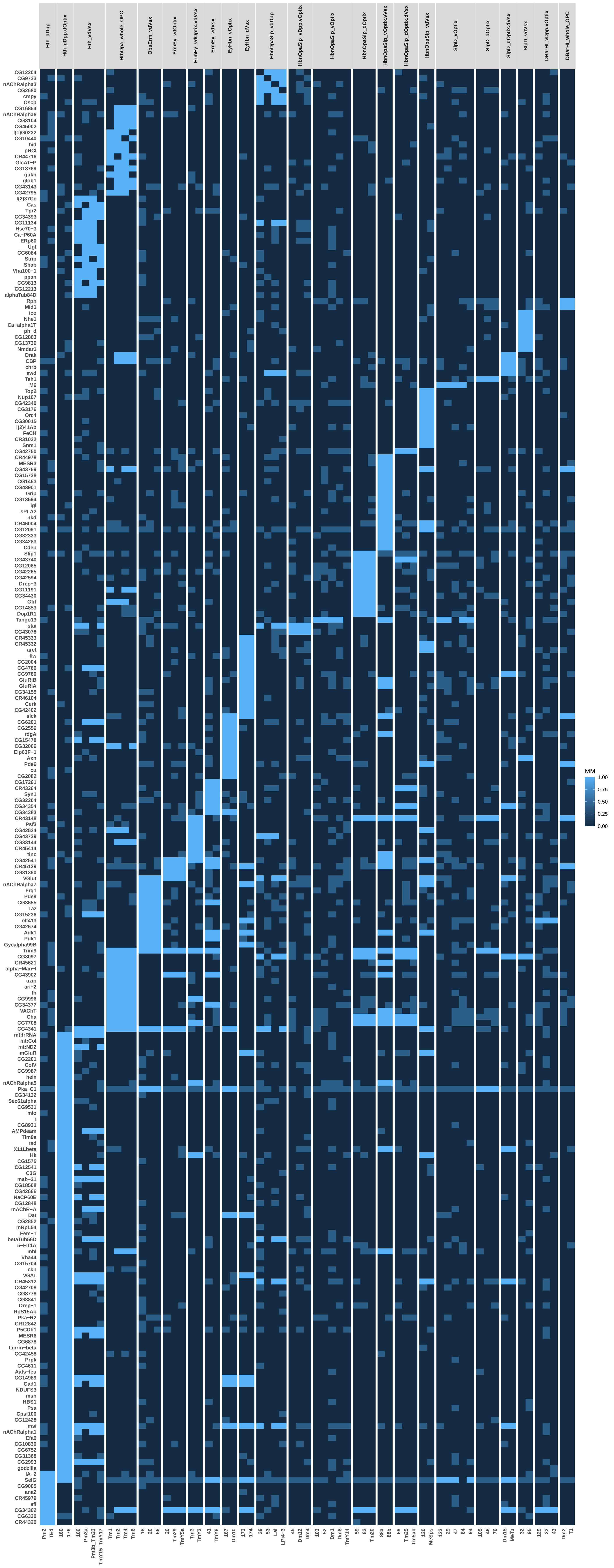

### P40_NO_non_TFs_CSSs_0.75ClusterFreq_0.5BinThreshold_0.5SpecificityThreshold_Simplified.pdf

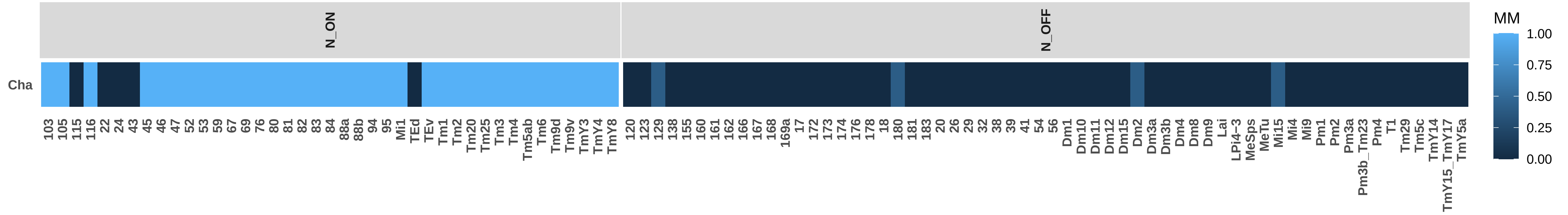

### P40_NO_SO_CSSs_0.75ClusterFreq_0.5BinThreshold_0.5SpecificityThreshold_Simplified.pdf

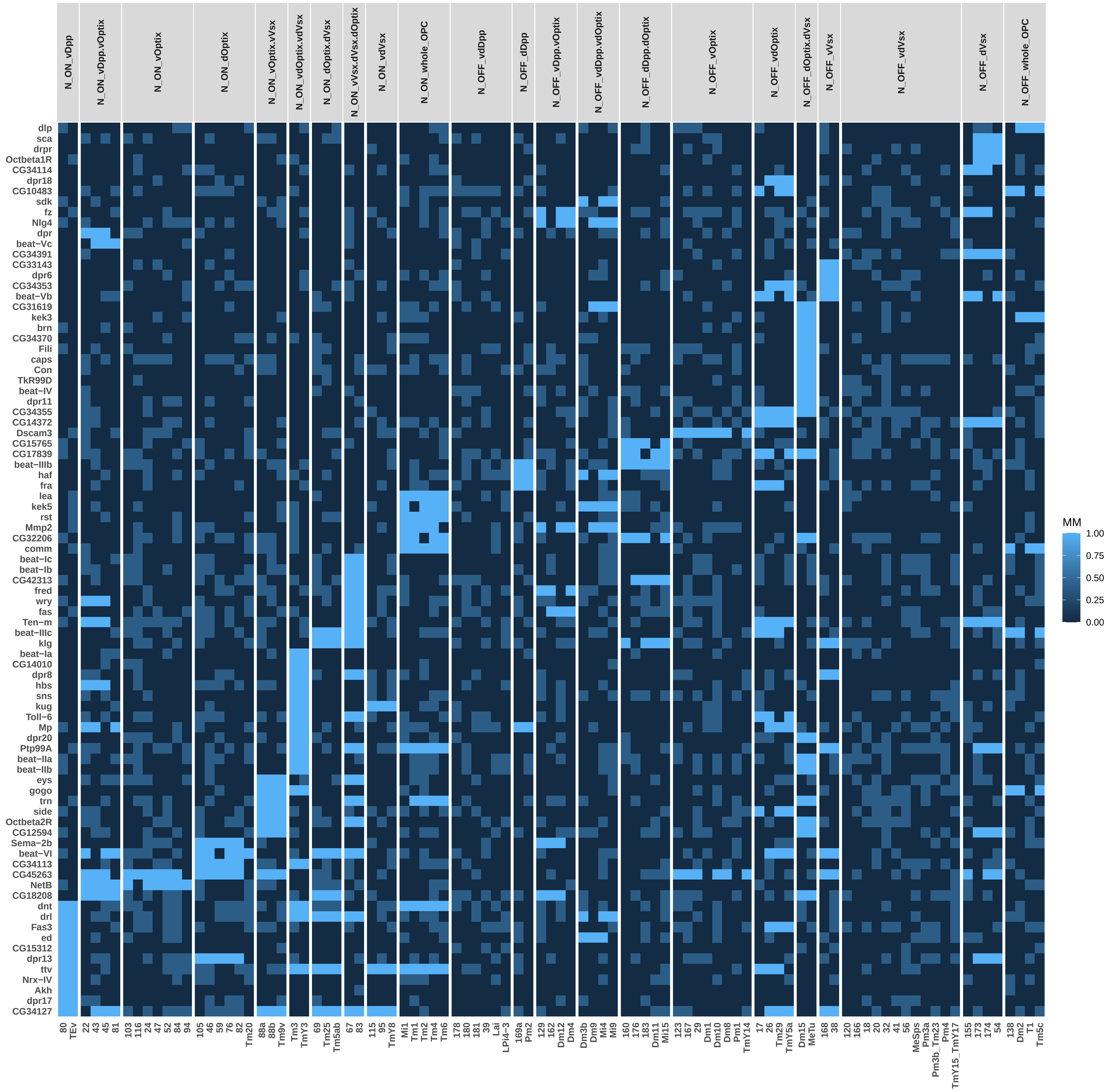

### P40_NO_SO_non_TFs_CSSs_0.75ClusterFreq_0.5BinThreshold_0.5SpecificityThreshold_Simplified.pdf

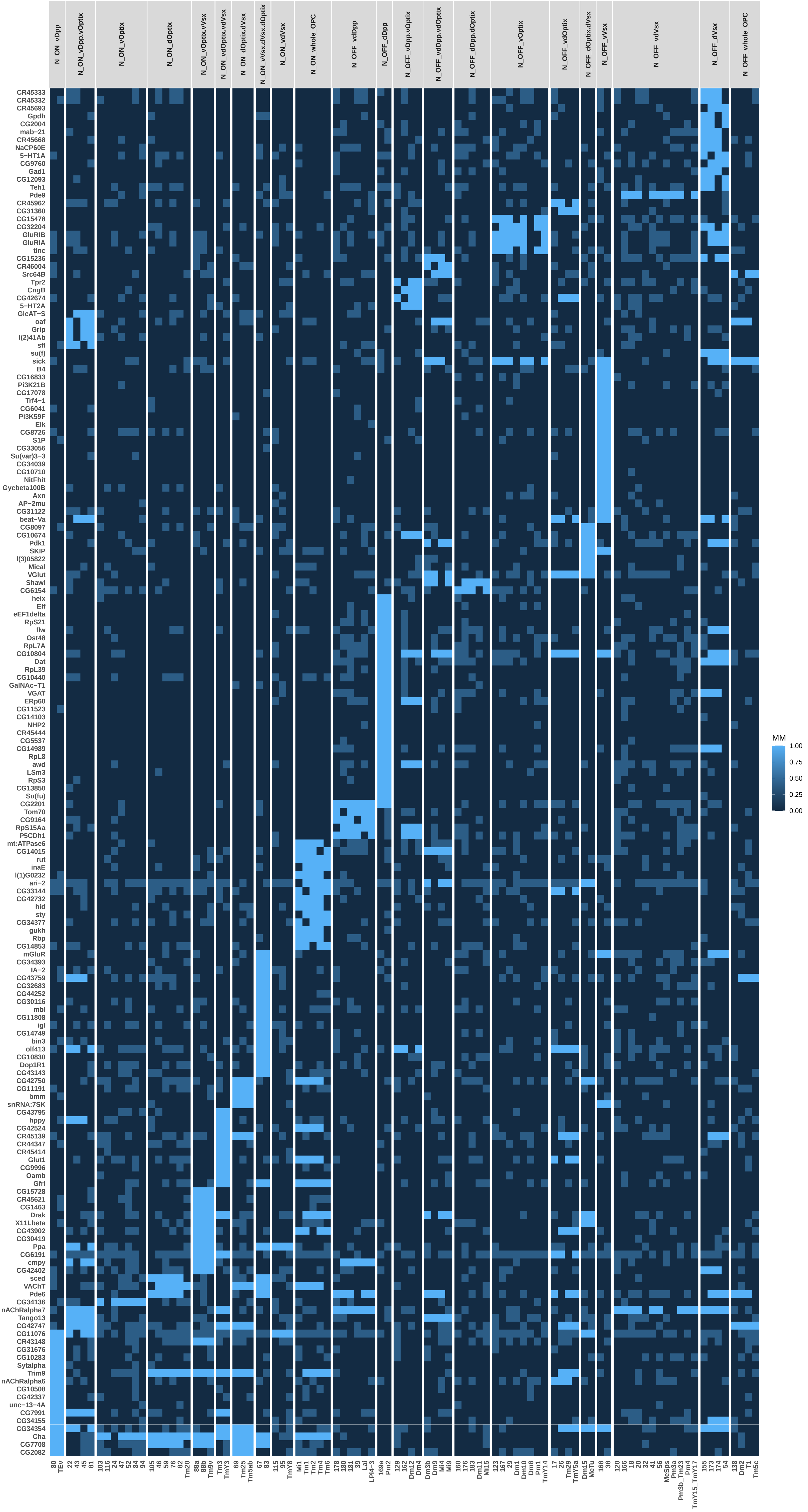

### P40_NO_SO_TFs_0.75ClusterFreq_0.5BinThreshold_0.5SpecificityThreshold_Simplified.pdf

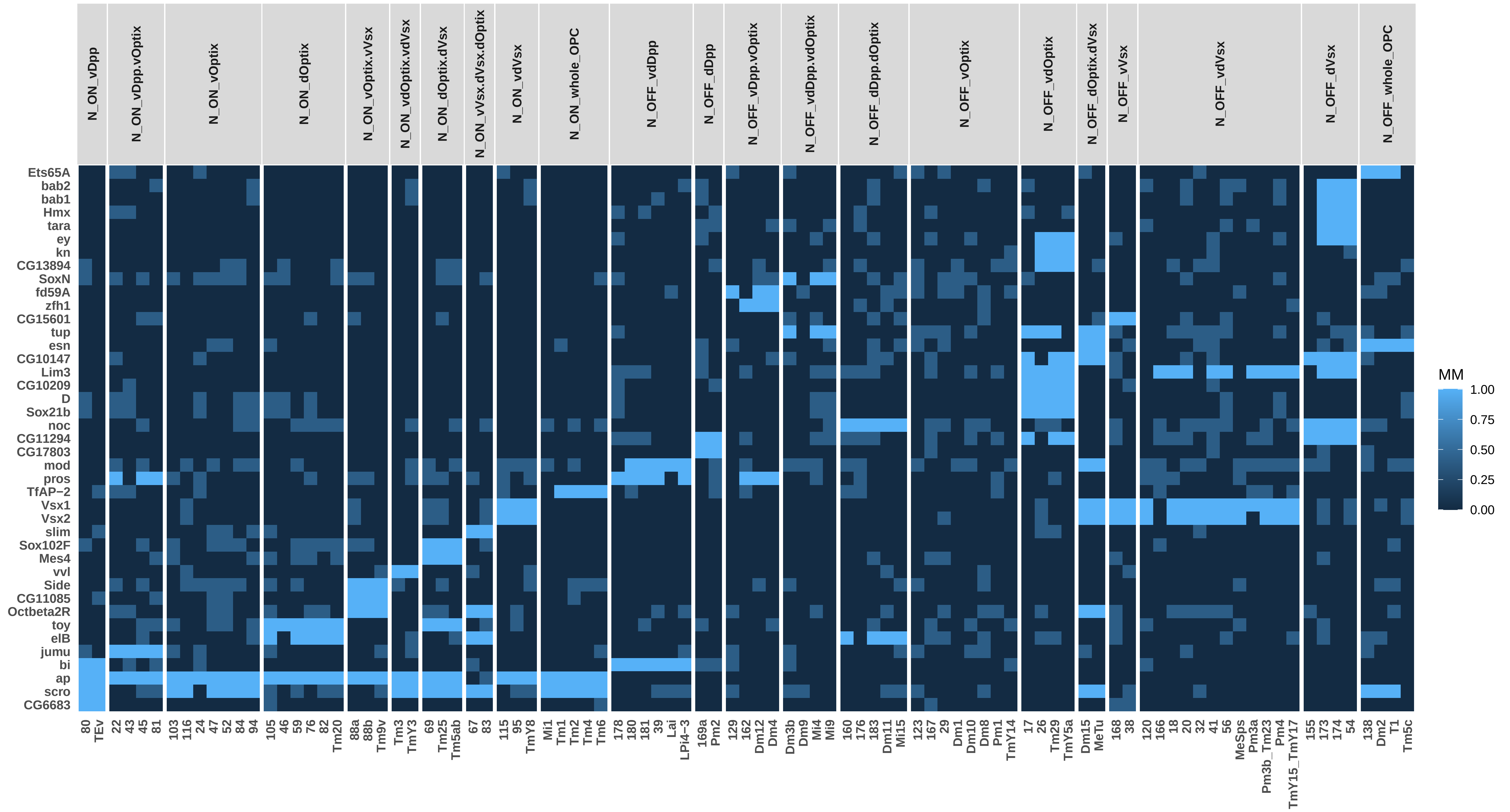

### P40_NO_TFs_0.75ClusterFreq_0.5BinThreshold_0.5SpecificityThreshold_Simplified.pdf

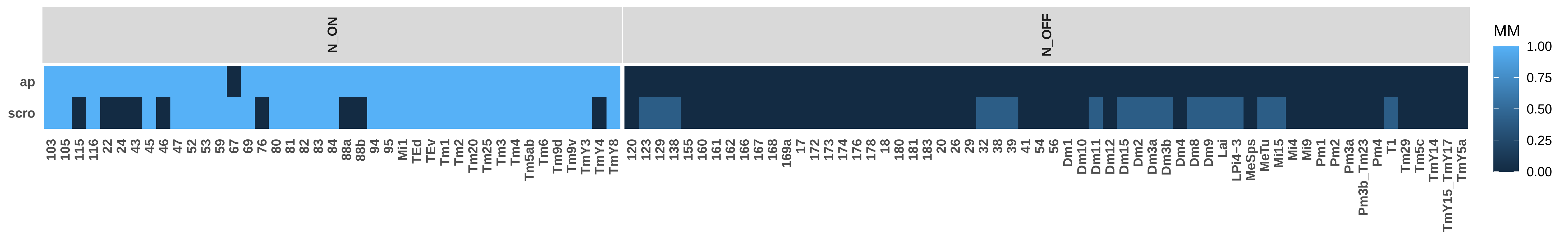

### P40_NO_TO_SO_CSSs_0.75ClusterFreq_0.5BinThreshold_0.5SpecificityThreshold_Simplified.pdf

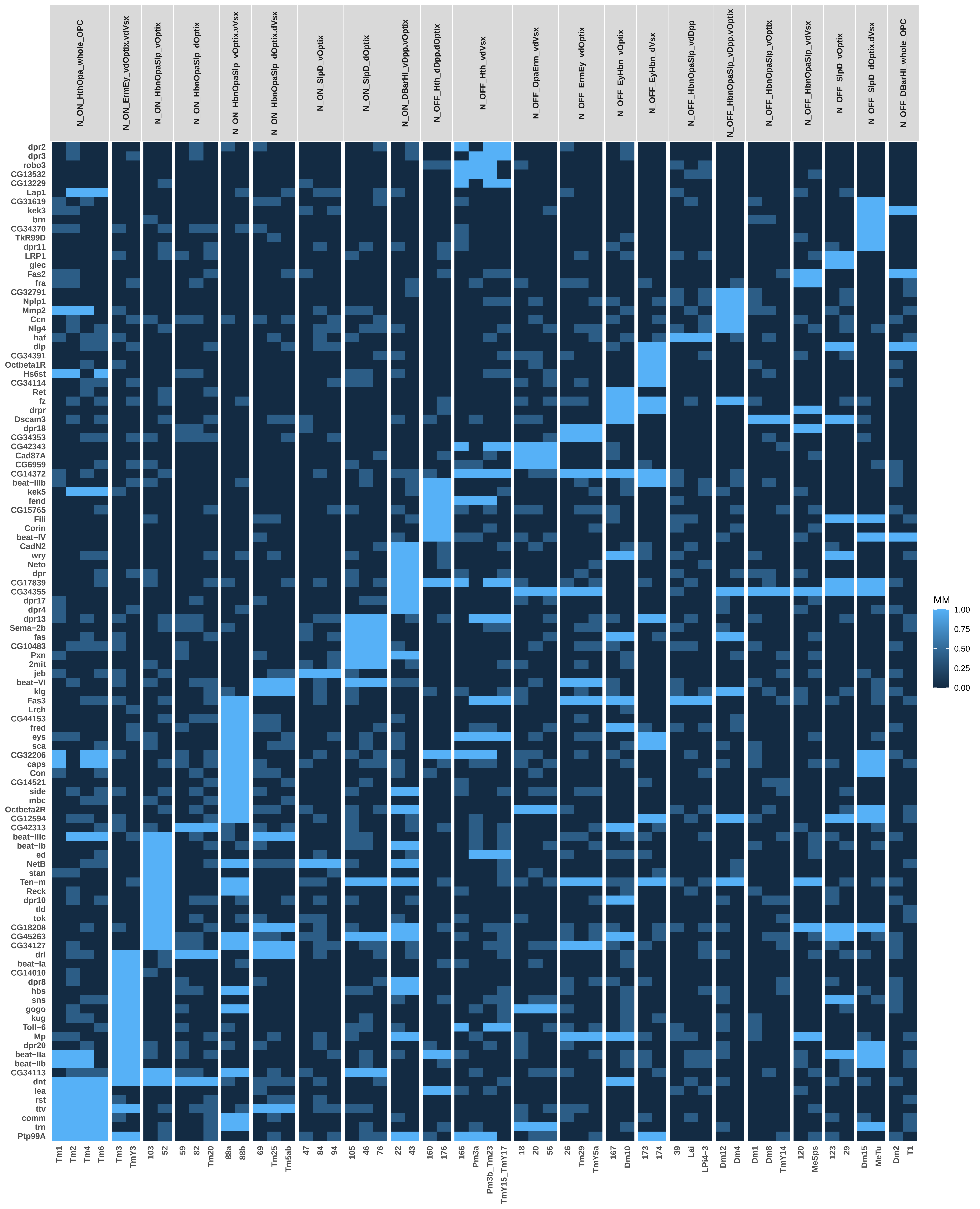

### P40_NO_TO_SO_non_TFs_CSSs_0.75ClusterFreq_0.5BinThreshold_0.5SpecificityThreshold_Simplified.pdf

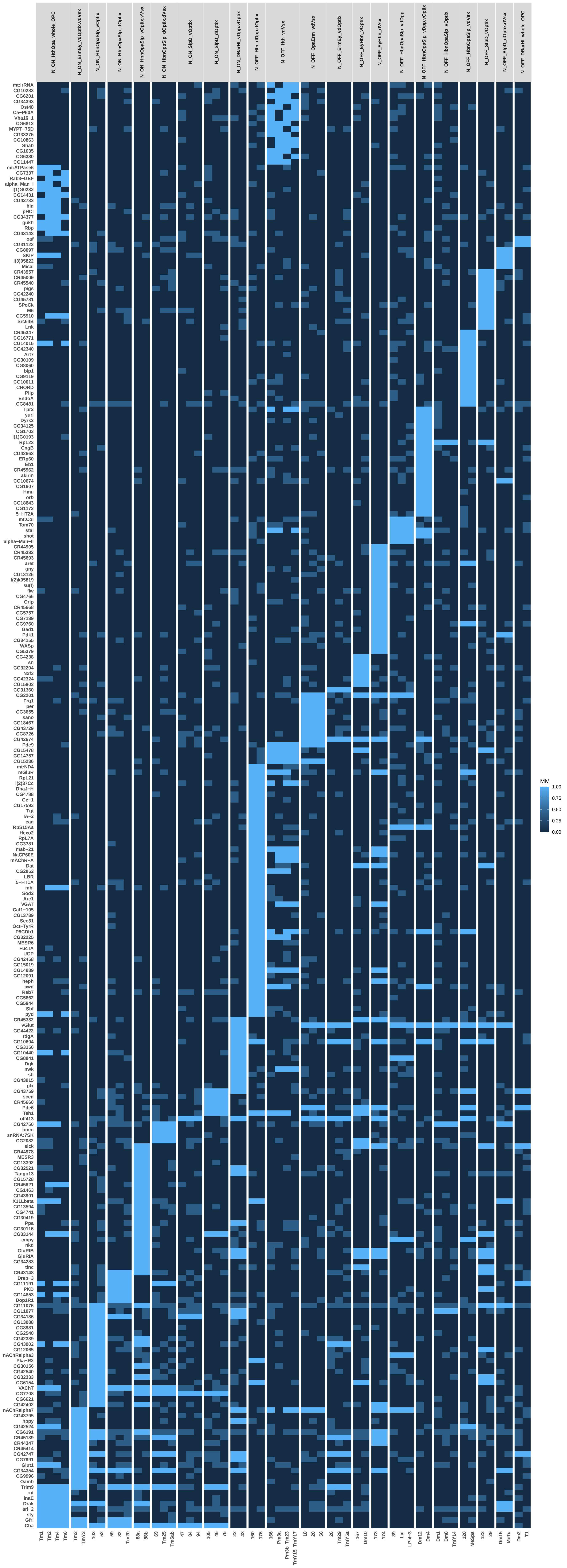

### P40_NO_TO_SO_TFs_0.75ClusterFreq_0.5BinThreshold_0.5SpecificityThreshold_Simplified.pdf

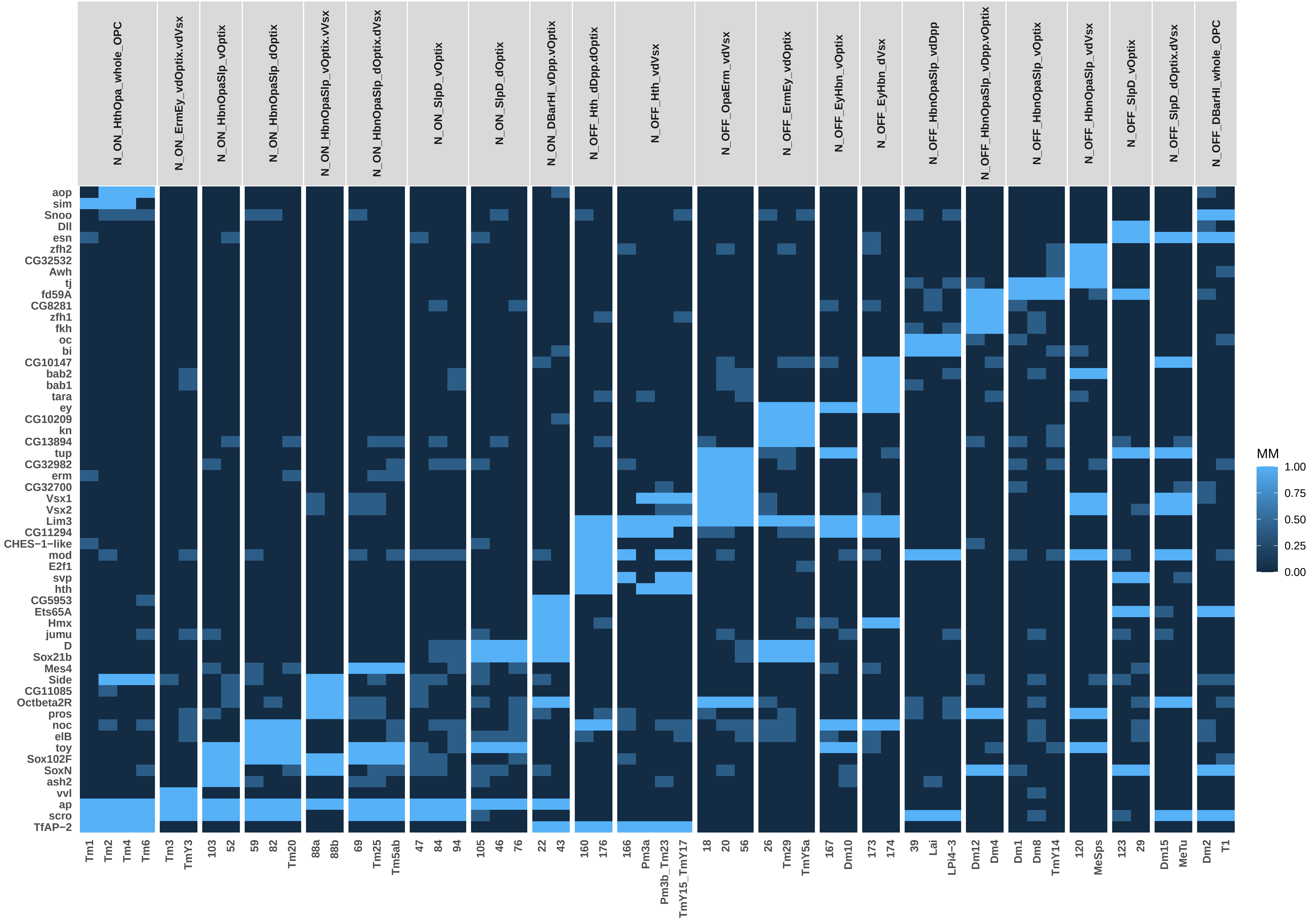

### P40_SO_CSSs_0.75ClusterFreq_0.5BinThreshold_0.5SpecificityThreshold_Simplified.pdf

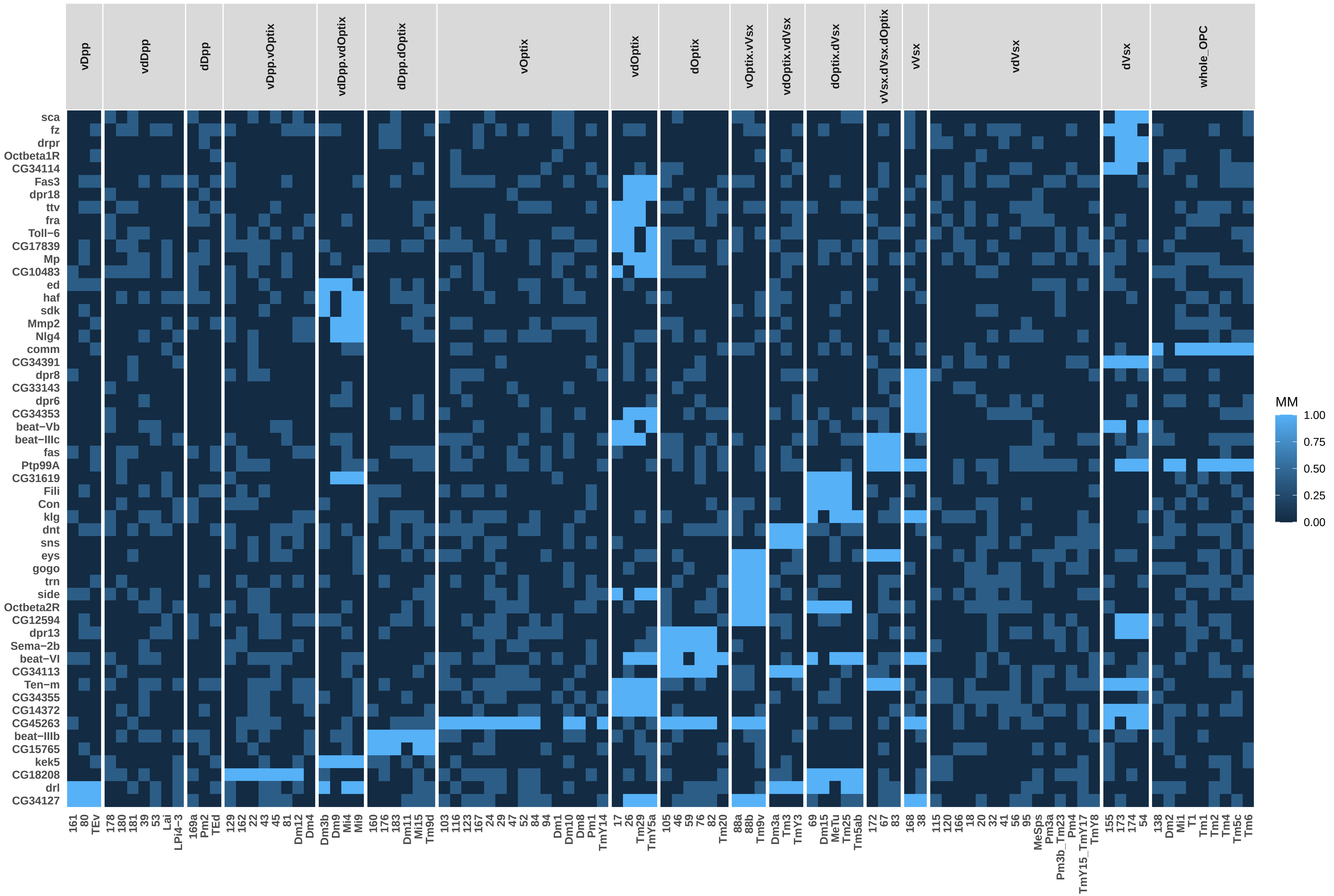

### P40_SO_non_TFs_CSSs_0.75ClusterFreq_0.5BinThreshold_0.5SpecificityThreshold_Simplified.pdf

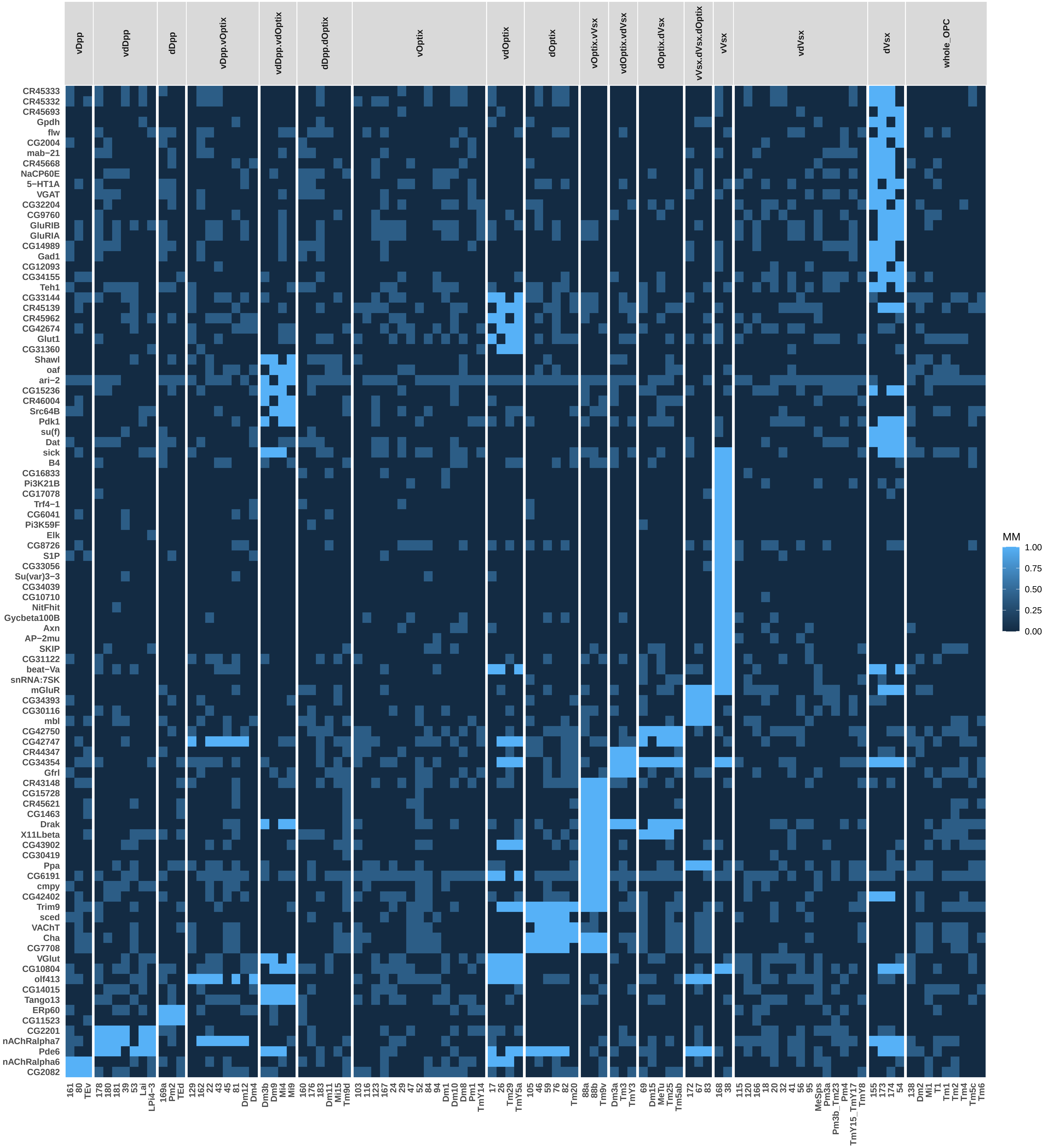

### P40_SO_TFs_0.75ClusterFreq_0.5BinThreshold_0.5SpecificityThreshold_Simplified.pdf

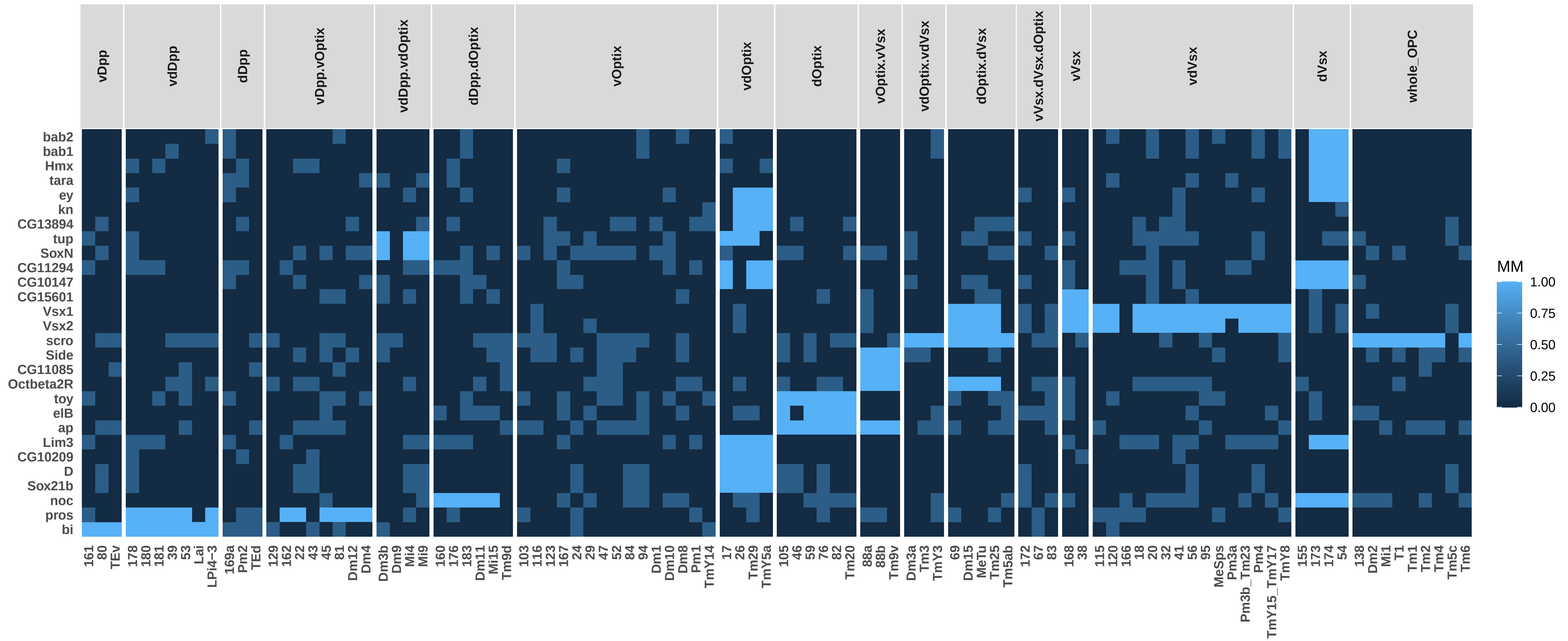

### P40_TO_CSSs_0.75ClusterFreq_0.5BinThreshold_0.5SpecificityThreshold_Simplified.pdf

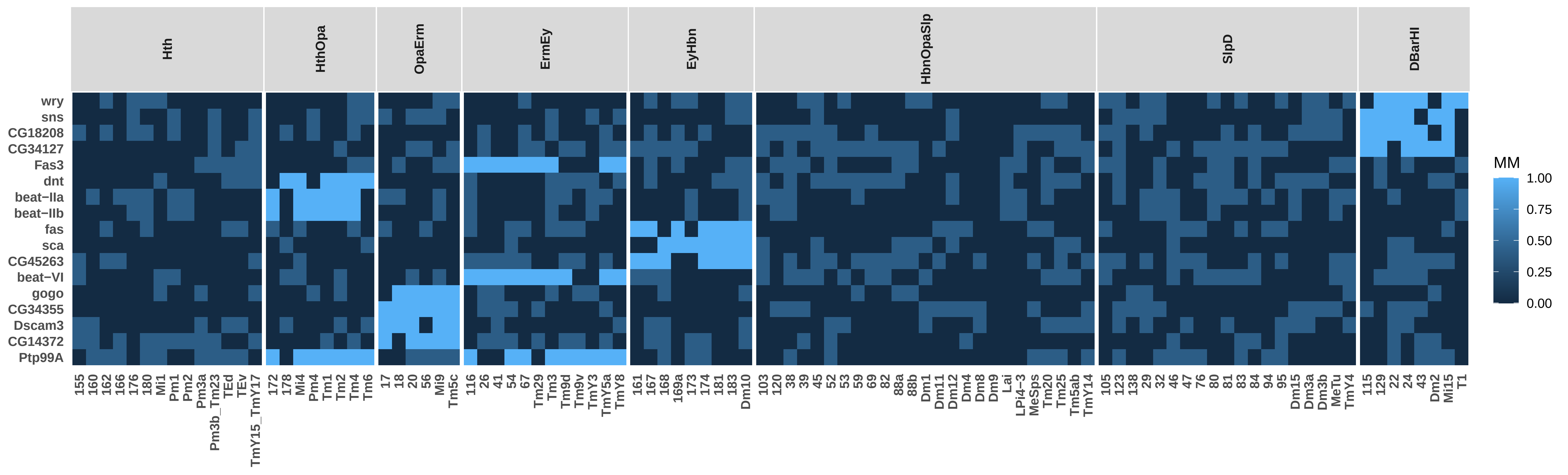

### P40_TO_NO_CSSs_0.75ClusterFreq_0.5BinThreshold_0.5SpecificityThreshold_Simplified.pdf

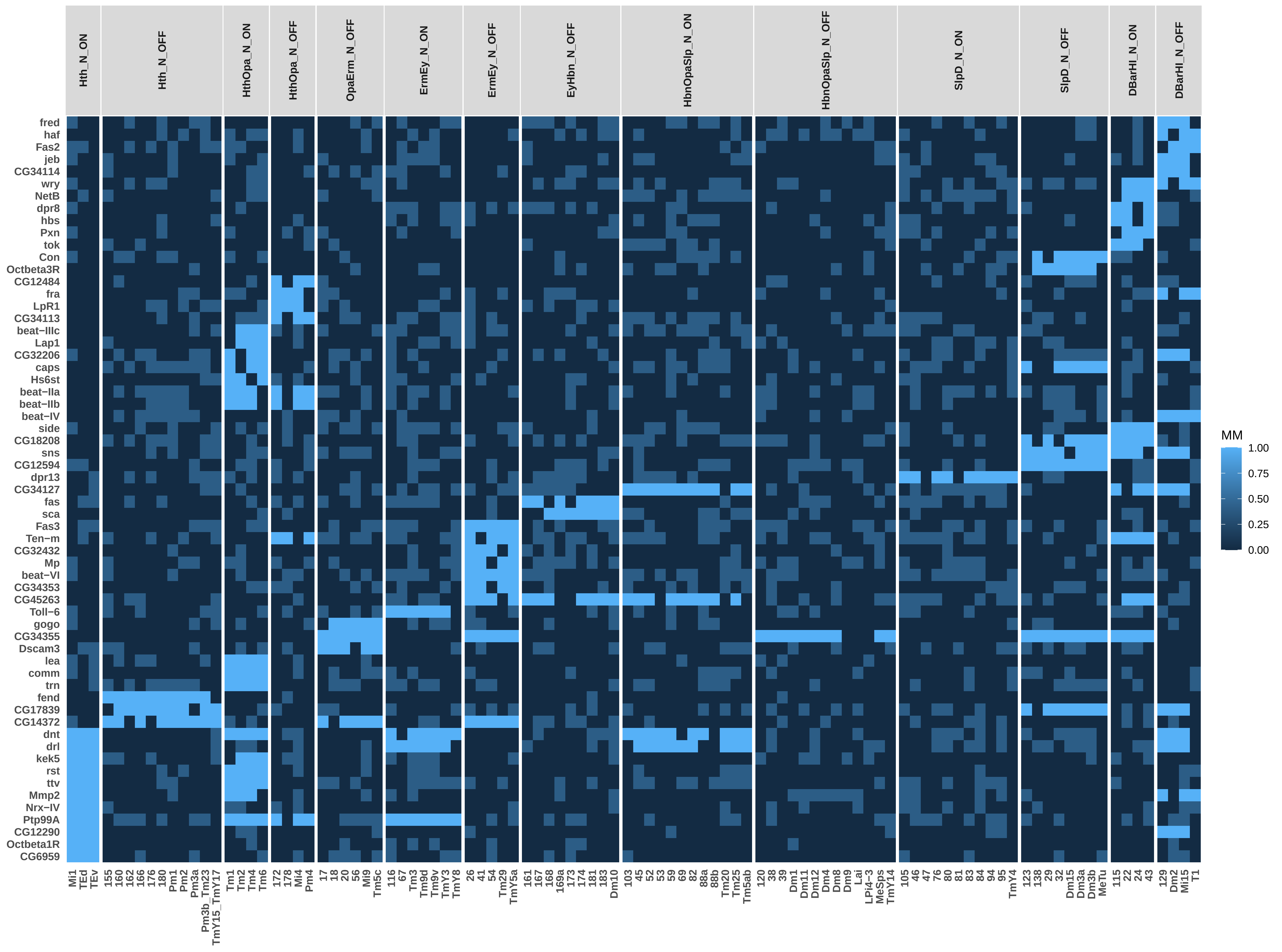

### P40_TO_NO_non_TFs_CSSs_0.75ClusterFreq_0.5BinThreshold_0.5SpecificityThreshold_Simplified.pdf

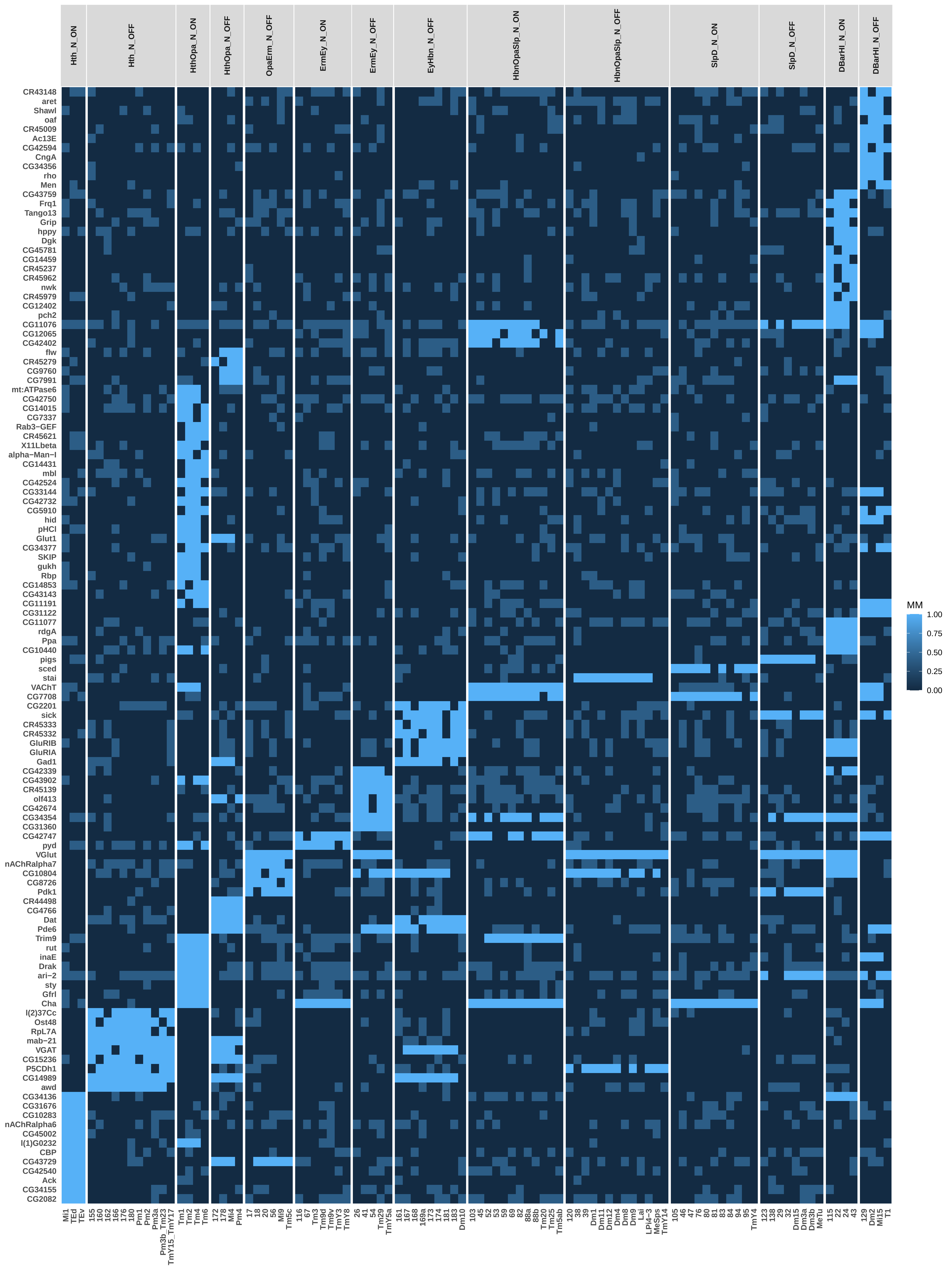

### P40_TO_NO_TFs_0.75ClusterFreq_0.5BinThreshold_0.5SpecificityThreshold_Simplified.pdf

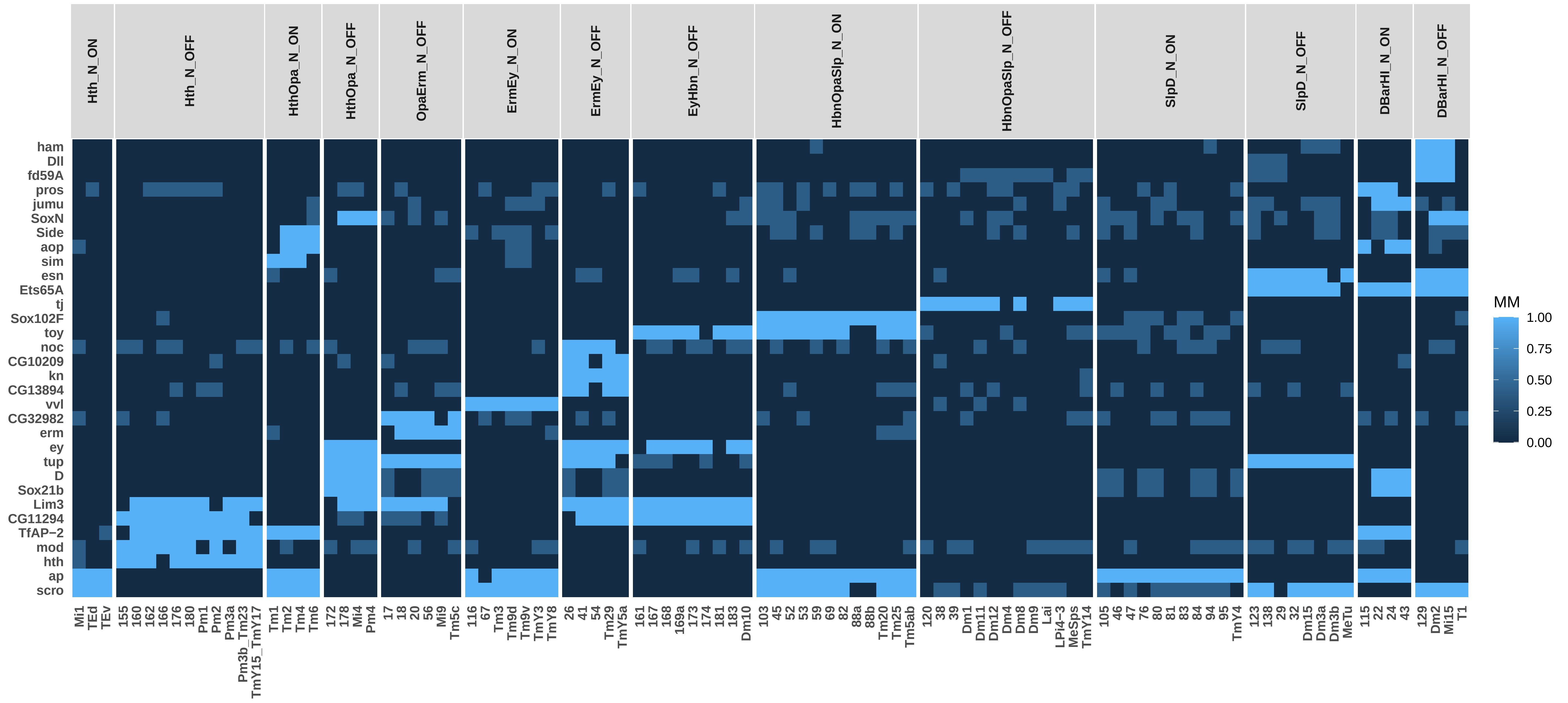

### P40_TO_non_TFs_CSSs_0.75ClusterFreq_0.5BinThreshold_0.5SpecificityThreshold_Simplified.pdf

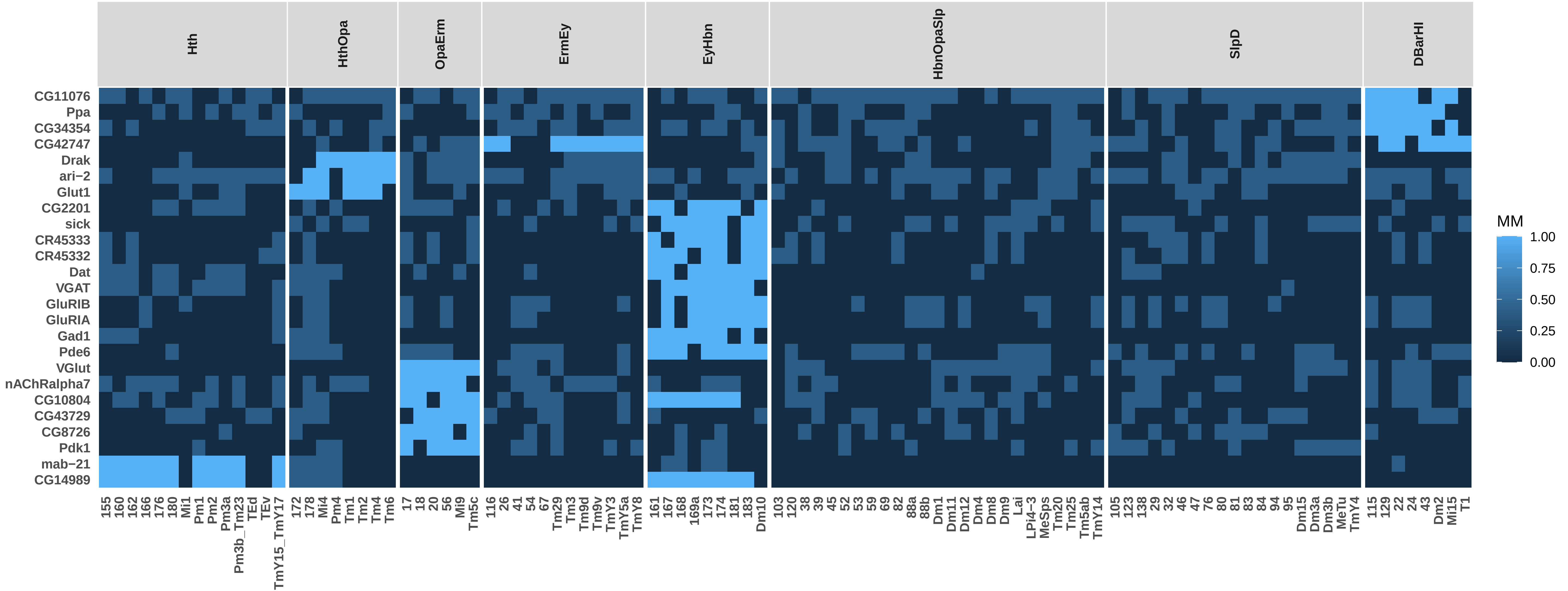

### P40_TO_SO_CSSs_0.75ClusterFreq_0.5BinThreshold_0.5SpecificityThreshold_Simplified.pdf

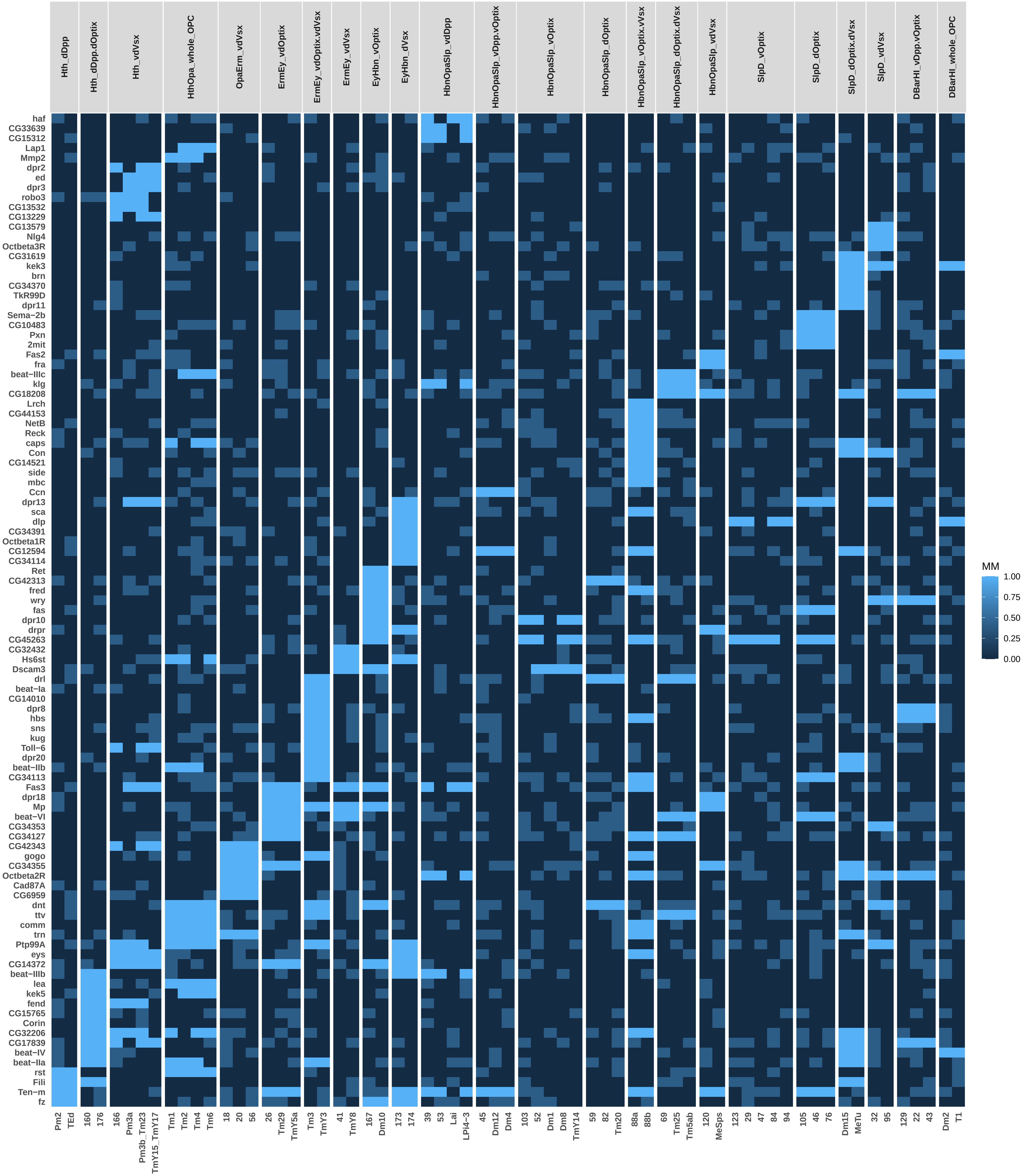

### P40_TO_SO_non_TFs_CSSs_0.75ClusterFreq_0.5BinThreshold_0.5SpecificityThreshold_Simplified.pdf

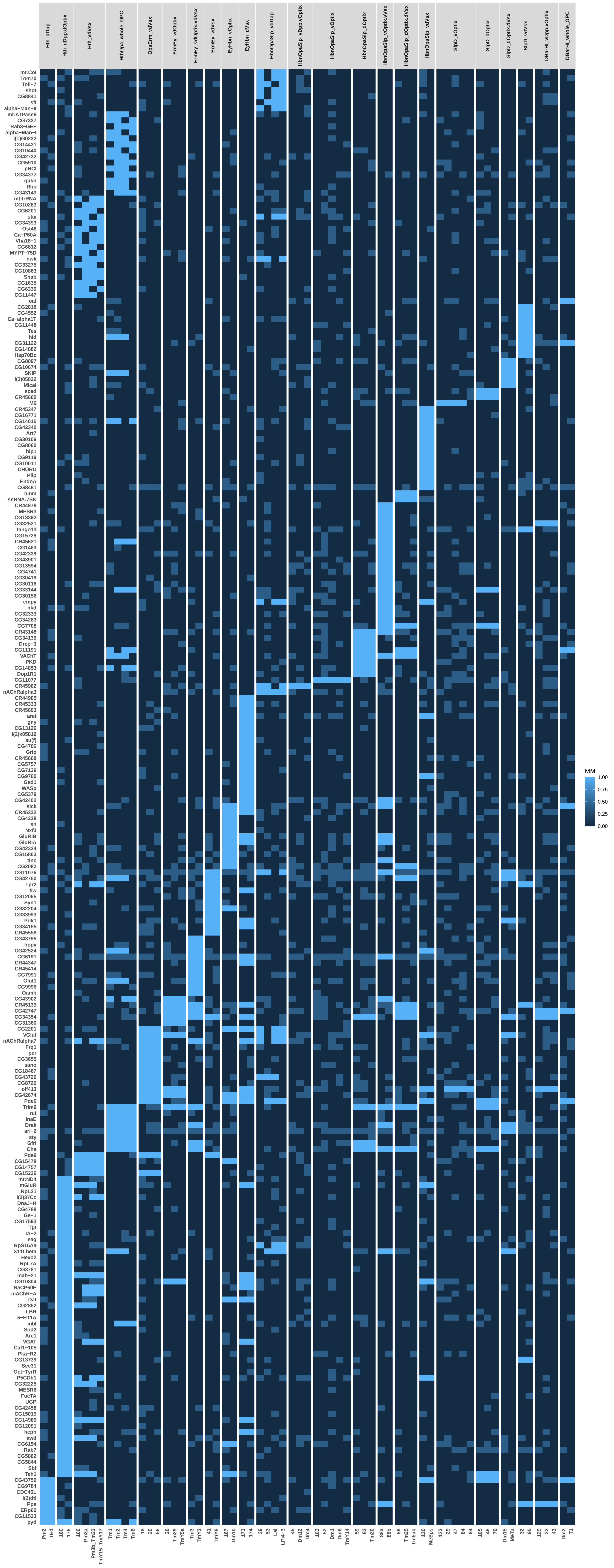

### P40_TO_SO_TFs_0.75ClusterFreq_0.5BinThreshold_0.5SpecificityThreshold_Simplified.pdf

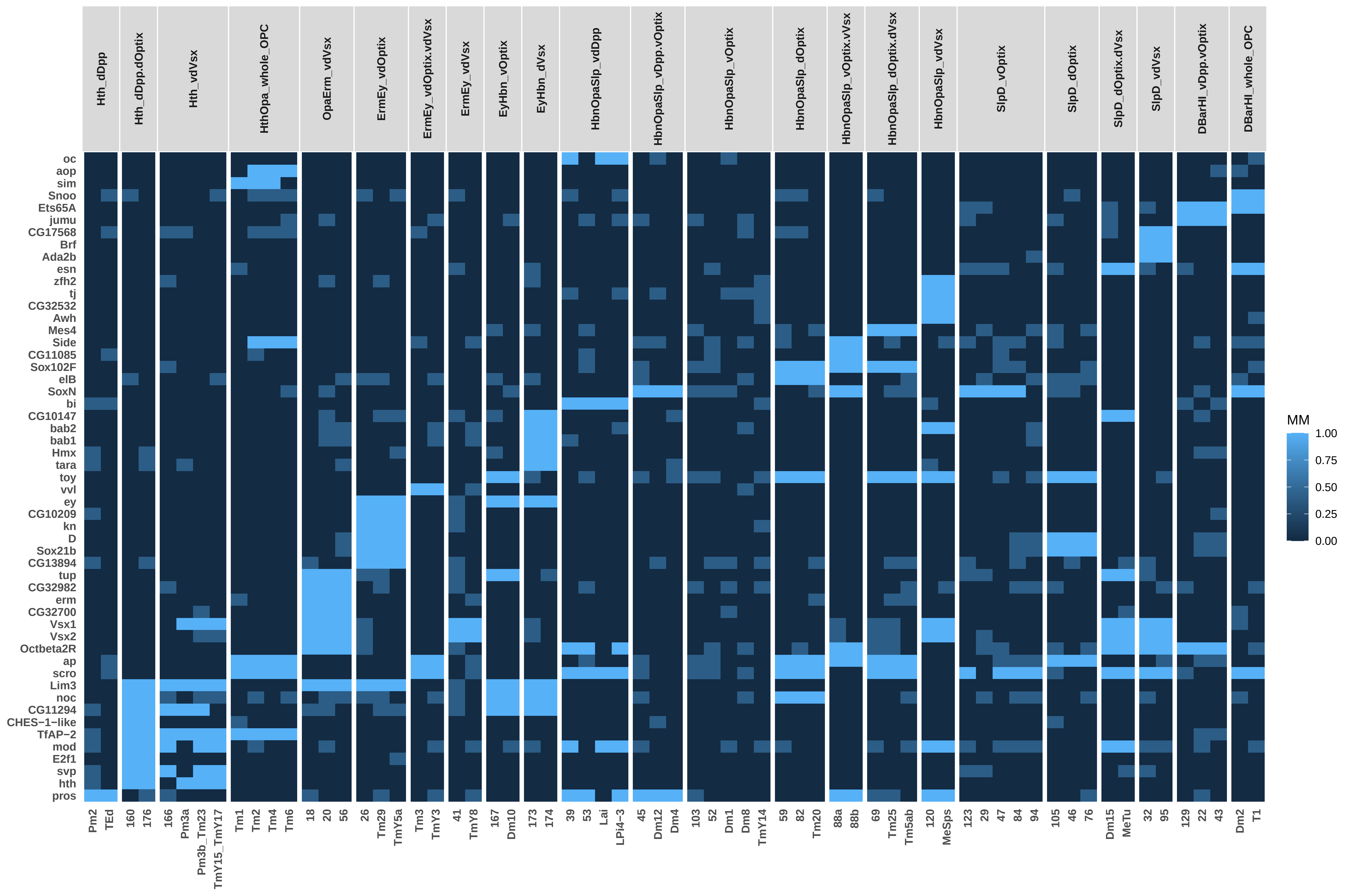

### P40_TO_TFs_0.75ClusterFreq_0.5BinThreshold_0.5SpecificityThreshold_Simplified.pdf

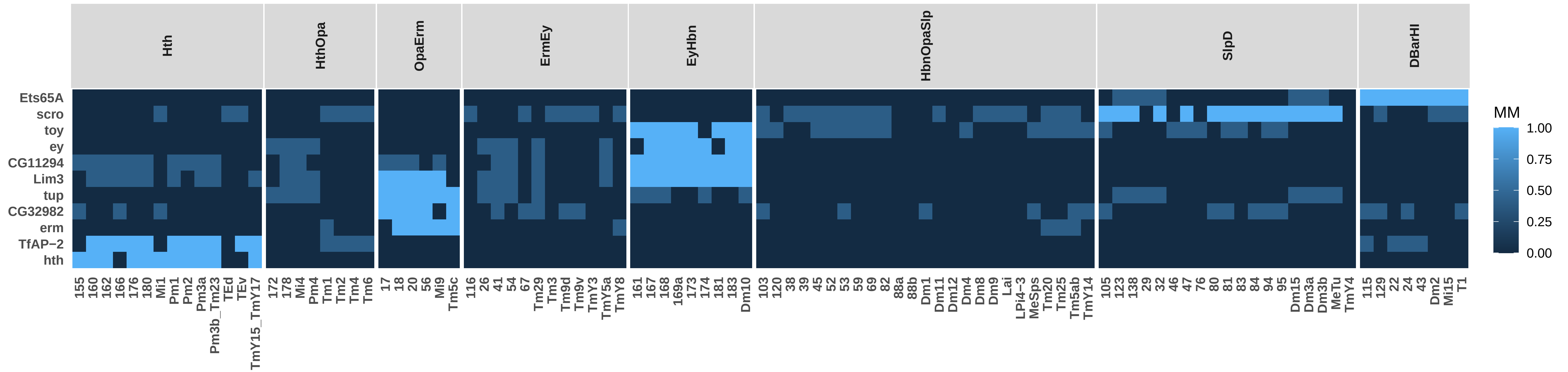

### P50_NO_non_TFs_CSSs_0.75ClusterFreq_0.5BinThreshold_0.5SpecificityThreshold_Simplified.pdf

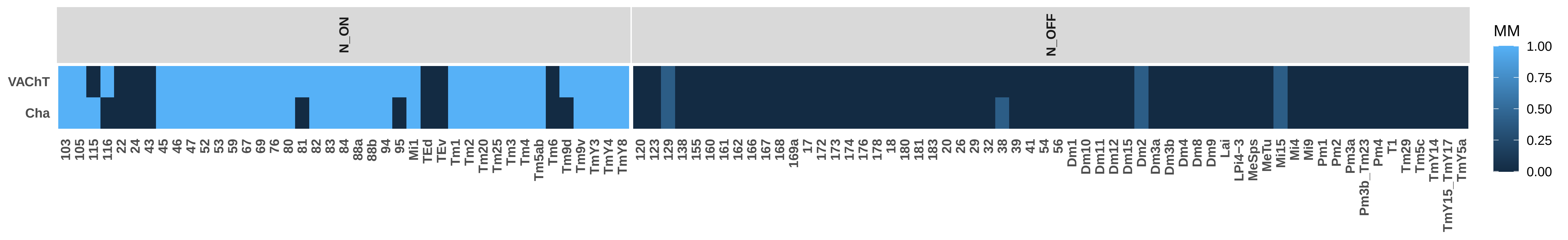

### P50_NO_SO_CSSs_0.75ClusterFreq_0.5BinThreshold_0.5SpecificityThreshold_Simplified.pdf

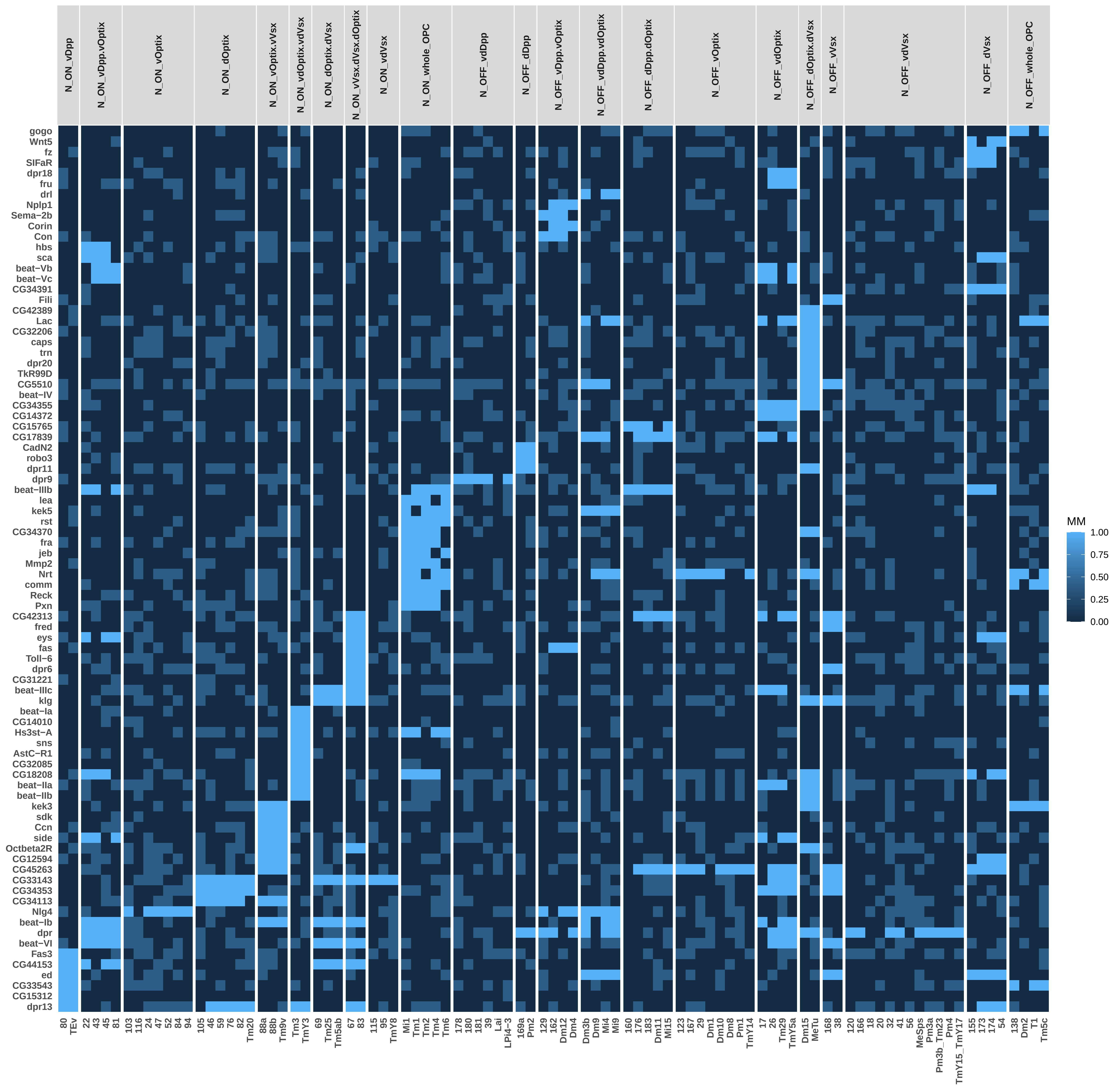

### P50_NO_SO_non_TFs_CSSs_0.75ClusterFreq_0.5BinThreshold_0.5SpecificityThreshold_Simplified.pdf

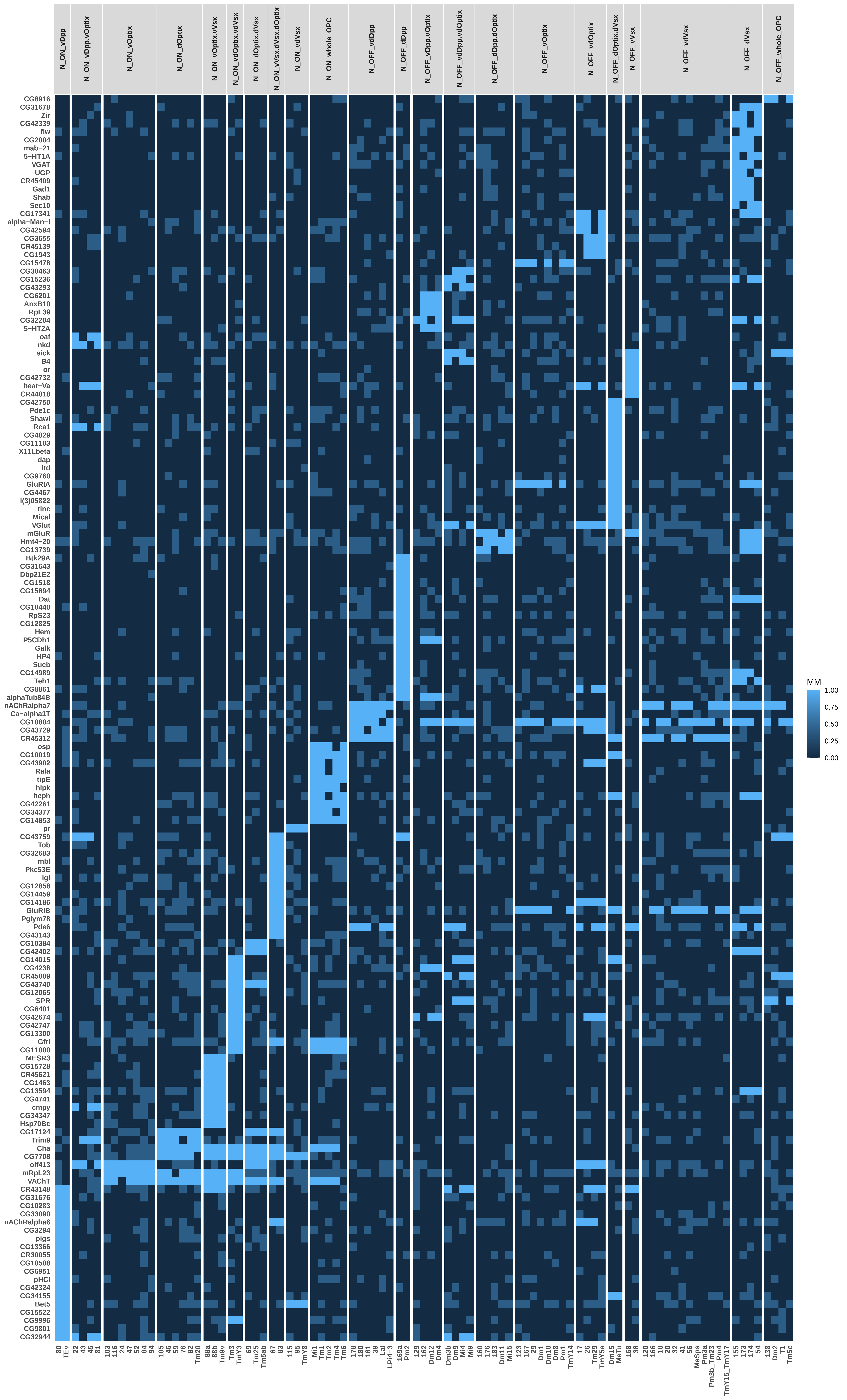

### P50_NO_SO_TFs_0.75ClusterFreq_0.5BinThreshold_0.5SpecificityThreshold_Simplified.pdf

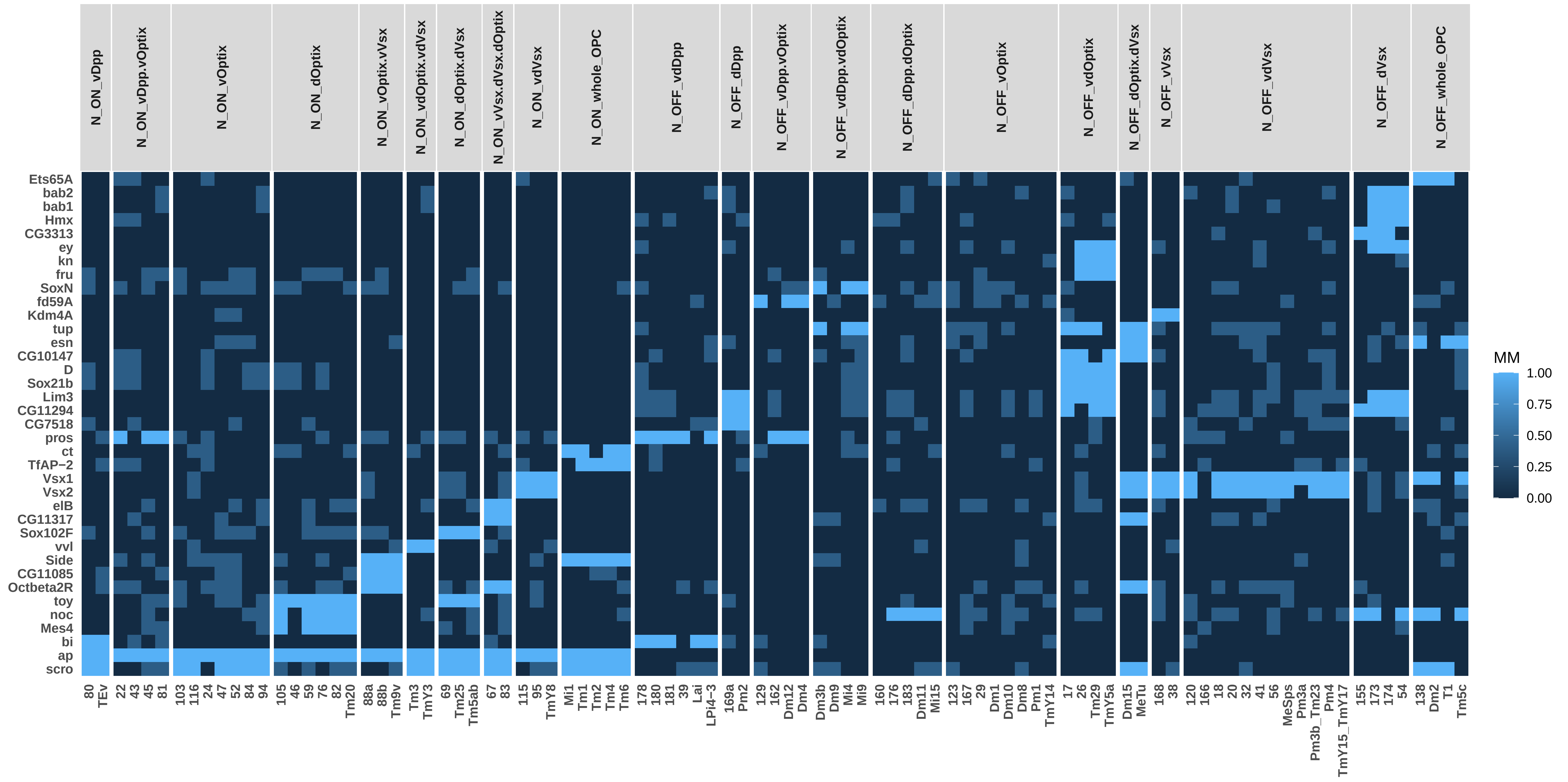

### P50_NO_TFs_0.75ClusterFreq_0.5BinThreshold_0.5SpecificityThreshold_Simplified.pdf

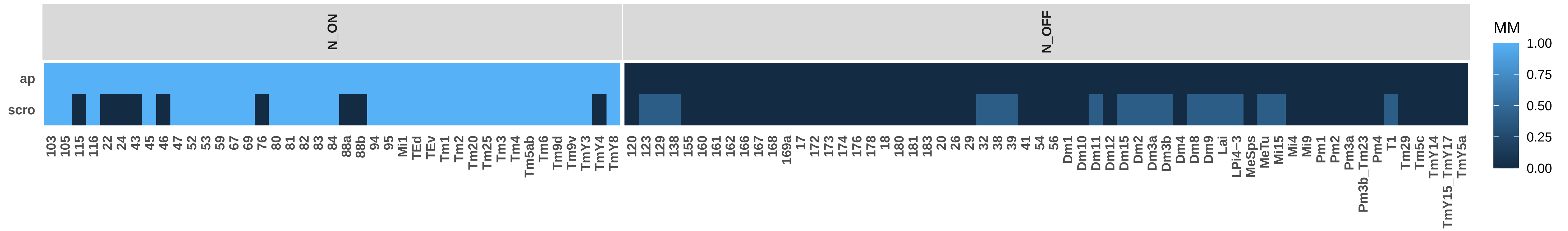

### P50_NO_TO_SO_CSSs_0.75ClusterFreq_0.5BinThreshold_0.5SpecificityThreshold_Simplified.pdf

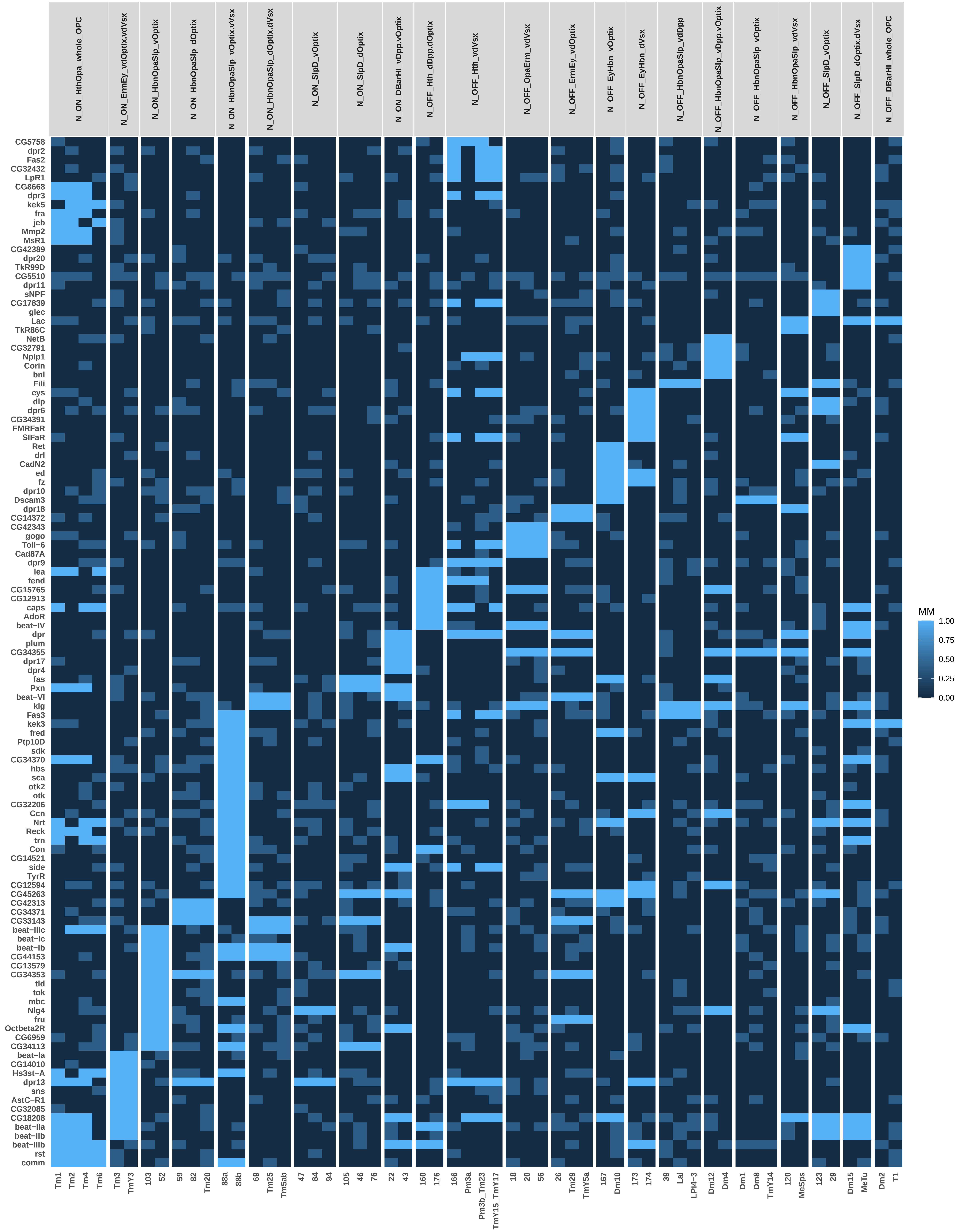

### P50_NO_TO_SO_non_TFs_CSSs_0.75ClusterFreq_0.5BinThreshold_0.5SpecificityThreshold_Simplified.pdf

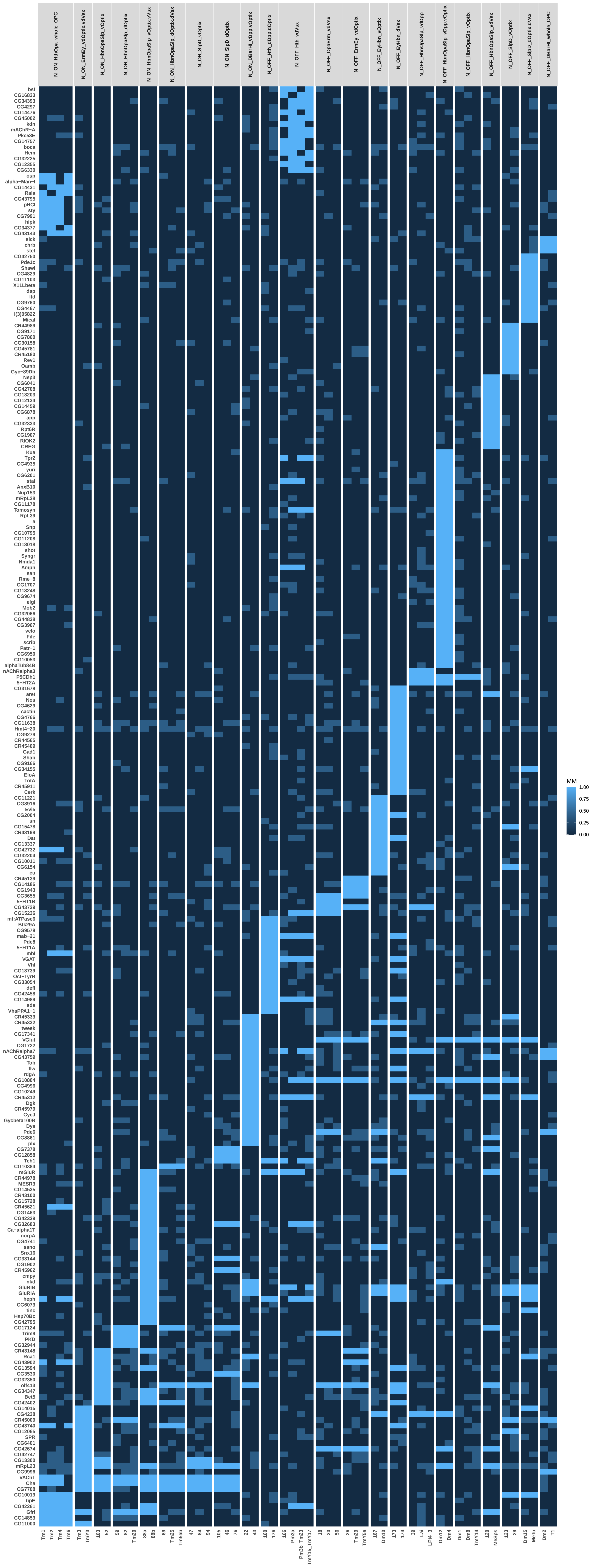

### P50_NO_TO_SO_TFs_0.75ClusterFreq_0.5BinThreshold_0.5SpecificityThreshold_Simplified.pdf

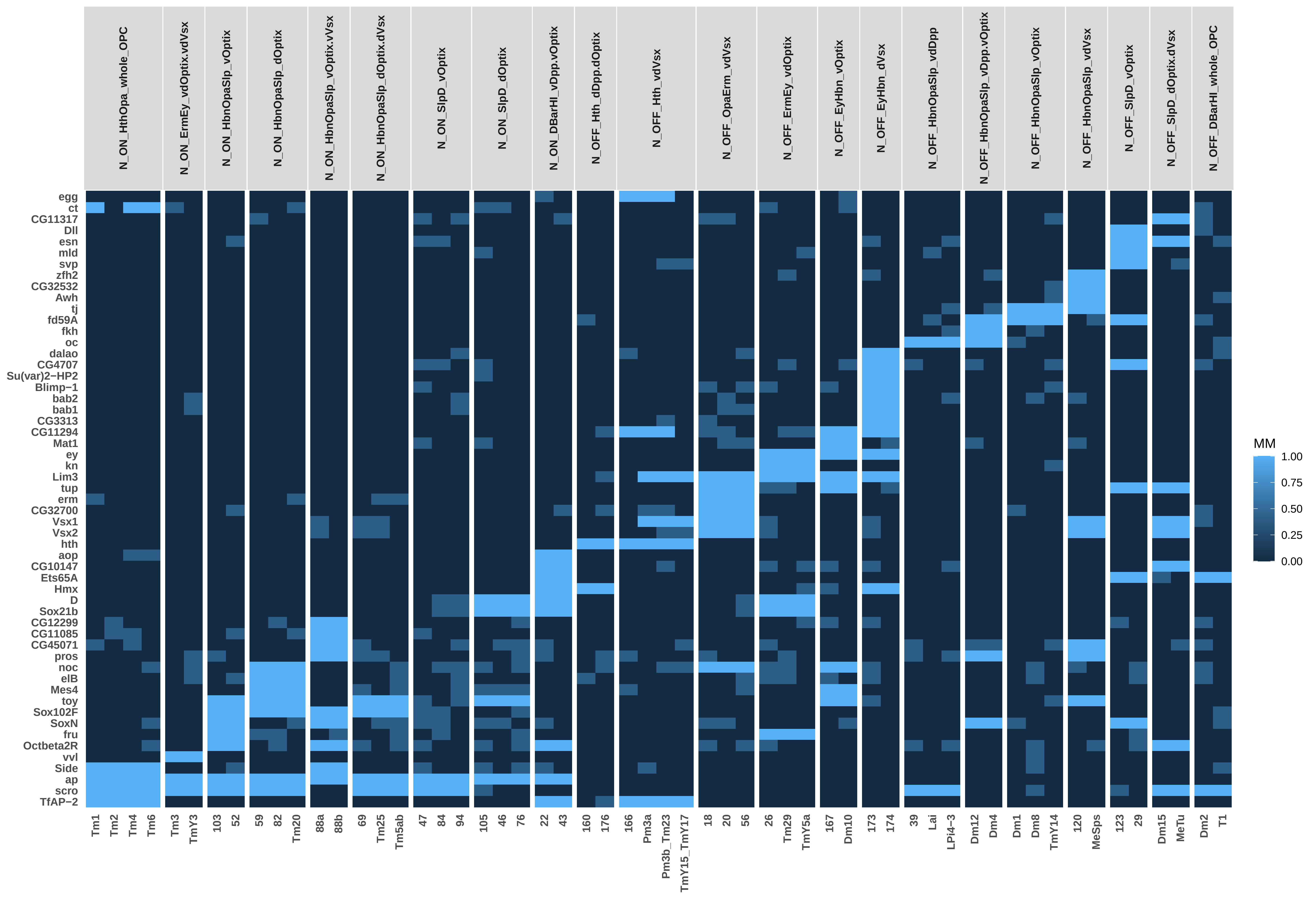

### P50_SO_CSSs_0.75ClusterFreq_0.5BinThreshold_0.5SpecificityThreshold_Simplified.pdf

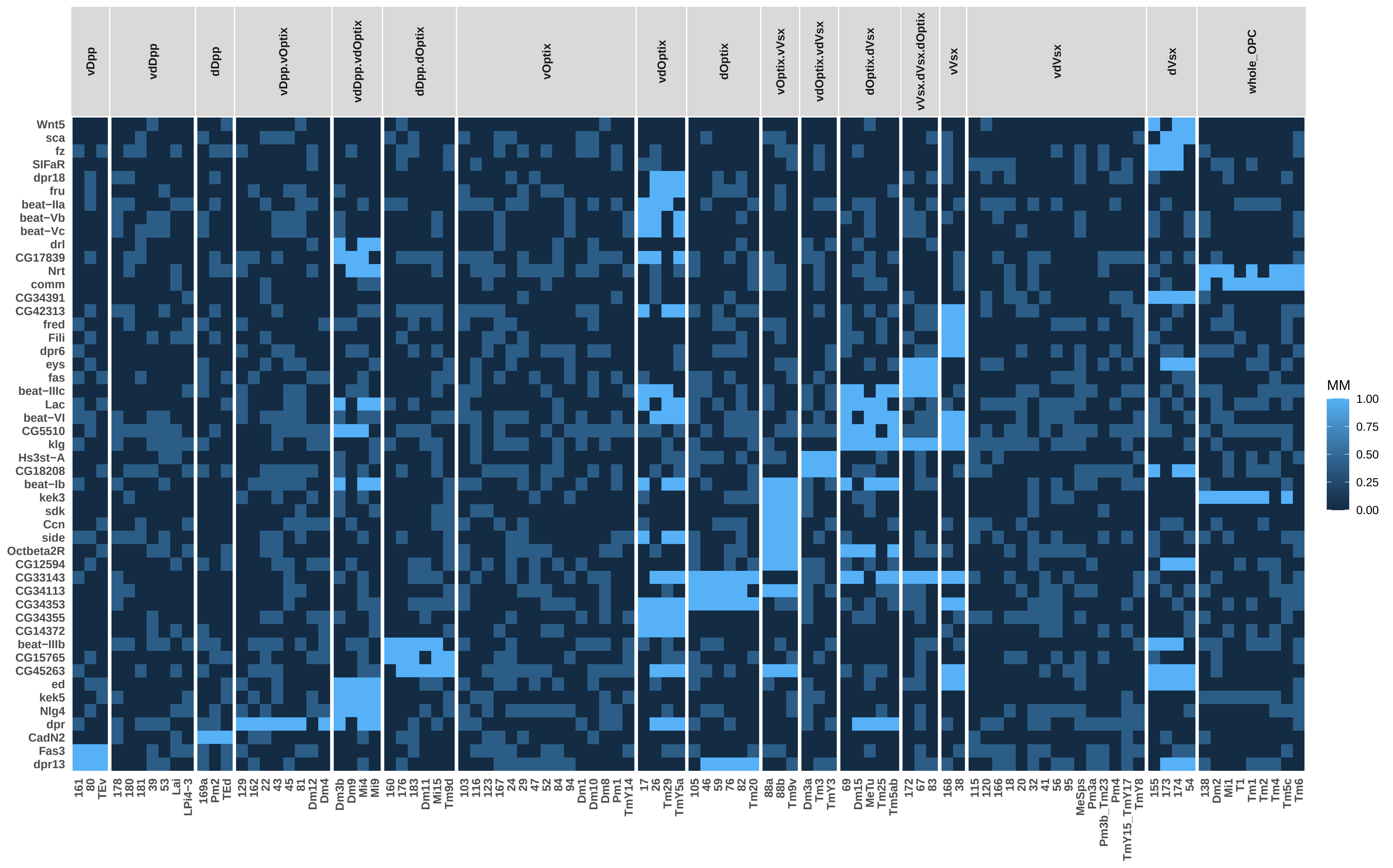

### P50_SO_non_TFs_CSSs_0.75ClusterFreq_0.5BinThreshold_0.5SpecificityThreshold_Simplified.pdf

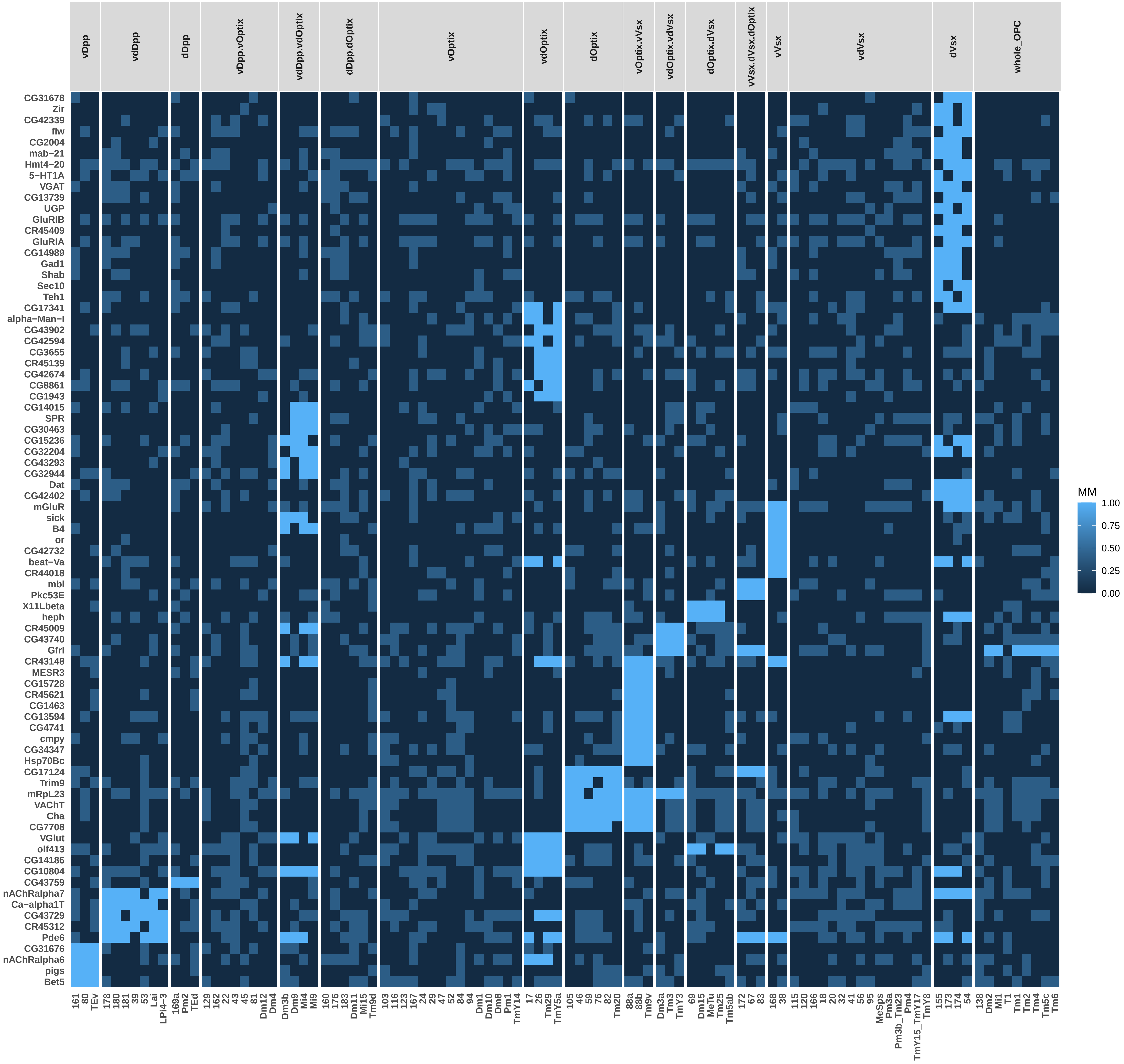

### P50_SO_TFs_0.75ClusterFreq_0.5BinThreshold_0.5SpecificityThreshold_Simplified.pdf

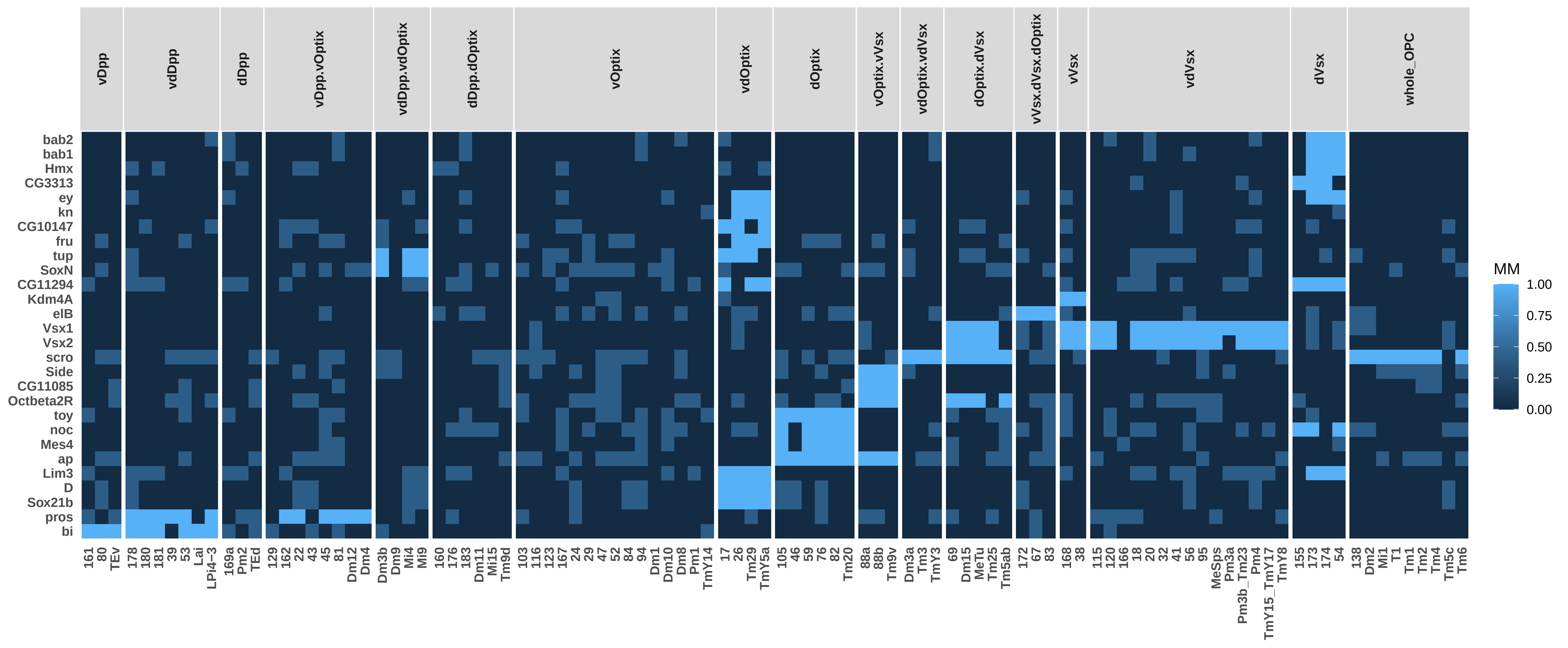

### P50_TO_CSSs_0.75ClusterFreq_0.5BinThreshold_0.5SpecificityThreshold_Simplified.pdf

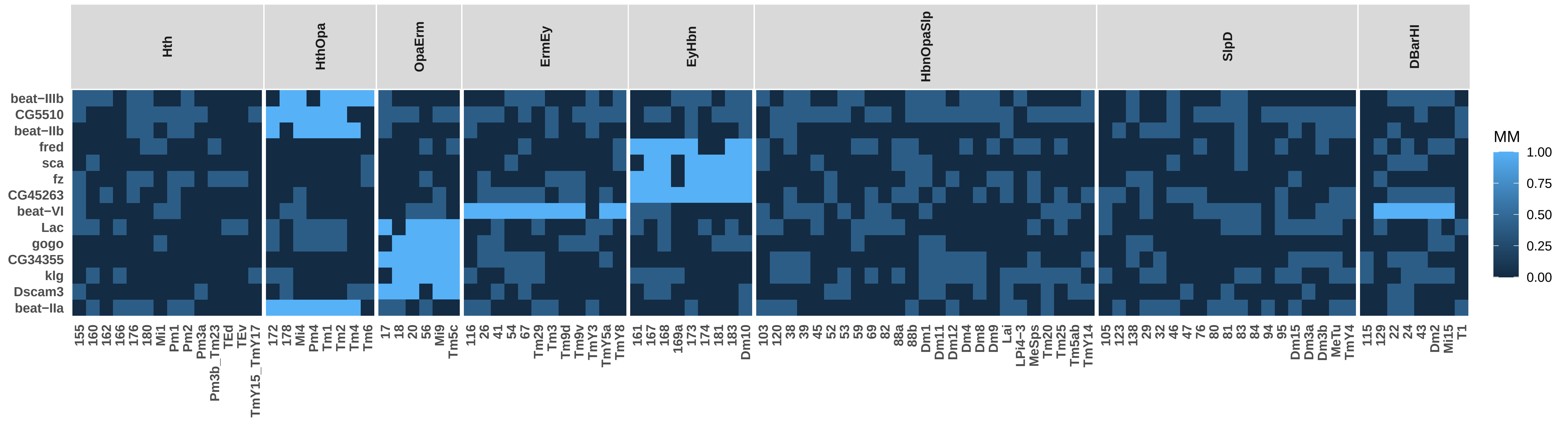

### P50_TO_NO_CSSs_0.75ClusterFreq_0.5BinThreshold_0.5SpecificityThreshold_Simplified.pdf

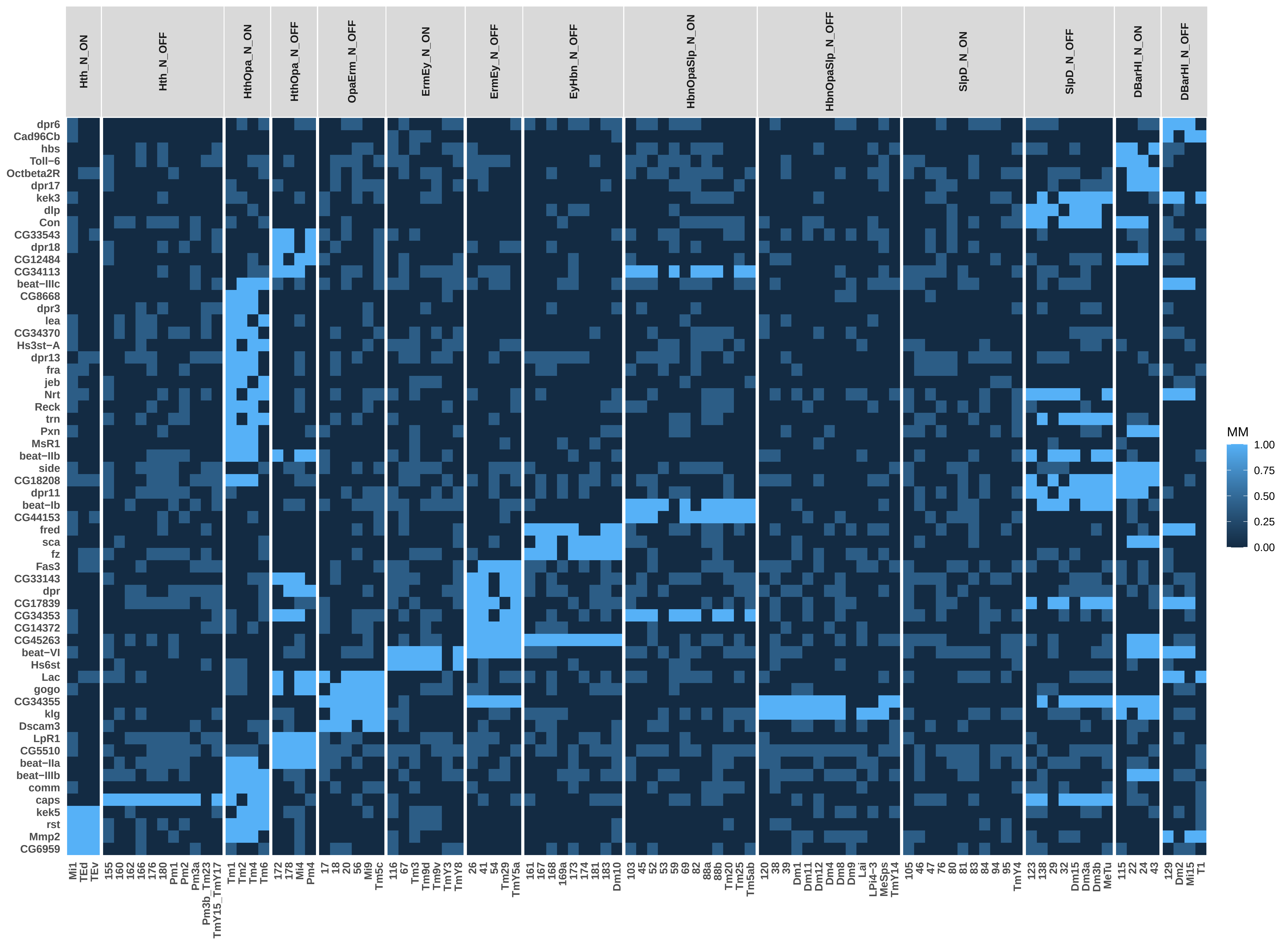

### P50_TO_NO_non_TFs_CSSs_0.75ClusterFreq_0.5BinThreshold_0.5SpecificityThreshold_Simplified.pdf

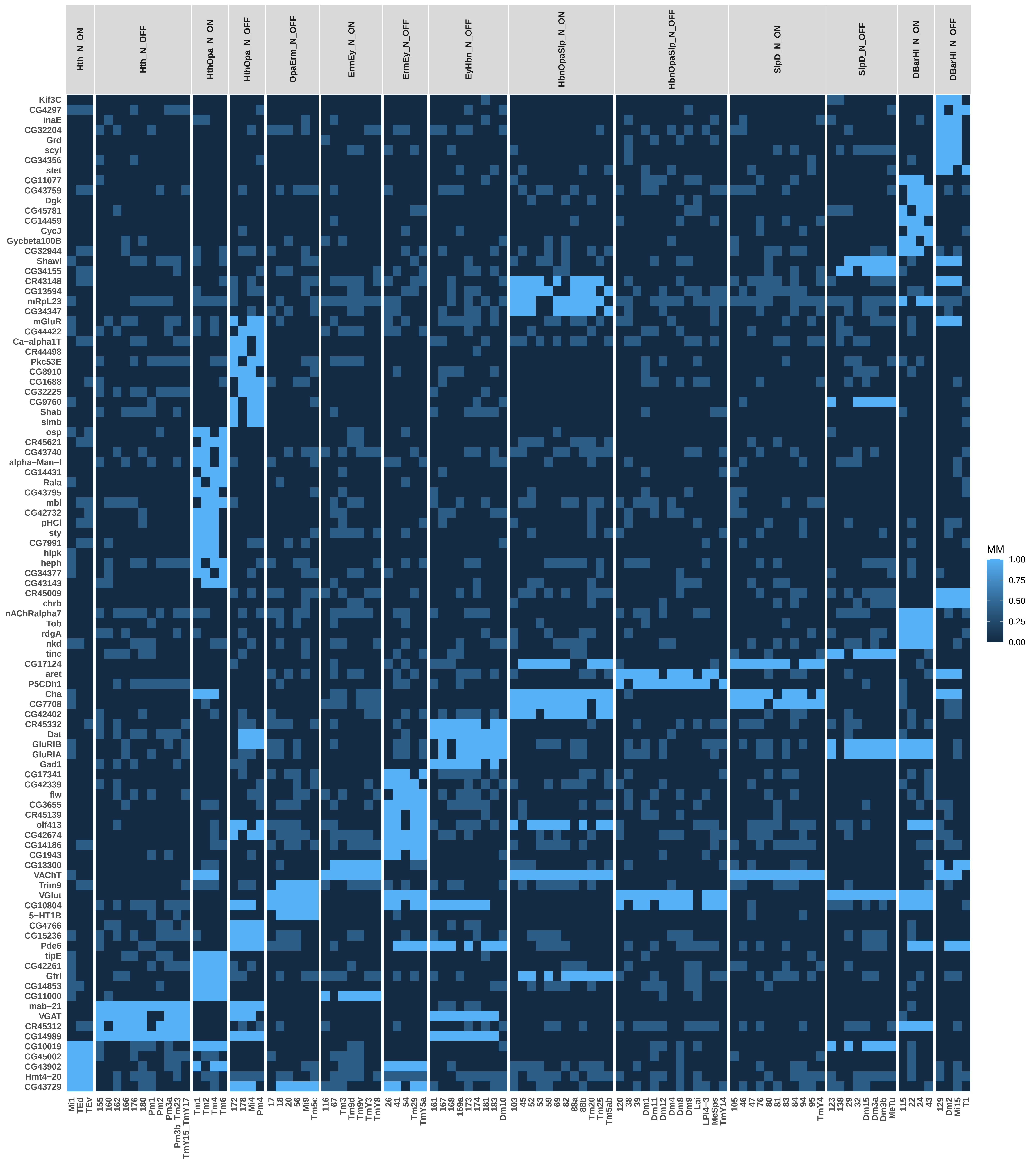

### P50_TO_NO_TFs_0.75ClusterFreq_0.5BinThreshold_0.5SpecificityThreshold_Simplified.pdf

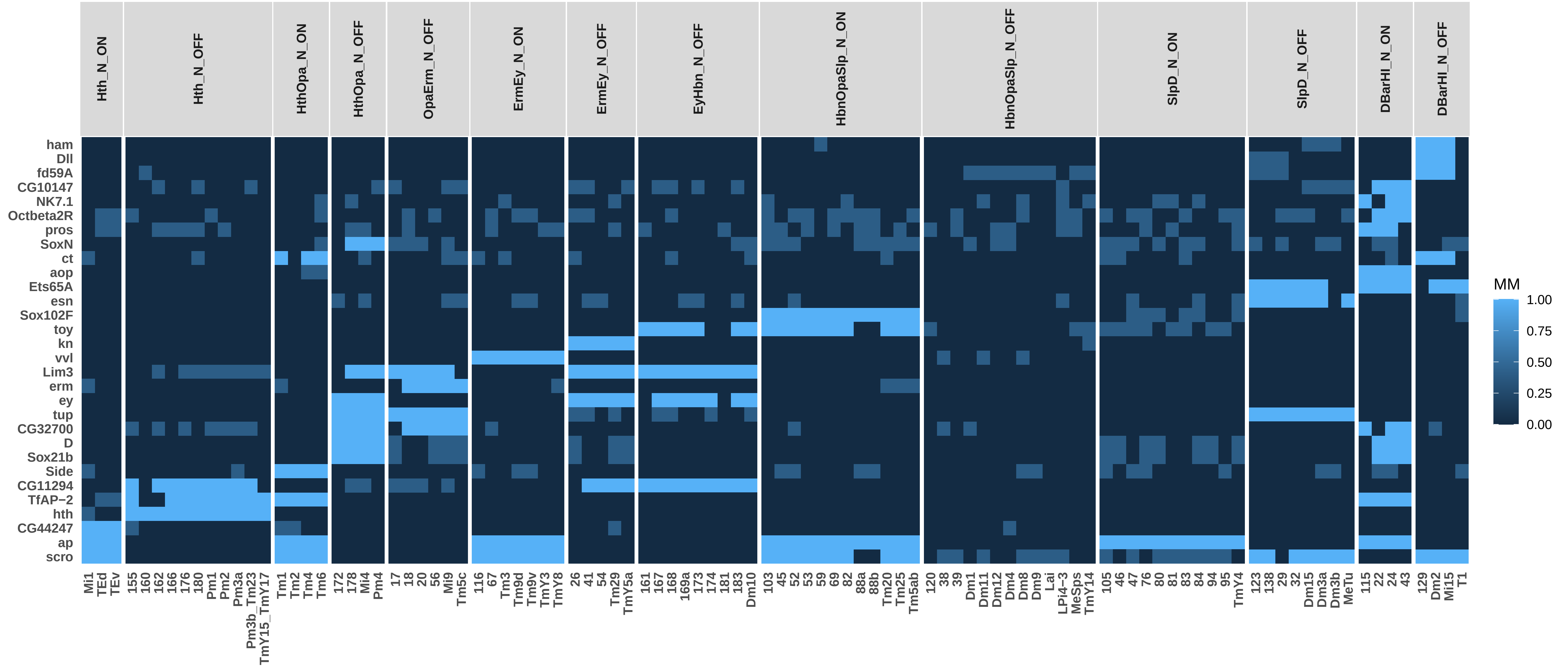

### P50_TO_non_TFs_CSSs_0.75ClusterFreq_0.5BinThreshold_0.5SpecificityThreshold_Simplified.pdf

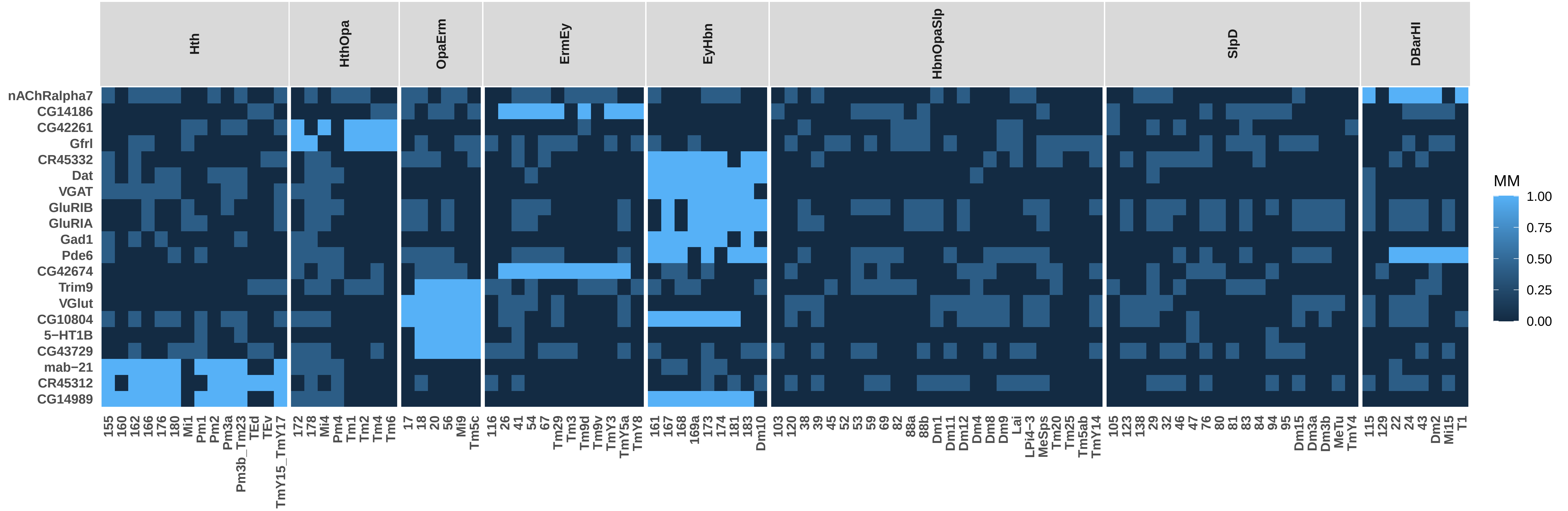

### P50_TO_SO_CSSs_0.75ClusterFreq_0.5BinThreshold_0.5SpecificityThreshold_Simplified.pdf

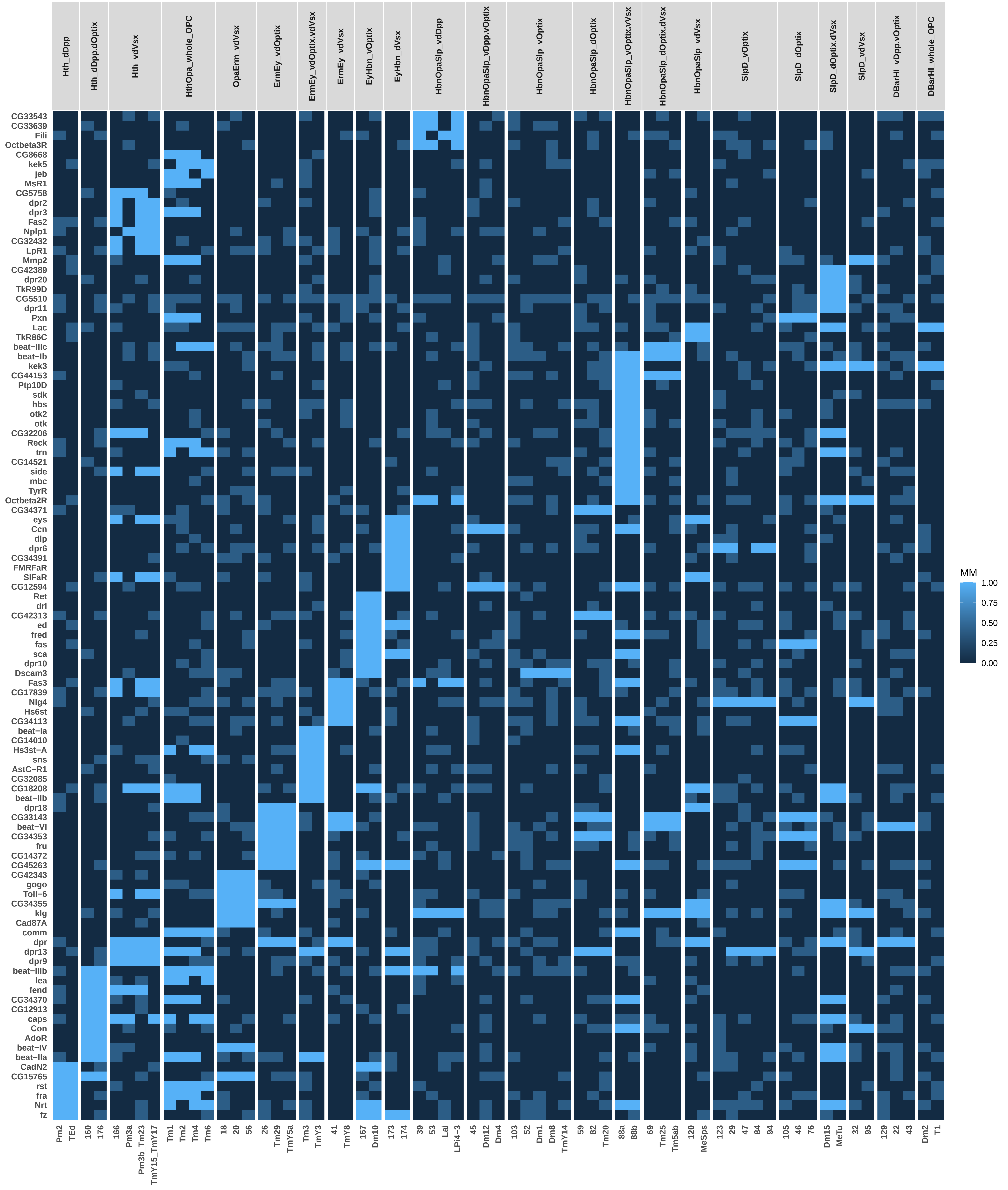

### P50_TO_SO_non_TFs_CSSs_0.75ClusterFreq_0.5BinThreshold_0.5SpecificityThreshold_Simplified.pdf

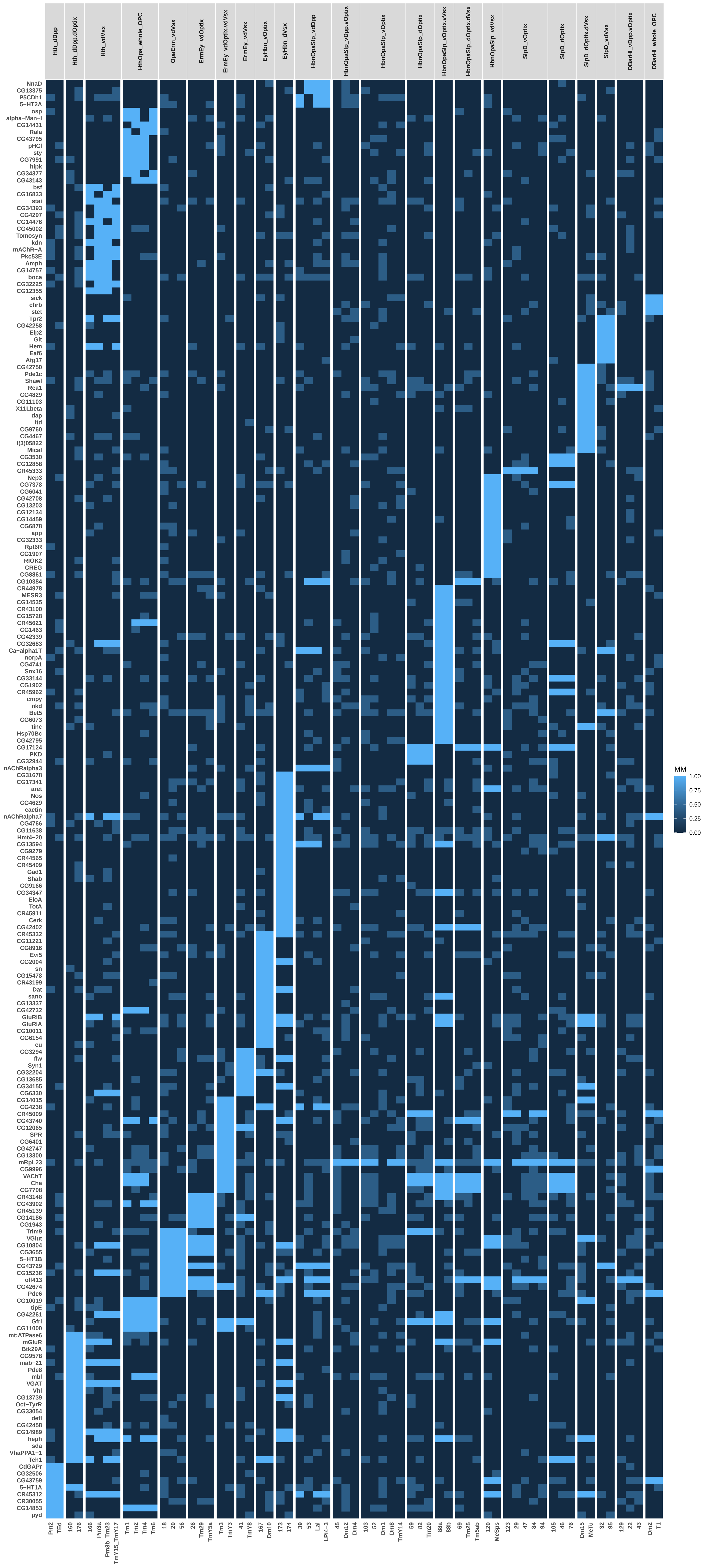

### P50_TO_SO_TFs_0.75ClusterFreq_0.5BinThreshold_0.5SpecificityThreshold_Simplified.pdf

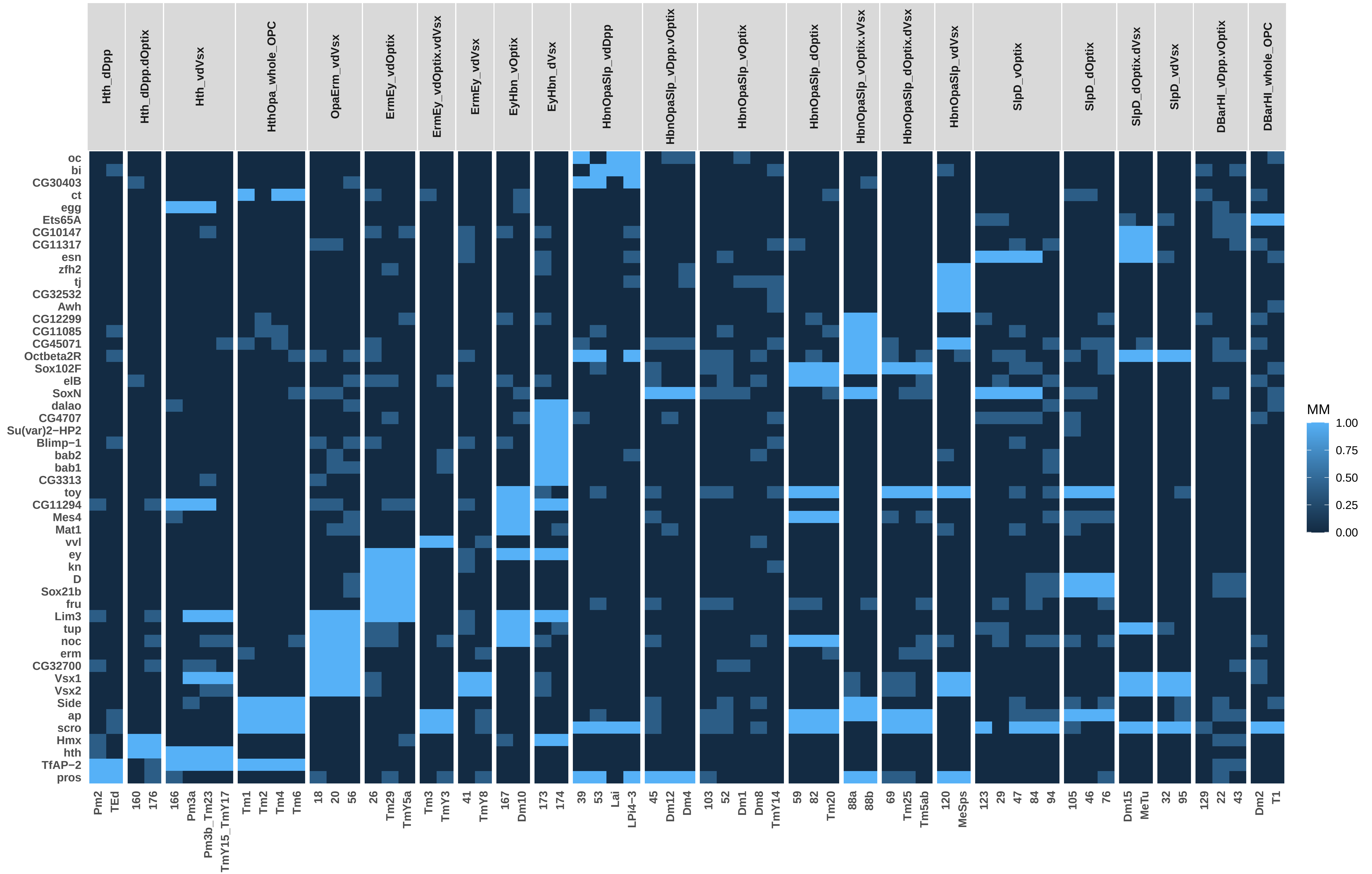

### P50_TO_TFs_0.75ClusterFreq_0.5BinThreshold_0.5SpecificityThreshold_Simplified.pdf

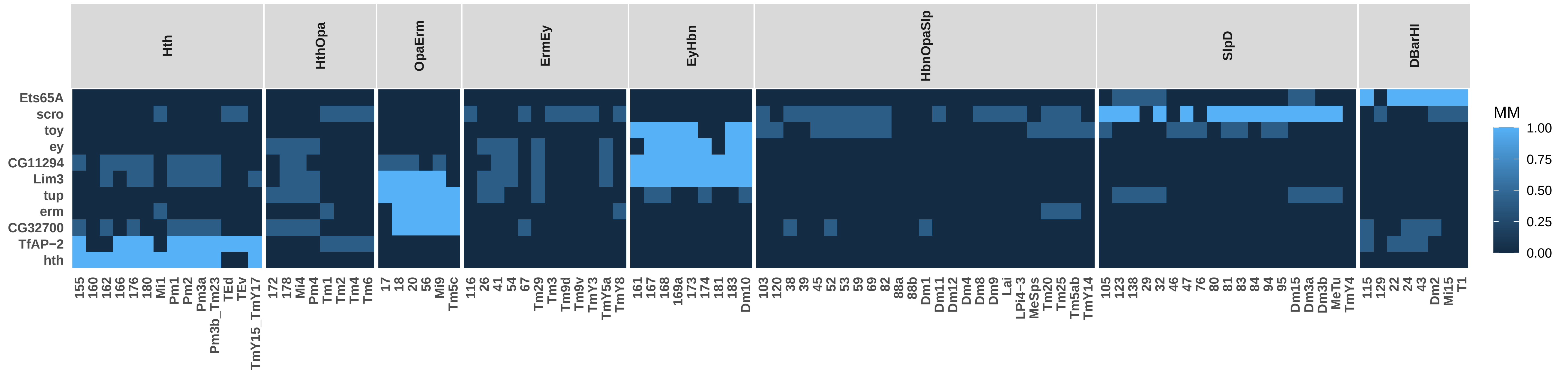

### P70_NO_CSSs_0.75ClusterFreq_0.5BinThreshold_0.5SpecificityThreshold_Simplified.pdf

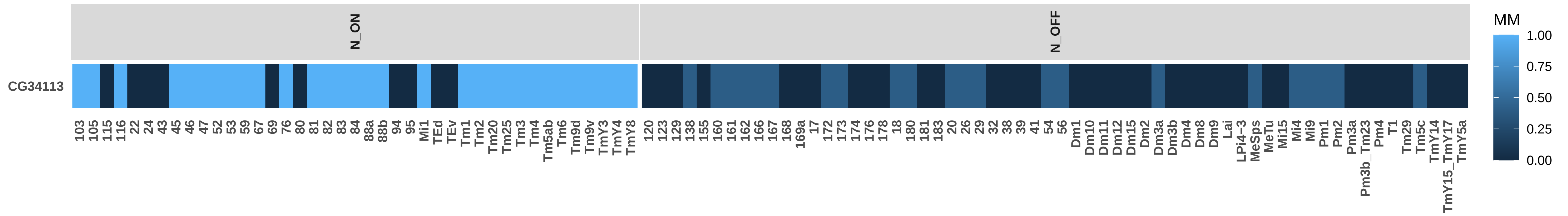

### P70_NO_non_TFs_CSSs_0.75ClusterFreq_0.5BinThreshold_0.5SpecificityThreshold_Simplified.pdf

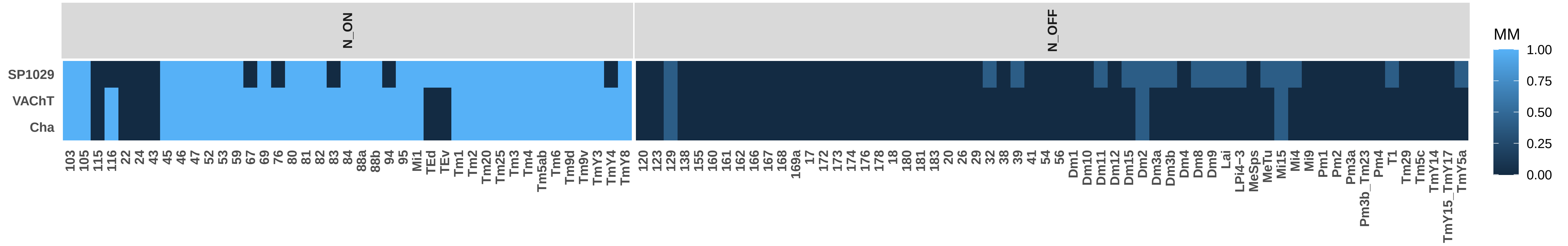

### P70_NO_SO_CSSs_0.75ClusterFreq_0.5BinThreshold_0.5SpecificityThreshold_Simplified.pdf

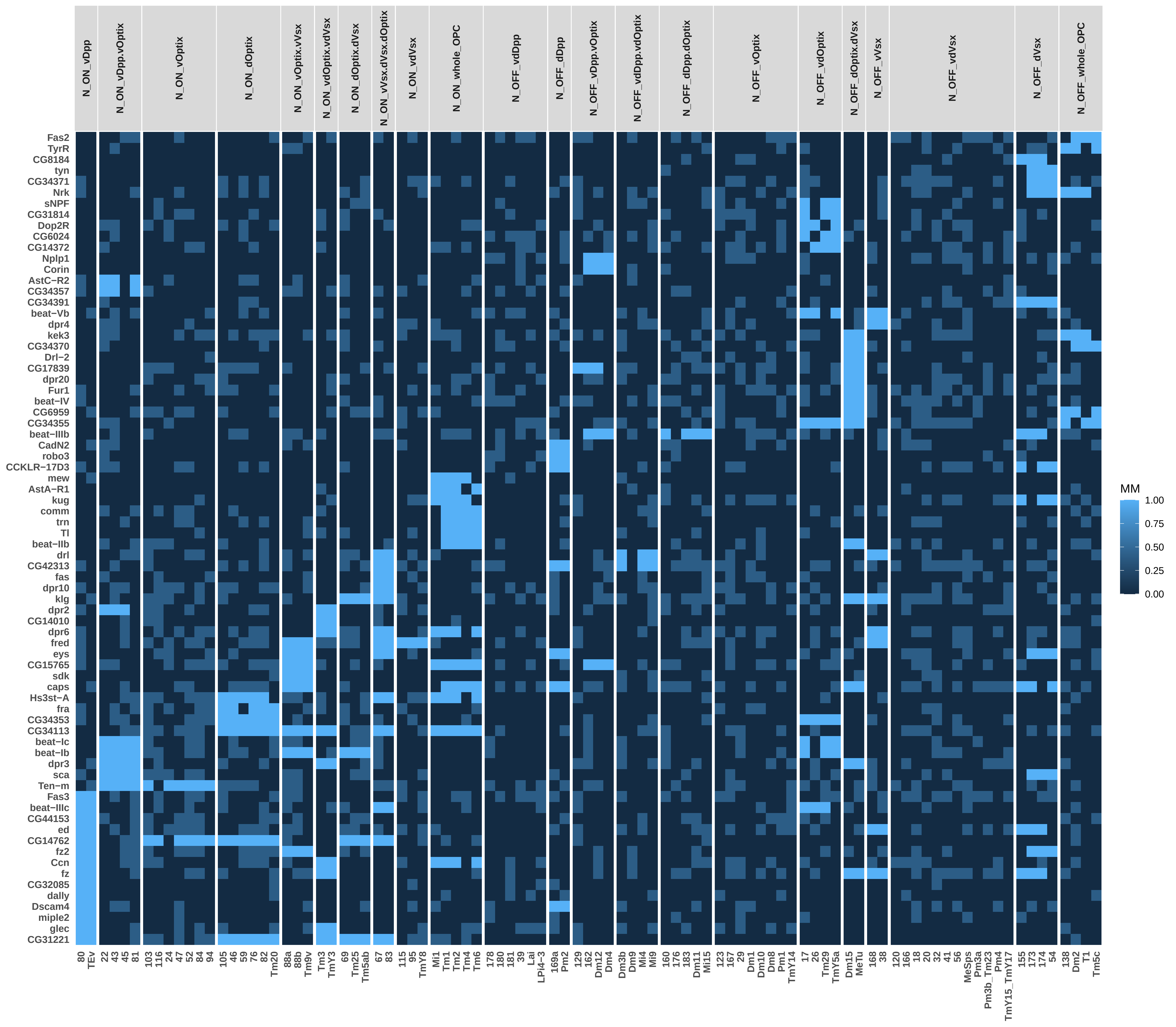

### P70_NO_SO_non_TFs_CSSs_0.75ClusterFreq_0.5BinThreshold_0.5SpecificityThreshold_Simplified.pdf

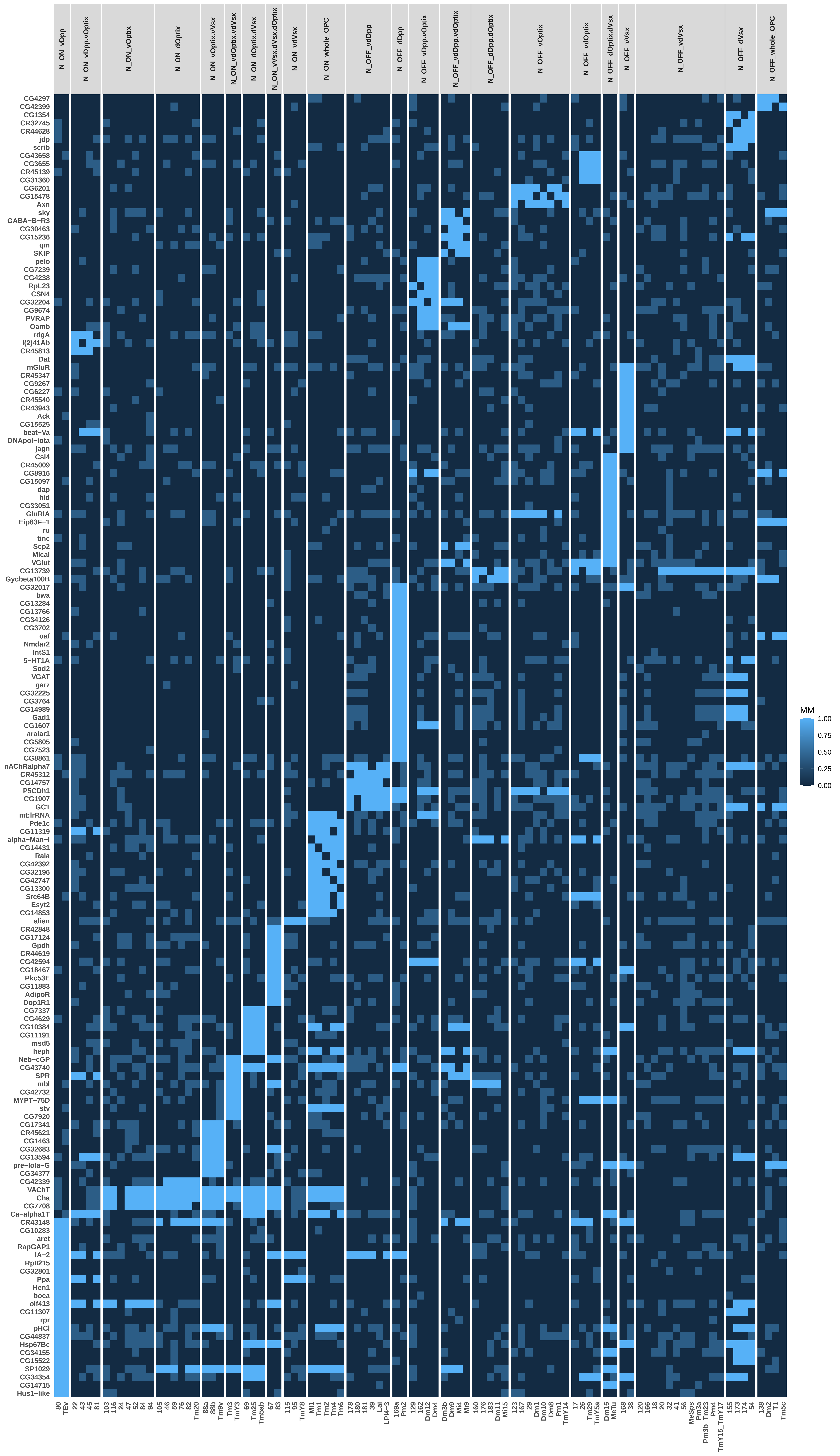

### P70_NO_SO_TFs_0.75ClusterFreq_0.5BinThreshold_0.5SpecificityThreshold_Simplified.pdf

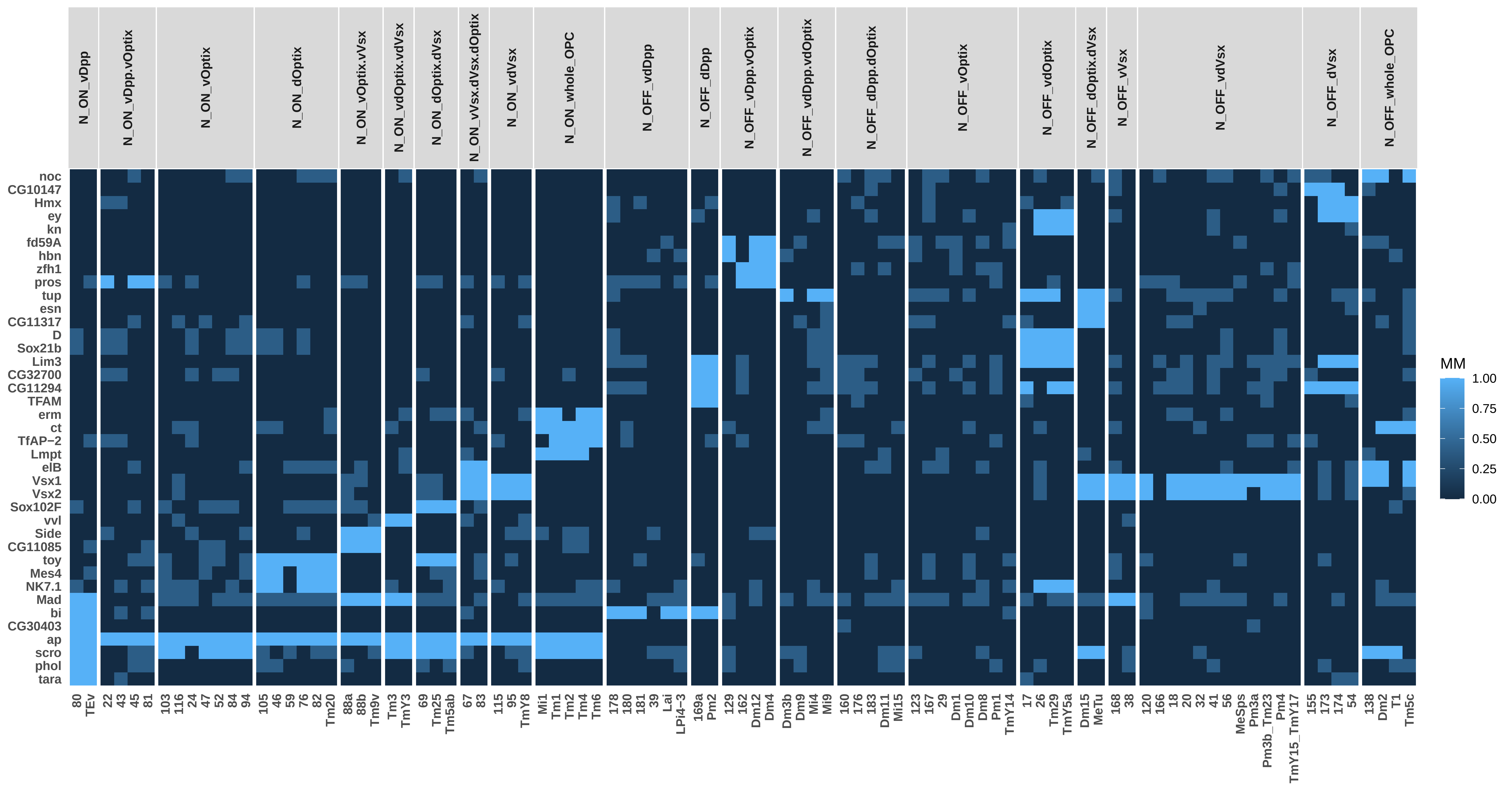

### P70_NO_TFs_0.75ClusterFreq_0.5BinThreshold_0.5SpecificityThreshold_Simplified.pdf

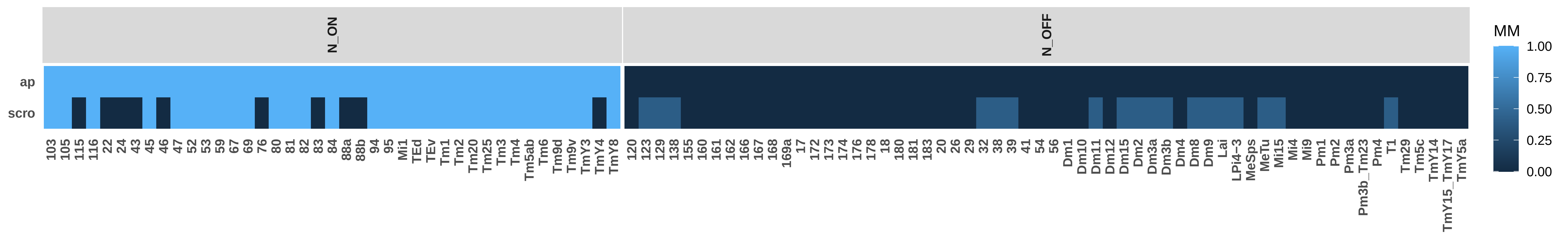

### P70_NO_TO_SO_CSSs_0.75ClusterFreq_0.5BinThreshold_0.5SpecificityThreshold_Simplified.pdf

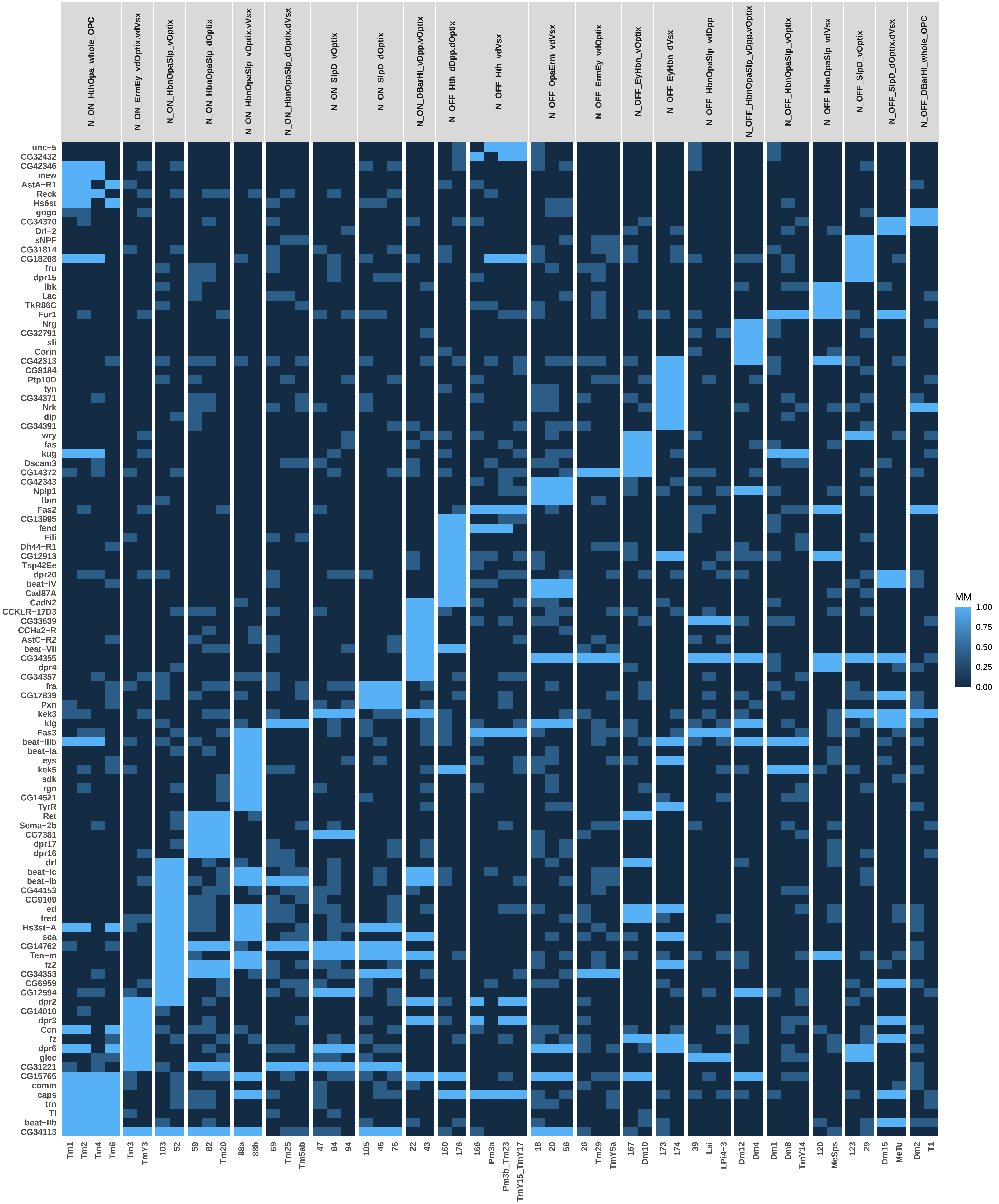

### P70_NO_TO_SO_non_TFs_CSSs_0.75ClusterFreq_0.5BinThreshold_0.5SpecificityThreshold_Simplified.pdf

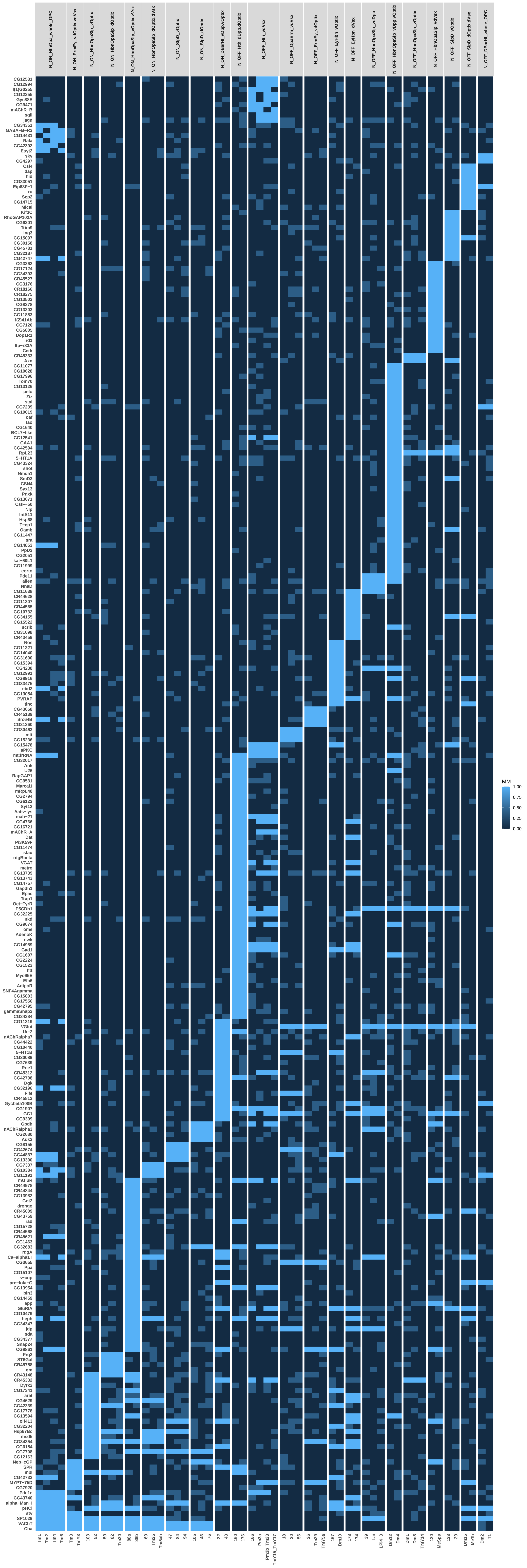

### P70_NO_TO_SO_TFs_0.75ClusterFreq_0.5BinThreshold_0.5SpecificityThreshold_Simplified.pdf

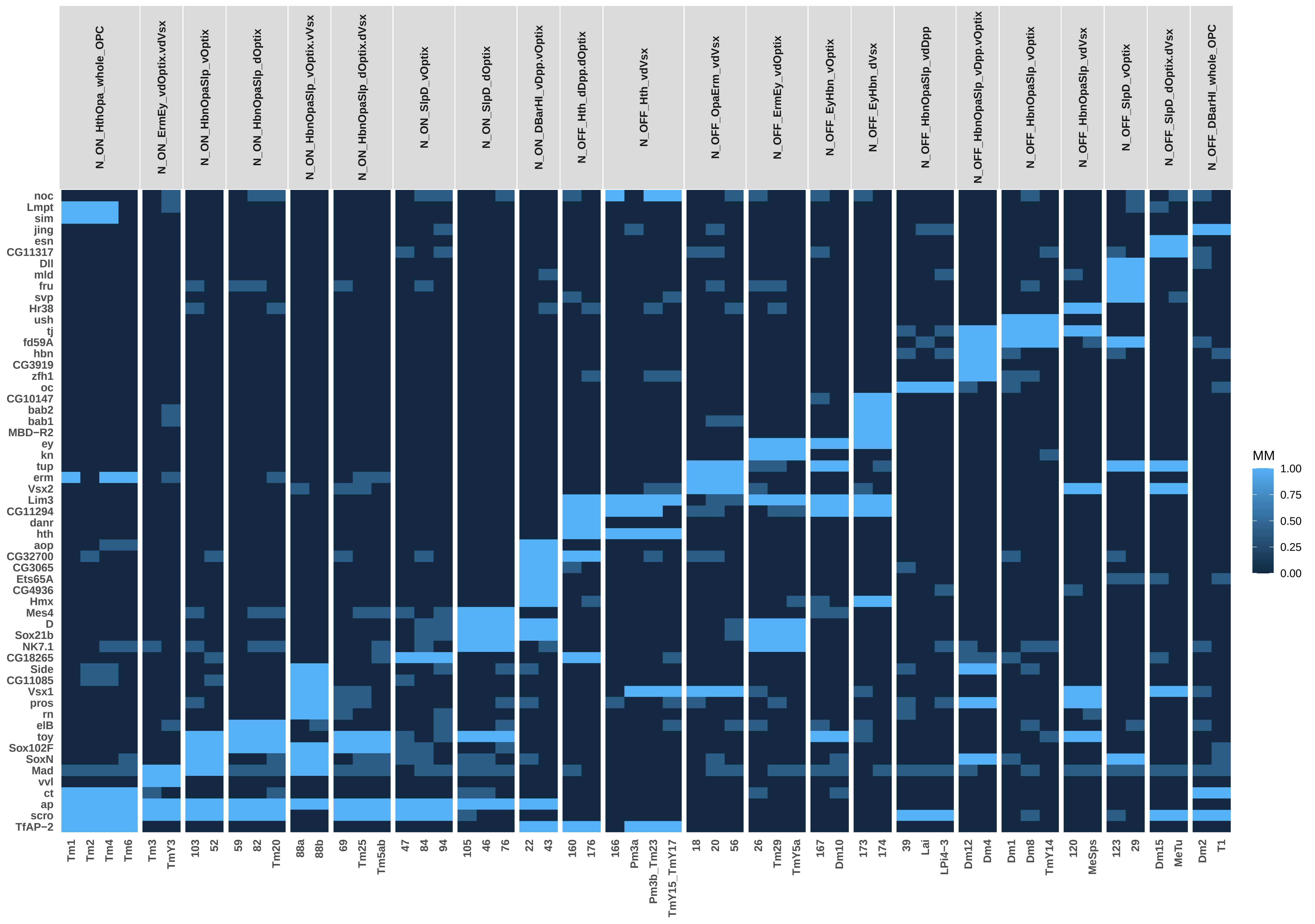

### P70_SO_CSSs_0.75ClusterFreq_0.5BinThreshold_0.5SpecificityThreshold_Simplified.pdf

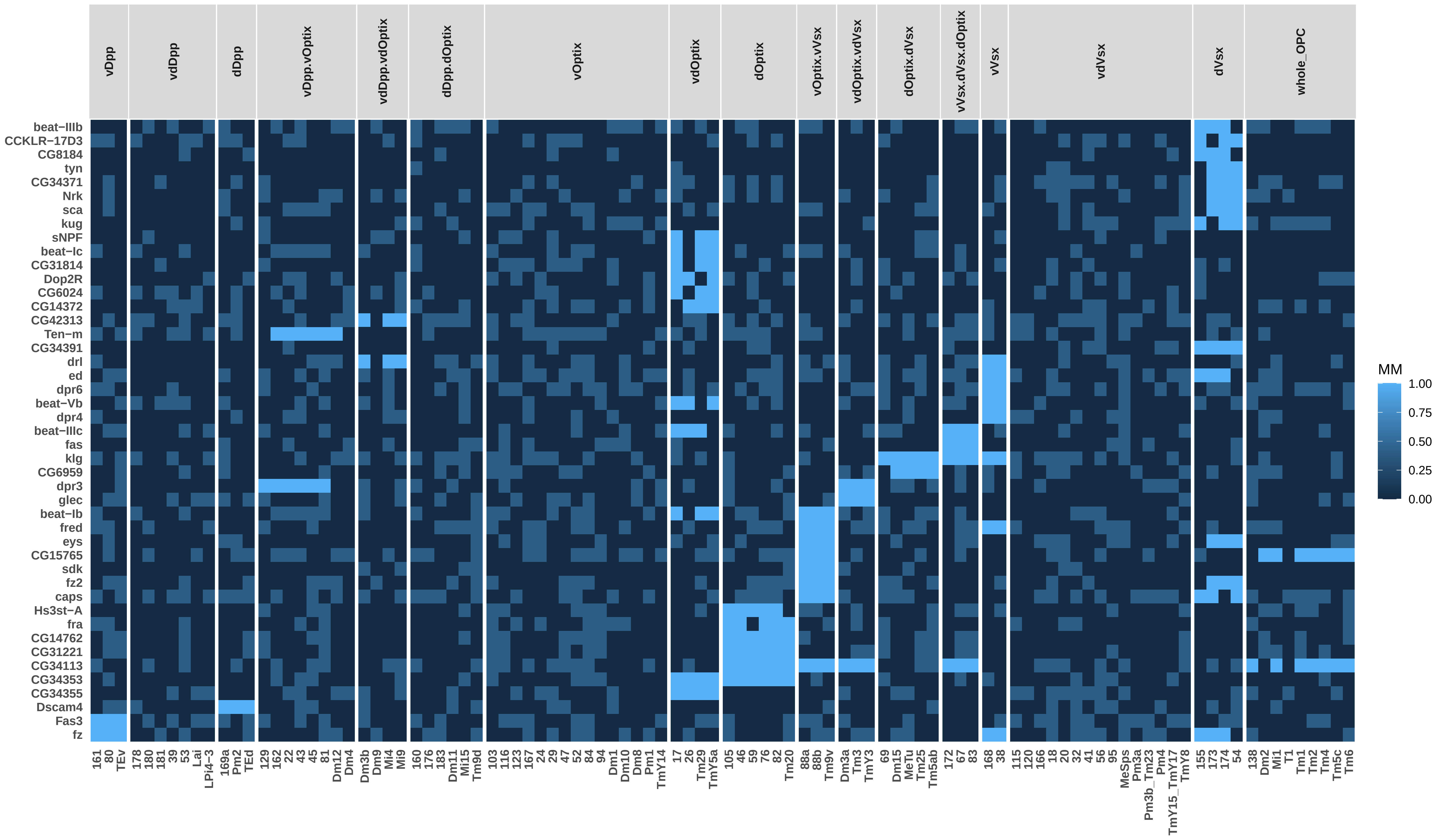

### P70_SO_non_TFs_CSSs_0.75ClusterFreq_0.5BinThreshold_0.5SpecificityThreshold_Simplified.pdf

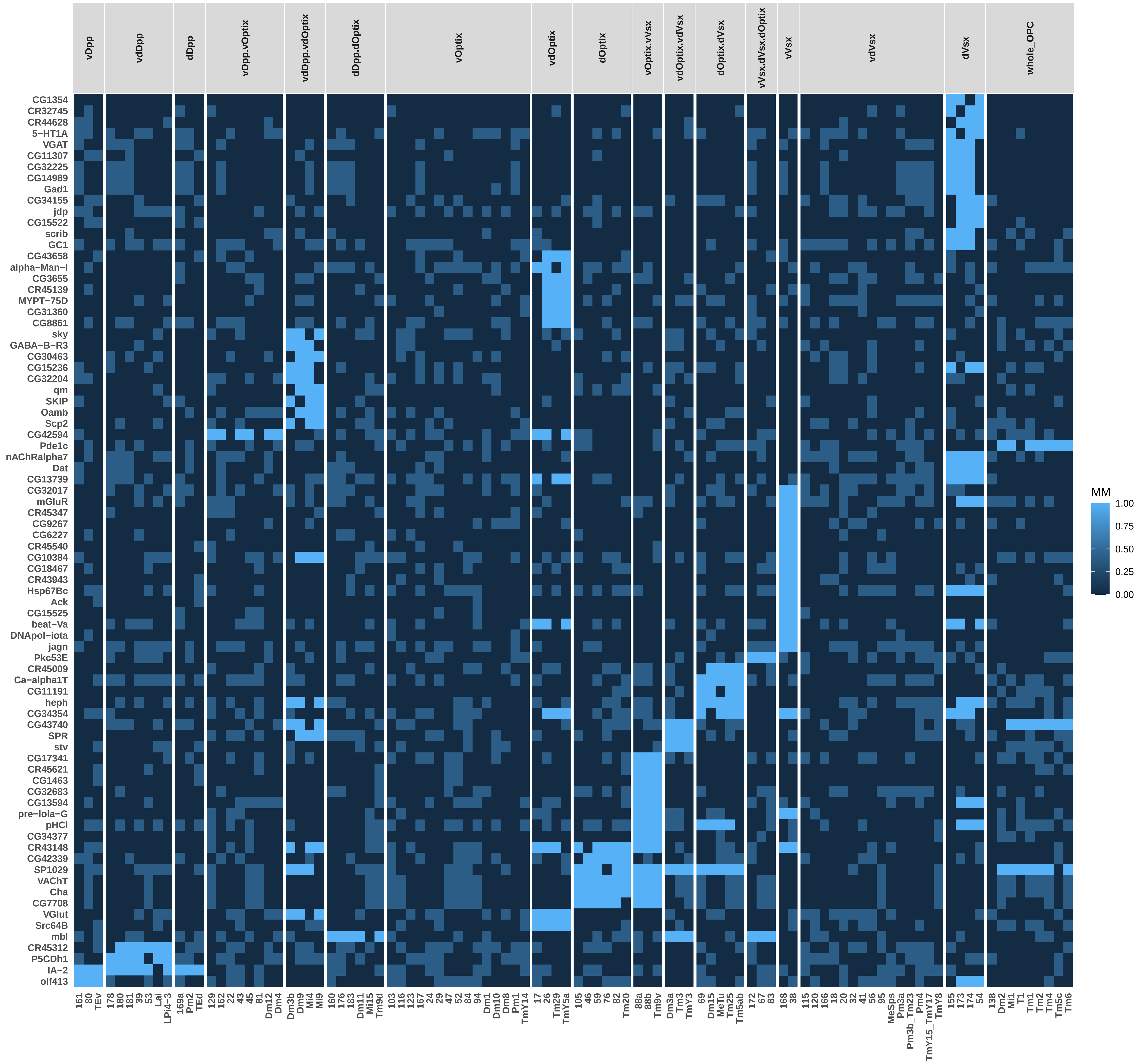

### P70_TO_CSSs_0.75ClusterFreq_0.5BinThreshold_0.5SpecificityThreshold_Simplified.pdf

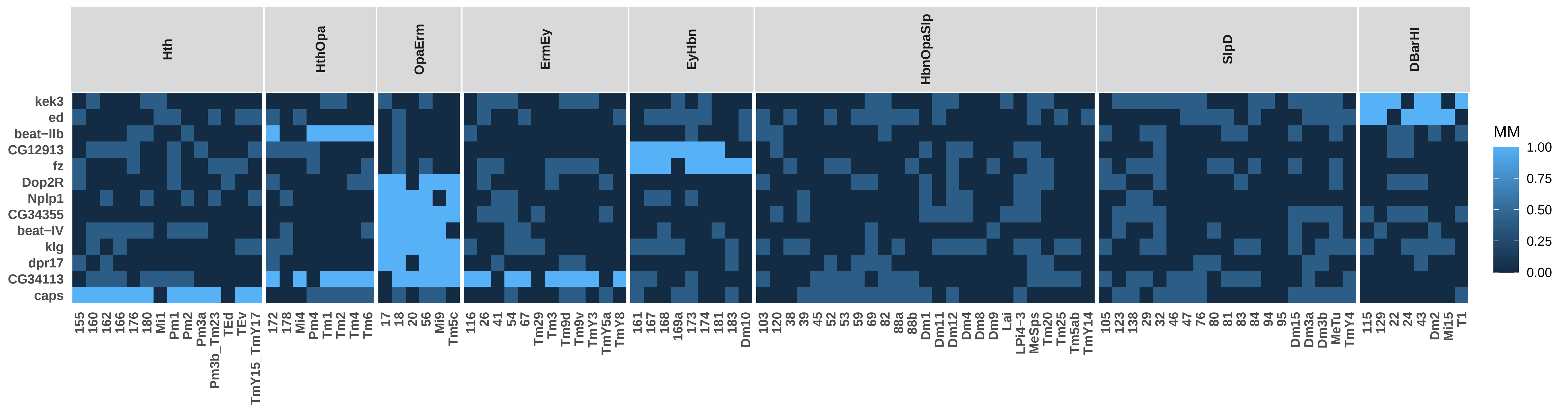

### P70_TO_NO_CSSs_0.75ClusterFreq_0.5BinThreshold_0.5SpecificityThreshold_Simplified.pdf

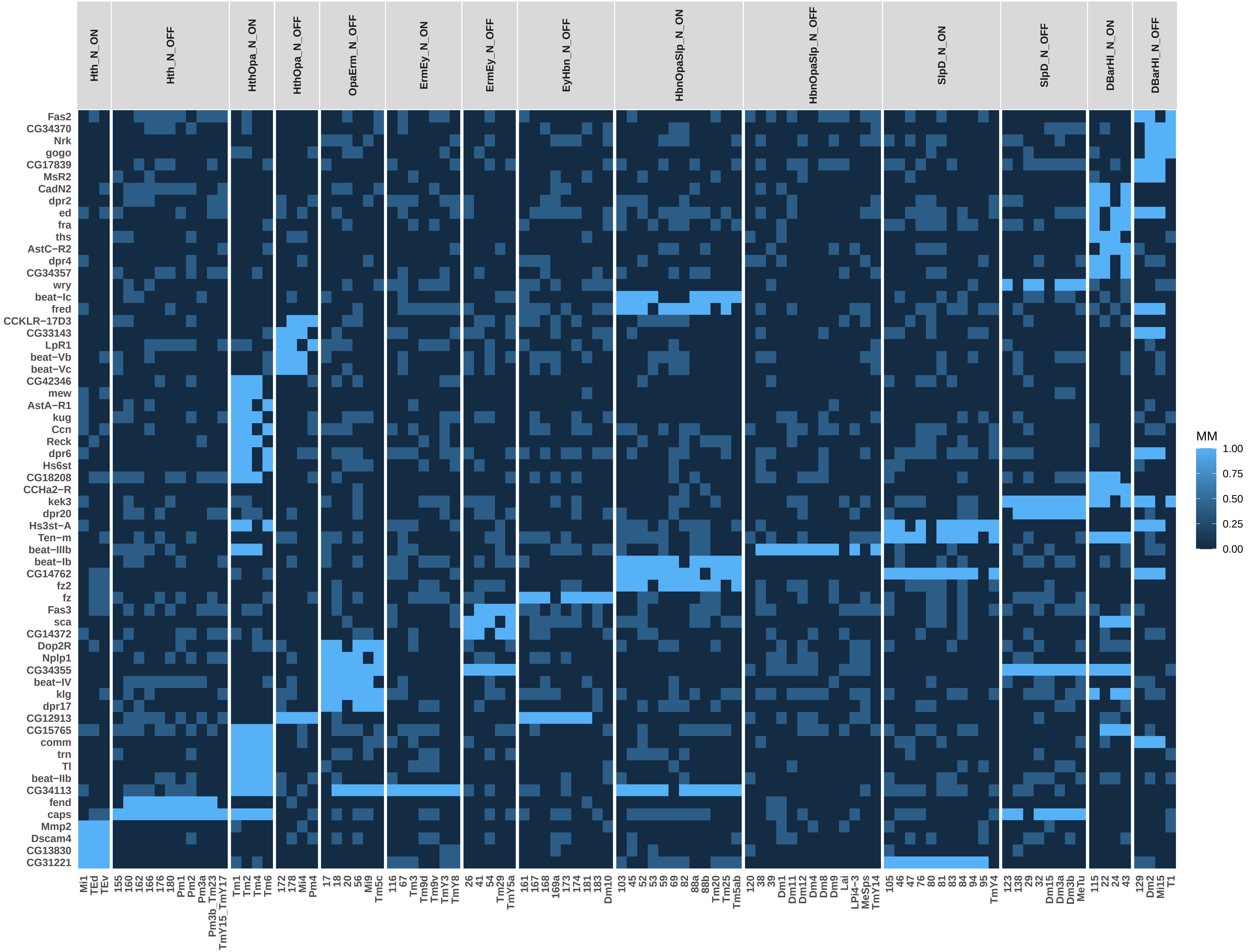

### P70_TO_NO_non_TFs_CSSs_0.75ClusterFreq_0.5BinThreshold_0.5SpecificityThreshold_Simplified.pdf

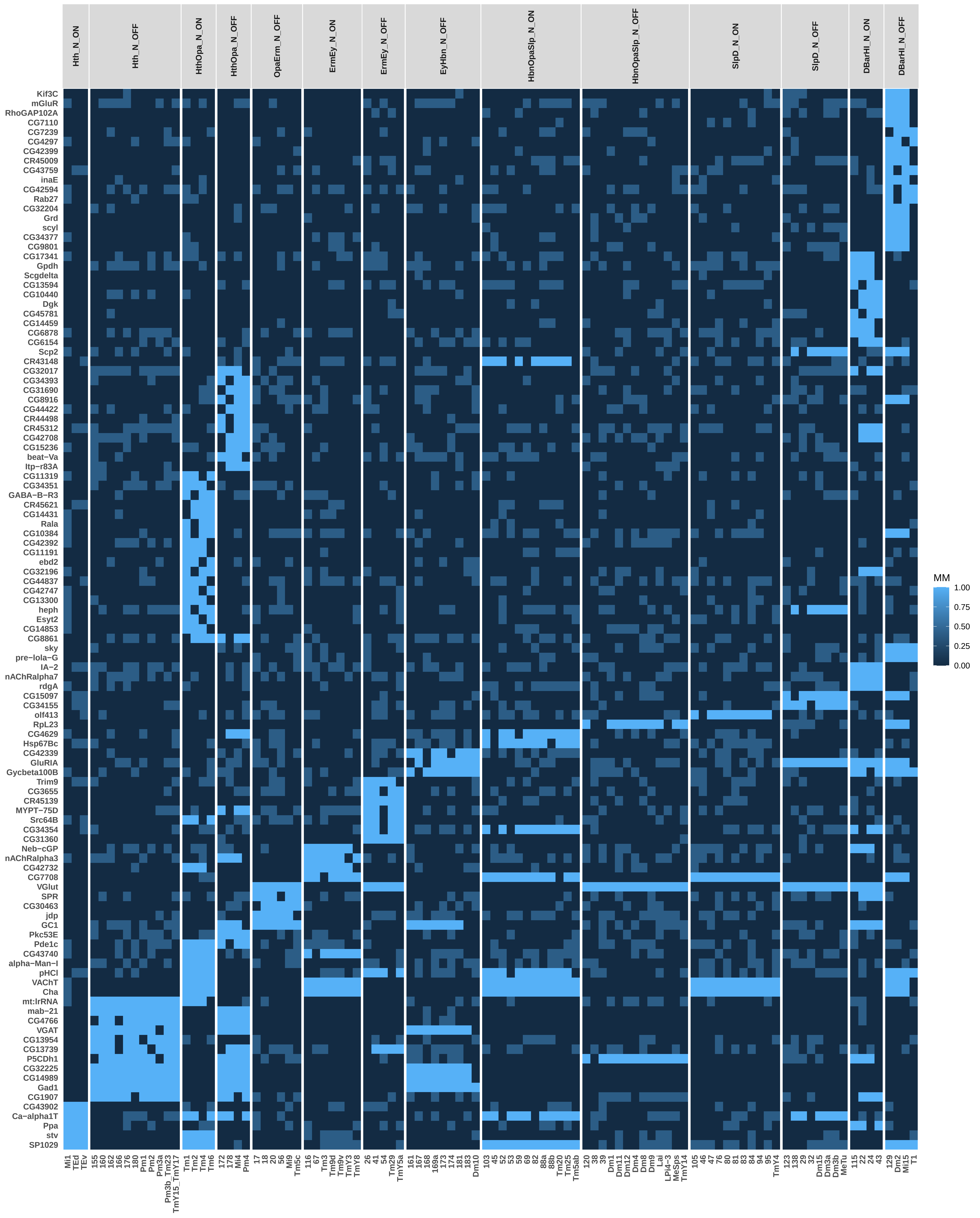

### P70_TO_NO_TFs_0.75ClusterFreq_0.5BinThreshold_0.5SpecificityThreshold_Simplified.pdf

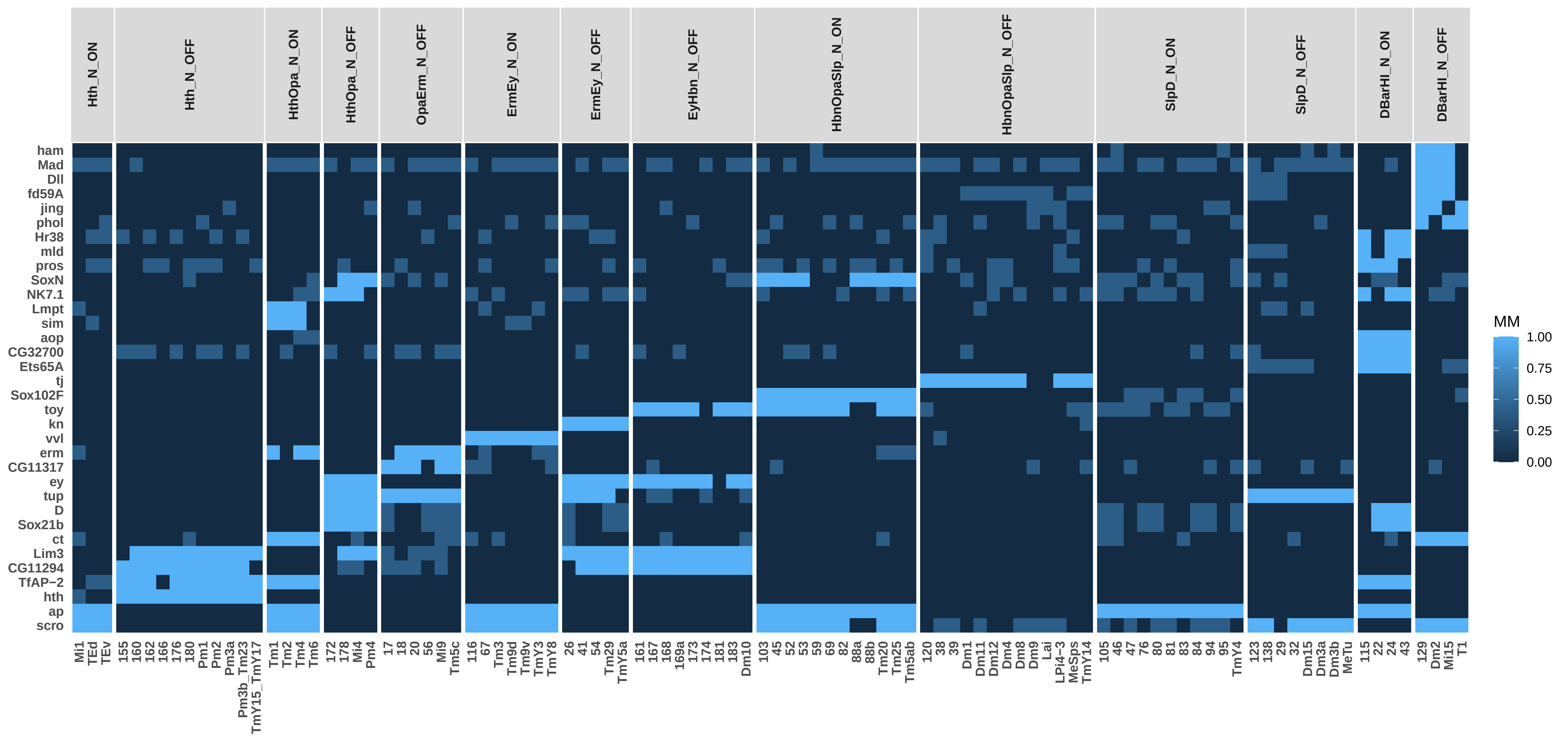

### P70_TO_non_TFs_CSSs_0.75ClusterFreq_0.5BinThreshold_0.5SpecificityThreshold_Simplified.pdf

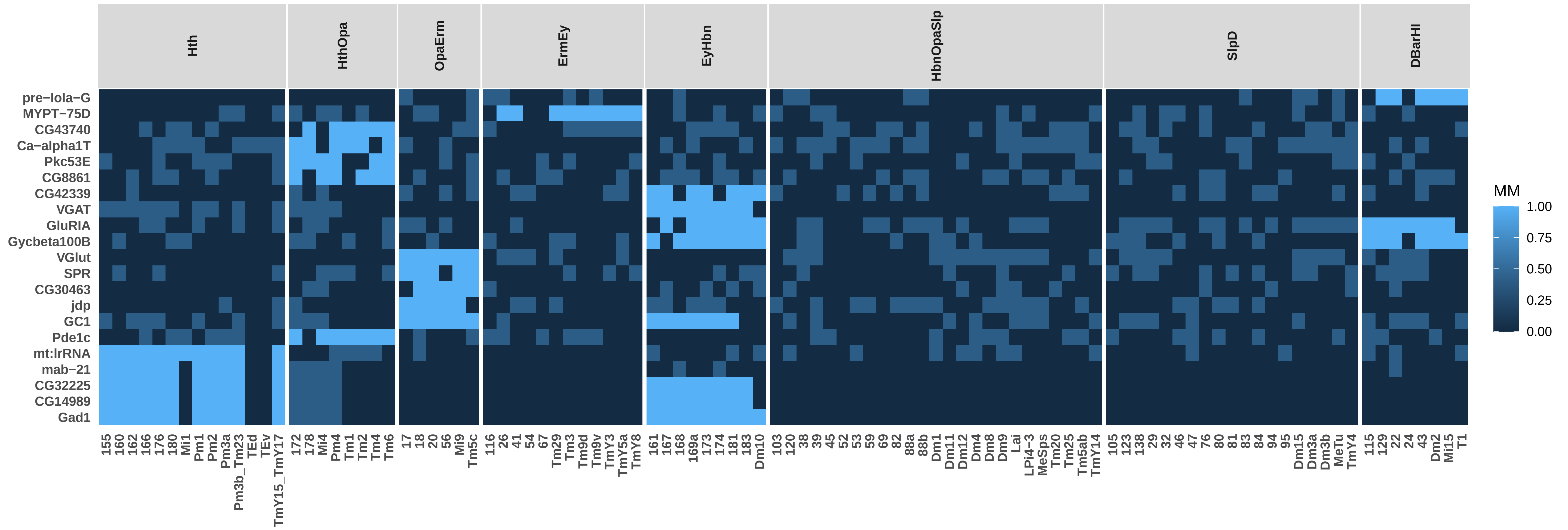

### P70_TO_SO_CSSs_0.75ClusterFreq_0.5BinThreshold_0.5SpecificityThreshold_Simplified.pdf

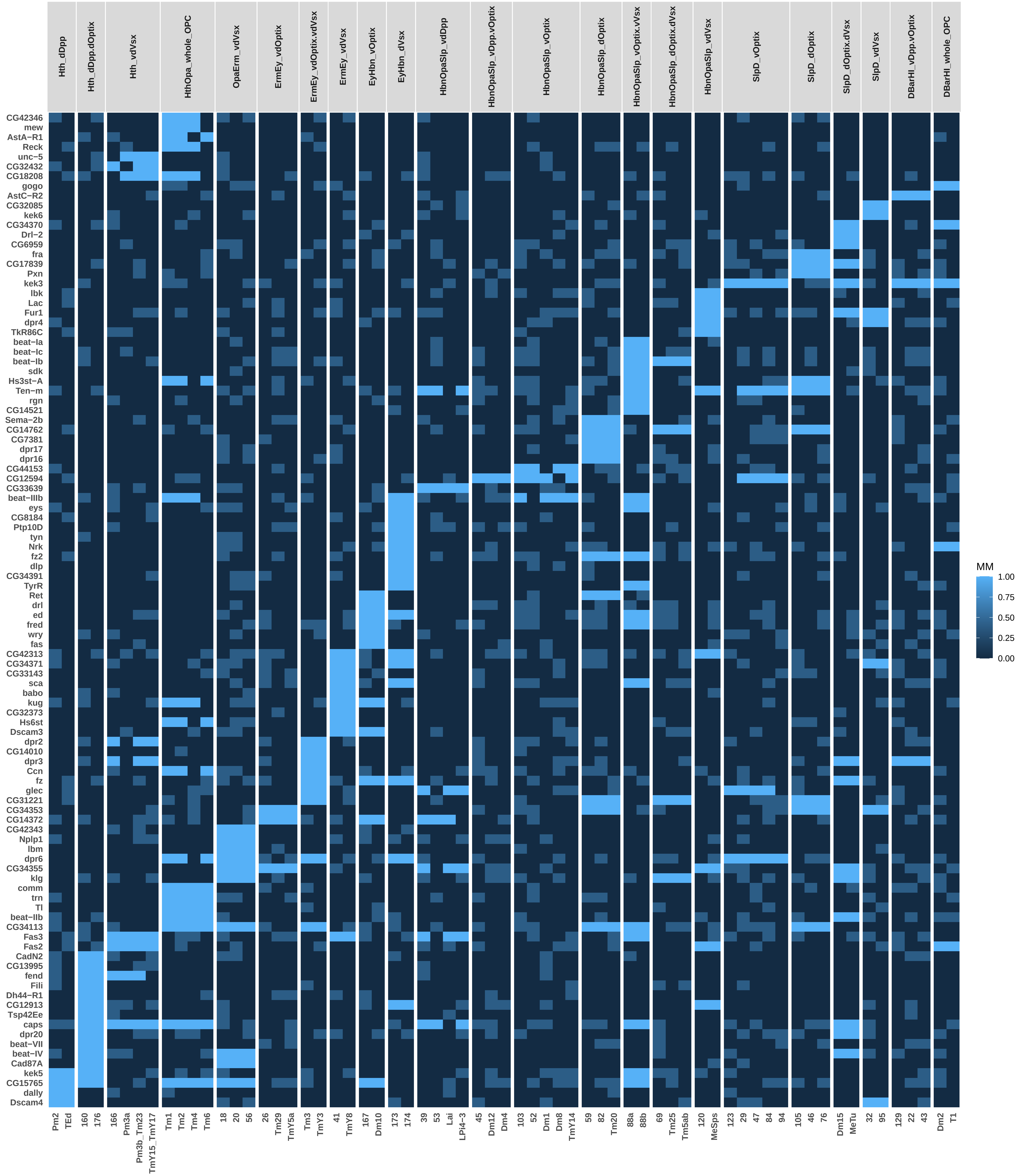

### P70_TO_SO_non_TFs_CSSs_0.75ClusterFreq_0.5BinThreshold_0.5SpecificityThreshold_Simplified.pdf

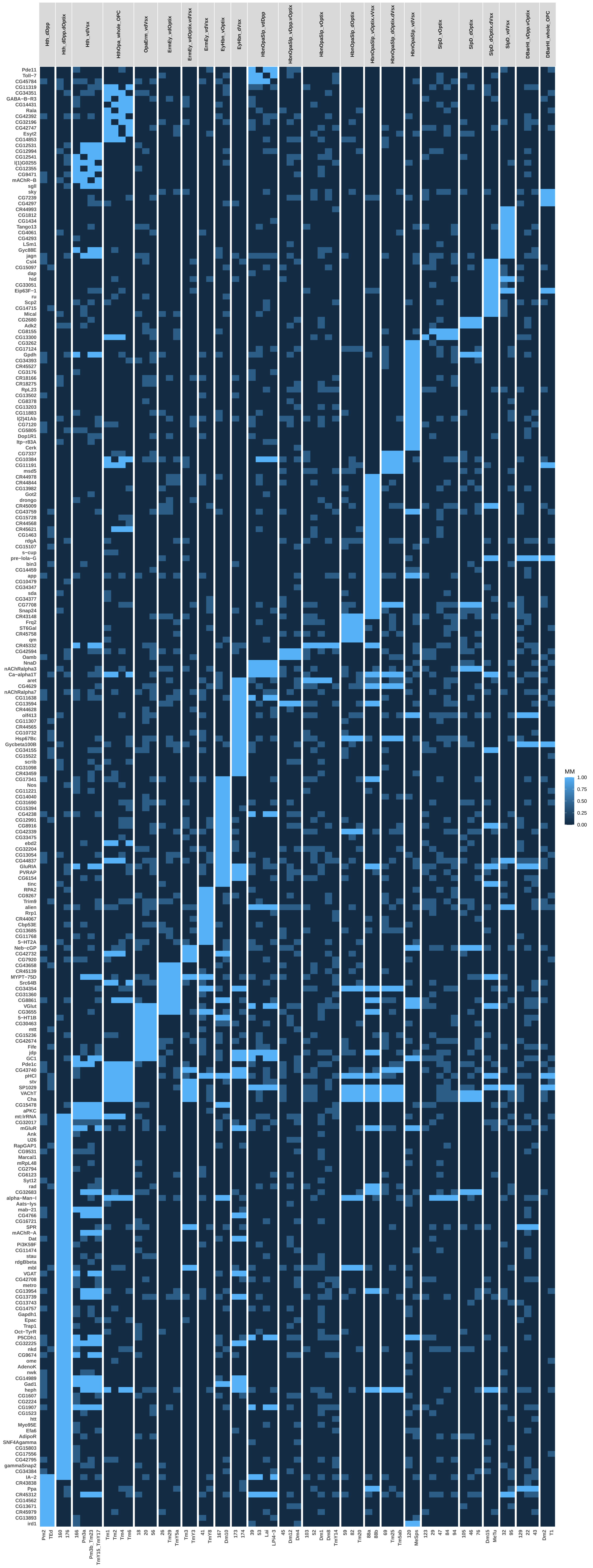

### P70_TO_SO_TFs_0.75ClusterFreq_0.5BinThreshold_0.5SpecificityThreshold_Simplified.pdf

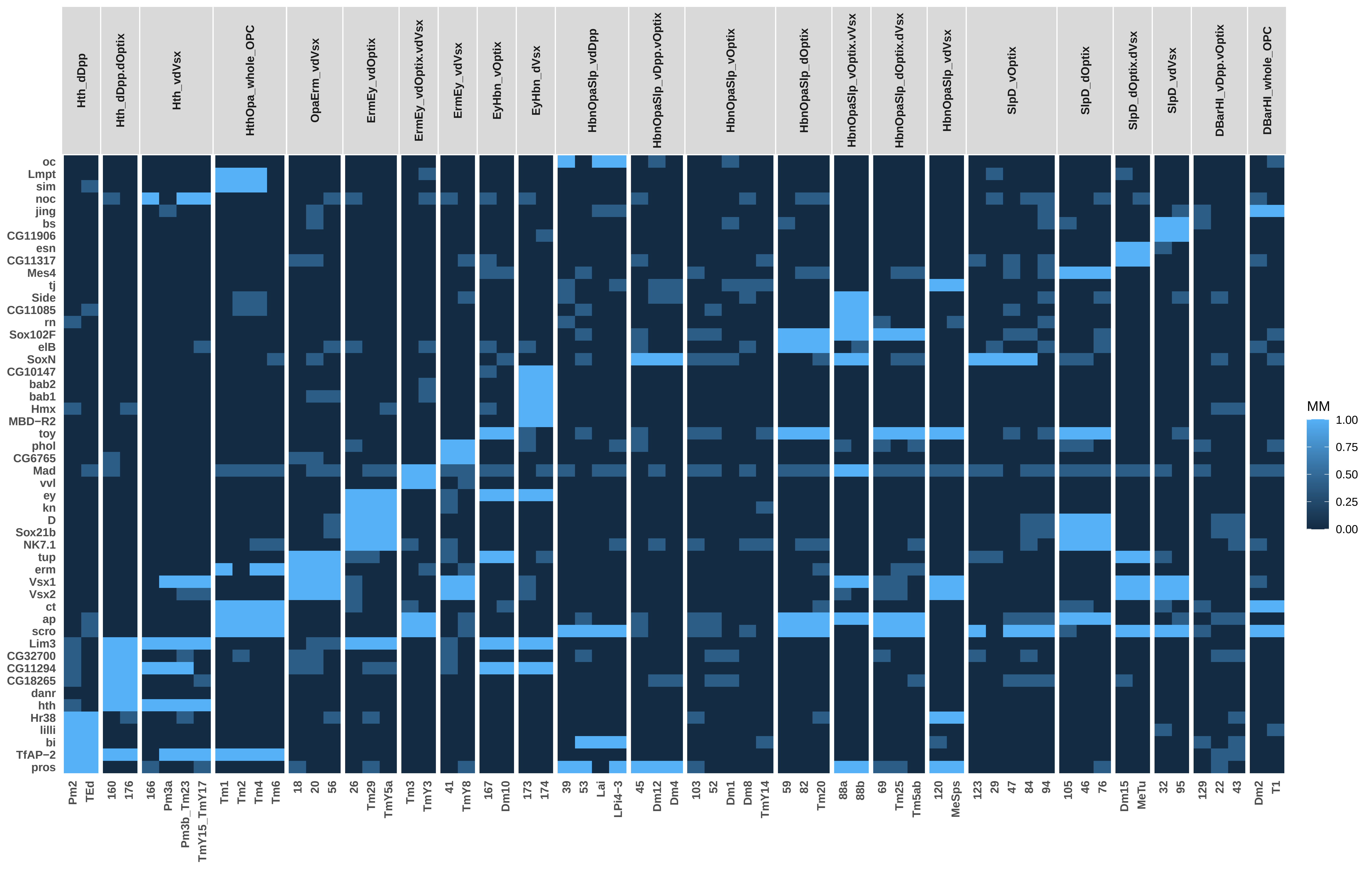

### P70_TO_TFs_0.75ClusterFreq_0.5BinThreshold_0.5SpecificityThreshold_Simplified.pdf

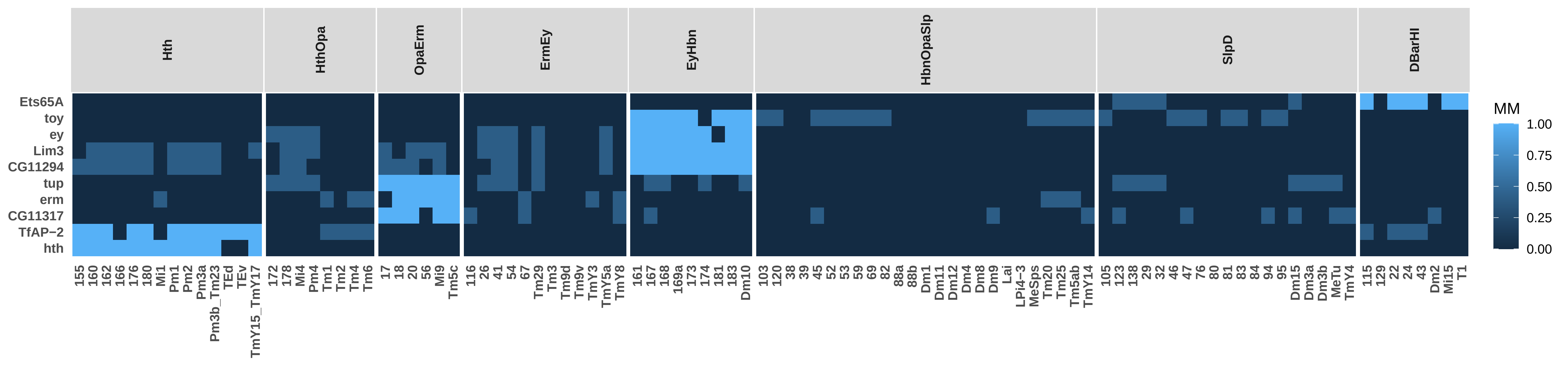
