## Supplementary material for "Establishment of terminal selector combinations in optic lobe neurons": Annex 3: Read_Me.txt

Tables listing the candidate regulators for molecular features (Folders names ending with "GeneExpr", file starting with "RFM_SevClustMarkers") or morphological features (folders names ending with "Morphology", files starting with "Morphology_QualitSevClust").

Please see Methods for explanation on how to read these tables.


The names of the files indicate the parameters used to run the random forest models:

 - P15, P30, etc. = indicates the stage of expression of the predictors (for morphological features) or of predictors and features (for molecular features) 

 - NO, SO, TO = indicates that Notch (NO), Spatial (SO) or Temporal (TO) origin were used as predictors

 - non_TSs = non-TSs TFs were used as predictors

 - TFs = TFs were used as predictors

 - TSs = TSs were used as predictors

 - Regular, Random = whether the expression of the predictors was randomized or not

 - mOPC, All: whether all clusters or only main OPC (mOPC) clusters were used to build and test the random forest models
